# Pervasive selection pressure in wild and domestic pigs

**DOI:** 10.1101/2020.09.09.289439

**Authors:** J. Leno-Colorado, S. Guirao-Rico, M. Pérez-Enciso, S. E. Ramos-Onsins

## Abstract

Animal domestication typically affected numerous polygenic quantitative traits, such as behaviour, development and reproduction. However, uncovering the genetic basis of quantitative trait variation is challenging, since it is probably caused by small allele-frequency changes. To date, only a few causative mutations related to domestication processes have been reported, strengthening the hypothesis that small effect variants have a prominent role. So far, the studies on domestication have been limited to the detection of the global effect of domestication on deleterious mutations and on strong beneficial variants, ignoring the importance of variants with small selective effects. In addition, very often, the study of the effects of selection are conducted on genome sequences that contain a non-negligible fraction of missing data, especially in non-model organisms. Hence, appropriate methods to account for these positions are needed. To overcome these difficulties, here we propose to estimate the proportion of beneficial variants using the asymptotic MacDonald-Kreitman (MK) method based on estimates of variability that summarizes the site frequency spectrum (SFS) while accounting for missing data and use them to perform an Approximate Bayesian Computation (ABC) analysis to infer the Distribution of Fitness Effects (DFE) of each population. We applied this approach to 46 genome sequences of pigs from three different populations, one wild and two domestics, with very different demographic histories and selective pressures. The obtained results showed that domestic and wild pig populations do not differ in nonsynonymous fixed mutations. Therefore, differences in *α* estimation among breeds are determined by their polymorphisms. The comparison of *α* for total and exclusive mutations suggests that the different domestic populations have suffered recent divergent changes in their functional versus neutral polymorphisms ratio, while the wild population is compatible with *α*=0. Besides, the DFE inferred with ABC indicates that both wild and domestic pigs display a large number of deleterious mutations at low frequency and a high number of neutral and/or nearly-neutral mutations that may have a significant effect on the evolution of domestic and wild populations. In addition, models not considering beneficial mutations have higher posterior probabilities, suggesting that beneficial mutations are difficult to detect or are scarce. Indeed, for all three populations, the median proportion of the strong favourable mutations are very low (≤ 0.1%) in those models that includes positive selection, with the average values of weak beneficial mutations around 0.6% for wild boar and 0.8-1.0% for the domestic pigs. Lastly, the analysis based on exclusive mutations showed that recent demographic changes may have severely affected the fitness of populations, especially that of the local Iberian breed.

## INTRODUCTION

Domestic animal histories are evolutionary experiments that have often lasted for millennia resulting in dramatic phenotypic changes to suit human needs. In addition, domestic species can be structured into subpopulations (breeds) that are partly or completely genetically isolated and can display a wide catalogue of specific phenotypes. Therefore, they offer a very valuable material of utmost interest to study the interplay between demography and accelerated adaptation. However, as their demographic history can be quite complex, many events remain unknown or poorly documented nowadays.

The pig (*Sus scrofa*) is a particularly interesting species because of its domestication history and its relatively well-annotated genome. *S. scrofa* originated in Southeast Asia ∼1-4 MYA and spread throughout Eurasia ∼0.2-1.2 MYA, colonizing all climates except the driest (Frantz et al. 2013, Zhang et al. 2021). Subsequently, the pig was domesticated from local wild boars (WB) independently in both Asia and Europe ∼9,000 years ago. To complicate the story, modern European domestic pig breeds were crossed with Asian domestic pigs during the late 17th century and onwards. In breeds such as Large White (LW), approximately 30% of the genome is estimated to be of Asian origin (Bosse, Megens, Madsen, et al. 2014). Nevertheless, some local European breeds, such as the Iberian breed (IB), were spared genetic contact with Asian pigs and no evidence of genetic introgression has been found in this breed (Alves et al. 2003, Esteve-Codina et al. 2013). Moreover, domestic breeds have different recent demographic histories. For instance, the IB breed suffered a dramatic reduction of its effective population size during the last century (Alves et al. 2006), whereas many commercial breeds such as Duroc or LW have been introgressed with Asian pigs (Bosse, Megens, Frantz, et al. 2014).

Differences in the effective population size, demographic histories and artificial selective pressures between pig breed or populations could result in differences among their evolutionary rates. In addition to possible differences in the evolutionary rates between populations, there may be differences in the evolutionary rate between genes within genomes. For instance, it is known that the strength of the selection is affected by the position of the genes in the networks in which they participate. Genes that are more central in a network and are more connected with other genes are more evolutionarily constrained, while peripheral genes are more prone to be under adaptive selection (Fraser et al. 2002; Hahn and Kern 2005; Montanucci et al. 2011; Alvarez-Ponce and Fares 2012). Furthermore, it has been observed that the evolutionary rate, within a metabolic pathway, increases as we move downstream, possibly because upstream genes are more pleiotropic, since they are involved in more functions and hence, these genes are probably more conserved (Rausher, Miller, and Tiffin 1999; Riley, Jin, and Gibson 2003; Livingstone and Anderson 2009; Ramsay, Rieseberg, and Ritland 2009).

So far, the nature of the underlying genetic changes caused by domestication and ensuing artificial breeding is still under debate. While the most prevalent view is that regulatory changes have been targeted (Anderson 2013), several other studies underline the influence of protein coding changes (Rubin et al. 2012). Some authors have reported an increase in the rate of deleterious mutations in domestic pigs compared to their wild counterparts (Cruz, Vilà, and Webster 2008; Renaut and Rieseberg 2015; Pérez-Enciso et al. 2017; Leno-Colorado et al. 2017). Others, as in Makino et al. (2018) detected a general pattern of reduction of variability in domestic populations in relation to their wild counterpart, and a higher nonsynonymous/synonymous ratio across the frequency spectrum. These patterns were compatible with the effect of strong bottlenecks in domestic populations and the higher accumulation of deleterious mutations. Interestingly, the same authors observed the opposite trend in pigs (e.g., higher variability levels in domestic pigs compared to their wild counterpart). Moreover, most of these previous studies have focused on genes of major effect with clear signals of selective sweeps. In those studies, the hallmarks of positive selection were detected as valleys of reduced variation and/or population differentiation that spans relatively large regions (e.g., Amaral et al. 2011, Rubin et al. 2012, Frantz et al. 2013, Wilkinsonet al. 2013), but also by the presence of haplotype structure and homozygosity blocks (e.g., Fang et al. 2011, Bosse et al. 2012, Li et al. 2013). Some of these studies have detected recent breed specific signals of selection attributed to the domestication process (Li et al. 2014, Kim et al. 2015). Nevertheless, the signals were too scarce to explain the whole picture of the domestication process. Other studies have tried to elucidate the effect that domestication has at the genomic scale and on the fitness of individuals of domestic populations (e.g., Cruz et al. 2008, MacEachern et al. 2009, Kono et al. 2016, Perez-Enciso et al. 2016, Makino et al. 2018, Chen et al. 2018, Orlando and Librado 2019). For instance, an excess of deleterious variants has been observed in a number of domestic animal and plants (e.g., contrasting nonsynonymous versus synonymous polymorphism ratios, Chen et al. 2018, using the MacDonald framework, MacEachern et al. 2009, contrasting the ancestors with ancient DNA, Orlando and Librado 2019, combining the frequency of polymorphisms with functional effects and divergence, Kono et al. 2016, Makino et al. 2018). Kono et al. (2016) and Perez-Enciso et al. (2016) found an excess of deleterious variants affecting phenotypes of interest, suggesting, as we previously mention above, that protein sequence may have a stronger influence than regulatory changes in the domestication process. Kono et al. (2016) also showed that null alleles are uncommon in domestic animal species (also reviewed by Anderson 2013), suggesting that phenotypic changes involved in domestication are produced by the accumulation of consecutive mutations that modify the gene functions under selection. Finally, the possible presence of beneficial mutations during the domestication process has also been reported (Perez-Enciso 2016).

Here, we are interested in determining the proportion and the selective effects of protein-coding variants in wild and domestic pig genomes to understand their role in the domestication process. Particularly, we aimed to test the role of both new and extant mutations in the domestication process and whether the phenotypes associated with domestic breeds are the product of a large number of variants with weak selective effects, as suggested by previous results. To achieve this, we have investigated the differential effects of selection on coding sequences at the different molecular scales (gene, metabolic pathway and whole-genome) in two domestic and one wild pig population using the McDonald-Kreitman framework (McDonald and Kreitman 1991, Eyre-Walker 2006, Fay 2011). We also have performed forward exploratory simulations and inferred the distribution of fitness effects (DFE) while taking into account the effect of different demographic scenarios. Interestingly, the analysis was performed using variability estimators that allow including positions with missing data (Ferretti, Raineri, and Ramos-Onsins 2012).

Our results support the hypothesis that changes in allele frequencies in coding variants with weak positive selective effect have been relevant for pig domestication, as evidenced by a relatively high number of nonsynonymous variants segregating at medium and high frequencies and by the obtained estimates of the DFE in domestic pig populations.

## MATERIALS AND METHODS

### Biological samples

We analyzed a sample of 46 pig (*Sus scrofa*) genomes (Table S1). These pigs correspond to European wild boars (WB, n = 20) and domestic pigs, which are represented by the Iberian Guadyerbas (IB, n = 6) and Large White (LW, n = 20) breeds. The two domestic breeds were selected because they have very different interesting features: IB is a local breed that has been under weak artificial selection intensity and with no documented evidence of Asian introgression. LW, in contrast, is a commercial breed undergoing strong artificial selection with a deliberate admixture with Asian pigs (Bosse, Megens, Madsen, et al. 2014; Groenen 2016). To analyze the divergence between the different breeds, we built the consensus ancestral reference sequence obtained from combining the genomic information from several *Sus* species (*S. barbatus, S. cebifrons, S. verrucosus, S. celebensi*, approximately 4.2 MYA of divergence) and the African warthog (*Phacochoerus africanus*, around ∼10 MYA of divergence) and used as an outgroup, as in Bianco et al. (2015). All sequences (see reference numbers at Table S1) are available in public databases: those generated in several previous works (Rubin et al. 2012; Ramírez *et al*. 2014; Bianco *et al*. 2015; Frantz et al. 2015; Moon *et al*. 2015, Esteve-Codina *et al*. 2013, Leno *et al*. 2017) and those generated in this work (WBES0231, WBES0252, WBES0288, WBES0291 and WBES0297, see Table S1). They can be downloaded from the short read archive (SRA, http://www.ncbi.nlm.nih.gov/sra, <u>see accession numbers in Table S1</u>).

### Mapping and genotyping analysis

For each pig genome, raw reads were mapped against the reference genome assembly (Sscrofa10.2, Groenen et al. 2012) using *BWA mem* option (H. Li and Durbin 2009). PCR duplicates were removed using *SAMtools rmdup* v. 0.1.19 (H. Li et al. 2009) and mapped reads were realigned around indels with the *GATK IndelRealigner* tool (McKenna et al. 2010). Genotype calling was performed with *SAMtools mpileup* and *bcftools call* v. 1.3.0 (H. Li et al. 2009) for each individual separately. We set a minimum (5x) and a maximum depth (twice the average sample’s depth plus one) to call a SNP. Base quality was set to 20 (*P*-value=1e-2). Homozygous blocks (regions of contiguous positions with the same nucleotide as the reference genome) were also called, following the same criteria as with the SNPs (i.e., minimum and maximum coverage and base quality) and using *samtools depth* utility, *BEDtools* (Quinlan 2014) and custom scripts (available at https://github.com/miguelperezenciso/NGSpipeline, Pérez-Enciso *et al*., 2017). This resulted in a *gVCF* file per individual with the information about variant calls and non-varying positions. Next, each *gVCF* file was converted into a fasta file and all fasta files were subsequently merged to obtain a multindividual *gVCF* file (Pérez-Enciso et al. 2017).

### Analysis of the population structure of the samples

A principal component analysis (PCA) was performed using the total number of SNPs to explore the population structure. First, genotypes were converted to alternative allele frequency, being 0 for the homozygous reference genotype (0/0), 0.5 for the heterozygous genotype (0/1) and 1 for the homozygous alternative genotype (1/1). For cases of missing genotype (./.), these were replaced by the average SNP frequency across all individuals. We used the function *tcrossprod()* from R v. 3.3.1 (2016) to obtain the matrix of covariates from the matrix of frequencies. Finally, we obtained the principal components from the Eigen-value decomposition with the R function *eigen()*. The software ADMIXTURE (Alexander et al. 2009) was also applied to analyze the population structure. The more suitable K was estimated using cross-validation procedure (Alexander et al. 2009) and the Evanno’s method (Evanno et al. 2005).

### Estimation of codon bias

We have used the single reference sequence of *S. scrofa* to estimate the codon bias at gene and genome level, assuming that polymorphic variants are not going to strongly modify the proportion of codons at this species. We have estimated gene and genomic codon usage with the Major Codon Usage statistic (MCU, the frequency of major codons among all codons in a sequence) and the Effective number of codons (*N*_*cw*_), using the python script following Fuglsang (2006). High values of MCU indicate a strong bias in codon usage, while low values suggest small bias. Instead, high values of *N*_*cw*_ indicate low codon bias because many codons are used. We have also calculated the correlation between MCU and *α* estimates (see below), considering all annotated coding regions and only coding regions having *α* values larger than zero.

### Estimation of levels and patterns of variability

Genetic diversity and divergence per pig population were estimated using *mstatspop* software (Nevado, Ramos-Onsins, and Perez-Enciso 2014; Bianco et al. 2015; Guirao-Rico et al. 2018, available from the authors, https://github.com/cragenomica/mstatspop). The *multi-VCF* file was converted into a *tfasta* (transposed *fasta*) file and *mstatspop* was run on i) the whole genome, ii) windows of 5-Mb size, and iii) gene coding regions. Note that, given the ubiquitous presence of missing data, the SFS with highest sample size that contained more coding SNPs, retained only around 20-25% of total coding variants. As the aim of this work is to detect the presence of (weak) beneficial selective effects and as not to lose power or bias the results of the analysis, we preferred to use an alternative method that consider the whole set of SNPs. We used four different estimators of nucleotide variability that takes into account missing data (Ferretti, Raineri, and Ramos-Onsins 2012): Watterson (Watterson 1975), Tajima (Tajima 1983), Fu&Li (Fu and Li 1993) and Fay&Wu’s estimators (Fay and Wu 2000). The variability was estimated for total, shared and exclusive variants, being shared and exclusive nucleotide variability counted regarding to the total positions (i.e., total = shared + exclusive).

### Filtering for artefactual effects

A preliminary analysis of the variability showed a moderate negative correlation (∼ 0.3) between the levels of variability and divergence and the proportion of missing data for each gene. To eliminate this artifactual correlation, we plotted the estimators of variability and divergence versus the ratio of missing data and eliminated those genes that showed a ratio of missing data greater than 0.3. Since this filtering was not enough to completely remove the bias, we also removed genes with extreme values of variability and divergence (higher than 99% quantile of the total genes). The remaining ∼13,500 genes (70% of the total annotated genes) showed a low or null correlation with missing data and were used in the present analysis (Table S2).

### Estimation of the proportion of adaptive of variants

Under the neutral model, the majority of polymorphisms segregating in a population are neutral and only a small number of selected variants segregates for a short time on their way to loss or fixation. Hence, most of the positive selected variants are only observed as fixed variants. In addition, functional positions (nonsynonymous positions) are constrained compared to non-functional positions (synonymous positions), and hence, their evolutionary ratios are smaller. In the neutral scenario, polymorphism and divergence (excluding the adaptive fixed variants) are proportional to the mutation rate and to the constriction factor in the case of nonsynonymous positions (McDonald and Kreitman 1991, Eyre-Walker 2006, Fay 2011). That is:

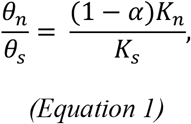

where *θ*_n_ is the nonsynonymous variability, *θ*_s_ is the synonymous variability, *K*_n_ is the nonsynonymous divergence, *K*_s_ is the synonymous divergence and *α* is the proportion of adaptive variants that have been fixed. To estimate the proportion of nonsynonymous substitutions that are adaptive (*α*) the previous expression is reordered (e. g., Eyre-Walker 2006):

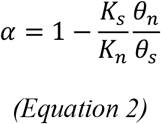

A higher ratio of nonsynonymous to synonymous divergence versus polymorphisms suggests that positive selection has fixed adaptive variants (*α* > 0) and the opposite case (*α* < 0) suggests the presence of deleterious mutations segregating in the population.

If we consider that weak deleterious mutations are segregating in the population, we expect that their relative proportion will be higher at lower frequency variants and low or zero for fixed deleterious mutations. Following the same notation as in equation 2:

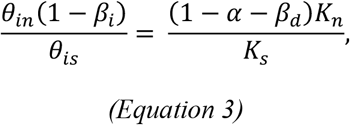

where *i* refers to the frequency at which the calculation of variability is estimated, *β*_*i*_ is the proportion of weakly deleterious polymorphic mutations at frequency *i, β*_*d*_ is the proportion of weakly deleterious fixed mutations. *β*_*d*_ < *β*_*i*_ was assumed at any frequency. Then, solving for the proportion of fixed adaptive variants (*α*):

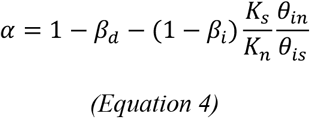

We see that in case of calculating *α* without considering the effects of deleterious mutations, this would be underestimated depending on the frequency at which the estimates of variability are calculated. If we assume that the deleterious variants would hardly be fixed, a good estimator of *α* using equation 2 would be the one that estimates variability based on high frequencies, as it would hardly contain deleterious mutations. This is in agreement with the asymptotic arguments used in Messer and Petrov (2013) and implemented in Haller and Messer (2017).

Similarly, if we also consider that weak positively selected variants are segregating in the population, we expect that their relative proportion, compared to neutral ones, is higher at higher frequencies:

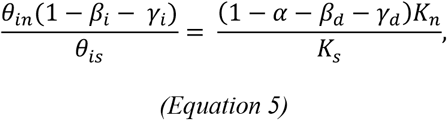

where *γ*_*i*_ is the proportion of weakly advantageous polymorphic mutations at frequency *i*, and *γ*_*d*_ is the proportion of weakly advantageous fixed mutations. Again, solving for the proportion of fixed adaptive variants (*α*+*γ*_*d*_):

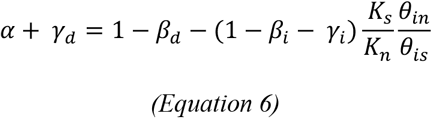

In this case, the presence of adaptive variants segregating in the population would affect the estimates of variability based on high frequency variants when using equation 2, which would result in an underestimation of the proportion of fixed adaptive variants (*α*). Note that adaptive variants stabilized at intermediate frequencies, which can be an important source of adaptation considering the infinitesimal model, are not considered in this approach.

If we focus on the effects of polymorphic weakly selected mutations, equation 5 suggests that the ratio of nonsynonymous to synonymous polymorphisms would increase due to mutations having both positive and negative effects. It is expected that the number of mutations with negative selection coefficients would rapidly decrease as we move to intermediate and high frequencies, while the opposite trend is expected for mutations with positive selection coefficients. Hence, higher ratios of nonsynonymous to synonymous polymorphisms at higher frequencies may be explained by the presence of advantageous mutations segregating in the population.

Furthermore, in cases where two populations are from the same species and there are no fixed mutations between them (e.g., they have equal divergence ratios versus the outgroup), we can estimate the possible differential effect of the selection (positive and negative together) at any frequency between populations from the ratios of synonymous to nonsynonymous polymorphisms of the two populations (*R*_*β γ*_*i*_):

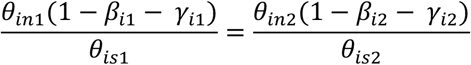

and

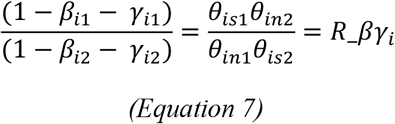

In addition, a comparison of the *R*_*βγ*_*i*_ values calculated using different variability estimators (hereafter *R*_*βγ*_*i*_ pattern) can be used to inform about the effects of selection. For example, values over 1 indicate that the population 2 has a higher ratio of nonsynonymous to synomymous polymorphisms compared to population 1, either produced by an accumulation of deleterious or of beneficial polymorphisms. Importantly, different demographic effects (e.g., bottlenecks) together with the presence of mutations with small selective effects may also disturb the ratios of variability and hence must be considered when interpreting the results. We include a couple of possible scenarios that can account for possible patterns: (i) after split of two the two populations, both populations have the same population size, but population 1 is affected by the action of positive selection on a quantitative trait (polygenic effect), which causes an increase in the frequencies of some of its variants without getting fixed. Under this scenario, we expect a R_βγ > 1 when this is calculated based on high frequencies. (ii) after split of two the two populations, the population 2 remains equal population size as before the split and the population 1 suffers a reduction in its effective population size, which causes that the slightly deleterious mutations become effectively neutral. Then, R_βγ is expected to be > 1 when it is calculated based on low frequency variants.

The effect of linkage disequilibrium between selective (deleterious or adaptive) and neutral variants should not overly affect the expected estimate of the proportion of adaptive variants, as it would affect both synonymous and nonsynonymous positions in similar proportion. On the contrary, the interaction of variants with opposite selective effects would possibly reduce the effect of selection and would have a significant consequence on the estimation of adaptive fixed variants (Hill and Robertson 1966; Booker and Keightley 2018).

### Bootstrap analysis

Nonparametric bootstrap analysis was performed to estimate the null distribution of the *α* statistic for each variability estimator and pig population. In each case, synonymous and nonsynonymous coding positions were randomly chosen with replacement and the *α* statistic was calculated as in equation 1. This process was repeated 100 times.

### Simulations

We carried out forward simulations using the software *SLiM* (Haller and Messer 2017) in order to explore the interaction between the different selective effects and demographic factors affecting the evolution of pig populations during domestication. We explored the expected values of nucleotide diversity, divergence, *α* and *R_βγ* statistics under 63 different scenarios. For each scenario, we simulated three populations corresponding to wild, domestic and an outgroup species. We first simulated nine different scenarios that were classified into three main groups: i) standard neutral model (SNM); ii) a model with negative selection (NS) and iii) a model with positive selection (PS). For the models with selection, we let that selection operate from the ancestral species to the present time. Each group of scenarios (SNM, NS and PS) was simulated with a constant effective population size for the three populations or with a reduction or an expansion of the effective population size in the branch leading to domestic pigs. A second group of simulations was performed under more complex scenarios. In those simulations, we incorporated the combined effect of negative and positive selective effects (using gamma and exponential distributions for the selective coefficients, respectively) plus demographic effects such as expansion and reduction of the effective population size in the domestic simulated populations and with or without migration from the wild into the domestic populations (in total 54 complex simulated scenarios). Figure S1 shows a general scheme for the simulated populations and Tables S3A-B show the parameter values used in these simulations. The obtained results were analyzed using *mstatspop* software (see above).

### Approximate Bayesian computation (ABC) analysis

We used the ratio of the estimates of nucleotide variability (*θn*/*θs*) per nucleotide for nonsynonymous versus synonymous positions (Fu&Li, Watterson, Tajima and Fay&Wu) and of divergence (*Kn/Ks*) as statistics to infer the distribution of fitness effects (DFE) in coding regions. We compared four evolutionary models that differ in the shape of the DFE using the algorithm proposed by Tataru et al. (2017), which are the following: (i) model A: a model with a deleterious gamma DFE with the mean and the shape of the gamma distribution as model parameters, (ii) model C: a model with a gamma distribution of deleterious variants with two parameters (shape and mean) and an exponential distribution of beneficial variants with one parameter (mean), and the additional parameter of the proportion of beneficial versus deleterious variants, (iii) model DN: a model with a discrete distribution of a priori values of negative selective coefficients with the proportion of negatively selected mutations for each of the negative selective coefficients as parameters. and (iii) model D: a model with a discrete distribution of a priori values of possible selective coefficients (positive and negative) with the proportion of positively and negatively selected mutations for each selected coefficients as parameters. Some of the additional parameters, such as demographic or linkage effects, were considered as nuisances. Nuisance parameters mimic the demographic effects and other parameters such as linkage effect by using the difference between the observations for the neutral dataset (i.e., synonymous sites) and the expected under the neutral model. Others, such as errors in the polarity of unfolded mutations, were fixed.

Table S4 shows the parameters of each model and the prior distributions used in the analysis. We used *polyDFEv2* (Tataru et al 2019) to obtain the expected unfolded site-frequency spectrum (SFS). The code of *polyDFEv2* was slightly modified in order to print the SFS and the parameters for a large number of conditions, which are needed to perform the ABC analysis using summary statistics. For each model, one million iterations were run using different parameter conditions and the resulting SFS for each condition were kept to later calculate the ratios of variability, divergence and the *α* statistic. The ABC analysis was performed using the R library *abc* (Csillery et al. 2012). We performed a cross validation analysis to evaluate the ability of the approach to distinguish between models using the *cv4postpr()* function, as suggested in the *abc* library documentation. The confusion matrix indicated that these three models were quite distinguishable with a probability of true classification from model A versus C/DN/D of 0.69, from model C versus A/DN/D of 0.65, from model DN versus A/C/D of 0.83 and from model D versus A/C/DN of 0.80, using a tolerance value of 0.05 (Table S5). Posterior probabilities of each model given the observed data (i.e., the probability assigned to each model relative to the other models of the analysis), were obtained using the *postpr()* function and considering a multinomial logistic and a rejection approach. Additionally, a goodness of fit analysis, which evaluates whether the prior distribution for model parameters are realistic, was also performed. The best model was selected based on posterior probabilities. Once the best model was chosen, the ability to infer the parameters of the model was assessed using the *cv4abc()* function. Prediction errors for the parameter inference of each model are shown in Table S6 and Figure S2. The parameters of the best model were inferred with the *abc()* function using a local linear regression and a rejection approach. Posterior predictive simulations were performed using the *α* statistic to determine whether the simulated data generated from the estimated parameter of our best model resembled the observed data (1000 replicates). Finally, the *α* values can be simply estimated using equation 10 from Tataru et al. (2017), as the proportion of positive selective coefficients (*s*) values in the case of the discrete distribution.

### Gene context and network topology analysis

We downloaded the complete list of pathways and genes of *S. scrofa* from KEGG v.20170213 (http://www.genome.jp/kegg/, Kanehisa et al. 2008). The list contained 471 pathways and 5,480 genes. The median and mean number of genes per pathway was 26 and 43, respectively, and ranged from 1 to 949. We filtered the pathways according to their size, removing pathways with less than 10 and more than 150 genes in order to discard pathways that were not informative or too generic and complex. The final list contained 171 pathways and 3,449 genes.

To analyze the selection pressure of each gene according to its position in the pathway, we obtained different topological parameters. For that, we first downloaded the XML file of each pathway from KEGG v.20170213. These files were analyzed with the *iGraph* R package (Csardi G. and Nepusz T. 2006) to obtain the topological descriptors of each gene in each pathway. For each gene, three different measures were computed: *betweenness* (number of shortest paths going through a vertex), *in-degree* (number of in-going edges) and *out-degree* (number of out-going edges). These parameters are measures of the importance of a gene within a pathway: *betweenness* is a centrality feature, *in-degree* suggests the facility of a protein to be regulated and *out-degree* reflects the regulatory role of a protein. We tested whether negatively and positively selected genes differed in any of these statistics using a nonparametric Wilcoxon rank test, due to the extreme leptokurtic distributions involved.

### Genomic context patterns

We have additionally tested whether there is a significant correlation between *α* and recombination, gene density, missing rate, %GC and CpG islands across genomes.

### Testing the differences in the estimates of *α* using whole-genome data versus the mean of gene estimates

We have studied the behaviour of the *α* statistic when it is estimated considering a single large dataset (*i.e*., genome) or when it is estimated using the mean of many subsets (*i.e*., genes). To do that, we made an R script (check_ratios_vs_meanratios.R) in which we simulated a hypothetical number of polymorphisms and substitutions per gene, following a Poisson distribution (we considered ∼10x more substitutions than polymorphisms, 2x more nonsynonymous positions than synonymous and 10x more functional constraint at nonsynonymous versus synonymous). We estimated *α* per window and per total. The distribution of *α* per gene can be strongly skewed to negative values when the windows become smaller, thus dragging the mean to negative values as well.

All the scripts used are available at Zenodo database (https://zenodo.org/record/6124306#.YlcVSy8RqLc).

## RESULTS

### Predominance of shared variants and global similar selective effects of mutations in genomic sequences of pig populations

We found a total of 6,684,142 SNPs in autosomes, with 149,440 SNPs located in coding regions. 12.5% of the SNPs in the coding regions are shared among the three populations, 32.2% are shared between at least two populations, 31.2% are exclusive to Large White (LW), 2.2% are exclusive to Iberian (IB) and 34.4% are exclusive to Wild boar (WB) (Table 1 and Table S7). The proportion of exclusive SNPs in each population is in accordance with its specific demographic history (Esteve-Codina et al. 2013, Bosse, Megens, Madsen, et al. 2014). Based on the PCA analysis and using the total number of SNPs, we found that the individuals of each breed cluster together and are well separated from other breeds (Figure S3A). The results from the ADMIXTURE analysis (Figure S3B) suggest that, K=2 is the most likely number of populations, where WB and IB are considered a single population. Under a K=2 scenario, only two LW and one WB individual show a significant percentage of admixture among groups. Nevertheless, for larger values of K, new groups emerge, being the IB one of these separated groups (Figure S3B). However, additional subgroups within WB and LW phenotypes appear and disappear when increasing the K value, making those subgroups apparently unreliable. Therefore, we decided to analyze separately the three breeds, LW, IB and WB, following the main patterns of population structure (Figure S3A) and the phenotypic features of the animals, which essentially separate domestic (commercial and local breed, separately) from wild animals.

**Table 1.**
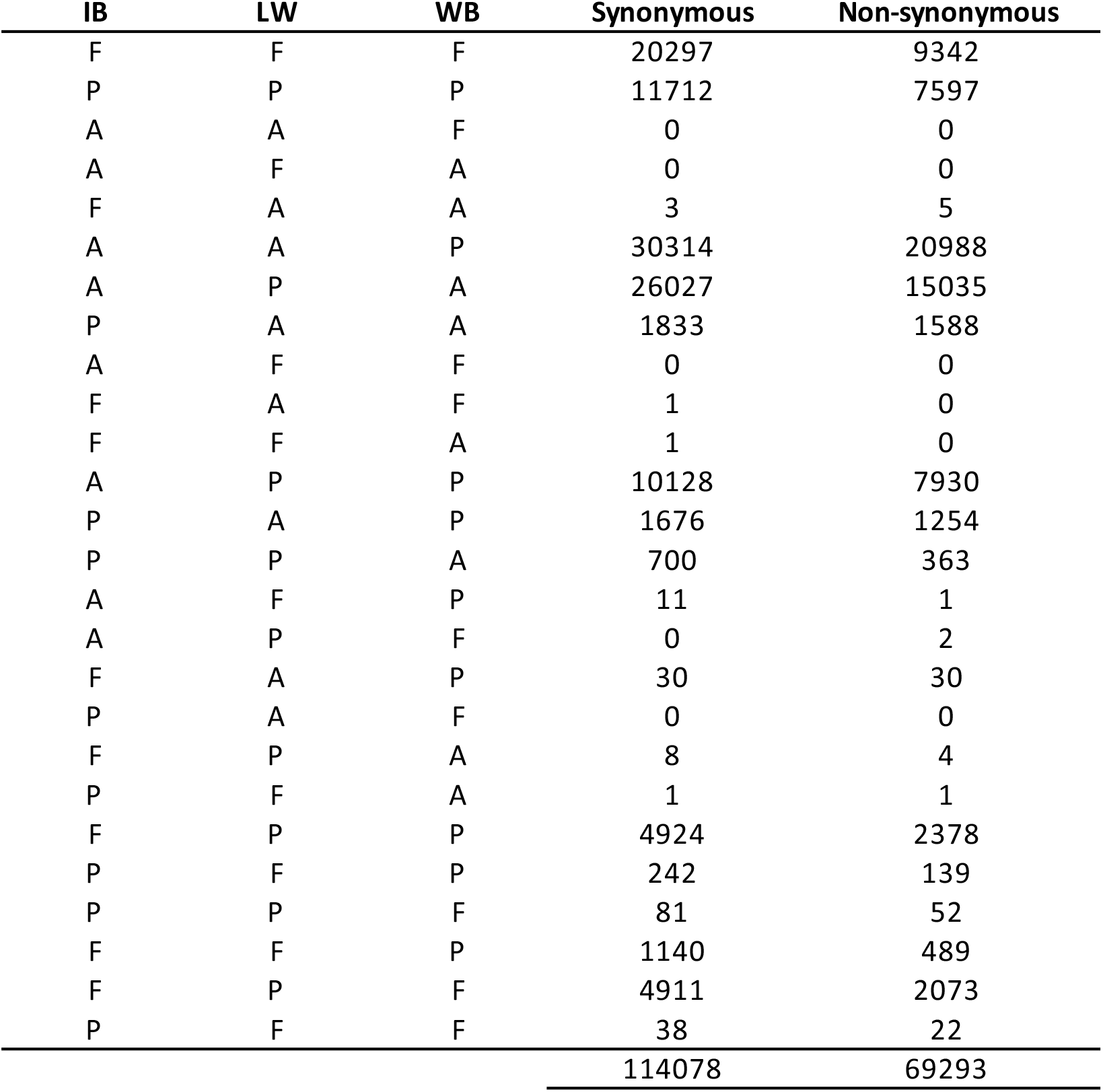
Number of synonymous and nonsynonymous SNPs according to its allelic status in each pig population. A: Ancestral allele, F: Fixed allele, P: Polymorphic allele. IB: Iberian; LW: Large White; WB: Wild boar. SNPs that are missing in any of the populations are not considered.

For each breed, coding positions were classified as polymorphic, fixed (i.e., different allele from the outgroup) or ancestral allele (i.e., same allele as in the outgroup), with the aim of identifying those variants that appeared previously or posteriorly to the domestication process (Table 1).

Surprisingly, we found very few fixed mutations between populations, indicating that the phenotypic traits of each population are not associated with fixed coding variants. Similarly, we found very few fixed coding variants in domestic (IB or LW) versus wild (WB). There are few variants fixed in the domestic breeds that are polymorphic in the wild population, suggesting that these variants were previously present in wild breeds or, alternatively, were introgressed into WB from domestic breeds. Most of the variants that are exclusive of a single breed are polymorphic, which is in agreement with the recent origin of these variants. We found a large number of fixed variants in the IB that are polymorphic in LW and WB, likely due to a reduction of the effective population size of the IB breed. The ratio of nonsynonymous to synonymous polymorphism was always lower than one and showed similar values for the three populations regardless of the variability estimator used (Table S9). This result suggests that, on average, there are no differential effects of selection between domestic and wild populations, although this might not be the case when individual genes are considered.

### Low codon bias at whole-genome scale

We estimated the level of codon bias at genome scale using MCU and *N*_*cw*_ statistics to control for the possible effect of selection on synonymous positions. Non-neutral synonymous mutations can have a large impact on the inference of the proportion of beneficial selection, and on the estimation of the Distribution of Fitness Effects (DFE). Indeed, the effect of bias in codon usage causes an overestimation of the beneficial proportion of variants that become fixed by increasing the ratio of synonymous polymorphisms versus synonymous fixations (Akashi, 1995, Matsumoto et al. 2016). For this species, we observed a low and large values of MCU and *N*_*cw*_, respectively, indicating low levels of codon bias at genome scale (mean MCU=0.485, Figure S5). However, it should be mentioned that positive selection could be acting on synonymous positions of some specific genes. We therefore have assessed whether there was a correlation between MCU and *a*, considering all coding regions or only coding regions showing positive *a* values. We observed no correlation between MCU and *a* values when considering only genes with positive *a* values (Figure S5) and slightly negative correlation when considering all genes regarding their respective *α* values (data not shown).

### Limited influence of genomic context and the network topology on selective patterns

The heterogeneity in the recombination rate, the gene density, the %GC and the distribution of CpG islands across the genome can affect the local levels of variability. A previous study on the IB breed detected a strong correlation between recombination and variability, although no correlation was observed between variability and gene density or GC content (Esteve-Codina et al. 2013). However, the effect of these factors on the estimation of the proportion of adaptive nonsynonymous mutations (*α*) has not been previously studied. When we assess whether there is a correlation between the estimated *α* and the above-mentioned factors, we found that there is no correlation between the estimated *α* and recombination, gene density, %GC and CpG in any of the three breeds (*P*-values > 0.01).

Next, we investigated the effect of gene network topology on the selective patterns. It has been claimed that topology limits the ‘evolvability’ of genes and that highly connected genes are more constrained and, consequently, less likely to be targets of positive selection. We compared the network topology features (*betweenness, out-degree* and *in-degree*) of genes within pathways regarding the estimates of *α*, grouping genes with positive versus negative *α* values. We found that genes with negative *α* values show significant large values of the *betweenness* statistic in the three pig breeds compared to genes with positive *α* values (*P*-value < 0.01; Figure S4). LW and WB showed significant values (*P*-values < 0.01) of the *in-degree* statistic for genes with negative *α* values compared to genes with positive *a* values. However, we did not observe significant differences in the *out-degree* values between genes with negative and positive *a* values in any of the three breeds (Figure S4). These results suggest that, in the three breeds, genes that are more central in a pathway are more evolutionary constrained compared to peripheral genes. In addition, in LW and WB, the genes that are more constrained tended to have a higher number of upstream genes that regulated them, which is also in agreement with the central position of these genes in the pathway. We did not observe significant differences in *in-degree* statistic in the IB breed between genes with negative and positive *a* values, likely because of a relaxation of functional constraints as a consequence of the reduction of its effective population size.

### Levels of nucleotide variation at protein coding regions are compatible with the history of the surveyed pig populations and with the presence of positive selection

To assess the selective effect of domestication, we first studied the pattern of variation at synonymous and nonsynonymous positions using four estimators of variability that differentially weight the SNP frequencies (See Material and Methods). Ideally, it would be more informative to analyze the whole genome Site Frequency Spectrum (SFS). Unfortunately, the relatively high number of positions with missing data discourages their use. A possible alternative would be to obtain the SFS from a reduced number of samples, and therefore use only a partial number of SNPs for subsequent analysis. However, by using this alternative we can either lose power or introduce some sort of bias, hence, we preferred to analyze the whole set of SNPs using those estimates of variability based on different frequencies of the spectrum that account for missing data. Nevertheless, in order to clarify the patterns of the SFS for these populations, we estimated the SFS for a subset of SNPs (around 25-30% of the available coding variants, depending on the breed) for a projection of variants on 38 haploid samples in both LW, WB and on 10 haploid samples in IB (Figure S6). The SFS profile for both synonymous and nonsynonymous showed a rapid decrease in the number of variants from lower to higher frequencies. We observed a slight increase in the number of polymorphisms at the highest frequencies at both synonymous and nonsynonymous sites and no apparent signals of admixture (i.e., no sudden peaks at specific ranges of frequencies). Estimates of whole-genome variability levels per nucleotide using different estimators are shown in Figure 1 and detailed in Table S8. We have considered the synonymous positions as neutral reference since no strong bias in codon usage has been detected (Figure S5). We expect that, under the Standard Neutral Model (SNM), the values for the different estimates of variability should be similar whereas differences among them may indicate demographic and/or selective effects. We observed that i) the levels of variability are different for each estimator within breeds and ii) the levels of variability are different for the same estimator for different breeds. However, for each breed, we observed a similar ratio of nonsynonymous to synonymous polymorphisms regardless of the used estimator, suggesting that demographic effects are responsible for the differences in the levels of variability (Figure 1). The less variable population is the IB breed, which shows far fewer singletons compared to WB and LW, probably as a consequence of the known reduction of its population size. Note than in all the three populations, high-frequency variants are proportionally more abundant than those at intermediate frequencies, which would be compatible with the accepted demographic history of the surveyed populations (i.e., introgression in LW, bottleneck in IB and some population reduction and introgression in WB) but also with the presence of pervasive positive selection in all three populations.

**Figure 1.**
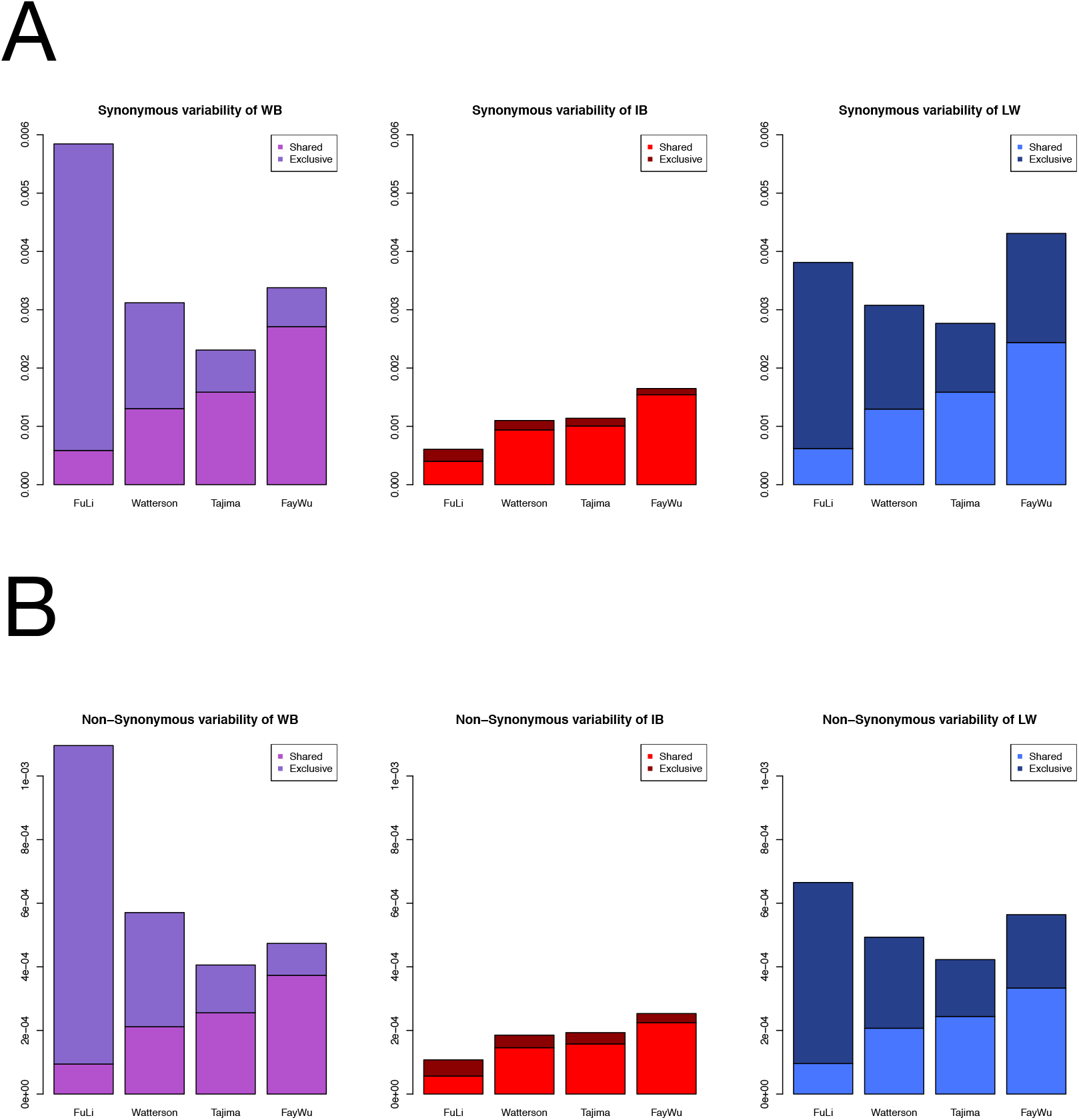
Estimates of the levels of variation at synonymous (A) and nonsynonymous (B) sites for each variability estimators and pig population and where variants were classified as shared and exclusive variants. WB; Wild boar; IB, Iberian; LW, Large White.

### *α*’s values and *R_βγ* ratios based on all SNPs might reflect a differential effect of selection due to domestication

The differential effect of selection in the domestic and wild populations can be studied by comparing their respective *α* values. Figure 2A and Table S9 show the genome-wide *α* values calculated using the four variability estimators for each population. As expected, the *α* values are negative when *α* is calculated using the estimate of variability based on low-frequency variants (*α*_Fu&Li_), probably reflecting the relatively high proportion of deleterious versus neutral mutations that are segregating at low frequencies. We observed a similar value of *α*_Fu&Li_ in all populations, suggesting a similar proportion of segregating deleterious mutations, irrespective of the domestication process or other demographic events (Figure 2A). Moreover, we observed milder negative values of *α*, or even positive for LW when *α* is calculated based on variants at high frequencies (Figure 2A), according to expectations, which point to a progressive elimination of deleterious mutations as we move towards higher frequencies. Nevertheless, the pattern of *α* (i.e., the comparative *a* value calculated using the four different variability estimators within each population) is very different in each population. WB and LW show positive or null *α* values when it is calculated based on high frequencies (Table S9). Instead, IB show very low negative *α* values for all estimators of variability. We found a compatible pattern when using the reduced subset of SNPs for the SFS estimation (Figure S6-A), where it can be observed that the estimates of *a* in all three populations are very similar among them (*a* ∼ -0.05), although their confidence intervals are quite wide.

**Figure 2.**
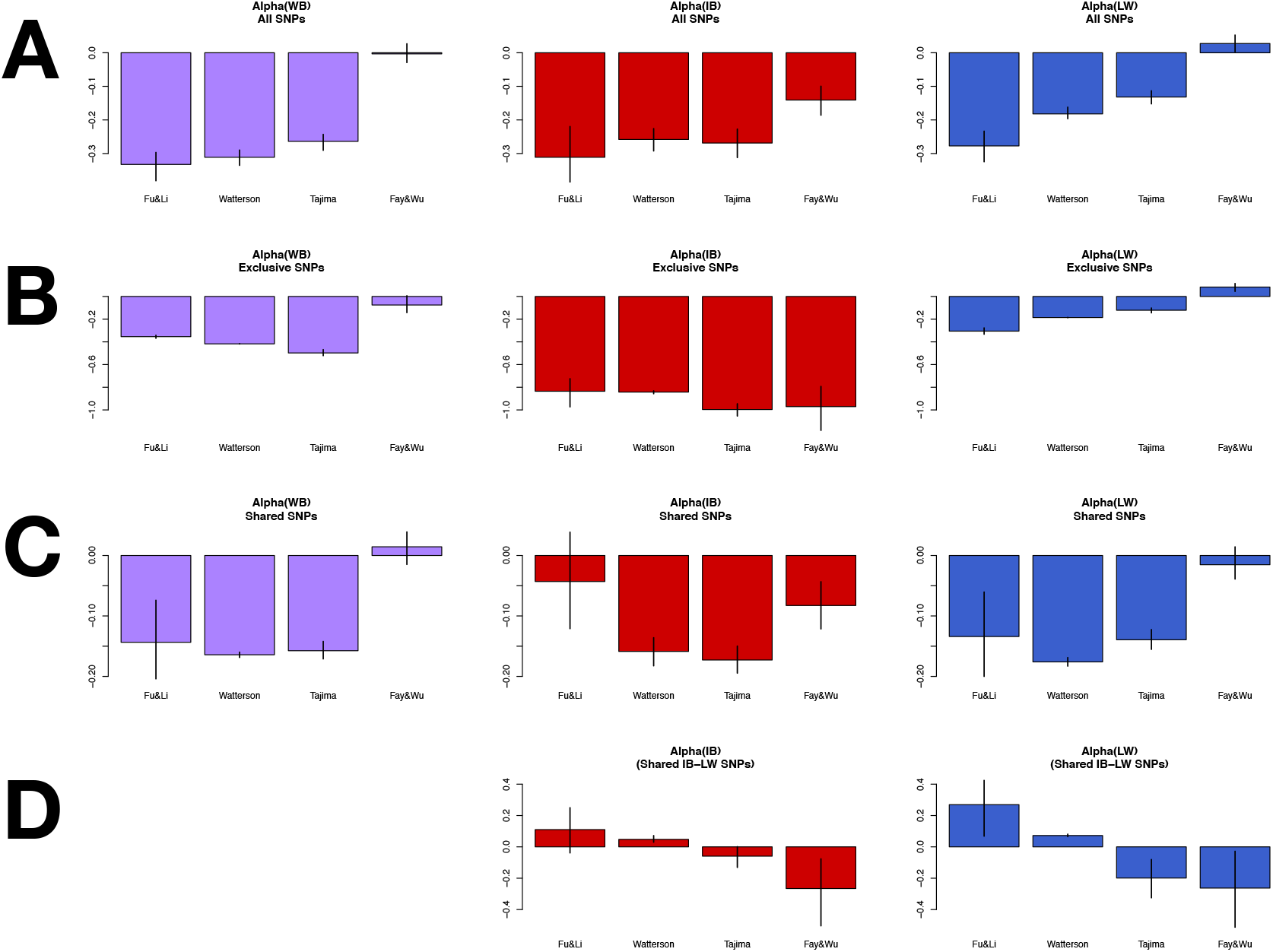
Estimates of *α* for each pig population based on different variability estimators. Total variants (A), exclusive variants (B), shared variants (C) and shared variants between IB and LW (D). Bootstrap intervals at 95% are indicated by a line at each bar. WB; Wild boar; IB, Iberian; LW, Large White.

The differences in the ratio of synonymous to nonsynonymous variability between the two different breeds is summarized by the *R_βγ* ratio (Figure 3). We observed that the largest deviations from *R_βγ* = 1 are observed when the ratio was calculated based on high-frequency variants (*α*_Fay&Wu_). Although the ratio of the two populations is difficult to interpret because of their different underlying demographic histories, some trends can be observed. WB shows an excess of nonsynonymous variants segregating at intermediate frequencies (WB-IB, WB-LW), which might be explained by a past bottleneck that increased deleterious mutations at intermediate frequencies. In addition, the *R_βγ* ratio in IB-LW shows an incremental pattern of this ratio from low to high frequencies, which is compatible with an increase of nonsynonymous beneficial variants on their way to fixation in LW.

**Figure 3.**
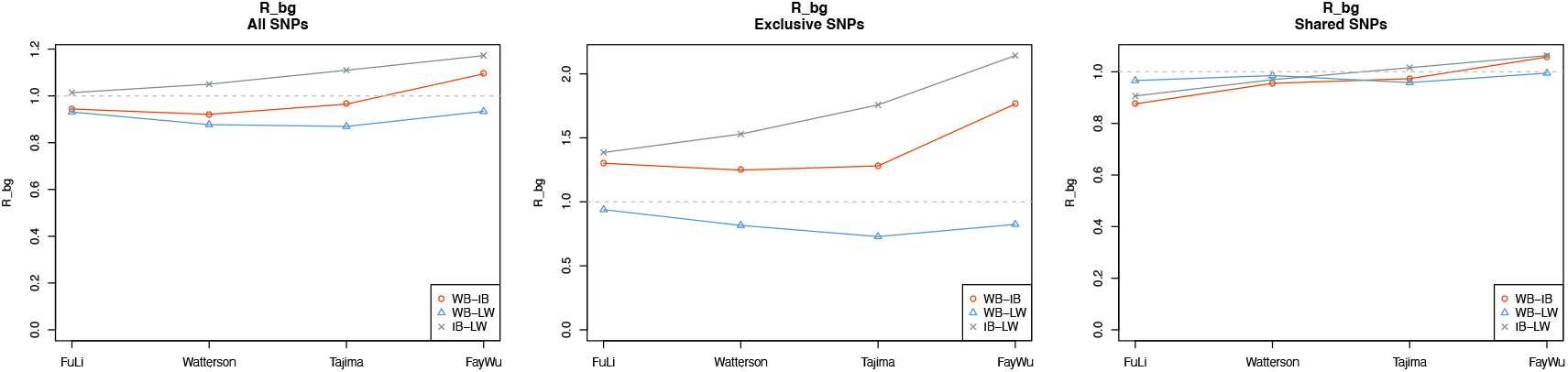
Estimates of *R_βγ* for all (left), exclusive (centre) and shared (right) variants. WB; Wild boar; IB, Iberian; LW, Large White.

### *α*’s and *R_βγ* ratios based on exclusive and shared polymorphisms might reflect changes in selective patterns before and after domestication

We observed a high ratio of nonsynonymous to synonymous singletons (*α*_Fu&Li_, Figure 2B) when the analysis was performed based on exclusive polymorphisms, suggesting that they have deleterious effects in all populations. Nevertheless, the values of *α* calculated based on intermediate frequency variants (*α*_Tajima_) in the WB and IB populations are lower than to those based on low-frequency variants, which point to a change in the selective pressure, maintaining nonsynonymous variants at relatively high frequencies. Nevertheless, the *α*_Fay&Wu_ values (−0.075, -0.971 and 0.083 for WB, IB and LW, respectively, Table S9) show a similar trend in relation to that based of Total SNPs, that is, close to 0 or positive for WB and LW, but strongly negative for IB. Concordant estimates are observed in the analysis of the SFS based on a reduced number of SNPs (0.155, -0.913 and 0.277 for WB, IB and LW, respectively, Figure 6B), with the difference that a clear positive and not 0 *α* values is observed in WB. The R_βγ statistic shows the same pattern as that calculated using all SNPs but with all over one (Figure 3). That indicates that WB has a higher proportion of nonsynonymous polymorphisms compared to IB, in contrast to what is observed when the analysis is performed based on all SNPs. This would suggest a recent change in the constraint of nonsynonymous positions likely at IB breed, as this ratio in IB-LW is also affected. This is also in agreement with the low *α* value in IB breed at exclusive variants regarding to Total SNPs.

On the other hand, the *α* values based on shared variants are in general more moderate (closer to zero) than those based on exclusive variants (Figure 2C), likely because shared nonsynonymous polymorphisms are older and hence, expected to be more functionally constrained than the exclusive ones. Additionally, the values of *α* based on singletons (*α*_Fu&Li_) are less negative than those based on intermediate-frequency variants. The *α* estimates based on shared variants in the analysis of the reduced subset of SNPs are very similar to Total SNPs and very close to zero (Figure 6C). The *R_βγ* statistic for shared variants shows similar patterns than those observed for all variants but with values much closer to 1, indicating a small or moderate selective effect on the shared variants compared to all variants (Figure 3).

When we calculated the *α* values from shared variants only between the two domestic breeds, we found an inverse pattern regarding to that calculated from all SNPs in each population, with high positive values of *α* based on low frequencies and very negative values when *α* is calculated based on high-frequency variants (Figure 2D). This could be due to i) the active elimination of new nonsynonymous variants to preserve differences among domestic breeds (*α*_Fu&Li_) and ii) the presence of nonsynonymous variants targeted by the process of domestication that shifts them toward high frequencies (*α*_Fay&wu_). Nevertheless, we cannot discard that this excess of nonsynonymous variants at high frequencies and the lack of nonsynonymous singletons at low frequency could be due to a more complex and not previously explored demographic scenario.

### Values of *α* are dependent of the molecular scale but the patterns of the estimated *α*’s are similar across the different molecular scales

In addition to the genome-wide analysis, *α* was calculated using three additional molecular scale levels: i) gene level, ii) genes within windows of 5 Mb, and iii) genes within the same pathway. Figure 4 shows the median of the distributions of the *α* values for each scale level. When the analysis was performed based on all SNPs, the pattern of *α* values estimated at the genome-wide level are concordant with those estimated at the gene level, genes within windows and genes within pathways for each breed. However, differences in the value of *α* within each breed are notorious depending on the scale level examined. The median estimates of *α* are generally lower at the gene scale level and most of them are very negative, while at the genome-wide scale, the *α* values are closer to zero. However, the distribution of *α* values can have a large variance at the gene scale since few variants are used for its estimation. We identified the regions and pathways that showed extreme *α* values (Table S10 and S11). We found a large number of genes showing *α* = 1 (highest value) because the number of polymorphic nonsynonymous variants per gene was zero. We also found a moderately high correlation of *α* values between breeds (rho ∼ 0.7, Pearson correlation when considering pathways) suggesting that in general, these breeds are under similar selective effects. When considering shared and exclusive variants, we generally observed the same pattern, from genes to whole-genome, that is, larger *α* values at the gene level and closer to zero *α* values at the larger scale. The differences in *α* values could be explained because of the distribution of this ratio statistic (*i.e*., skewed distribution to negative values) and the uneven distribution of the functional variants, in which the mean can be displaced to more negative values (see Materials and Methods).

**Figure 4.**
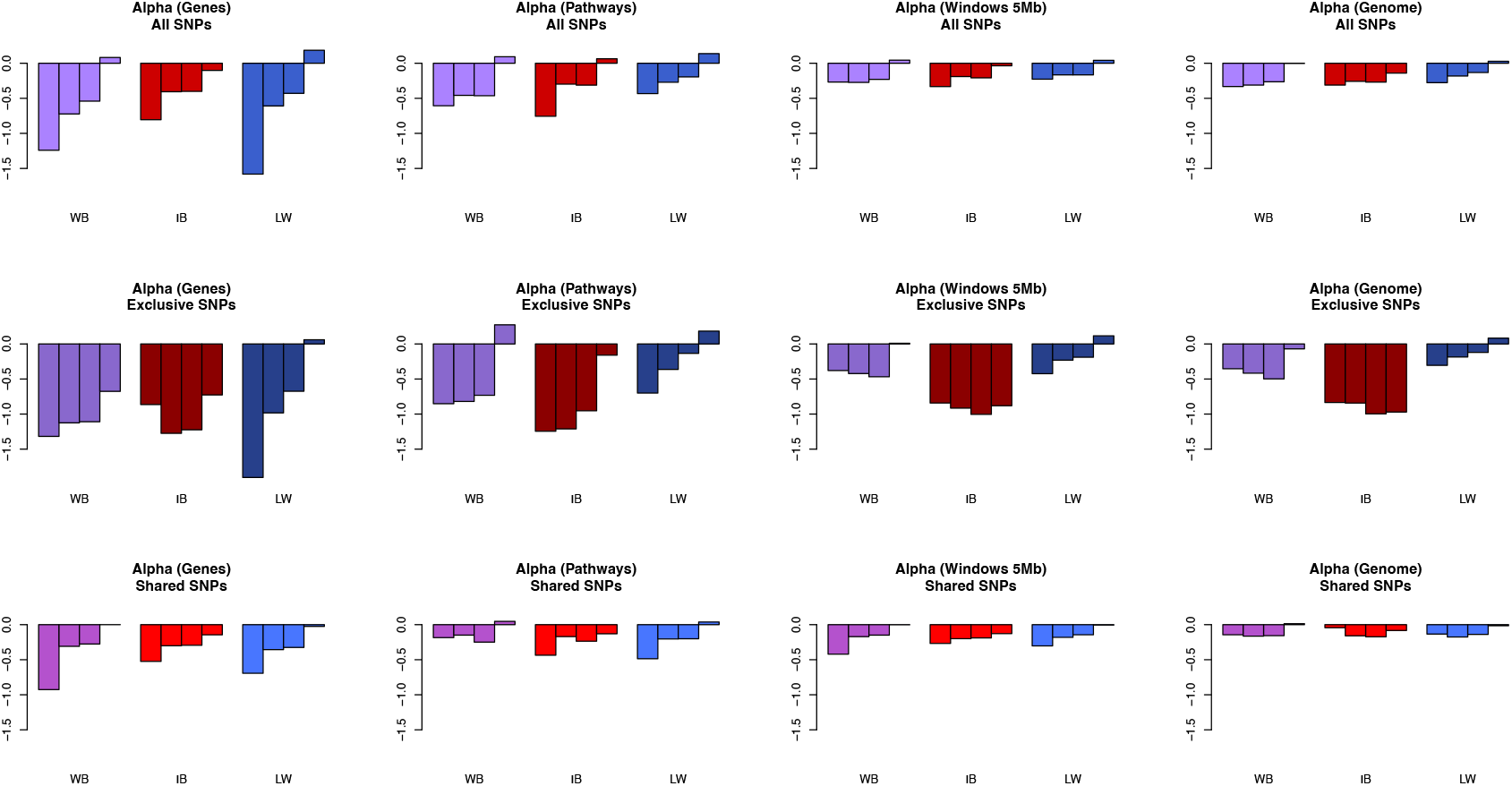
Estimates of the median values of *α* based on different variability estimators and for each pig population at different molecular scales and for all, exclusive and shared variants. Within each population, the order of different *α*’s is: Fu&Li, Watterson, Tajima and Fay&Wu. WB; wild boar; IB, Iberian; LW, Large White.

### Simulated data under different scenarios that include the joint effect of demography and selective events were more concordant to the observed data

We used computer simulations to study how the different demographic and selective events occurred during domestication process shaped nucleotide variation present in these populations. We simulated populations mimicking the process of domestication using *SLiM* software (Haller and Messer 2017), coupled with several demographic events, including changes of the population size and/or migration. We analyzed the genome-wide patterns of *α* and the *R_βγ* statistic produced by 63 simulated scenarios that included different demographic events and selective forces acting separately (simple scenarios) or jointly (complex scenarios). The results of the simulation study are summarized in Figures S7-S48. The observed patterns of *α* based on all variants in the surveyed populations are not compatible with simple scenarios that only consider demographic or positive selection forces (Figure S7). Rather, *α* patterns from simulated data (irrespective of the magnitude of *α*) fit a scenario with a predominant effect of negative selection (Figure S7). However, the *R_βγ* statistic do not fit any of the simulated simple scenarios (Figure S8). When more complex scenarios were considered (i.e., including a bottleneck, positive/negative selection and/or migration, Figures S9-S14), the general *α* patterns generated by those scenarios that include both negative and positive selection resembled those observed in WB and LW (with negative *α*’s at low frequency values to slightly positive *α* values at high frequency). The scenarios that also include some migration events are the ones that showed more concordance for these two breeds (Figures S12, S14). On the other hand, the IB population is more compatible with a scenario without positive selection and with a recent population size reduction (Figure S13). The trends in the *R_βγ* statistic are, in broad strokes, concordant with the conclusions extracted from the comparison between the observed and simulated patterns of *α* (Figures S15-S20).

The observed patterns of *α* values based on exclusive variants are similar to those based on total variants but only for the LW population (Figure S21). These patterns cannot be fully explained by any of the complex simulated scenarios that are concordant when considering all variants, although surprisingly, they would be more compatible with those including a population size reduction (Figures S24-S28). The observed *R_βγ*’s are compatible with the scenarios that combine both types of selection and a population size reduction (WB-IB) or with scenarios that include migration (WB-LW; Figures S29-S34). Finally, the observed patterns of *α* and *R_βγ* statistics calculated from shared variants are also compatible with scenarios which includes both types of selection (Figures S35-S48), and being quite compatible with those including expansion demographic events. Overall, the simulations data showed that complex scenarios, including demography, migration, and positive and negative selection, may be necessary to explain the observed data.

### Models that assume a discrete distribution of beneficial and deleterious mutations would fit better the observed data

We used an approximate Bayesian computation (ABC) analysis to infer the DFE separately for each population using the ratios of nonsynonymous to synonymous variants (i.e., polymorphism and divergence) obtained from the whole-genome analysis (see Materials and Methods). Four different models implemented in *polyDFE2* software (Tataru et al. 2019) were tested. These models overcome the inference of the demographic parameters (and others such as linkage effects) by the inclusion of nuisance parameters (see Material and Methods). The four models were: model A, which assumes a gamma distribution of deleterious mutations; model C, which assumes a gamma distribution for deleterious mutations and an exponential distribution for beneficial mutations; model DN, that assumes a discrete distribution of only deleterious and neutral mutations, and model D, that assumes a discrete distribution of deleterious, neutral and beneficial, mutations. Goodness of fit (GoF) analysis revealed that the simulated data under the different models fits differentially to the observed data, although the used range of parameters for priors are compatible with the observed data for all the four models (Table S12). Posterior probabilities showed that the DN is the most likely model for all three populations when using total number of variants (Table 2). The posterior probability for this model is especially high for the IB breed (0.97). Nevertheless, note that the model D is just below the DN model by less than half of the probability in the case of WB and LW. Finally, the posterior predictive analysis indicated that the observed *α* values for the three populations are within the range determined by the minimum and maximum simulated *α* values (i.e., Q1-1.5*IQR, Q3+1.5*IQR, respectively, being IQR the Interquantile Range Q3-Q1) under both models DN and D, although not always inside the Q1 and Q3 quantiles (Figure 5). The mean parameters of the DFE inferred for each population are shown in Table 3 (see also Table S13). The obtained results indicated that the DFE is quite similar among all three populations, which is not entirely surprising because they share a long-term history. According to model DN, and despite there is a lot of uncertainty in the inferred estimates (Table S13), the obtained results show that the DFE contains a large fraction of very deleterious variants, with approximately 75% of the variants being strongly deleterious (S =−2000), and with approximately 12% of the variants being neutral or slightly deleterious (approx. 4%). The model D infers a higher proportion of weak deleterious mutations compared to the neutral ones, although the sum of both is similar to model DN. Finally, the inferred contribution of positive selection is relatively low (around 0.7-1.1%).

**Table 2.**
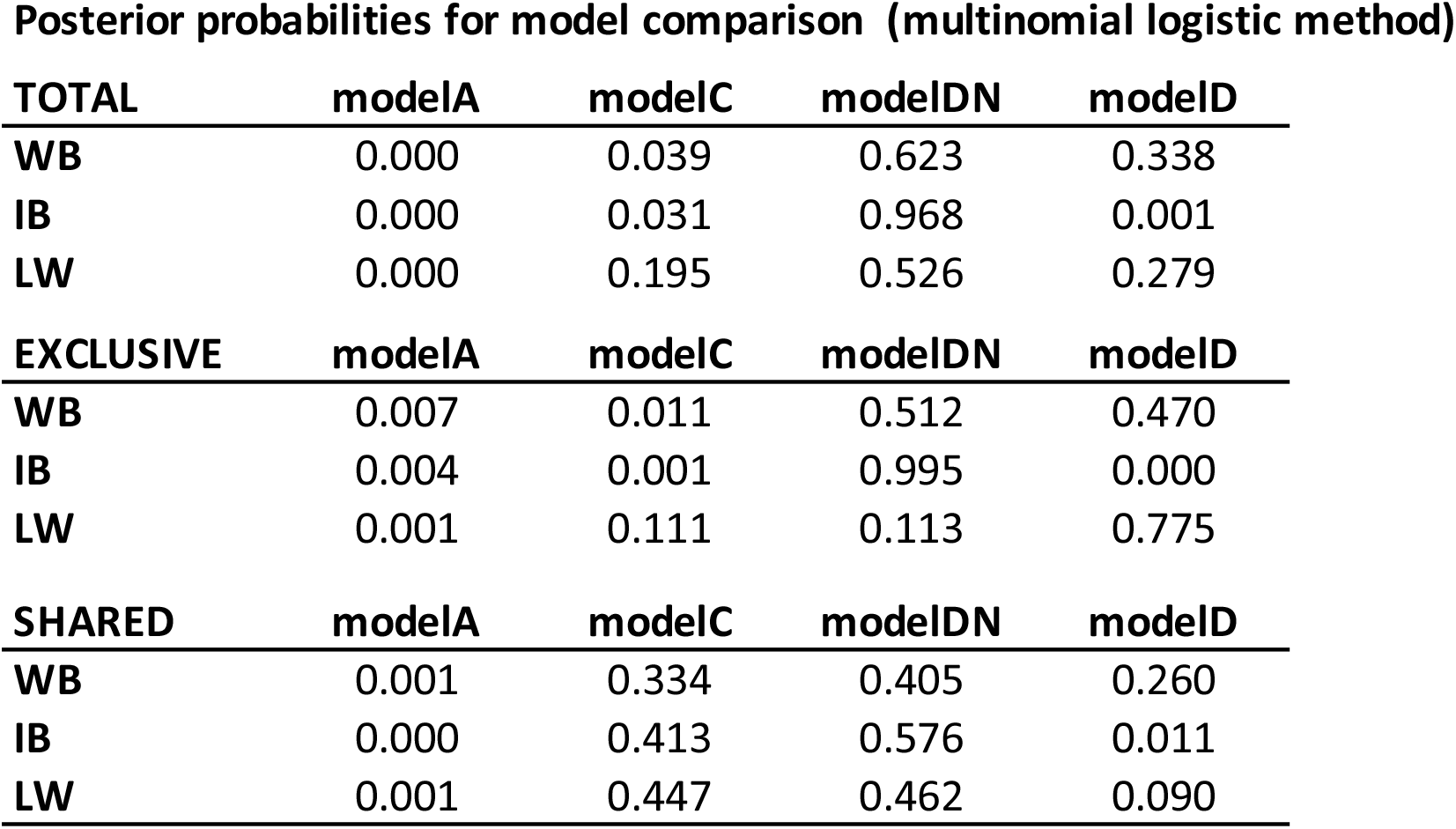
Posterior Probabilities for each ABC model (multinomial logistic method with tolerance 0.01) and for each pig population for Total variants, Exclusive variants. and Shared variants.

**Table 3.**
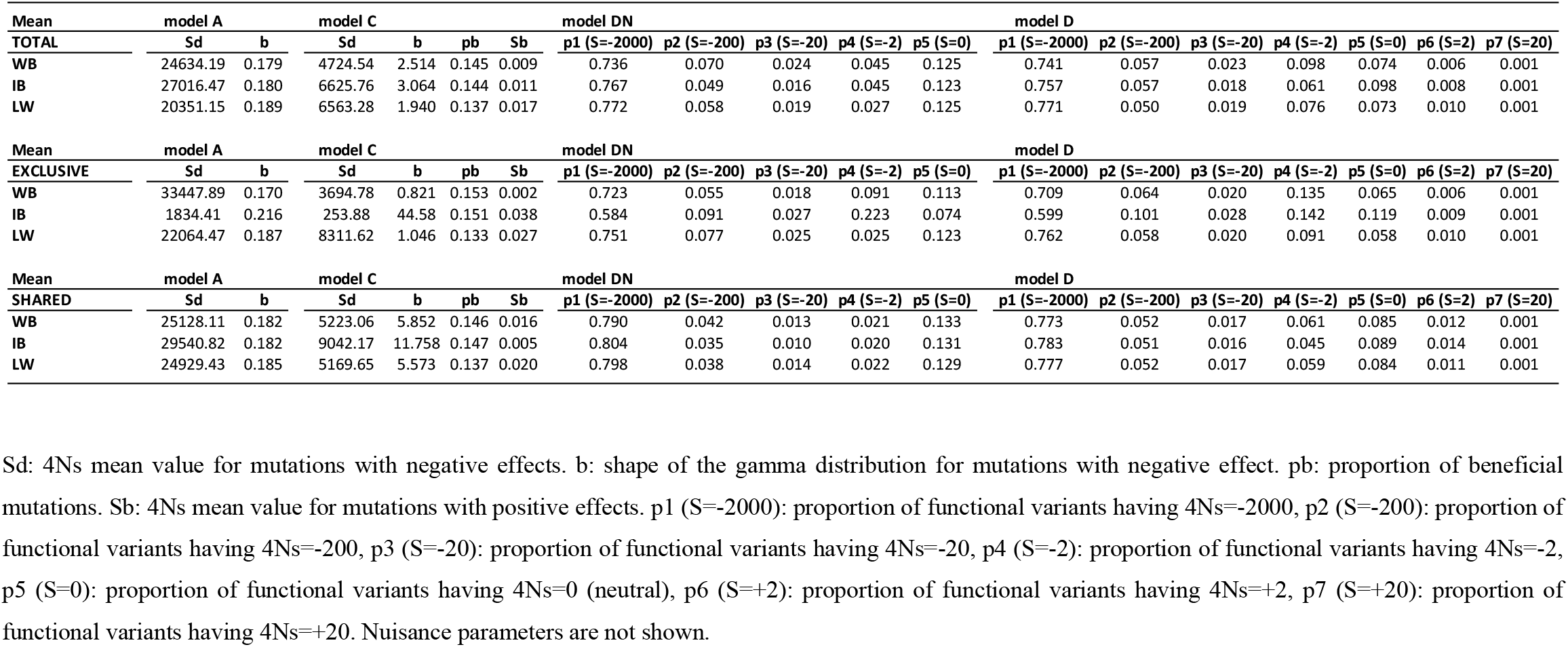
Mean Inference of parameters for each of the four analyzed models. Inferred selective parameters (weighted mean) for each ABC model and pig population. (A) Total variants. (B) Exclusive polymorphisms. (C) Shared polymorphisms. Note that for model DN and D, the proportion of each discrete S value is relative to the total S values, considering negative inferred values as zero.

**Figure 5.**
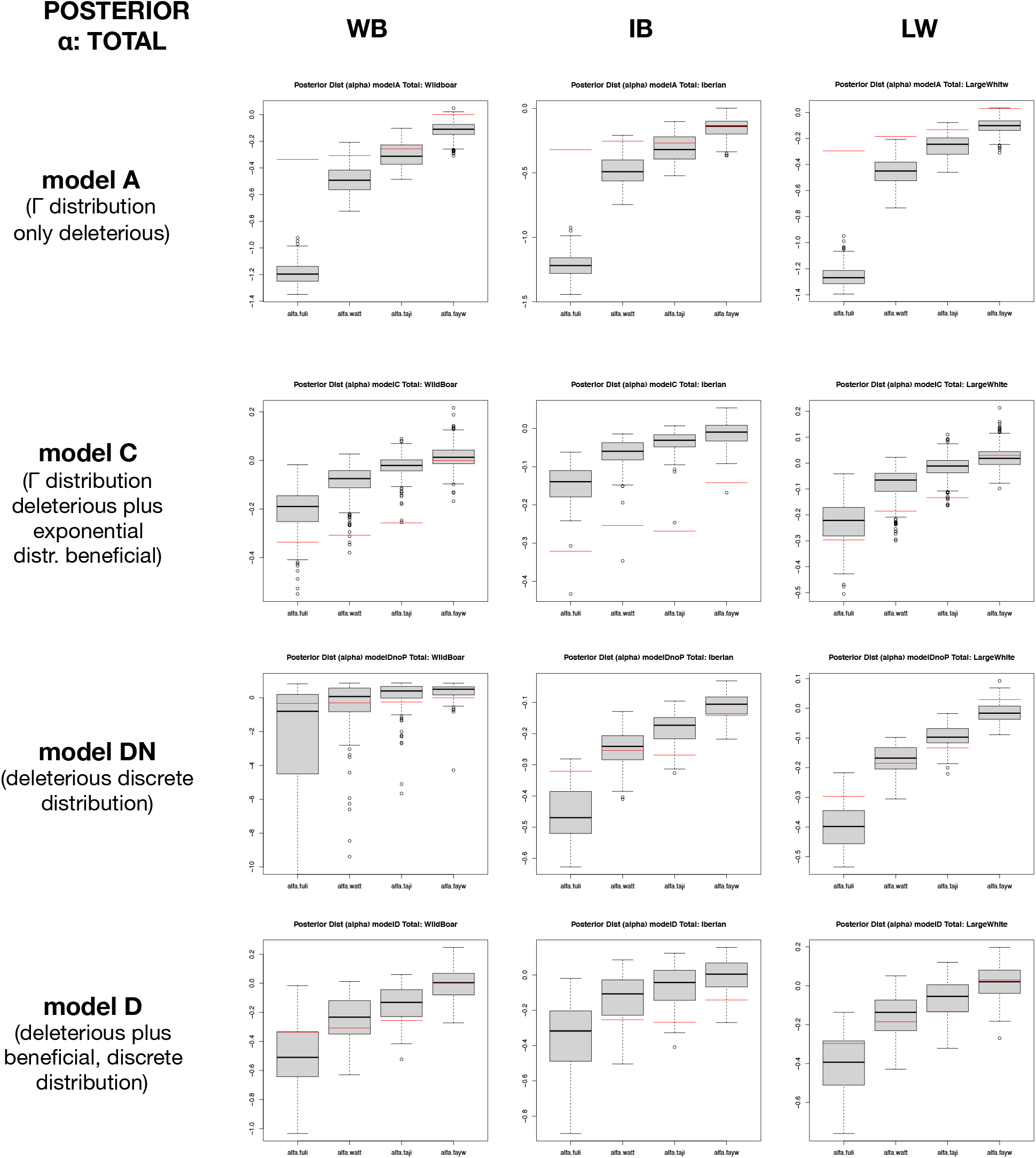
Posterior distributions of the *α* values for total variants based on different variability estimators (Fu&Li, Watterson, Tajima and Fay&Wu). Box plots indicate simulated distributions of *α* values. Red lines indicate observed *α* values.

### The differential patterns of DFE based on exclusive versus shared variants may indicate selective differences after the split of the populations

We are aware that the inference of the SFS is going to be highly distorted by choosing only a subsection of the variants (e.g., exclusive variants are mostly very recent and have no time to reach high frequencies and shared variants show no or few singletons per populations). However, the nuisance parameters incorporated in polyDFE should account for this effect (Tataru et al. 2017). The classification of polymorphisms in exclusive or shared are dependent of the relationship between two populations, and are a priori not related to the selective effect of these polymorphisms across their frequencies, although shared (mostly older) and exclusive (mostly recent) variants are chronologically related to the selection of variants. Then, we considered that the inference of DFE from exclusive and shared polymorphisms can give some clues about recent and past events related to the domestication processes. The models with higher posterior probabilities in the case of WB and IB breeds based on exclusive variants are the same than those for total variants (Table 2). However, for LW and based on exclusive variants, the model D has higher posterior probabilities, in contrast to what was obtained based on total variants (Table 2). The posterior predictive simulations showed that the models DN and D yielded similar estimated *a* values to those from the observed data, with the IB breed exhibiting the posterior distributions of *a* values more distant to the observed data (Figure 6). The estimates of the parameters of the models indicate that, for all populations, the exclusive segregating variants exhibit less strongly deleterious effects compared to those based on total and shared variants (Table 3). Indeed, the IB breed shows significantly lower proportions of strong deleterious mutations, according to its assumed recent population decline. As in the analysis based on total variants, posterior predictive analysis based on exclusive variants showed that models DN and D are those generating *a* values more similar to the observed ones, but in this case, the observed *a* values for the IB breed were slightly closer to the simulated *a*’s, compared to those based on total variants (Figure 6).The results obtained based on shared variants also show that the DN is the most likely model for all populations, although the model C shows closer probabilities (Table 2), especially in the case of LW, which might suggest that shared variants may have played a significant role as a substrate for adaptive process. However, posterior predictive distribution of *a* values for this breed under this model resembled less the observed data compared to those for the most likely models (Figure 7).

**Figure 6.**
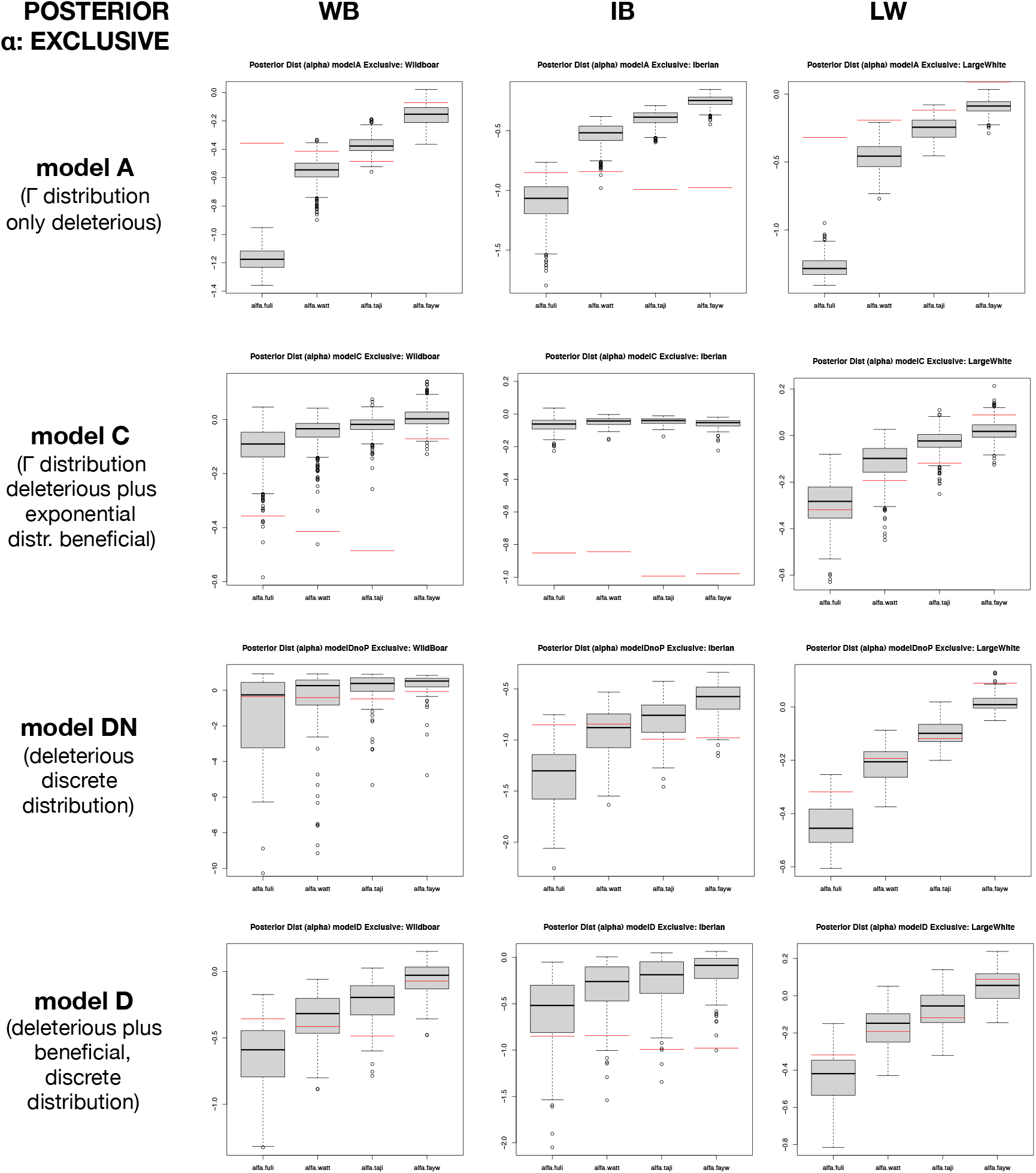
Posterior distribution of the *α* values for exclusive variants based on different variability estimators (Fu&Li, Watterson, Tajima and Fay&Wu). Box plots indicate the simulated distributions of *α* values. Red lines indicate observed *α* values.

**Figure 7.**
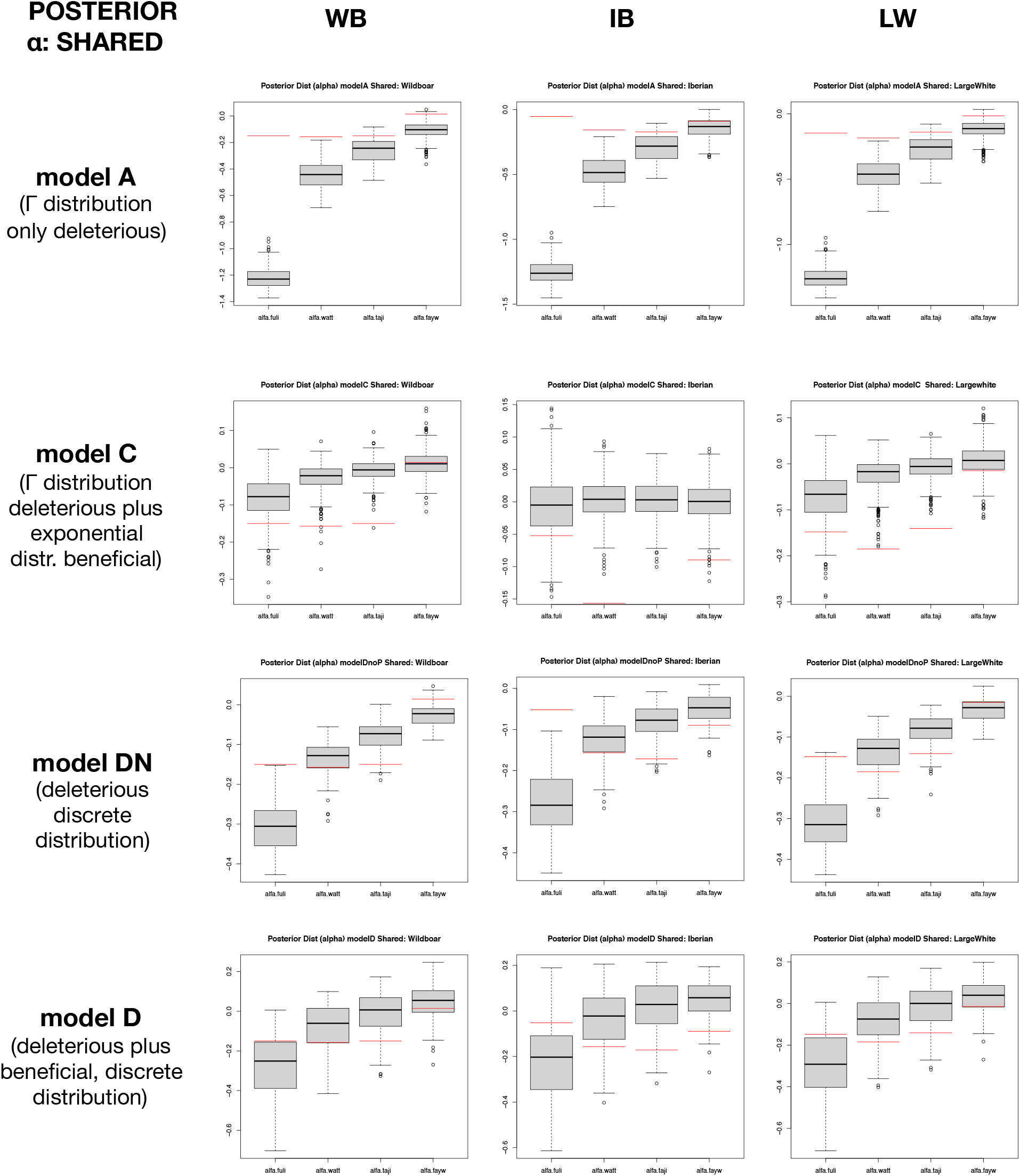
Posterior distribution of the *α* values for shared variants based on different variability estimators (Fu&Li, Watterson, Tajima and Fay&Wu). Box plots indicate the simulated distributions of *α* values. Red lines indicate observed *α* values..

## DISCUSSION

The study of the genetic effects produced by domestication can be challenged for many reasons. First, the current domesticated species have been severely manipulated by humans, which means that most of individuals were not crossed randomly, and consequently, different complex domestication scenarios can be found such as a high degree of structuration, a fast creation of new lineages from highly inbreeding crosses, a forced introgression between far related populations or from close species and a very divergent selective events across time and space, among others (e.g., Mignon-Grasteau et al. 2005, Ross-Ibarra et al. 2007, Ramos-Onsins et al. 2014, Gaut et al. 2018). Moreover, in animals, the polygenic nature of the domestication traits often precludes identifying their underlying genes since the domestic phenotypes might be probably caused by subtle allele-frequency changes of variants distributed throughout the genome, and hence, very difficult to be detect. In addition, the study of the effects of selection using genome sequences that contain a nonnegligible fraction of missing data, such as those from non-model organisms, is challenging and needs the use of appropriate methods to account for these positions. Statistics that exploit the frequency of the variants while accounting for missing data are particularly appropriate for such analyses (Ferretti, Raineri, and Ramos-Onsins 2012). Despite all this inconvenience, domestic populations are an excellent model to study the effect of strong and recent selection (e.g., Doebley, Gaut and Smith 2006, Groenen 2016).

One of the main goals of this work is to provide a novel approach that combines the use of different estimators of variability that account for missing data with the asymptotic approach proposed by Messer and Petrov (2013) and Uricchio et al. (2019) in order to take into account some of these issues. This approach is designed to be used as an alternative in case of the estimation of the full SFS is compromised by large amounts of missing data. Although it can be less precise (we used only four statistics to capture the entire trend of *α*’s across the SFS), it allows analyzing a larger number of positions, helps to reduce the variance (by summarizing the SFS into few statistics) and facilitates the visual interpretation. To illustrate the utility of the proposed approach, we used this methodology to perform an exhaustive comparative study of the observed patterns of functional versus neutral diversity and divergence in domestic pigs and wild boars. In addition, and to delve deeper into the domestication process of this species, we also performed a forward simulation study, including several diverse evolutionary scenarios and the inference of DFE parameters given different selective models. Note that the DFE was inferred using Bayesian calculations (ABC) instead of exact Bayesian or Likelihood methods since despite ABC requires additional steps and validation analysis and is in general less precise, it allows contrasting models and inferring parameters from complex datasets or data containing missing information (Beaumont et al. 2002). Finally, we have also analysed exclusive and shared polymorphisms separately to extract information about the new and the past domestication events of the demographic history of these populations, but also about the role of population admixture.

### General selection pressures on pigs and the process of domestication

Like us, others have been already performed several analyses to shed light on the process of domestication using the MacDonald and Kreitman extension methods or using other estimates such as variability or divergence at functional or synonymous positions (MacEachern et al. 2009, Kono et al. 2016, Makino et al. 2018). As in Makino et al. (2018), we do not observe an increase in functional diversity in domestic versus wild populations. This may be explained by several recent events occurring in these populations: (i) differences in the recent history our local and commercial domesticated populations (i.e., high inbreeding degree in Iberian local pigs and recent gene flow from Asian pigs into the commercial pigs); (ii) demographic effects in the wild boar population that may have reduced their diversity (Groenen et al 2016) or have increased the confidence intervals of the patterns of *α*’s (Figures S5-S46); (iii) differential adaptive forces in local (IB) versus commercial pigs (LW), with a recent high selective pressure in this last population.

Accordingly, with their recent history, the IB breed shows the lowest levels of synonymous and nonsynonymous variation among the breeds studied. Note that the two domestic breeds analyzed here have very different recent histories: the IB is a local Spanish breed (Guadyerbas) that suffered a strong bottleneck during the 1970s (Esteve-Codina et al. 2013) and with no evidence of introgression whereas the LW breed was admixed with pigs of Asian origin (Bosse, Megens, Madsen, et al. 2014). Therefore, our observations are perfectly compatible with the small effective population size and the close relatedness of the individuals expected for this population. However, for the other two breeds, the obtained results do not seem to conform to what was expected. since we detected very similar levels of variability between LW and WB, even though we expected to find higher levels of variation in the first due to the documented introgression of Asian germplasm into LW (approximately 20–35% of the genome has been estimated to be of Asian origin; Groenen et al. 2012, Bosse et al. 2012, Bosse, Megens, Madsen, et al. 2014, Frantz et al. 2015, Bianco et al. 2015, Ai et al. 2015). Interestingly, the high levels of variability were observed for variants that belonged to different frequency ranges in these two populations: singletons in WB and in high-frequency derived alleles in LW. Although these differences may be mainly due to the effects of gene flow in LW, we cannot discard an important effect of the selective programs applied to this commercial breed.

### Domestication hallmarks at pig coding regions

Another main goal of this work was finding the hallmarks of positive selection produced as a consequence of the domestication since this process implies a process of positive (human mediated) selection for traits that benefit both humans and the species of interest. The paucity of fixed variants found at coding positions in the three breeds indicates that the observed heritable phenotypic differences among the breeds are either due to: i) very few selective sweeps, ii) positive selection at noncoding functional regions that were not analyzed in here, iii) changes in the frequencies of nonsynonymous variants without being fixed. If the first hypothesis is true, we expect that domestication process should fix the adaptive variants for those genes underlaying the phenotypes of interest. However, we found no fixed variants between domestic breeds and wild pigs. Although this might be a consequence of some genetic exchange among populations, we found that the individuals were classified in groups by their location and according to their respective phenotype (Figure S3A), which suggest that the domestication features of the different breeds, even if admixed, are maintained. When we checked the *α* values for those genes that were previously reported to show signals of positive selection using other approaches (Groenen 2016), we found that these genes show little or no nonsynonymous polymorphisms or fixed variants (Table S14). This absence of variability is typical from regions under selective sweeps, although not necessarily implicating that these genes are the targets of domestication since there are no variants fixed or close to fixation at their coding regions. We only found significant values of *α* over zero at the gene KIT in the IB breed, the genes IGF2R and JMJD1C in the LW breed and the gene LRRTM3 in the WB population. This low number of genes with positive *α* values would make the first hypothesis unlikely. The second hypothesis implies that the functional regions implicated in domestication would be out of coding regions (promotors, enhancers, and others. (e.g., Li et al 2018, Rubin et al 2012, Anderson 2012). However, although being a promising hypothesis, we did not analyze those regions because it requires a very accurate analysis of homology and their associated functionality, which is very complicated at the genome level, especially for non-model species with a high proportion of missing data. The third hypothesis suggests that the domesticated phenotype is caused by a moderate change in the frequency of a relatively large number of variants with small selective effects. In this case, depending on the size of the selective effect there would only be changes in the frequencies of the variants without reaching fixation. In the last case, the functional variants involved in domestication should be segregating in the analyzed populations. These positively selected variants segregating at high frequencies, together with the presence of deleterious mutations also segregating at low frequencies, would be reflected as an excess of non-neutral polymorphism compared to divergence (*i.e*., negative *α* statistic at high frequencies). Hence, in cases where there is a significant proportion of positive selection variants that have not yet being fixed, we expect to observe a trend in the *α* slope showing more negative *α* values at intermediate-high frequencies. However, we did not observe this pattern in any of the three populations examined when the analysis was performed based on all coding positions, although it was observed for *α* values estimated based on exclusive and shared mutations, which suggest that different types of variants (total, shared and exclusive) could be capturing different aspects of the domestication process (demography versus selection).

The estimation of the DFE from the ABC analysis showed that the most likely evolutionary model for all three populations based on total variants was that consisting in a discrete DFE without significant positive selection effect (model DN; Table 2), showing a clear genome-wide effect of the action of purifying selection. We also observed a reduced effect of purifying selection in IB and in less extend in WB when the analysis was performed based on exclusive variants, which suggest a reduction of the population size of these two populations. However, for LW and when the analysis was performed based on exclusive positions, the most likely model was that with a discrete distribution that includes the effect of positive selection (model D; Table 2), which may reflect the increase of new Asian variants which increased in frequency by artificial selection.. Nevertheless, the differences between models including or excluding beneficial variants were relatively small, suggesting that, in general, a few proportion of beneficial mutations contributed to the domestication process. In fact, under model D, the estimated global proportion of beneficial mutations (weak and strong) was relatively small and slightly higher when the analysis is based on shared variants (0.1% based on total and exclusive SNPs and 1.4% based on shared SNPs; Table 3) and similar in wild and domestic populations. Nevertheless, this proportion of mutations may be substantial in absolute numbers (*i.e*., several thousand mutations).

Although an excess of nonsynonymous shared variants compared to the synonymous ones can be explained by some demographic scenarios such as bottlenecks, they may also reflect biological constraints at the species level. For instance, the phenotypic variation in a polygenic selective scenario could be caused by subtle changes in the frequencies of many genes (in an infinitesimal scenario) which would result in the observed phenotypic differences among the breeds. On the other hand, exclusive variants may reflect recent and breed-specific selective hallmarks and hence, would be responsible for the observed differences between domesticated and wild breeds. In both cases, shared and exclusive polymorphisms are contributing to the differences in the SFS between functional and non-functional positions. Nevertheless, the differences of the DFE when the analysis was performed based on total or shared variants is very small, suggesting that exclusive variants would be more informative to detect the effects of the change of selective effects.

In addition, our simulated domestication scenarios indicate that the effect of positive selection irrespective of being either strong and affecting a small percentage of variants or weak and affecting a large percentage of variants is not reflected as marked changes in the estimated patterns of *α*. This could be due to the short time since the change in the fitness effects of variants occurred but also by the interaction of positive and negative selection and demographic processes in the case of the complex scenarios, which are the most compatible with the observed data. In fact, the observed *α* patterns are compatible with the simulated demographic effects (population size reduction in WB and IB and gene flow in LW) but also, as in the ABC analysis results, with the effect of positive selection in LW when the analysis was based on exclusive variants.

We are aware that the evolutionary models used here are very simple and contain few parameters and the real observations contain high heterogeneity that could not be fitted to these models. The reasons for this heterogeneity may be technical (e.g., not adequate filtering of raw sequences), conceptual (undetected correlations that distort model assumptions) or biological (too simplistic models to explain the real data). In any case, the model that assumes a discrete distribution of deleterious mutations (model DN) seems to generally explain better the observed data, together with the model D (model DN but including the effect of beneficial selection) in a minor degree and also that exclusive variants seemed to be more informative to detect the changes of the selective effects.

### Final remarks

The observed patterns of variability are compatible with the presence of deleterious mutations segregating in all three breeds but also with weak signals of positive selection. In addition, when the variants are split into shared and exclusive, we observed patterns that are in line with the simulated data under different demographic scenarios with the joint action of positive and negative selection. We found a clear effect of deleterious mutations at low-frequency variants and a possible mild effect of positive selection at higher frequencies. However, additional analyses contrasting evolutionary models that consider the effects of standing variation, whose effect change under the domestication process, may shed more light and will help to understand the patterns of variation shaped by the domestication process.

## Supporting information

Comparison_1st_vs_2nd_submissiion

## ACKNOWLEDGMENTS

We acknowledge L. Silió and M. C. Rodríguez for helpful comments on the manuscript. S.E.R.O. is supported by grants AGL2016-78709-R (MEC, Spain) and PID2020-119255GB-I00 (MICINN, Spain), by the CERCA Programme/Generalitat de Catalunya and acknowledges financial support from the Spanish Ministry of Economy and Competitiveness, through the Severo Ochoa Programme for Centres of Excellence in R&D 2016-2019 and 2020-2023 (SEV-2015-0533, CEX2019-000917) and the European Regional Development Fund (ERDF). S.G-R. was supported by a Beatriu de Pinós postdoctoral fellowship (AGAUR; 2014 BP-B 00027).

## SUPPLEMENTARY MATERIAL

See Supplementary Material file added to see the Tables (S1-S14) and Figures (S1-S48).

