## Supplementary material for "Pervasive selection pressure in wild and domestic pigs": Comparison_1st_vs_2nd_submissiion

**Keywords**: Domestication, Distribution of fitness effects, Proportion of beneficial mutations, Approximate Bayesian calculation, Polygenic selection, Population genomics

**Deleted:** behavior
**Deleted:** they are
**Deleted:** approaches
**Deleted:** based on
**Deleted:**
**Deleted:** , according to
**Deleted:** based on
**Deleted:** .
**Deleted:** the pig species, analyzing
**Deleted:** complete
**Deleted:** 20 European wild boars, 6 Iberian and 20 Large White pigs at …
**Deleted:** molecular scales: gene, metabolic pathway and whole-genome.
Descriptive variability analyses on pig
**Deleted:** indicate
**Deleted:** Instead, most variants are shared
**Deleted:** them, despite that the phenotypes of wild and
**Deleted:** individuals are clearly
**Deleted:** . Additionally, asymptotic MK plots based on summary statistics show that small effects variants may affect the final calculation of $\alpha$, the proportion of beneficial mutations. The distribution of fitness effects inferred with Approximate Bayesian Computation analysis
**Deleted:** an important quantity
**Deleted:** (~83% of total mutations)
**Deleted:** (~17%)
**Deleted:** Exclusive
**Deleted:** show that recent demographic changes
**Deleted:** Finally, the median proportion of the strong favorable mutations are very scarce in all cases (≤ 0.2%). The median estimated alpha values (weak and strong favorable) are 0.9% for wild and domestic pigs.
**Deleted:** ¶
¶

**Deleted:** Raw reads for
**Deleted:** as with the SNPs
**Deleted:** as with the SNPs

using *samtools depth* utility, *BEDtools* (Quinlan 2014) and custom scripts (available at , Pérez-Enciso *et al*., 2017). This resulted in a *gVCF* file per individual with the information about variant calls and non-varying positions. Next, each *gVCF* file was converted into a fasta file and all fasta files were subsequently merged to obtain a multindividual *gVCF* file (Pérez-Enciso et al. 2016).

**Deleted:** either

**Deleted:** using 5 Mb

**Deleted:** at each functional

**Deleted:** region.

**Deleted:** Specifically,

**Deleted:** using the Ferretti, Raineri, and Ramos-Onsins (2012) expression:

$$\hat{\theta} = \frac{1}{L}\sum_{x=1}^{L}\sum_{i=1}^{n_x-1} i\omega_{i,n_x}\xi_i(x), \qquad \frac{1}{L}\sum_{x=1}^{L}$$

*(Equation 1)*
where ($\omega_i$) is the weight

**Deleted:** the different

**Deleted:** estimators such as $\omega$

**Deleted:** $= n/(i(n-i)(1+\partial_{i,n-i}))$ for the Watterson estimator, $\omega_i = n/(1+\partial_{i,n-i})$ for the Tajima estimator (both for folded spectrum), $\omega_i = i$ for the Fay&Wu estimator and $\omega_i = 1$, $\omega_{i>1} = 0$ for the Fu&Li estimator …

**Deleted:** .

**Deleted:** artifactual

**Deleted:** positively

addition, functional positions (nonsynonymous positions) are constrained compared to non-functional positions (synonymous positions), and hence, their evolutionary ratios are smaller. In the neutral scenario, polymorphism and divergence (excluding the adaptive fixed variants) are proportional to the mutation rate and to the constriction factor in the case of nonsynonymous positions (McDonald and Kreitman 1991, Eyre-Walker 2006, Fay 2011). That is:

$$\frac{\theta_n}{\theta_s} = \frac{(1-\alpha)K_n}{K_s},$$

*(Equation 1)*

where $\theta_n$ is the nonsynonymous variability, $\theta_s$ is the synonymous variability, $K_n$ is the nonsynonymous divergence, $K_s$ is the synonymous divergence and $\alpha$ is the proportion of adaptive variants that have been fixed. To estimate the proportion of nonsynonymous substitutions that are adaptive ($\alpha$) the previous expression is reordered (e. g., Eyre-Walker 2006):

$$\alpha = 1 - \frac{K_s}{K_n}\frac{\theta_n}{\theta_s}$$

*(Equation 2)*

A higher ratio of nonsynonymous to synonymous divergence versus polymorphisms suggests that positive selection has fixed adaptive variants ($\alpha > 0$) and the opposite case ($\alpha < 0$) suggests the presence of deleterious mutations segregating in the population.

If we consider that weak deleterious mutations are segregating in the population, we expect that their relative proportion will be higher at lower frequency variants and low or zero for fixed deleterious mutations. Following the same notation as in equation 2:

$$\frac{\theta_{in}(1-\beta_i)}{\theta_{is}} = \frac{(1-\alpha-\beta_d)K_n}{K_s}, \frac{\theta_{in}(1-\beta i)}{\theta_{is}} \frac{(1-\alpha-\beta d)K_n}{K_s}$$

*(Equation 3)*

where $i$ refers to the frequency at which the calculation of variability is estimated, $\beta_i$ is the proportion of weakly deleterious polymorphic mutations at frequency $i$, $\beta_d$ is the proportion of weakly deleterious fixed mutations. $\beta_d < \beta_i$ was assumed at any frequency. Then, solving for the proportion of fixed adaptive variants ($\alpha$):

**Deleted:** nonfunctional

**Deleted:** *2*

**Deleted:** we reorder

**Deleted:** :

**Deleted:** *3*

**Deleted:** null

**Deleted:** 3

**Deleted:** =

**Deleted:** ,

**Deleted:** *4*

**Deleted:** $\beta d$

$$\alpha = 1 - \beta_d - (1 - \beta_i)\frac{K_s}{K_n}\frac{K_s}{K_n}\frac{\theta_{in}}{\theta_{is}}$$

*(Equation 4)*

Deleted: $\beta d -$ (
Deleted: $\beta i$)
Deleted: 5

We see that in case of calculating $\alpha$ without considering the effects of deleterious mutations, this would be underestimated depending on the frequency at which the estimates of variability are calculated. If we assume that the deleterious variants would hardly be fixed, a good estimator of $\alpha$ using equation 2 would be the one that estimates variability based on high frequencies, as it would hardly contain deleterious mutations. This is in agreement with the asymptotic arguments used in Messer and Petrov 2013) and implemented in Haller and Messer (2017).

Deleted: it
Deleted: detrimental
Deleted: never
Deleted: 3
Deleted: detrimental
Deleted: .

Similarly, if we also consider that weak positively selected variants are segregating in the population, we expect that their relative proportion, compared to neutral ones, is higher at higher frequencies:

$$\frac{\theta_{in}(1 - \beta_i - \gamma_i)}{\theta_{is}} = \frac{(1 - \alpha - \beta_d - \gamma_d)K_n}{K_s}, \frac{\theta_{in}(1 - \beta i - \gamma i)}{\theta_{is}} \frac{(1 - \alpha - \beta\, d - \gamma d)K_n}{K_s}$$

*(Equation 5)*

Deleted: =
Deleted: ,
Deleted: 6

where $\gamma_i$ is the proportion of weakly advantageous polymorphic mutations at frequency $i$, and $\gamma_d$ is the proportion of weakly advantageous fixed mutations. Again, solving for the proportion of fixed adaptive variants ($\alpha$+$\gamma_d$):

Deleted: $\gamma i$
Deleted: $\gamma d$
Deleted: ):

$$\alpha + \gamma_d = 1 - \beta_d - (1 - \beta_i - \gamma_i)\frac{K_s}{K_n}\frac{K_s}{K_n}\frac{\theta_{in}}{\theta_{is}}$$

*(Equation 6)*

Deleted: $\gamma d =$
Deleted: $\beta d -$ (
Deleted: $\beta i - \gamma i$)
Deleted: 7

In this case, the presence of adaptive variants segregating in the population would affect the estimates of variability based on high frequency variants when using equation 2, which would result in an underestimation of the proportion of fixed adaptive variants ($\alpha$). Note that adaptive variants stabilized at intermediate frequencies, which can be an important source of adaptation considering the infinitesimal model, are not considered in this approach.

Deleted: 3
Deleted: are not considered in this approach

If we focus on the effects of polymorphic weakly selected mutations, equation 5 suggests that the ratio of nonsynonymous to synonymous polymorphisms would increase due to mutations having both positive and negative effects. It is expected that the number of mutations with negative selection coefficients would rapidly decrease as we move to intermediate and high frequencies, while the opposite trend is expected for mutations with positive selection coefficients. Hence, higher ratios of nonsynonymous to synonymous polymorphisms at higher frequencies may be explained by the presence of advantageous mutations segregating in the population.

Furthermore, in cases where two populations are from the same species and there are no fixed mutations between them (e.g., they have equal divergence ratios versus the outgroup), we can estimate the possible differential effect of the selection (positive and negative together) at any frequency between populations from the ratios of synonymous to nonsynonymous polymorphisms of the two populations ($R\_\beta\gamma_i$):

$$\frac{\theta_{in1}(1-\beta_{i1}-\gamma_{i1})}{\theta_{is1}} = \frac{\theta_{in2}(1-\beta_{i2}-\gamma_{i2})}{\theta_{is2}} \frac{\theta_{in1}(1-\beta i1-\gamma i1)}{\theta_{is1}} \frac{\theta_{in2}(1-\beta i2-\gamma i2)}{\theta_{is2}}$$

and

$$\frac{(1-\beta_{i1}-\gamma_{i1})}{(1-\beta_{i2}-\gamma_{i2})} = \frac{\theta_{is1}\theta_{in2}}{\theta_{in1}\theta_{is2}} = R\_\beta\gamma_i \frac{(1-\beta i1-\gamma i1)}{(1-\beta i2-\gamma i2)} \frac{\theta_{is1}\theta_{in2}}{\theta_{in1}\theta_{is2}}$$

$$\frac{(1-\beta i1-\gamma i1)}{(1-\beta i2-\gamma i2)} \frac{\theta_{is1}\theta_{in2}}{\theta_{in1}\theta_{is2}}$$

*(Equation 7)*

In addition, a comparison of the $R\_\beta\gamma_i$ values calculated using different variability estimators (hereafter $R\_\beta\gamma_i$ pattern) can be used to inform about the effects of selection. For example, values over 1 indicate that the population 2 has a higher ratio of nonsynonymous to synomymous polymorphisms compared to population 1, either produced by an accumulation of deleterious or of beneficial polymorphisms. Importantly, different demographic effects (e.g., bottlenecks) together with the presence of mutations with small selective effects may also disturb the ratios of variability and hence must be considered when interpreting the results. We include a couple of possible scenarios that can account for possible patterns: (i) after split of two the two populations, both populations have the same population size, but population 1 is affected by the action of

**Deleted:** on the polymorphisms
**Deleted:** 6
**Deleted:** among
**Deleted:** $\beta\gamma i$):
**Deleted:** =
**Deleted:** and
**Deleted:** =
**Deleted:** =
**Deleted:** = $R\_\beta\gamma i$
**Deleted:** = $R\_\beta\gamma i$
**Deleted:** *8*
**Deleted:** the different types of selection.
**Deleted:** taken into account

positive selection on a quantitative trait (polygenic effect), which causes an increase in the frequencies of some of its variants without getting fixed. Under this scenario, we expect a R_βγ > 1 when this is calculated based on high frequencies. (ii) after split of two the two populations, the population 2 remains equal population size as before the split and the population 1 suffers a reduction in its effective population size, which causes that the slightly deleterious mutations become effectively neutral. Then, R_βγ is expected to be > 1 when it is calculated based on low frequency variants.

The effect of linkage disequilibrium between selective (deleterious or adaptive) and neutral variants should not overly affect the expected estimate of the proportion of adaptive variants, as it would affect both synonymous and nonsynonymous positions in similar proportion. On the contrary, the interaction of variants with opposite selective effects would possibly reduce the effect of selection and would have a significant consequence on the estimation of adaptive fixed variants (Hill and Robertson 1966; Booker and Keightley 2018).

**Deleted:** detrimental

**Deleted:** the same

### Bootstrap analysis

Nonparametric bootstrap analysis was performed to estimate the null distribution of the $\alpha$ statistic for each variability estimator and pig population. In each case, synonymous and nonsynonymous coding positions were randomly chosen with replacement and the $\alpha$ statistic was calculated as in equation 1. This process was repeated 100 times.

**Gene context and network topology analysis**

**Deleted:** , while others

**Deleted:**

**Deleted:** With the aim of performing the ABC using summary statistics, the

**Deleted:** .

**Deleted:** 70

**Deleted:** 60 and

**Deleted:** D

**Deleted:** 67, with a

**Deleted:** compares the median of the distance between the accepted summary statistics and the observed ones

**Deleted:** to select the

**Deleted:** .

**Deleted:** with

**Deleted:** (instead of with the ratios of variability and of divergence, to avoid circularity in the analysis)

**Deleted:** which is a simple

**Deleted:** **Pathway**

We downloaded the complete list of pathways and genes of S. *scrofa* from KEGG v.20170213 (http://www.genome.jp/kegg/, Kanehisa et al. 2008). The list contained 471 pathways and 5,480 genes. The median and mean number of genes per pathway was 26 and 43, respectively, and ranged from 1 to 949. We filtered the pathways according to their size, removing pathways with less than 10 and more than 150 genes in order to discard pathways that were not informative or too generic and complex. The final list contained 171 pathways and 3,449 genes.

**Deleted:** *Sus*

**Deleted:** 5480

**Deleted:** 3449

To analyze the selection pressure of each gene according to its position in the pathway, we obtained different topological parameters. For that, we first downloaded the XML file of each pathway from KEGG v.20170213. These files were analyzed with the *iGraph* R package (Csardi G. and Nepusz T. 2006) to obtain the topological descriptors of each gene in each pathway. For each gene, three different measures were computed: *betweenness* (number of shortest paths going through a vertex), *in-degree* (number of in-going edges) and *out-degree* (number of out-going edges). These parameters are measures of the importance of a gene within a pathway: *betweenness* is a centrality feature, *in-degree* suggests the facility of a protein to be regulated and *out-degree* reflects the regulatory role of a protein. We tested whether negatively and positively selected genes differed in any of these statistics using a nonparametric Wilcoxon rank test, due to the extreme leptokurtic distributions involved.

**Genomic context patterns**

We have additionally tested whether there is a significant correlation between $\alpha$ and recombination, gene density, missing rate, %GC and CpG islands across genomes.

**Testing the differences in the estimates of $\alpha$ using whole-genome data versus the mean of gene estimates.**

**Deleted:** ------------------Page Break------------------ ¶

**Deleted:** on

**Deleted:** of these

**Deleted:** We found that

**Deleted:** ,

**Deleted:** ). Based on the PCA analysis and using the total number of SNPs, we found that the individuals of each breed cluster together and are well separated from other breeds (Figure S3).

**Deleted:** private

**Deleted:** .

**Table 1.** Number of synonymous and nonsynonymous SNPs according to its allelic status in each pig population. A: Ancestral allele, F: Fixed allele, P: Polymorphic allele. IB: Iberian; LW: Large White; WB: Wild boar. SNPs that are missing in any of the populations are not considered.

| IB | LW | WB | Synonymous | Non-synonymous |
|---|---|---|---|---|
| F | F | F | 20297 | 9342 |
| P | P | P | 11712 | 7597 |
| A | A | F | 0 | 0 |
| A | F | A | 0 | 0 |
| F | A | A | 3 | 5 |
| A | A | P | 30314 | 20988 |
| A | P | A | 26027 | 15035 |
| P | A | A | 1833 | 1588 |
| A | F | F | 0 | 0 |
| F | A | F | 1 | 0 |
| F | F | A | 1 | 0 |
| A | P | P | 10128 | 7930 |
| P | A | P | 1676 | 1254 |
| P | P | A | 700 | 363 |
| A | F | P | 11 | 1 |
| A | P | F | 0 | 2 |
| F | A | P | 30 | 30 |
| P | A | F | 0 | 0 |
| F | P | A | 8 | 4 |
| P | F | A | 1 | 1 |
| F | P | P | 4924 | 2378 |
| P | F | P | 242 | 139 |
| P | P | F | 81 | 52 |
| F | F | P | 1140 | 489 |
| F | P | F | 4911 | 2073 |
| P | F | F | 38 | 22 |
| | | | 114078 | 69293 |

**Deleted:** value

**Deleted:** (but not IB, possibly because the low sample size) show

**Deleted:** perhaps

**Levels of nucleotide variation at protein coding regions are compatible with the history of the surveyed pig populations and with the presence of positive selection**

**Deleted:** is

**Deleted:** Estimates of

**Deleted:** levels

**Deleted:** at the genome level

**Deleted:** S7.

**Deleted:** all

**Deleted:** while

**Deleted:** also

**Deleted:** for all estimators

**Deleted:** variability

which would be compatible with the accepted demographic history of the surveyed populations (i.e., introgression in LW, bottleneck in IB and some population reduction and introgression in WB) but also with the presence of pervasive positive selection in all three populations.

Deleted: the LW

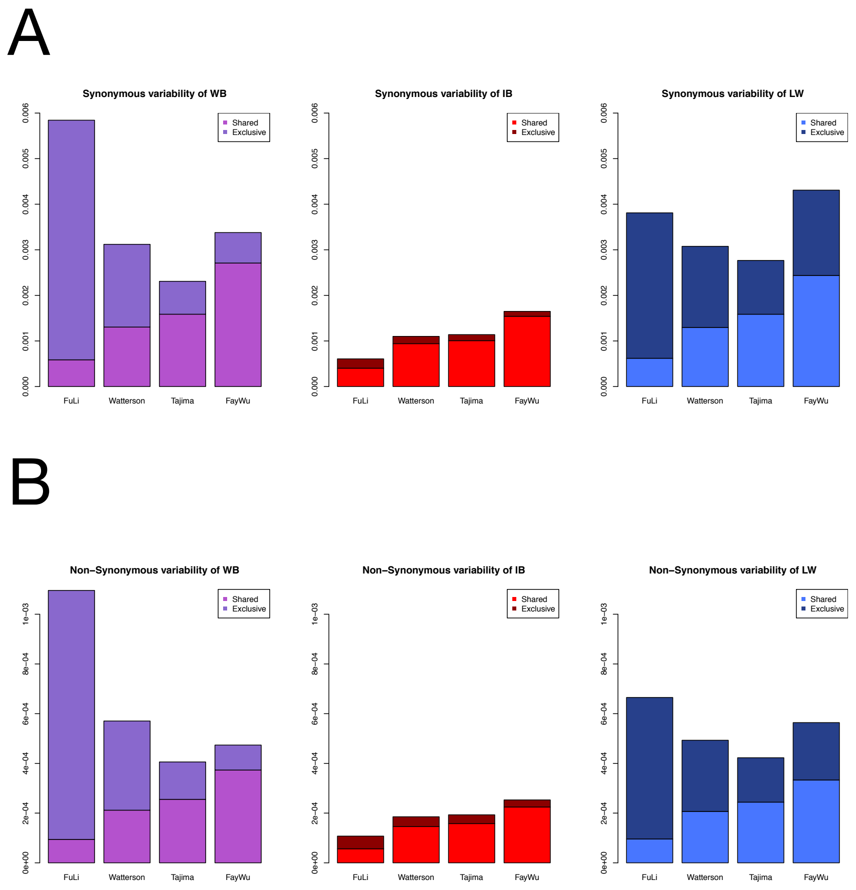


**Figure 1.** Estimates of the levels of variation at synonymous (A) and nonsynonymous (B) sites for each variability estimators and pig population and where variants were classified as shared and exclusive variants. WB; Wild boar; IB, Iberian; LW, Large White.

**$\alpha$'s values and $R\_\beta\gamma$ ratios based on all SNPs might reflect a differential effect of selection due to domestication**

The differential effect of selection in the domestic and wild populations can be studied by comparing their respective $\alpha$ values. Figure 2A and Table S9 show the genome-wide $\alpha$ values calculated using the four variability estimators for each population. As expected, the $\alpha$ values are negative when $\alpha$ is calculated using the estimate of variability based on low-frequency variants ($\alpha_{Fu\&Li}$), probably reflecting the relatively high proportion of deleterious versus neutral mutations that are segregating at low frequencies. We observed a similar value of $\alpha_{Fu\&Li}$ in all populations, suggesting a similar proportion of segregating deleterious mutations, irrespective of the domestication process or other demographic events (Figure 2A). Moreover, we observed milder negative values of $\alpha$, or even positive for LW when $\alpha$ is calculated based on variants at high frequencies (Figure 2A), according to expectations, which point to a progressive elimination of deleterious mutations as we move towards higher frequencies. Nevertheless, the pattern of $\alpha$ (i.e., the comparative $\alpha$ value calculated using the four different variability estimators within each population) is very different in each population. WB and LW show positive or null $\alpha$ values when it is calculated based on high frequencies (Table S9). Instead, IB show very low negative $\alpha$ values for all estimators of variability. We found a compatible pattern when using the reduced subset of SNPs for the SFS estimation (Figure S6-A), where it can be observed that the estimates of $\alpha$ in all three populations are very similar among them ($\alpha \sim$ -0.05), although their confidence intervals are quite wide.

The differences in the ratio of synonymous to nonsynonymous variability between the two different breeds is summarized by the $R\_\beta\gamma$ ratio (Figure 3). We observed that the largest deviations from $R\_\beta\gamma$ = 1 are observed when the ratio was calculated based on high-frequency variants ($\alpha_{Fay\&Wu}$). Although the ratio of the two populations is difficult to interpret because of their different underlying demographic histories, some trends can be observed. WB shows an excess of nonsynonymous variants segregating at intermediate frequencies (WB-IB, WB-LW), which might be explained by a past bottleneck that increased deleterious mutations at intermediate frequencies. In addition, the $R\_\beta\gamma$ ratio in IB-LW shows an incremental pattern of this ratio from low to high

**Deleted:** show
**Deleted:** the
**Deleted:** 2
**Deleted:** S8
**Deleted:** less
**Deleted:** (
**Deleted:** in Figure 2A),
**Deleted:** ,
**Deleted:** that
**Deleted:** at
**Deleted:** values
**Deleted:** WB and
**Deleted:** of $\alpha$
**Deleted:** most of the
**Deleted:** , except
**Deleted:** $\alpha_{Fay\&Wu}$ in WB, which is zero. LW is the only breed that shows a linear increase of the $\alpha$ negative values across
**Deleted:** , being even positive when calculated based on high frequencies ($\alpha_{Fay\&Wu}$).
**Deleted:** We did not observe similar patterns of $\alpha$ between domestic breeds compared to WB (Figure 2A).
**Deleted:** 3A
**Deleted:** demographics
**Deleted:** LW-

frequencies, which is compatible with an increase of nonsynonymous beneficial variants on their way to fixation in LW.

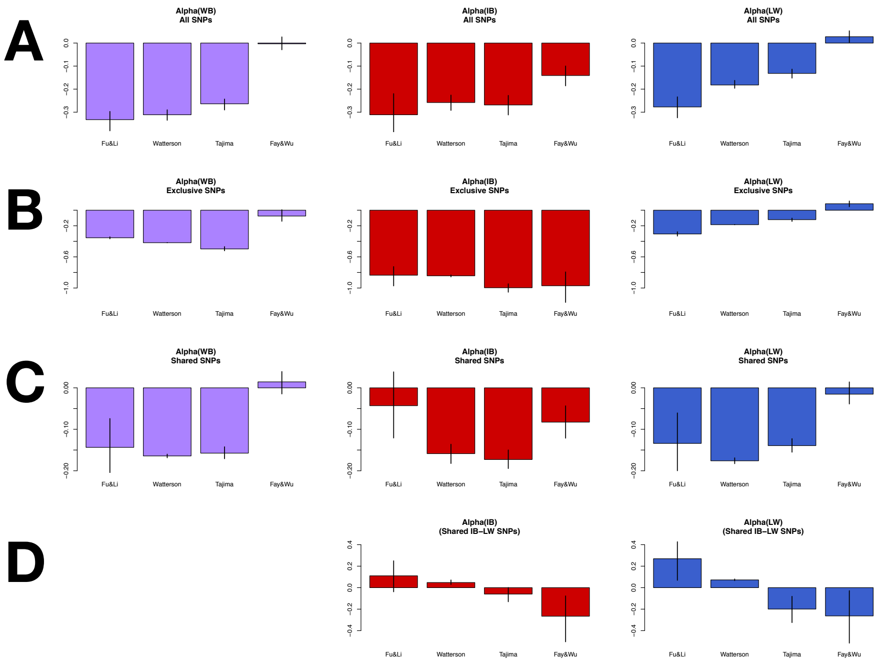


**Figure 2.** Estimates of $\alpha$ for each pig population based on different variability estimators. Total variants (A), exclusive variants (B), shared variants (C) and shared variants between IB and LW (D). Bootstrap intervals at 95% are indicated by a line at each bar. WB; Wild boar; IB, Iberian; LW, Large White.

**Moved (insertion) [1]**

***α*'s and *R_βγ* ratios based on exclusive and shared polymorphisms might reflect changes in selective patterns before and after domestication**

**Deleted:** from

**Deleted:** selection

We observed a high ratio of nonsynonymous to synonymous singletons ($\alpha_{Fu\&Li}$, Figure 2B) when the analysis was performed based on exclusive polymorphisms, suggesting that they have

**Deleted:** exclusive

**Deleted:** ),

deleterious effects in all populations. Nevertheless, the values of $\alpha$ calculated based on intermediate frequency variants ($\alpha_{Tajima}$) in the WB and IB populations are lower than to those based on low-frequency variants, which point to a change in the selective pressure, maintaining nonsynonymous variants at relatively high frequencies. Nevertheless, the $\alpha_{Fay\&Wu}$ values (-0.075, -0.971 and 0.083 for WB, IB and LW, respectively, Table S9) show a similar trend in relation to that based of Total SNPs, that is, close to 0 or positive for WB and LW, but strongly negative for IB. Concordant estimates are observed in the analysis of the SFS based on a reduced number of SNPs (0.155, -0.913 and 0.277 for WB, IB and LW, respectively, Figure 6B), with the difference that a clear positive and not 0 $\alpha$ values is observed in WB. The R_$\beta\gamma$ statistic shows the same pattern as that calculated using all SNPs but with all over one (Figure 3). That indicates that WB has a higher proportion of nonsynonymous polymorphisms compared to IB, in contrast to what is observed when the analysis is performed based on all SNPs. This would suggest a recent change in the constraint of nonsynonymous positions likely at IB breed, as this ratio in IB-LW is also affected. This is also in agreement with the low $\alpha$ value in IB breed at exclusive variants regarding to Total SNPs.

On the other hand, the $\alpha$ values based on shared variants are in general more moderate (closer to zero) than those based on exclusive variants (Figure 2C), likely because shared nonsynonymous polymorphisms are older and hence, expected to be more functionally constrained than the exclusive ones. Additionally, the values of $\alpha$ based on singletons ($\alpha_{Fu\&Li}$) are less negative than those based on intermediate-frequency variants. The $\alpha$ estimates based on shared variants in the analysis of the reduced subset of SNPs are very similar to Total SNPs and very close to zero (Figure 6C). The $R\_\beta\gamma$ statistic for shared variants shows similar patterns than those observed for all variants but with values much closer to 1, indicating a small or moderate selective effect on the shared variants compared to all variants (Figure 3).

When we calculated the $\alpha$ values from shared variants only between the two domestic breeds, we found an inverse pattern regarding to that calculated from all SNPs in each population, with high positive values of $\alpha$ based on low frequencies and very negative values when $\alpha$ is calculated based on high-frequency variants (Figure 2D). This could be due to i) the active elimination of new

**Deleted:** the action of positive selection

**Deleted:** higher

**Deleted:** in the case of WB

**Deleted:** to an attenuated effect of deleterious mutations in IB due to a population size decline. In the case of

**Deleted:** the observed pattern of $\alpha$ is

**Deleted:** calculated with all SNPs (

**Deleted:** 2B). Likewise, the

**Deleted:** a higher magnitude of its value

**Deleted:** 3B).

**Deleted:** or all SNPs

**Deleted:** Again, this pattern might indicate that selection is involve

**Deleted:** increase

**Deleted:** ratio nonsynonymous

**Deleted:** synonymous polymorphisms up

**Deleted:** intermediate frequencies.

**Deleted:** 3C

**Deleted:** Some possible explanations might be

nonsynonymous variants to preserve differences among domestic breeds ($\alpha_{Fu\&Li}$) and ii) the presence of nonsynonymous variants targeted by the process of domestication that shifts them toward high frequencies ($\alpha_{Fay\&wu}$). Nevertheless, we cannot discard that this excess of nonsynonymous variants at high frequencies and the lack of nonsynonymous singletons at low frequency could be due to a more complex and not previously explored demographic scenario.

Deleted: either the effect of the ancestral population structure (wild versus domestic), or

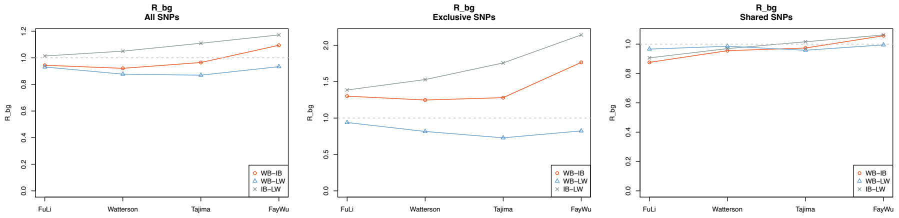


**Figure 3.** Estimates of $R\_\beta\gamma$ for all (left), exclusive (centre) and shared (right) variants. WB; Wild boar; IB, Iberian; LW, Large White.

**Values of $\alpha$ are dependent of the molecular scale but the patterns of the estimated $\alpha$'s are similar across the different molecular scales**

Deleted: Absolute values

Deleted: remain

In addition to the genome-wide analysis, $\alpha$ was calculated using three additional molecular scale levels: i) gene level, ii) genes within windows of 5 Mb, and iii) genes within the same pathway. Figure 4 shows the median of the distributions of the $\alpha$ values for each scale level. When the analysis was performed based on all SNPs, the pattern of $\alpha$ values estimated at the genome-wide level are concordant with those estimated at the gene level, genes within windows and genes within pathways for each breed. However, differences in the value of $\alpha$ within each breed are notorious depending on the scale level examined. The median estimates of $\alpha$ are generally lower at the gene scale level and most of them are very negative, while at the genome-wide scale, the $\alpha$ values are closer to zero. However, the distribution of $\alpha$ values can have a large variance at the gene scale since few variants are used for its estimation. We identified the regions and pathways that showed extreme $\alpha$ values (Table S10 and S11). We found a large number of genes showing $\alpha = 1$ (highest value) because the number of polymorphic nonsynonymous variants per gene was zero. We also

Deleted: estimated

Deleted: In general,

Deleted: distribution

Deleted: pathway level in

Deleted: S9

Deleted: S10

Deleted: having

found a moderately high correlation of $\alpha$ values between breeds (rho ~ 0.7, Pearson correlation when considering pathways) suggesting that in general, these breeds are under similar selective effects. When considering shared and exclusive variants, we generally observed the same pattern, from genes to whole-genome, that is, larger $\alpha$ values at the gene level and closer to zero $\alpha$ values at the larger scale. The differences in $\alpha$ values could be explained because of the distribution of this ratio statistic (*i.e.*, skewed distribution to negative values) and the uneven distribution of the functional variants, in which the mean can be displaced to more negative values (see Materials and Methods).

**Deleted:** using

**Deleted:** For

**Deleted:** Only shared variants of the IB breed exhibited similar $\alpha$ values from genes to the whole-genome scale.

**Deleted:** absolute

**Deleted:** nature

**Deleted:** more

**Deleted:** due to a reduced number of functional variants (*i.e.,* few or null segregating nonsynonymous variants).

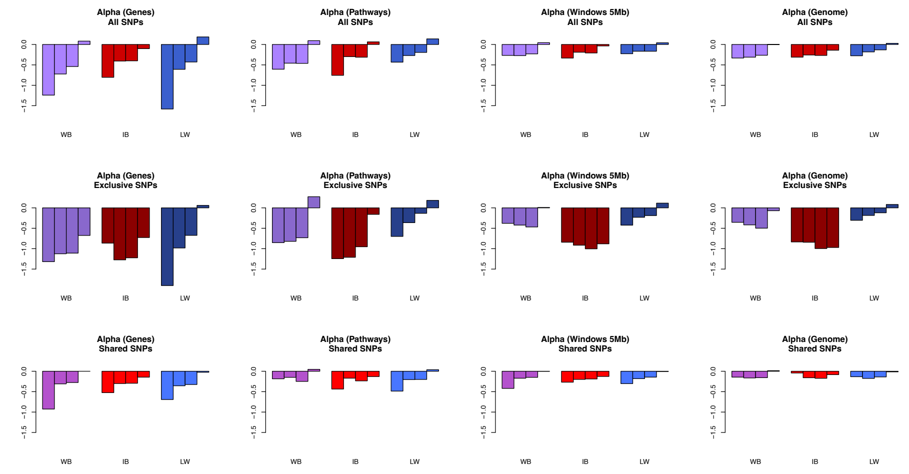


**Figure 4.** Estimates of the median values of $\alpha$ based on different variability estimators and for each pig population at different molecular scales and for all, exclusive and shared variants. Within each population, the order of different $\alpha$'s is: Fu&Li, Watterson, Tajima and Fay&Wu. WB; wild boar; IB, Iberian; LW, Large White.

**Simulated data under different scenarios that include the joint effect of demography and selective events were more concordant to the observed data**

**Deleted: similar**

We used computer simulations to study how the different demographic and selective events occurred during domestication process shaped nucleotide variation present in these populations. We simulated populations mimicking the process of domestication using *SLiM* software (Haller

**Deleted:** affected

**Deleted:** .

and Messer 2017), coupled with several demographic events, including changes of the population size and/or migration. We analyzed the genome-wide patterns of $\alpha$ and the $R\_\beta\gamma$ statistic produced by 63 simulated scenarios that included different demographic events and selective forces acting separately (simple scenarios) or jointly (complex scenarios). The results of the simulation study are summarized in Figures S7-S48. The observed patterns of $\alpha$ based on all variants in the surveyed populations are not compatible with simple scenarios that only consider demographic or positive selection forces (Figure S7). Rather, $\alpha$ patterns from simulated data (irrespective of the magnitude of $\alpha$) fit a scenario with a predominant effect of negative selection (Figure S7). However, the $R\_\beta\gamma$ statistic do not fit any of the simulated simple scenarios (Figure S8). When more complex scenarios were considered (i.e., including a bottleneck, positive/negative selection and/or migration, Figures S9-S14), the general $\alpha$ patterns generated by those scenarios that include both negative and positive selection resembled those observed in WB and LW (with negative $\alpha$'s at low frequency values to slightly positive $\alpha$ values at high frequency). The scenarios that also include some migration events are the ones that showed more concordance for these two breeds (Figures S12, S14). On the other hand, the IB population is more compatible with a scenario without positive selection and with a recent population size reduction (Figure S13). The trends in the $R\_\beta\gamma$ statistic are, in broad strokes, concordant with the conclusions extracted from the comparison between the observed and simulated patterns of $\alpha$ (Figures S15-S20).

The observed patterns of $\alpha$ values based on exclusive variants are similar to those based on total variants but only for the LW population (Figure S21). These patterns cannot be fully explained by any of the complex simulated scenarios that are concordant when considering all variants, although surprisingly, they would be more compatible with those including a population size reduction (Figures S24-S28). The observed $R\_\beta\gamma$'s are compatible with the scenarios that combine both types of selection and a population size reduction (WB-IB) or with scenarios that include migration (WB-LW; Figures S29-S34). Finally, the observed patterns of $\alpha$ and $R\_\beta\gamma$ statistics calculated from shared variants are also compatible with scenarios which includes both types of selection (Figures S35-S48), and being quite compatible with those including expansion demographic events. Overall, the simulations data showed that complex scenarios, including demography, migration, and positive and negative selection, may be necessary to explain the observed data.

- **Deleted:** S5-S46
- **Deleted:** calculated from
- **Deleted:** S5
- **Deleted:** $\alpha$ trends
- **Deleted:** S5
- **Deleted:** S6
- **Deleted:** ,
- **Deleted:** trends of the
- **Deleted:** of $\alpha$
- **Deleted:** with
- **Deleted:** Figures S7-S13). Notice that this is true only for those
- **Deleted:** demographic or
- **Deleted:** S10-
- **Deleted:** would fit
- **Deleted:** S11
- **Deleted:** S13-S18
- **Deleted:** from
- **Deleted:** different from
- **Deleted:** the
- **Deleted:** in WB and IB populations
- **Deleted:** S19
- **Deleted:** proposed
- **Deleted:** except for the IB population, which might
- **Deleted:** scenario with negative selection and
- **Deleted:** S22-S26
- **Deleted:** $\beta\gamma$
- **Deleted:** also
- **Deleted:** combining
- **Deleted:** S27-S32
- **Deleted:** having deleterious plus beneficial mutations
- **Deleted:** S33-S46).

1012

**Table 2.** Posterior Probabilities for each ABC model (multinomial logistic method with tolerance 0.01) and for each pig population for Total variants, Exclusive variants. and Shared variants.

**Posterior probabilities for model comparison (multinomial logistic method)**

| TOTAL | modelA | modelC | modelDN | modelD |
|---|---|---|---|---|
| WB | 0.000 | 0.039 | 0.623 | 0.338 |
| IB | 0.000 | 0.031 | 0.968 | 0.001 |
| LW | 0.000 | 0.195 | 0.526 | 0.279 |
| **EXCLUSIVE** | **modelA** | **modelC** | **modelDN** | **modelD** |
| WB | 0.007 | 0.011 | 0.512 | 0.470 |
| IB | 0.004 | 0.001 | 0.995 | 0.000 |
| LW | 0.001 | 0.111 | 0.113 | 0.775 |
| **SHARED** | **modelA** | **modelC** | **modelDN** | **modelD** |
| WB | 0.001 | 0.334 | 0.405 | 0.260 |
| IB | 0.000 | 0.413 | 0.576 | 0.011 |
| LW | 0.001 | 0.447 | 0.462 | 0.090 |

**Models that assume a discrete distribution of beneficial and deleterious mutations would fit better the observed data**

We used an approximate Bayesian computation (ABC) analysis to infer the DFE separately for each population using the ratios of nonsynonymous to synonymous variants (i.e., polymorphism and divergence) obtained from the whole-genome analysis (see Materials and Methods). Four different models implemented in *polyDFE2* software (Tataru et al. 2019) were tested. These models overcome the inference of the demographic parameters (and others such as linkage effects) by the inclusion of nuisance parameters (see Material and Methods). The four models were: model A, which assumes a gamma distribution of deleterious mutations; model C, which assumes a gamma distribution for deleterious mutations and an exponential distribution for beneficial mutations; model DN, that assumes a discrete distribution of only deleterious and neutral mutations, and model D, that assumes a discrete distribution of deleterious, neutral and beneficial, mutations. Goodness of fit (GoF) analysis revealed that the simulated data under the different models fits differentially to the observed data, although the used range of parameters for priors are compatible with the observed data for all the four models (Table S12). Posterior probabilities

**Deleted:** A model

**Deleted:** assumes

**Deleted:** in the ABC analysis

**Deleted:** Three

**Deleted:** Model

**Deleted:** and model D

**Deleted:** and deleterious mutations. In these models, we included demographic and linkage effects as nuisance parameters (Tataru et al. 2017).

**Deleted:** , which is a measure of the adjustment of the prior chosen models to the data,

**Deleted:** do not always fit well

**Deleted:** (Table S11A), suggesting that the real data fit only to a very restrictive parameter showed that the DN is the most likely model for all three populations when using total number of variants (Table 2). The posterior probability for this model is especially high for the IB breed (0.97). Nevertheless, note that the model D is just below the DN model by less than half of the probability in the case of WB and LW. Finally, the posterior predictive analysis indicated that the observed $\alpha$ values for the three populations are within the range determined by the minimum and maximum simulated $\alpha$ values (i.e., Q1-1.5*IQR, Q3+1.5*IQR, respectively, being IQR the Interquantile Range Q3-Q1) under both models DN and D, although not always inside the Q1 and Q3 quantiles (Figure 5). The mean parameters of the DFE inferred for each population are shown in Table 3 (see also Table S13). The obtained results indicated that the DFE is quite similar among all three populations, which is not entirely surprising because they share a long-term history. According to model DN, and despite there is a lot of uncertainty in the inferred estimates (Table S13), the obtained results show that the DFE contains a large fraction of very deleterious variants, with approximately 75% of the variants being strongly deleterious ($S = -2000$), and with approximately 12% of the variants being neutral or slightly deleterious (approx. 4%). The model D infers a higher proportion of adaptive mutations compared to the neutral ones, although the sum of both is similar to model DN. Finally, the **inferred** contribution of positive selection is relatively low (around 0.7-1.1%).

**Deleted:** each model. This

**Deleted:** pronounced for exclusive variants, where the simulations under the different models fit only marginally to the observed data. Indeed, the model with a wider parameter range versus the real data is usually the discrete

**Deleted:** . However, if we also take into account the values

**Deleted:** posterior probabilities, which is

**Deleted:** probability assigned to each model relative to

**Deleted:** other models of the analysis, we found that the best fit model differed among populations (model C to WB and LW and model D to the IB breed; Table S11A). Finally,

**Deleted:** ) and variability nonsynonymous/synonymous ratios (Figure S47) cannot be obtained using the estimated parameters of any model, although they were closer to model D. The

**Deleted:** S12.

**Deleted:** indicate

**Deleted:** Despite

**Deleted:** ,

**Deleted:** 83

**Deleted:** $\leq -200$; considering the posterior distribution and using the rejection method

**Deleted:** 17

**Deleted:** , weak beneficial and weak

**Deleted:** mutations ($-2 \leq S \leq +2$).

**Deleted:** median

**Deleted:** weak deleterious

**Deleted:** ($\alpha$) estimated from the discrete distribution (by summing weak and strong beneficial proportion of mutations in Model C) is approximately 0.9% (Table S12A).

**The**

**Deleted: DFE is different when based on exclusive and shared variants**

Although the inference of the DFE is going to be distorted by choosing only a subsection

**Deleted:** the variants (e.g., exclusive variants are mostly very recent), we considered that it can give some clues about past events that could be more related to the domestication processes. Regarding to exclusive variants, the simulations under the three different models show a

**Deleted:** fit to the observed data. Indeed, in some cases, the GoF is less than 1% (Table S11B). Although the posterior probability is higher for model C, the posterior predictive simulations show that none of the models can reproduce well the observed data. In general, the posterior predictive simulations under model D yield more similar values to the observed data. However, only the posterior predictive simulations under this model are reasonably similar to the observed data for the LW breed (Figure 6, Figure S48). The results obtained for shared variants are very similar to those considering the total variants, supporting the hypothesis that shared mutations have a predominant effect (compared to exclusive variants) at the whole genome. Finally, the median proportion of adaptive variants ($\alpha$) estimated from the discrete distribution (Model C) gave similar results to the total mutations, but surprisingly estimated a slightly higher proportion for the Iberian breed (Table S12C; Figure S49-S50)....

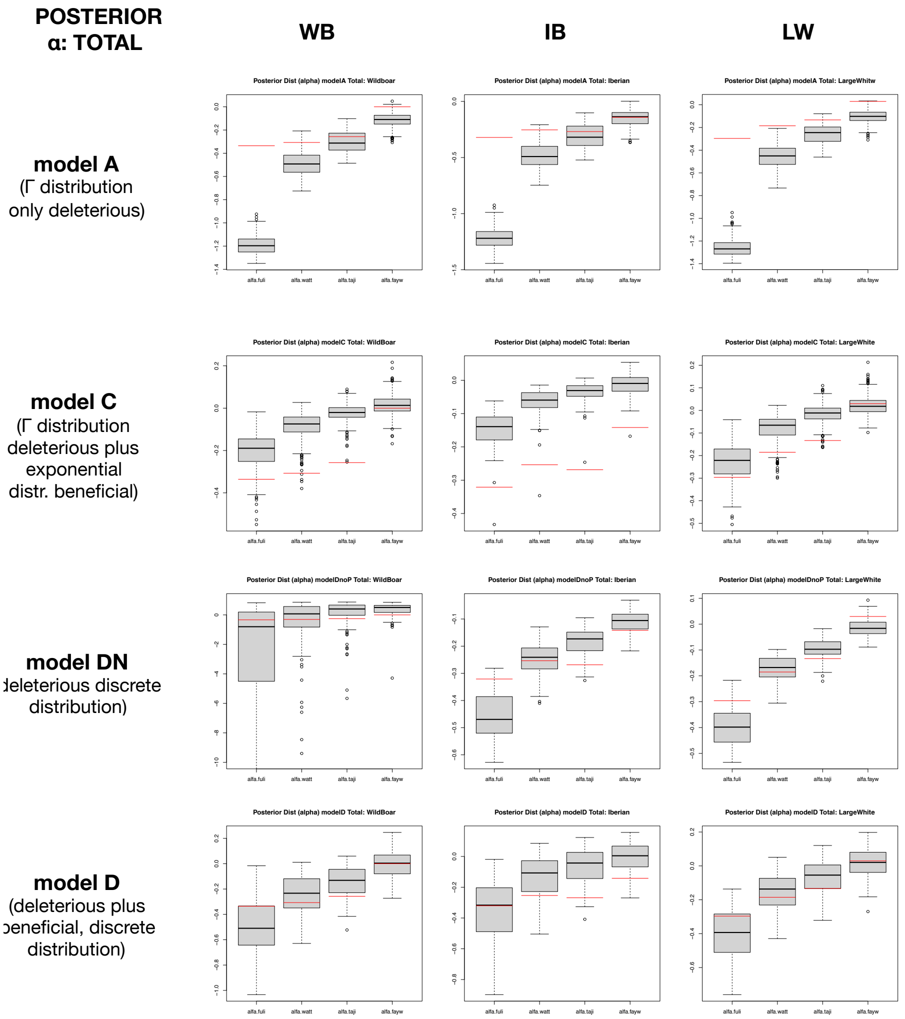


**Figure 5**. Posterior distributions of the $\alpha$ values for total variants based on different variability estimators (Fu&Li, Watterson, Tajima and Fay&Wu). Box plots indicate simulated distributions of $\alpha$ values. Red lines indicate observed $\alpha$ values.

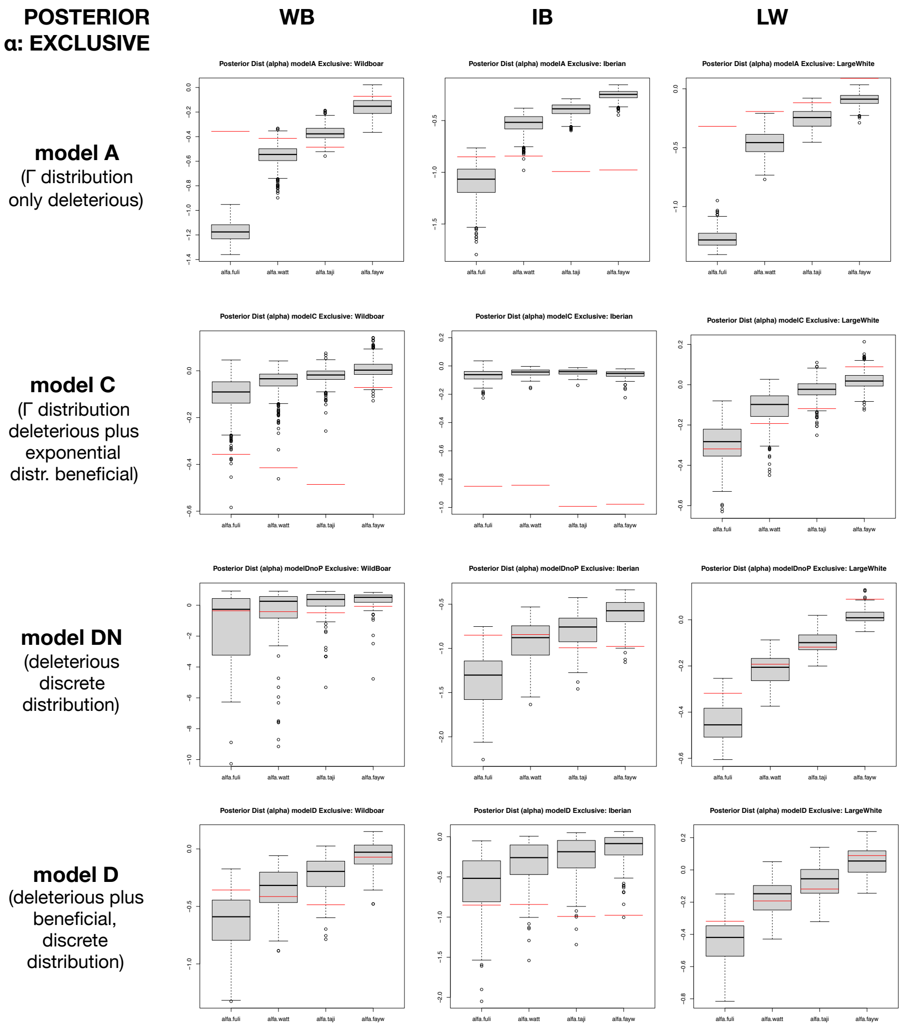


**Figure 6**. Posterior distribution of the $\alpha$ values for exclusive variants based on different variability estimators (Fu&Li, Watterson, Tajima and Fay&Wu). Box plots indicate the simulated distributions of $\alpha$ values. Red lines indicate observed $\alpha$ values.

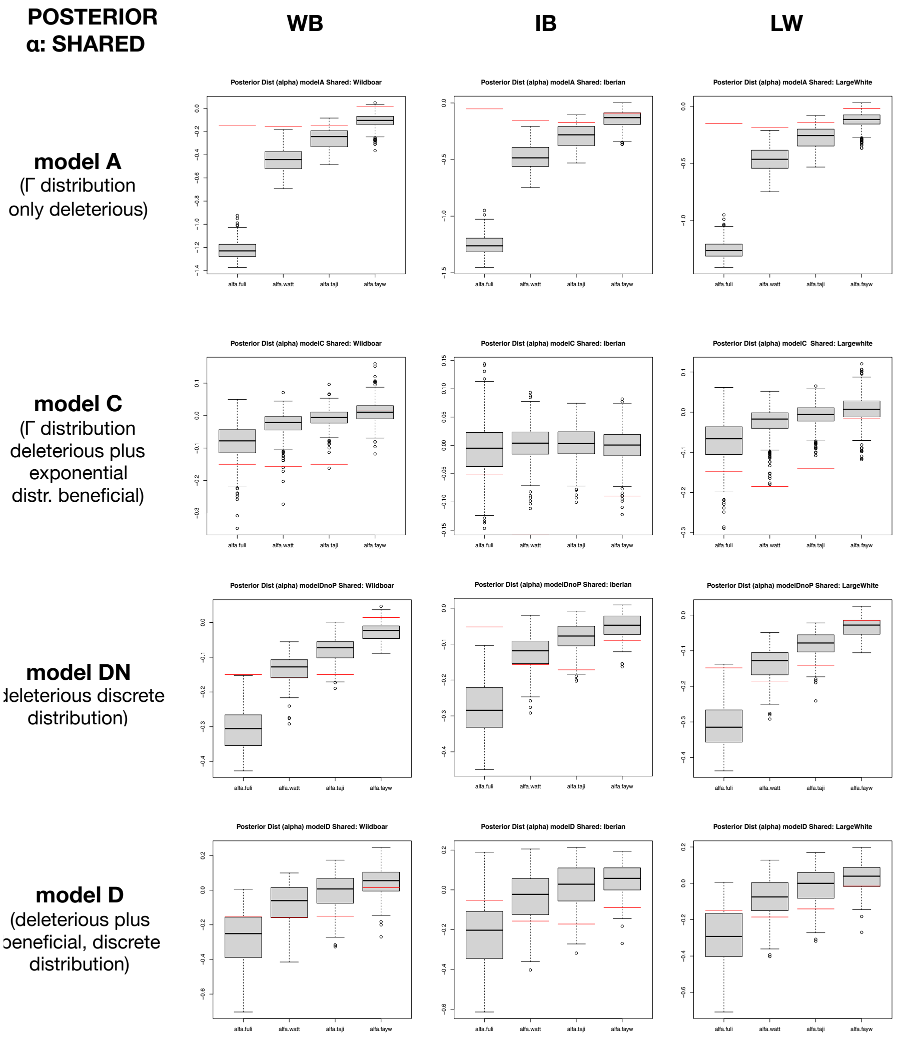


**Figure 7**. Posterior distribution of the $\alpha$ values for shared variants based on different variability estimators (Fu&Li, Watterson, Tajima and Fay&Wu). Box plots indicate the simulated distributions of $\alpha$ values. Red lines indicate observed $\alpha$ values. .

**Table 3.** Inferred selective parameters (weighted mean) for each ABC model and pig population. (A) Total variants. (B) Exclusive polymorphisms. (C) Shared polymorphisms. Note that for model DN and D, the proportion of each discrete S value is relative to the total S values, considering negative inferred values as zero.

**Table 3. Mean Inference of parameters for each of the four analyzed models**

| Mean | model A | | model C | | | | model DN | | | | | model D | | | | | | |
|---|---|---|---|---|---|---|---|---|---|---|---|---|---|---|---|---|---|---|
| **TOTAL** | **Sd** | **b** | **Sd** | **b** | **pb** | **Sb** | **p1 (S=-2000)** | **p2 (S=-200)** | **p3 (S=-20)** | **p4 (S=-2)** | **p5 (S=0)** | **p1 (S=-2000)** | **p2 (S=-200)** | **p3 (S=-20)** | **p4 (S=-2)** | **p5 (S=0)** | **p6 (S=2)** | **p7 (S=20)** |
| **WB** | 24634.19 | 0.179 | 4724.54 | 2.514 | 0.145 | 0.009 | 0.736 | 0.070 | 0.024 | 0.045 | 0.125 | 0.741 | 0.057 | 0.023 | 0.098 | 0.074 | 0.006 | 0.001 |
| **IB** | 27016.47 | 0.180 | 6625.76 | 3.064 | 0.144 | 0.011 | 0.767 | 0.049 | 0.016 | 0.045 | 0.123 | 0.757 | 0.057 | 0.018 | 0.061 | 0.098 | 0.008 | 0.001 |
| **LW** | 20351.15 | 0.189 | 6563.28 | 1.940 | 0.137 | 0.017 | 0.772 | 0.058 | 0.019 | 0.027 | 0.125 | 0.771 | 0.050 | 0.019 | 0.076 | 0.073 | 0.010 | 0.001 |
| **Mean** | **model A** | | **model C** | | | | **model DN** | | | | | **model D** | | | | | | |
| **EXCLUSIVE** | **Sd** | **b** | **Sd** | **b** | **pb** | **Sb** | **p1 (S=-2000)** | **p2 (S=-200)** | **p3 (S=-20)** | **p4 (S=-2)** | **p5 (S=0)** | **p1 (S=-2000)** | **p2 (S=-200)** | **p3 (S=-20)** | **p4 (S=-2)** | **p5 (S=0)** | **p6 (S=2)** | **p7 (S=20)** |
| **WB** | 33447.89 | 0.170 | 3694.78 | 0.821 | 0.153 | 0.002 | 0.723 | 0.055 | 0.018 | 0.091 | 0.113 | 0.709 | 0.064 | 0.020 | 0.135 | 0.065 | 0.006 | 0.001 |
| **IB** | 1834.41 | 0.216 | 253.88 | 44.58 | 0.151 | 0.038 | 0.584 | 0.091 | 0.027 | 0.223 | 0.074 | 0.599 | 0.101 | 0.028 | 0.142 | 0.119 | 0.009 | 0.001 |
| **LW** | 22064.47 | 0.187 | 8311.62 | 1.046 | 0.133 | 0.027 | 0.751 | 0.077 | 0.025 | 0.025 | 0.123 | 0.762 | 0.058 | 0.020 | 0.091 | 0.058 | 0.010 | 0.001 |
| **Mean** | **model A** | | **model C** | | | | **model DN** | | | | | **model D** | | | | | | |
| **SHARED** | **Sd** | **b** | **Sd** | **b** | **pb** | **Sb** | **p1 (S=-2000)** | **p2 (S=-200)** | **p3 (S=-20)** | **p4 (S=-2)** | **p5 (S=0)** | **p1 (S=-2000)** | **p2 (S=-200)** | **p3 (S=-20)** | **p4 (S=-2)** | **p5 (S=0)** | **p6 (S=2)** | **p7 (S=20)** |
| **WB** | 25128.11 | 0.182 | 5223.06 | 5.852 | 0.146 | 0.016 | 0.790 | 0.042 | 0.013 | 0.021 | 0.133 | 0.773 | 0.052 | 0.017 | 0.061 | 0.085 | 0.012 | 0.001 |
| **IB** | 29540.82 | 0.182 | 9042.17 | 11.758 | 0.147 | 0.005 | 0.804 | 0.035 | 0.010 | 0.020 | 0.131 | 0.783 | 0.051 | 0.016 | 0.045 | 0.089 | 0.014 | 0.001 |
| **LW** | 24929.43 | 0.185 | 5169.65 | 5.573 | 0.137 | 0.020 | 0.798 | 0.038 | 0.014 | 0.022 | 0.129 | 0.777 | 0.052 | 0.017 | 0.059 | 0.084 | 0.011 | 0.001 |

Sd: 4Ns mean value for mutations with negative effects. b: shape of the gamma distribution for mutations with negative effect. pb: proportion of beneficial mutations. Sb: 4Ns mean value for mutations with positive effects. p1 (S=-2000): proportion of functional variants having 4Ns=-2000, p2 (S=-200): proportion of functional variants having 4Ns=-200, p3 (S=-20): proportion of functional variants having 4Ns=-20, p4 (S=-2): proportion of functional variants having 4Ns=-2, p5 (S=0): proportion of functional variants having 4Ns=0 (neutral), p6 (S=+2): proportion of functional variants having 4Ns=+2, p7 (S=+20): proportion of functional variants having 4Ns=+20. Nuisance parameters are not shown.

**Deleted:** The
**Deleted:** in animals
**Deleted:** are
**Deleted:** that are
**Deleted:** .
**Deleted:** Here, we
**Deleted:** ).
**Deleted:** allowed us to study and to interpret the effects of the domestication process on genomic variation
**Deleted:** to
**Deleted:** use
**Deleted:** .
**Deleted:** $\alpha$
**Deleted:** The analyses performed here include
**Deleted:** ,

study, including several diverse evolutionary scenarios and the inference of DFE parameters given different selective models.

**Deleted:** for

**Deleted:** ¶

**Deleted:** . Despite

**Deleted:** 2002).

**Deleted:** Selection pressure

**Deleted:** A number of analyses with the aim

**Deleted:** explain

**Deleted:** have been performed already,

**Deleted:** ,

**Deleted:** For instance, Kono et al. (2016) analyzed derived frequencies in domesticated barley populations at different functional classes (deleterious, tolerated) and observed a higher quantity of deleterious variants at low frequencies compared to tolerated or to synonymous mutations. Makino et al. (2018) investigated the ratio of functional to neutral variants at different frequencies in domestic and in wild populations of several animal and plant species, including Asian and European pigs. They observed generally lower levels of synonymous and nonsynonymous variation and a higher ratio of functional to neutral variants in the domestic species compared with their wild counterparts. This ratio was negatively correlated with the frequency of the variants, consistent with a higher number of detrimental variants in domestic populations. The authors claimed that these patterns were compatible with the expected effect of a bottleneck as a consequence of domestication, and with an increase in nonsynonymous variants produced by the lesser efficacy of purifying selection at smaller population sizes, although the presence of positive selection (hitchhiking) was not discarded. However, the opposite pattern was observed in European wild boars and domestic pigs (a higher variability and a lower ratio of nonsynonymous to synonymous in domestic pig populations compared to wild boars). The authors argue that this can be explained by the highly variable patterns produced by bottlenecks, the strong population contraction of European wild boars during the last glaciation and the presence of gene flow between wild and domestic populations.¶
¶

**Deleted:** result

**Deleted:** also

**Deleted:** of

**Deleted:** , that is,

**Deleted:** but

**Deleted:** of

**Deleted:** with Asian pigs;

**Deleted:** increase

**Deleted:** variance

**Deleted:** (see simulations,

**Deleted:** ,

**Deleted:**

**Deleted:** ¶
The

we detected very similar levels of variability between LW and WB, even though we expected to find higher levels of variation in the first due to the documented introgression of Asian germplasm into LW approximately 20–35% of the genome has been estimated to be of Asian origin; Groenen et al. 2012, Bosse et al. 2012, Bosse, Megens, Madsen, et al. 2014, Frantz et al. 2015, Bianco et al. 2015, Ai et al. 2015). Interestingly, the high levels of variability were observed for variants that belonged to different frequency ranges in these two populations: singletons in WB and in high-frequency derived alleles in LW. Although these differences may be mainly due to the effects of gene flow in LW, we cannot discard an important effect of the selective programs applied to this commercial breed.

**Domestication hallmarks at pig coding regions**

Another main goal of this work was finding the hallmarks of positive selection produced as a consequence of the domestication since this process implies a process of positive (human mediated) selection for traits that benefit both humans and the species of interest. The paucity of fixed variants found at coding positions in the three breeds indicates that the observed heritable phenotypic differences among the breeds are either due to: i) very few selective sweeps, ii) positive selection at noncoding functional regions that were not analyzed in here, iii) changes in the frequencies of nonsynonymous variants without being fixed. If the first hypothesis is true, we expect that domestication process should fix the adaptive variants for those genes underlaying the phenotypes of interest. However, we found no fixed variants between domestic breeds and wild pigs. Although this might be a consequence of some genetic exchange among populations, we found that the individuals were classified in groups by their location and according to their respective phenotype (Figure S3A), which suggest that the domestication features of the different breeds, even if admixed, are maintained. When we checked the $\alpha$ values for those genes that were previously reported to show signals of positive selection using other approaches (Groenen 2016), we found that these genes show little or no nonsynonymous polymorphisms or fixed variants (Table S14). This absence of variability is typical from regions under selective sweeps, although not necessarily implicating that these genes are the targets of domestication since there are no variants fixed or close to fixation at their coding regions. We only found significant values of $\alpha$ over zero at the gene KIT in the IB breed, the genes IGF2R and JMJD1C in the LW breed and the

**Deleted:** Currently,
**Deleted:** LW
**Deleted:** (
**Deleted:** Accordingly, the IB breed shows the lowest levels of synonymous and nonsynonymous variation among the breeds studied, probably because of its small effective population size and because the individuals from the IB sample come from a very closed population of pigs. However, we expected to find a higher variability in LW compared to WB due to the process of Asian introgression that this breed has undergone. Surprisingly, we detected very similar levels of variability between them. However
**Deleted:** This difference
**Deleted:** due
**Deleted:** but also in some extend to
**Deleted:** the
**Deleted:** which
**Deleted:** this work
**Deleted:** In
**Deleted:** , although they show the right breed
**Deleted:** . We
**Deleted:** . We
**Deleted:** S13

gene LRRTM3 in the WB population.

**Deleted:** etc

**Deleted:** .

**Deleted:** , the number of fixed shared variants may be significant at the genome level, or alternatively,

**Deleted:** be

**Deleted:** . Hence,

**Deleted:** $\alpha$ values calculated using statistics based on

**Deleted:** variants should be observed

**Deleted:** is

**Deleted:** (~

**Deleted:** 9%)

**Deleted:** all

**Deleted:** is

**Deleted:** . Although speculative, these mutations may change the fate of these populations that are affected by natural or artificial selection.

**Deleted:** ¶
We expected that shared variants between populations would be enriched by selective pressures that predate domestication.

**Deleted:** , these shared polymorphisms can also be the source of

**Deleted:** such that a change

**Deleted:** their

**Deleted:** Furthermore, private variants (those segregating only in one breed)

**Deleted:** nonfunctional positions. We expect that adaptive changes

**Deleted:** increase the ratio of nonsynonymous to synonymous polymorphisms and that this should be reflected as an increase in the negative value of the $\alpha$ statistic. Our

**Deleted:** process indicates

**Deleted:** DFE

**Deleted:** .

**Deleted:** . Finally, the obtained results in the ABC analysis based on total variants show a clear genome-wide effect of the action of purifying selection. We also observed a minor effect of purifying selection in IB and WB when the analysis was performed based on exclusive variants, which suggest a reduction of the population size of these two populations. Nevertheless, we had some difficulties in adjusting the observed data to pertinent DFE models, especially when the analysis was performed based on exclusive variants. Although the

**Deleted:** ,

**Deleted:** contained

**Deleted:** ,

models to explain the real data). In any case, the model that assumes a discrete distribution of deleterious mutations (model DN) seems to generally explain better the observed data, together with the model D (model DN but including the effect of beneficial selection) in a minor degree and also that exclusive variants seemed to be more informative to detect the changes of the selective effects.

### Final remarks

The observed patterns of variability are compatible with the presence of deleterious mutations segregating in all three breeds but also with weak signals of positive selection. In addition, when the variants are split into shared and exclusive, we observed patterns that are in line with the simulated data under different demographic scenarios with the joint action of positive and negative selection. We found a clear effect of deleterious mutations at low-frequency variants and a possible mild effect of positive selection at higher frequencies. However, additional analyses contrasting evolutionary models that consider the effects of standing variation, whose effect change under the domestication process, may shed more light and will help to understand the patterns of variation shaped by the domestication process.

### ACKNOWLEDGMENTS

We acknowledge L. Silió and M. C. Rodríguez for helpful comments on the manuscript. S.E.R.O. is supported by grants AGL2016-78709-R (MEC, Spain) and PID2020-119255GB-I00 (MICINN, Spain), by the CERCA Programme/Generalitat de Catalunya and acknowledges financial support from the Spanish Ministry of Economy and Competitiveness, through the Severo Ochoa Programme for Centres of Excellence in R&D 2016-2019 and 2020-2023 (SEV-2015-0533, CEX2019-000917) and the European Regional Development Fund (ERDF). S.G-R. was supported by a Beatriu de Pinós postdoctoral fellowship (AGAUR; 2014 BP-B 00027).

### CONFLICT OF INTEREST DISCLOSURE

**Deleted:** a
**Deleted:** beneficial and
**Deleted:** D
**Deleted:** the observed data
**Deleted:** . The change of
**Deleted:** DFE when
**Deleted:** analysis was performed based on shared variants is undistinguishable from that based on all variants, indicating
**Deleted:** should
**Deleted:** useful
**Deleted:** effects
**Deleted:** change of
**Deleted:** and
**Deleted:** Nevertheless
**Deleted:** joint
**Deleted:** high
**Deleted:** Additional
**Deleted:** due to
**Deleted:** This work was
**Deleted:** Ministerio de Economia y Competitividad
**Deleted:** AGL2013-41834-R (MEC, Spain),
**Deleted:** . We acknowledge
**Deleted:** “
**Deleted:** ”
**Deleted:** -
**Deleted:** -
**Deleted:** ).
**Deleted:** Beautriu

The authors of this article declare that they have no financial conflict of interest with the content of this article. Sebastian E. Ramos-Onsins is one of the PCIEvolBiol recommenders.

Berkshire (European native pig) provides insights into its origin and domestication." *Sci Rep.* 4:4678. doi: 10.1038/srep04678.

Livingstone, Kevin, and Stephanie Anderson. 2009. “Patterns of Variation in the Evolution of Carotenoid Biosynthetic Pathway Enzymes of Higher Plants.” *Journal of Heredity* 100 (6): 754–61. https://doi.org/10.1093/jhered/esp026.

MacEachern S, McEwan J, McCulloch A, Mather A, Savin K, Goddard M. 2009 "Molecular evolution of the Bovini tribe (Bovidae, Bovinae): is there evidenceof rapid evolution or reduced selective constraint in Domestic cattle?" *BMC Genomics* (10) *179;* https:// doi: 10.1186/1471-2164-10-179.

Makino T, Rubin CJ, Carneiro M, Axelsson E, Andersson L, Webster MT. 2018 "Elevated Proportions of Deleterious Genetic Variation in Domestic Animals and Plants." *Genome Biol Evol.* 10(1):276-290. https://doi: 10.1093/gbe/evy004.

McDonald, J H, and M Kreitman. 1991. “Accelerated Protein Evolution at the Adh Locus in Drosophila.” *Nature* 351: 652–54.

McKenna, Aaron, Matthew Hanna, Eric Banks, Andrey Sivachenko, Kristian Cibulskis, Andrew Kernytsky, Kiran Garimella, et al. 2010. “The Genome Analysis Toolkit: A MapReduce Framework for Analyzing next-Generation DNA Sequencing Data.” *Genome Research* 20 (9): 1297–1303. https://doi.org/10.1101/gr.107524.110.

Messer, Philipp W., and Dmitri A. Petrov. 2013. “Frequent Adaptation and the McDonald-Kreitman Test.” *Proceedings of the National Academy of Sciences of the United States of America* 110 (21): 8615–20. https://doi.org/10.1073/pnas.1220835110.

Montanucci, Ludovica, Hafid Laayouni, Giovanni Marco Dall’Olio, and Jaume Bertranpetit. 2011. “Molecular Evolution and Network-Level Analysis of the N-Glycosylation Metabolic Pathway across Primates.” *Molecular Biology and Evolution* 28 (1): 813–23. https://doi.org/10.1093/molbev/msq259.

Moon, Sunjin, Tae-Hun Kim, Kyung-Tai Lee, Woori Kwak, Taeheon Lee, Si-Woo Lee, Myung-Jick Kim, et al. 2015. “A Genome-Wide Scan for Signatures of Directional Selection in Domesticated Pigs.” *BMC Genomics* 16 (1): 1–12. https://doi.org/10.1186/s12864-015-1330-x.

Nevado, Bruno, Sebastian E. Ramos-Onsins, and Miguel Perez-Enciso. 2014. “Resequencing Studies of Nonmodel Organisms Using Closely Related Reference Genomes: Optimal Experimental Designs and Bioinformatics Approaches for Population Genomics.” *Molecular Ecology* 23 (7): 1764–79. https://doi.org/10.1111/mec.12693.

Orlando L, Librado P. 2019 "Origin and Evolution of Deleterious Mutations in Horses." *Genes* 10, 649; doi:10.3390/genes10090649

Pérez-Enciso, M., G. de los Campos, N. Hudson, J. Kijas, and A. Reverter. 2016. “The ‘Heritability’ of Domestication and Its Functional Partitioning in the Pig.” *Heredity* 118: 160–68. https://doi.org/10.1038/hdy.2016.78.

Quinlan, Aaron R. 2014. “BEDTools: The Swiss-Army Tool for Genome Feature Analysis.” *Current Protocols in Bioinformatics / Editoral Board, Andreas D. Baxevanis ... [et Al.]* 47 (January): 11.12.1-11.12.34. https://doi.org/10.1002/0471250953.bi1112s47.

Ramírez, Oscar, William Burgos-Paz, Encarna Casas, Maria Ballester, Erica Bianco, Iñigo Olalde, Gabriel Santpere, et al. 2014. “Genome Data from a Sixteenth Century Pig Illuminate Modern Breed Relationships.” *Heredity* 114 (2): 175–84. https://doi.org/10.1038/hdy.2014.81.

Ramsay, Heather, Loren H Rieseberg, and Kermit Ritland. 2009. “The Correlation of Evolutionary Rate with Pathway Position in Plant Terpenoid Biosynthesis.” *Molecular Biology and Evolution* 26 (5): 1045–53. https://doi.org/10.1093/molbev/msp021.

Rausher, Mark D, Richard E Miller, and Peter Tiffin. 1999. “Patterns of Evolutionary Rate Variation Among Genes of the Anthocyanin Biosynthetic Pathway.” *Molecular Biology and Evolution* 16 (2): 266–74. https://watermark.silverchair.com/mbev_16_02_0266.pdf?token=AQECAHi208BE49Ooan9kkhW_Ercy7Dm3ZL_9Cf3qfKAc485ysgAAAdwwggHYBgkqhkiG9w0BBwagggHJMIIBxQIBADCCAb4GCSqGSIb3DQEHATAeBglghkgBZQMEAS4wEQQMB_IVBQyX1n-F9I4pAgEQgIIBj8r0BkU_bdeIWEuY_7bUxMFToA8Yo-26snF_yPY.

Renaut, Sebastien, and Loren H. Rieseberg. 2015. “The Accumulation of Deleterious Mutations as a Consequence of Domestication and Improvement in Sunflowers and Other Compositae Crops.” *Molecular Biology and Evolution* 32 (9): 2273–83.

https://doi.org/10.1093/molbev/msv106.

Riley, Rebecca M., WeI Jin, and Greg Gibson. 2003. "Contrasting Selection Pressures on Components of the Ras-Mediated Signal Transduction Pathway in Drosophila." *Molecular Ecology* 12 (5): 1315–23. https://doi.org/10.1046/j.1365-294X.2003.01741.x.

Rubin, Carl-Johan, Hendrik-Jan Megens, Alvaro Martinez Barrio, Khurram Maqbool, Shumaila Sayyab, Doreen Schwochow, Chao Wang, et al. 2012. "Strong Signatures of Selection in the Domestic Pig Genome." *Proceedings of the National Academy of Sciences of the United States of America* 109 (48): 19529–36. https://doi.org/10.1073/pnas.1217149109.

Tajima, Fumio. 1983. "Evolutionary Relationship of DNA Sequences in Finite Populations." *Genetics* 105: 437–60. https://www.ncbi.nlm.nih.gov/pmc/articles/PMC1202167/pdf/437.pdf.

Uricchio LH, Petrov DA, Enard D. 2019 "Exploiting selection at linked sites to infer the rate and strength of adaptation." *Nat Ecol Evol*. 3(6):977-984. doi: 10.1038/s41559-019-0890-6.

Watterson, G.A. 1975. "On the Number of Segregating Sites in Genetical Models without Recombination." *Theoretical Population Biology* 7 (2): 256–76. https://doi.org/10.1016/0040-5809(75)90020-9.

Wilkinson S, Lu ZH, Megens HJ, Archibald AL, Haley C, Jackson IJ, Groenen MA, Crooijmans RP, Ogden R, Wiener P. 2013 "Signatures of diversifying selection in European pig breeds." PLoS Genet. 2013 Apr;9(4):e1003453. doi: 10.1371/journal.pgen.1003453.

Zeder, Melinda A. 2012. "The Domestication of Animals." *Journal of Anthropological Research* 68 (2): 161–90. https://doi.org/10.3998/jar.0521004.0068.201.

### SUPPLEMENTARY MATERIAL

See Supplementary Material file added to see the Tables (S1-S14) and Figures (S1-S48).

**Moved up [1]:** . Total variants (A), exclusive variants (B), shared variants (C) and shared variants between IB and LW (D). Bootstrap intervals at 95% are indicated by a line at each bar.

**Deleted:**
**TABLES**
***Table 1.*** *Number of SNPs (in whole genome, in genes and in coding regions), classified by its presence in each population.*
***Table 2.*** *Combinations of SNPs from coding regions according to its allelic status in each population (A: Ancestral allele, F: Fixed allele, P: Polymorphic allele). SNPs that are missing in any of the populations are not considered in this table.*

**FIGURES**
**Figure 1.** Levels of variation at synonymous (A) and nonsynonymous (B) sites for each pig population and variability estimates and for shared and exclusive variants. WB; wild boar population; IB, Iberian breed; LW, Large White breed.
**Figure 2.** Estimates of $\alpha$ for each pig population

**Moved up [2]:** **Figure 5**.

**Moved up [3]:** Box plots indicate simulated distributions of $\alpha$ values.

**Moved up [4]:** Box plots indicate the simulated distributions of $\alpha$ values.

**Deleted:** WB; wild boar population; IB, Iberian breed; LW, Large White breed.
**Figure 3.** Estimates of R_βγ for each pig population and for all, exclusive and shared variants. WB; wild boar population; IB, Iberian breed; LW, Large White breed.
**Figure 4.** Estimates of the median values of $\alpha$ for each pig population, different molecular scales and for all, exclusive and shared variants. For each population, the order of different $\alpha$'s is: Fu&Li, Watterson, Tajima and Fay&Wu. WB; wild boar population; IB, Iberian breed; LW, Large White breed.

**Deleted:** Posterior distribution of the α values for total variants. Four different estimators of alpha (Fu&Li, Watterson, Tajima and Fay&Wu, see Materials and Methods) are used.

**Deleted:** Red line indicates observed $\alpha$ values.
**Figure 6**. Posterior distribution of the α values for exclusive variants. Four different estimators of alpha (Fu&Li, Watterson, Tajima and Fay&Wu, see Materials and Methods) are used.

**Deleted:** Red line indicates observed $\alpha$ values.

**Deleted:** S13

**Deleted:** S50
