## Supplementary_Material for "Pervasive selection pressure in wild and domestic pigs"

**Table S1.** List of samples.

| Continent | Status | Breed | Origin | Sample | Accession | Depth |
|---|---|---|---|---|---|---|
| Europe | Domestic | IB | Spain | IBGM0327 | SRR1513307 | 13.0 |
| Europe | Domestic | IB | Spain | IBGU1330 | SRR765849 | 7.2 |
| Europe | Domestic | IB | Spain | IBGU1802 | SRX245748 | 12.4 |
| Europe | Domestic | IB | Spain | IBGU1803 | SRR3745079 | 13.0 |
| Europe | Domestic | IB | Spain | IBGU1804 | SRR1917381 | 14.5 |
| Europe | Domestic | IB | Spain | IBGU1805 | SRR5515065 | 12.4 |
| Europe | Domestic | LW | International | LW22F02 | ERR173186 | 10.0 |
| Europe | Domestic | LW | International | LW22F03 | ERR173187 | 10.1 |
| Europe | Domestic | LW | International | LW22F04 | ERR173188 | 10.1 |
| Europe | Domestic | LW | International | LW22F06 | ERR173189 | 9.3 |
| Europe | Domestic | LW | International | LW22F07 | ERR173190 | 11.6 |
| Europe | Domestic | LW | International | LW22M07 | ERR173192 | 10.4 |
| Europe | Domestic | LW | International | LW36F01 | ERR173193 | 9.8 |
| Europe | Domestic | LW | International | LW36F04 | ERR173196 | 9.4 |
| Europe | Domestic | LW | International | LWGB0348 | SRR5513124 | 12.0 |
| Europe | Domestic | LW | International | LWNA3577 | SRR1581108, SRR1581107 | 9.9 |
| Europe | Domestic | LW | International | LWNA3579 | SRR1581111, SRR1581110 | 10.1 |
| Europe | Domestic | LW | International | LWNA3582 | SRR1581121, SRR1581119 | 9.0 |
| Europe | Domestic | LW | International | LWNA3584 | SRR1581128, SRR1581127 | 11.8 |
| Europe | Domestic | LW | International | LWNA3594 | SRR1581137 | 11.8 |
| Europe | Domestic | LW | International | LWNA3595 | SRR1581139 | 12.8 |
| Europe | Domestic | LW | International | LWNA3596 | SRR1581138 | 12.0 |
| Europe | Domestic | LW | International | LWNA3597 | SRR1581140 | 12.5 |
| Europe | Domestic | LW | International | LWNA3599 | SRR1581141 | 12.3 |
| Europe | Domestic | LW | International | LWNA37F01 | ERR977060 | 17.9 |
| Europe | Domestic | LW | International | LWNA38M02 | ERR977062 | 19.6 |
| Europe | Wild | WB | Switzerland | WBCH26M09 | ERR173218 | 14.4 |
| Europe | Wild | WB | Spain | WBES0231 | SRR15698718 | 11.2 |
| Europe | Wild | WB | Spain | WBES0252 | SRR15698980,SRR15686514 | 12.2 |
| Europe | Wild | WB | Spain | WBES0288 | SRR15699754 | 5.2 |
| Europe | Wild | WB | Spain | WBES0291 | SRR15699650 | 11.5 |
| Europe | Wild | WB | Spain | WBES0297 | SRR15699771 | 5.4 |
| Europe | Wild | WB | Spain | WBES0494 | SRR3745077 | 12.6 |
| Europe | Wild | WB | Spain | WBES0717 | SRR1513306 | 13.0 |
| Europe | Wild | WB | France | WBFR25U11 | ERR173217 | 9.4 |
| Europe | Wild | WB | Netherlands | WBNL21M03 | ERR173214 | 11.4 |
| Europe | Wild | WB | Netherlands | WBNL21F04 | ERR977317 | 15.3 |
| Europe | Wild | WB | Netherlands | WBNL22M02 | ERR977342 | 16.6 |
| Europe | Wild | WB | Tunisia | WBTN0965 | SRR3745078 | 12.4 |
| Europe | Wild | WB | Greece | WBGR32F07 | ERR977364 | 10.7 |
| Europe | Wild | WB | Greece | WBGR32U05 | ERR977367 | 10.2 |
| Europe | Wild | WB | Italy | WBIT44U06 | ERR977380 | 13.2 |
| Europe | Wild | WB | Italy | WBIT44U07 | ERR977383 | 11.8 |
| Europe | Wild | WB | Italy | WBIT28F31 | ERR977355 | 17.3 |
| Europe | Wild | WB | Italy | WBIT28M39 | ERR977356 | 12.6 |
| Europe | Wild | WB | Italy | WBIT42M09 | ERR977377 | 13.6 |

**Table S2.** Correlation between nucleotide variation and divergence and the ratio of missing data for a dataset of filtered genes. Genes in the dataset have a proportion of missing data less than 0.3 and variability and divergence values lower than the 99% quantile of the total genes. *P*-values are shown in parenthesis.

| | | **Missing** | |
|---|---|---|---|
| | | **Synonymous** | **Non-Synonymous** |
| IB | Fu&Li | -0.0023 (0.785) | -0.0142 (0.101) |
| | Watterson | -0.0115 (0.180) | -0.0204 (0.017) |
| | Tajima | -0.0087 (0.310) | -0.0178 (0.038) |
| | Fay&Wu | -0.0163 (0.058) | -0.0205 (0.017) |
| | Divergence | -0.0123 (0.154) | -0.0523 (1.2e-09) |
| LW | Fu&Li | -0.0585 (1.1e-10) | -0.0089 (0.327) |
| | Watterson | -0.0228 (0.012) | -0.0131 (0.148) |
| | Tajima | 0.0109 (0.228) | -0.0110 (0.222) |
| | Fay&Wu | -0.0048 (0.593) | -0.0248 (0.006) |
| | Divergence | 0.0125 (0.167) | -0.0203 (0.024) |
| WB | Fu&Li | -0.1851 (<2.2e-16) | -0.1251 (<2.2e-16) |
| | Watterson | -0.1440 (<2.2e-16) | -0.0970 (<2.2e-16) |
| | Tajima | -0.0600 (2.4e-12) | -0.0459 (8.4e-08) |
| | Fay&Wu | -0.0457 (1.0e-07) | -0.0536 (4.1e-10) |
| | Divergence | -0.0369 (1.7e-05) | -0.0515 (1.9e-09) |

**Table S3.** Simulated scenarios and their associated parameters

# A

| Model name | Scenarios | s deleterious mutations | s beneficial mutations | Proportion mutations (deleterious/neutral/beneficial) | Migration Rate: ** Wild to Domestic | Mutation rate | Recombination rate | Total Length | Number of individuals per population | Number of replicates |
|---|---|---|---|---|---|---|---|---|---|---|
| SNM | * Stat/Exp/Red | -1 | 0 | 0.1/0.9/0 | 0 | | | | | |
| NS1 | Stat/Exp/Red | -0.01 | 0 | 0.1/0.9/0 | 0 | 2.50E-07 | 1.17E-08 | 10000bp x 100 loci | 10000 | 100 |
| PS1 | Stat/Exp/Red | -0.1 | 0.0005 | 0.903/0.093/0.093 | 0 | | | | | |

# B

| Model name | Scenario | mean Gamma distrib. (b=0.2) deleterious mutations | mean Exponential distrib. beneficial mutations | Proportion mutations (deleterious/neutral/beneficial) | Migration Rate: ** Wild to Domestic | Mutation rate | Recombination rate | Total Length | Number of individuals per population | Number of replicates |
|---|---|---|---|---|---|---|---|---|---|---|
| SNM_M00 | * Stat/Exp/Red MIG 0 | 0 | 0 | 0/1/0 | 0 | | | | | |
| SNM_M05 | Stat/Exp/Red MIG 0.05 | 0 | 0 | 0/1/0 | 0.05 | | | | | |
| PS010_M00 | Stat/Exp/Red MIG 0 | 0 | 0.01 | 0/0.99/0.01 | 0 | | | | | |
| PS010_M05 | Stat/Exp/Red MIG 0.05 | 0 | 0.01 | 0/0.99/0.01 | 0.05 | | | | | |
| PS001_M00 | Stat/Exp/Red MIG 0 | 0 | 0.001 | 0/0.90/0.10 | 0 | | | | | |
| PS001_M05 | Stat/Exp/Red MIG 0.05 | 0 | 0.001 | 0/0.90/0.10 | 0.05 | | | | | |
| NS030_M00 | Stat/Exp/Red MIG 0 | -0.03 | 0 | 1/0/0 | 0 | | | | | |
| NS030_M05 | Stat/Exp/Red MIG 0.05 | -0.03 | 0 | 1/0/0 | 0.05 | | | | | |
| NS003_M00 | Stat/Exp/Red MIG 0 | -0.003 | 0 | 1/0/0 | 0 | 2.50E-06 | 1.17E-08 | 10000bp x 100 loci | 500 | 100 |
| NS003_M05 | Stat/Exp/Red MIG 0.05 | -0.003 | 0 | 1/0/0 | 0.05 | | | | | |
| PS010_NS030_M00 | Stat/Exp/Red MIG 0 | -0.03 | 0.01 | 0.99/0/0.01 | 0 | | | | | |
| PS010_NS030_M05 | Stat/Exp/Red MIG 0.05 | -0.03 | 0.01 | 0.99/0/0.01 | 0.05 | | | | | |
| PS010_NS003_M00 | Stat/Exp/Red MIG 0 | -0.003 | 0.01 | 0.99/0/0.01 | 0 | | | | | |
| PS010_NS003_M05 | Stat/Exp/Red MIG 0.05 | -0.003 | 0.01 | 0.99/0/0.01 | 0.05 | | | | | |
| PS001_NS030_M00 | Stat/Exp/Red MIG 0 | -0.03 | 0.001 | 0.90/0/0.10 | 0 | | | | | |
| PS001_NS030_M05 | Stat/Exp/Red MIG 0.05 | -0.03 | 0.001 | 0.90/0/0.10 | 0.05 | | | | | |
| PS001_NS003_M00 | Stat/Exp/Red MIG 0 | -0.003 | 0.001 | 0.90/0/0.10 | 0 | | | | | |
| PS001_NS003_M05 | Stat/Exp/Red MIG 0.05 | -0.003 | 0.001 | 0.90/0/0.10 | 0.05 | | | | | |

* Stat: No change in Ne, Exp: 10x Increase Ne, Red: 0.1x Reduction Ne that started 0.4 Ne generations in domestic population before present.
** Mgration started just after the split of Wild with Domestic Populations until present.

**Table S4.** Prior values and distributions for the ABC models using polyDFE.

| Fixed General Parameters | Name Parameter | Value / Distribution | Description |
|---|---|---|---|
| | eps | 0.0001 | probability to misidentify ancestral allele |
| | T | 10 | Divergence time in 4N generations |
| | theta | 0.001 | Population mutation rate (4Nu) |
| **Parameters model A (deleterious)** | | | |
| | Sd | logUniform(100,10000) | mean of the gamma distribution for deleterious mutations (-4Ns) |
| | b | logUniform(0.01,10) | shape of the gamma distribution for deleterious mutations |
| | Smax | 0 | maximum value of 4Ns |
| | r2-13 (nuisance) | logUniform(0.1,10) | distorsion of frequencies 2 to 13 values of the SFS due to demography, linkage, ascertainment bias, non-random sampling, In relation of singletons |
| | r14-26 (nuisance) | logUniform(0.1,10) | distorsion of frequencies 14 to 26 values of the SFS due to demography, etc., in relation of singletons |
| | r27-39 (nuisance) | logUniform(0.1,10) | distorsion of frequencies 27 to 39 values of the SFS due to demography, etc., in relation of singletons |
| | r40 (nuisance) | logUniform(0.1,10) | distorsion of divergence due to demography, etc.. In relation of singletons |
| **Parameters model C (del + beneficial)** | | | |
| | Sd | logUniform(100,10000) | mean of the gamma distribution for deleterious mutations (-4Ns) |
| | b | logUniform(0.01,10) | shape of the gamma distribution for deleterious mutations |
| | pb | Uniform(0,0.25) | proportion of beneficial mutations |
| | Sb | logUniform(0.01,100) | mean of the exponential distribution for beneficial mutations (+4Ns) |
| | r2-13 (nuisance) | logUniform(0.1,10) | distorsion of frequencies 2 to 13 values of the SFS due to demography, linkage, ascertainment bias, non-random sampling, In relation of singletons |
| | r14-26 (nuisance) | logUniform(0.1,10) | distorsion of frequencies 14 to 26 values of the SFS due to demography, etc., in relation of singletons |
| | r27-39 (nuisance) | logUniform(0.1,10) | distorsion of frequencies 27 to 39 values of the SFS due to demography, etc., in relation of singletons |
| | r40 (nuisance) | logUniform(0.1,10) | distorsion of divergence due to demography, etc.. In relation of singletons |
| **Parameters model DN (discrete deleterious)** | | | |
| | p1 | Uniform(0,1) | proportion of mutations with 4Ns=-2000 |
| | p2 | Uniform(0,1-p1) | proportion of mutations with 4Ns=-200 |
| | p3 | Uniform((0,1-p1-p2) | proportion of mutations with 4Ns=-20 |
| | p4 | Uniform(0,1-p1-p2-p3) | proportion of mutations with 4Ns=-2 |
| | p5 | Uniform(0,1-p1-p2-p3-p4) | proportion of mutations with 4Ns= 0 |
| | r2-13 (nuisance) | logUniform(0.1,10) | distorsion of frequencies 2 to 13 values of the SFS due to demography, linkage, ascertainment bias, non-random sampling, In relation of singletons |
| | r14-26 (nuisance) | logUniform(0.1,10) | distorsion of frequencies 14 to 26 values of the SFS due to demography, etc., in relation of singletons |
| | r27-39 (nuisance) | logUniform(0.1,10) | distorsion of frequencies 27 to 39 values of the SFS due to demography, etc., in relation of singletons |
| | r40 (nuisance) | logUniform(0.1,10) | distorsion of divergence due to demography, etc.. In relation of singletons |
| **Parameters model D (discrete del + beneficial)** | | | |
| | p1 | Uniform(0,1) | proportion of mutations with 4Ns=-2000 |
| | p2 | Uniform(0,1-p1) | proportion of mutations with 4Ns=-200 |
| | p3 | Uniform((0,1-p1-p2) | proportion of mutations with 4Ns=-20 |
| | p4 | Uniform(0,1-p1-p2-p3) | proportion of mutations with 4Ns=-2 |
| | p5 | Uniform(0,1-p1-p2-p3-p4) | proportion of mutations with 4Ns= 0 |
| | p6 | Uniform(0,1-p1-p2-p3-p4-p5) | proportion of mutations with 4Ns=+2 |
| | p7 | 1-p1-p2-p3-p4-p5-p6 | proportion of mutations with 4Ns=+20 |
| | r2-13 (nuisance) | logUniform(0.1,10) | distorsion of frequencies 2 to 13 values of the SFS due to demography, linkage, ascertainment bias, non-random sampling, In relation of singletons |
| | r14-26 (nuisance) | logUniform(0.1,10) | distorsion of frequencies 14 to 26 values of the SFS due to demography, etc., in relation of singletons |
| | r27-39 (nuisance) | logUniform(0.1,10) | distorsion of frequencies 27 to 39 values of the SFS due to demography, etc., in relation of singletons |
| | r40 (nuisance) | logUniform(0.1,10) | distorsion of divergence due to demography, etc.. In relation of singletons |

**Table S5.** Confusion matrix to validate Model selection in ABC analyses

**Cross Validation Probabilities for Model Selection**

| **tolerance=0.05** | | modelA | modelC | modelDN | modelD |
|---|---|---|---|---|---|
| | modelA | **69.00** | 7.00 | 24.00 | 0.00 |
| | modelC | 18.00 | **65.00** | 9.00 | 8.00 |
| | modelDN | 13.00 | 3.00 | **83.00** | 1.00 |
| | modelD | 4.00 | 10.00 | 6.00 | **80.00** |

| **tolerance=0.001** | | modelA | modelC | modelDN | modelD |
|---|---|---|---|---|---|
| | modelA | **76.00** | 2.00 | 21.00 | 1.00 |
| | modelC | 16.00 | **69.00** | 9.00 | 6.00 |
| | modelDN | 7.00 | 5.00 | **87.00** | 1.00 |
| | modelD | 1.00 | 12.00 | 16.00 | **71.00** |

**Table S6.** Prediction error for parameter estimation of the ABC models based on a cross-validation sample of size 100

| model A | tolerance | Sd | b | r.2. | r.14. | r.28. |
|---|---|---|---|---|---|---|
| | 0.005 | 0.199 | 0.288 | 0.387 | 0.436 | 0.633 |
| | 0.01 | 0.206 | 0.298 | 0.432 | 0.488 | 0.654 |
| | 0.05 | 0.223 | 0.318 | 0.527 | 0.551 | 0.704 |

| model C | tolerance | Sd | b | pb | Sb | r.2-13 | r.14-26 | r.27-39 |
|---|---|---|---|---|---|---|---|---|
| | 0.005 | 0.702 | 0.649 | 0.449 | 0.142 | 0.817 | 0.851 | 0.932 |
| | 0.010 | 0.704 | 0.656 | 0.458 | 0.143 | 0.833 | 0.858 | 0.957 |
| | 0.050 | 0.703 | 0.669 | 0.484 | 0.153 | 0.895 | 0.880 | 1.005 |

| model DN | tolerance | p1 | p2 | p3 | p4 | p5 | r.2-13 | r.14-26 | r.27-39 |
|---|---|---|---|---|---|---|---|---|---|
| | 0.005 | 0.318 | 1.040 | 0.147 | 0.158 | 0.029 | 0.326 | 0.371 | 0.652 |
| | 0.01 | 0.322 | 1.051 | 0.147 | 0.163 | 0.030 | 0.366 | 0.402 | 0.666 |
| | 0.05 | 0.326 | 1.060 | 0.145 | 0.166 | 0.031 | 0.463 | 0.496 | 0.691 |

| model D | tolerance | p1 | p2 | p3 | p4 | p5 | p6 | p7 | r.2-13 | r.14-26 | r.27-39 |
|---|---|---|---|---|---|---|---|---|---|---|---|
| | 0.005 | 0.195 | 0.516 | 0.099 | 0.315 | 0.583 | 0.320 | 0.002 | 0.407 | 0.601 | 0.785 |
| | 0.01 | 0.198 | 0.523 | 0.100 | 0.309 | 0.580 | 0.347 | 0.002 | 0.439 | 0.619 | 0.791 |
| | 0.05 | 0.204 | 0.542 | 0.108 | 0.295 | 0.574 | 0.438 | 0.003 | 0.529 | 0.695 | 0.812 |

**Table S7.** Number of SNPs present in the whole genome, genes and coding regions and classified according to their presence in pig populations.

| | Number of SNPs | Shared between WB, IB and LW | Shared between WB and IB | Shared between WB and LW | Shared between IB and LW | Exclusive of WB | Exclusive of IB | Exclusive of LW |
|---|---|---|---|---|---|---|---|---|
| **Whole-genome** | 24,869,699 | 7,293,787 | 666,927 | 4,017,107 | 138,378 | 4,239,052 | 385,504 | 8,128,944 |
| **Genes** | 6,684,142 | 1,964,562 | 98,433 | 1,152,555 | 48,351 | 1,138,370 | 100,550 | 2,181,321 |
| **Coding regions** | 149,440 | 18,611 | 3,252 | 25,044 | 1,177 | 51,432 | 3,356 | 46,568 |

**Table S8.** Levels of variability at synonymous and nonsynonymous sites for all breeds for Total, Shared and Exclusive polymorphic variants and for the divergence versus the outgroup.

| | | WB | | IB | | LW | |
|---|---|---|---|---|---|---|---|
| | | Ps | Pn | Ps | Pn | Ps | Pn |
| Total Variants | FuLi | 0.0059 | 0.0011 | 0.0006 | 0.0001 | 0.0038 | 0.0007 |
| | Watterson | 0.0031 | 0.0006 | 0.0011 | 0.0002 | 0.0031 | 0.0005 |
| | Tajima | 0.0023 | 0.0004 | 0.0011 | 0.0002 | 0.0028 | 0.0004 |
| | FayWu | 0.0034 | 0.0005 | 0.0016 | 0.0003 | 0.0043 | 0.0006 |
| Shared Polymorphisms | FuLi | 0.0006 | 0.0001 | 0.0004 | 0.0001 | 0.0006 | 0.0001 |
| | Watterson | 0.0013 | 0.0002 | 0.0009 | 0.0001 | 0.0013 | 0.0002 |
| | Tajima | 0.0016 | 0.0003 | 0.0010 | 0.0002 | 0.0016 | 0.0002 |
| | FayWu | 0.0027 | 0.0004 | 0.0015 | 0.0002 | 0.0024 | 0.0003 |
| Exclusive Polymorphisms | FuLi | 0.0053 | 0.0010 | 0.0002 | 0.0001 | 0.0032 | 0.0006 |
| | Watterson | 0.0018 | 0.0004 | 0.0002 | 0.0000 | 0.0018 | 0.0003 |
| | Tajima | 0.0007 | 0.0002 | 0.0001 | 0.0000 | 0.0012 | 0.0002 |
| | FayWu | 0.0007 | 0.0001 | 0.0001 | 0.0000 | 0.0019 | 0.0002 |
| | | Ds | Dn | Ds | Dn | Ds | Dn |
| Divergence | FuLi | 0.0088 | 0.0012 | 0.0086 | 0.0012 | 0.0085 | 0.0012 |
| | Watterson | 0.0087 | 0.0012 | 0.0085 | 0.0011 | 0.0085 | 0.0011 |
| | Tajima | 0.0087 | 0.0012 | 0.0085 | 0.0011 | 0.0084 | 0.0011 |
| | FayWu | 0.0086 | 0.0012 | 0.0085 | 0.0011 | 0.0084 | 0.0011 |

**Table S9.** Estimates of $\alpha$ for All, Exclusive and Shared variants for each pig population

| | | WB | IB | LW |
|---|---|---|---|---|
| All SNPs | FuLi | -0.332 | -0.311 | -0.277 |
| | Watterson | -0.311 | -0.258 | -0.182 |
| | Tajima | -0.264 | -0.269 | -0.132 |
| | FayWu | -0.003 | -0.141 | 0.028 |
| Exclusive SNPs | FuLi | -0.354 | -0.835 | -0.305 |
| | Watterson | -0.417 | -0.843 | -0.186 |
| | Tajima | -0.498 | -0.997 | -0.121 |
| | FayWu | -0.075 | -0.971 | 0.083 |
| Shared SNPs | FuLi | -0.144 | -0.043 | -0.134 |
| | Watterson | -0.164 | -0.159 | -0.176 |
| | Tajima | -0.157 | -0.173 | -0.139 |
| | FayWu | 0.014 | -0.083 | -0.015 |
| Shared IB-LW SNPs | FuLi | | 0.110 | 0.269 |
| | Watterson | | 0.048 | 0.072 |
| | Tajima | | -0.059 | -0.199 |
| | FayWu | | -0.266 | -0.263 |

**Table S10.** Pathways showing the highest $\alpha_{Fay\&Wu}$ values.

| WB | | IB | | LW | |
|---|---|---|---|---|---|
| **Pathway** | $\alpha_{Fay\&Wu}$ | **Pathway** | $\alpha_{Fay\&Wu}$ | **Pathway** | $\alpha_{Fay\&Wu}$ |
| Maturity onset diabetes of the young | 0.99230 | Maturity onset diabetes of the young | 1.00000 | Phototransduction | 0.91997 |
| Protein export | 0.90743 | Nitrogen metabolism | 1.00000 | Folate biosynthesis | 0.84830 |
| Phototransduction | 0.90399 | Protein export | 1.00000 | Spliceosome | 0.81299 |
| Apelin signaling pathway | 0.82168 | Basal transcription factors | 0.96922 | Thyroid cancer | 0.81254 |
| Basal transcription factors | 0.81404 | Hypertrophic cardiomyopathy | 0.96097 | mRNA surveillance pathway | 0.78237 |
| Carbohydrate digestion and absorption | 0.78207 | Beta-Alanine metabolism | 0.86393 | Morphine addiction | 0.75427 |
| GnRH signaling pathway | 0.77080 | Carbohydrate digestion and absorption | 0.85869 | Antifolate resistance | 0.72263 |
| Circadian rhythm | 0.74982 | Mitophagy-animal | 0.85324 | Basal transcription factors | 0.71137 |
| Mineral absorption | 0.73443 | MAPK (JNK) signaling | 0.83623 | Choline metabolism in cancer | 0.60802 |
| Amyotrophic lateral sclerosis | 0.72434 | Viral myocarditis | 0.80584 | Arginine and proline metabolism | 0.58645 |

**Table S11.** Analysis of the genomic windows of 5-Mb showing the highest $\alpha_{\mathrm{Fay\&Wu}}$ values.

| WB | Window (Chromosome:Initial-Final Positions | alpha(Fay&Wu) |
|---|---|---|
| | 15:10000001-15000000 | 1.00000 |
| | 1:170000001-175000000 | 1.00000 |
| | 15:45000001-50000000 | 0.99734 |
| | 11:65000001-70000000 | 0.98494 |
| | 1:215000001-220000000 | 0.96279 |
| | 15:100000001-105000000 | 0.96238 |
| | 10:1-5000000 | 0.95963 |
| | 16:15000001-20000000 | 0.95849 |
| | 11:30000001-35000000 | 0.95648 |
| | 13:190000001-195000000 | 0.94697 |
| | 1:50000001-55000000 | 0.94079 |
| | 11:55000001-60000000 | 0.92279 |
| | 1:190000001-195000000 | 0.91743 |
| | 16:65000001-70000000 | 0.91263 |
| | 5:110000001-111506441 | 0.91231 |
| | 13:45000001-50000000 | 0.91105 |
| | 16:80000001-85000000 | 0.90642 |
| | 16:10000001-15000000 | 0.90341 |
| | 7:110000001-115000000 | 0.90178 |
| | 6:5000001-10000000 | 0.89058 |
| | 13:65000001-70000000 | 0.87955 |
| | 4:65000001-70000000 | 0.85453 |
| | 4:25000001-30000000 | 0.84057 |
| | 9:85000001-90000000 | 0.82182 |
| | 16:1-5000000 | 0.81841 |
| | 6:20000001-25000000 | 0.81645 |
| | 2:35000001-40000000 | 0.80054 |
| | 1:195000001-200000000 | 0.79284 |
| | 12:30000001-35000000 | 0.78398 |
| | 10:35000001-40000000 | 0.78328 |

| IB | Window (Chromosome:Initial-Final Positions | alpha(Fay&Wu) |
|---|---|---|
| | 9:150000001-153670197 | 1.00000 |
| | 8:20000001-25000000 | 1.00000 |
| | 7:110000001-115000000 | 1.00000 |
| | 4:55000001-60000000 | 1.00000 |
| | 2:95000001-100000000 | 1.00000 |
| | 2:115000001-120000000 | 1.00000 |
| | 16:85000001-86898991 | 1.00000 |
| | 16:15000001-20000000 | 1.00000 |
| | 15:80000001-85000000 | 1.00000 |
| | 15:5000001-10000000 | 1.00000 |
| | 15:45000001-50000000 | 1.00000 |
| | 15:30000001-35000000 | 1.00000 |
| | 15:155000001-157681621 | 1.00000 |
| | 15:10000001-15000000 | 1.00000 |
| | 15:100000001-105000000 | 1.00000 |
| | 13:65000001-70000000 | 1.00000 |
| | 11:60000001-65000000 | 1.00000 |
| | 11:30000001-35000000 | 1.00000 |
| | 10:1-5000000 | 1.00000 |
| | 16:1-5000000 | 0.99712 |
| | 14:125000001-130000000 | 0.99439 |
| | 1:240000001-245000000 | 0.99190 |
| | 16:65000001-70000000 | 0.99184 |
| | 10:60000001-65000000 | 0.98889 |
| | 3:140000001-144787322 | 0.97741 |
| | 16:40000001-45000000 | 0.96543 |
| | 1:215000001-220000000 | 0.95732 |
| | 1:40000001-45000000 | 0.95056 |
| | 8:35000001-40000000 | 0.94947 |
| | 11:80000001-85000000 | 0.94127 |

| LW | Window (Chromosome:Initial-Final Positions) | alpha(Fay&Wu) |
|---|---|---|
| | 16:10000001-15000000 | 1.00000 |
| | 15:45000001-50000000 | 0.99930 |
| | 11:30000001-35000000 | 0.98920 |
| | 15:100000001-105000000 | 0.97879 |
| | 1:170000001-175000000 | 0.95827 |
| | 14:125000001-130000000 | 0.95233 |
| | 1:65000001-70000000 | 0.95093 |
| | 14:150000001-153851969 | 0.94730 |
| | 13:85000001-90000000 | 0.94534 |
| | 15:10000001-15000000 | 0.92277 |
| | 7:110000001-115000000 | 0.91288 |
| | 11:55000001-60000000 | 0.90170 |
| | 9:85000001-90000000 | 0.89437 |
| | 13:65000001-70000000 | 0.89346 |
| | 1:145000001-150000000 | 0.86119 |
| | 15:95000001-100000000 | 0.86061 |
| | 2:130000001-135000000 | 0.82900 |
| | 18:20000001-25000000 | 0.82542 |
| | 3:140000001-144787322 | 0.82330 |
| | 16:15000001-20000000 | 0.80676 |
| | 15:50000001-55000000 | 0.80520 |
| | 14:85000001-90000000 | 0.80366 |
| | 13:190000001-195000000 | 0.79835 |
| | 1:315000001-315321322 | 0.78167 |
| | 1:130000001-135000000 | 0.77286 |
| | 6:95000001-100000000 | 0.77138 |
| | 18:60000001-61220071 | 0.76378 |
| | 13:40000001-45000000 | 0.76357 |
| | 5:85000001-90000000 | 0.74643 |
| | 6:30000001-35000000 | 0.72954 |

**Table S12.** Goodness of fit probabilities for each ABC model and for each pig population for Total, Exclusive and Shared variants.

| TOTAL | modelA | modelC | modelDN | modelD |
|---|---|---|---|---|
| WB | 0.970 | 0.570 | 0.170 | 0.620 |
| IB | 0.940 | 0.500 | 0.610 | 0.210 |
| LW | 0.950 | 0.540 | 0.760 | 0.370 |

| EXCLUSIVE | modelA | modelC | modelDN | modelD |
|---|---|---|---|---|
| WB | 0.950 | 0.600 | 0.210 | 0.420 |
| IB | 0.840 | 0.720 | 0.160 | 0.300 |
| LW | 0.960 | 0.540 | 0.410 | 0.730 |

| SHARED | modelA | modelC | modelDN | modelD |
|---|---|---|---|---|
| WB | 0.940 | 0.550 | 0.250 | 0.680 |
| IB | 0.910 | 0.510 | 0.280 | 0.630 |
| LW | 0.940 | 0.600 | 0.360 | 0.730 |

**Table S13.** Inferred selective parameters for each ABC model and pig population (breed). Sd: mean 4Ns value for mutations with negative effects. b: shape of the gamma distribution for mutations with negative effect. pb: proportion of beneficial mutations. Sb: mean 4Ns value for mutations with positive effects. p1 (S=-2000): proportion of functional variants having 4Ns=-2000, p2 (S=-200): proportion of functional variants having 4Ns=-200, p3 (S=-20): proportion of functional variants having 4Ns=-20, p4 (S=-2): proportion of functional variants having 4Ns=-2, p5 (S=0): proportion of functional variants having 4Ns=0 (neutral), p6 (S=+2): proportion of functional variants having 4Ns=+2, p7 (S=+20): proportion of functional variants having 4Ns=+20. Nuisance parameters are not shown. (A) Total variants. (B) Exclusive variants. (C) Shared variants. For models A and C, we used the local regression method and tolerance=0.0005. For models DN and D, we used lower tolerance (0.0001) and the rejection method, to limit the parameter estimates proportions between 0 and 1.

A

**TOTAL VARIANTS: Posterior distributions for each parameter and breed**

| WB mode lA | | | WB model C | | | | |
|---|---|---|---|---|---|---|---|
| | Sd | b | | Sd | b | pb | Sb |
| Min. | 24416.1 | 0.18 | Min. | 1514.6 | 0.422 | 0.141 | 0.002 |
| Weighted 2.5% | 24513.7 | 0.18 | Weighted 2.5% | 1746.2 | 0.585 | 0.143 | 0.002 |
| Weighted Median | 24643.7 | 0.18 | Weighted Median | 3722.4 | 1.603 | 0.145 | 0.007 |
| Weighted Mean | 24634.2 | 0.18 | Weighted Mean | 4724.5 | 2.514 | 0.145 | 0.009 |
| Weighted Mode | 24666.9 | 0.18 | Weighted Mode | 2813.2 | 1.098 | 0.146 | 0.004 |
| Weighted 97.5% | 24709.3 | 0.18 | Weighted 97.5% | 11669.6 | 7.425 | 0.146 | 0.026 |
| Max. | 24724.0 | 0.18 | Max. | 14514.5 | 12.903 | 0.147 | 0.057 |

| WB model DN | | | | | | WB model D | | | | | | | |
|---|---|---|---|---|---|---|---|---|---|---|---|---|---|
| | p1 (S=-2000) | p2 (S=-200) | p3 (S=-20) | p4 (S=-2) | p5 (S=0) | | p1 (S=-2000) | p2 (S=-200) | p3 (S=-20) | p4 (S=-2) | p5 (S=0) | p6 (S=2) | p7 (S=20) |
| Min.: | 0.559 | 0.000 | 0.001 | 0.001 | 0.100 | Min.: | 0.564 | 0.003 | 0.000 | 0.003 | 0.001 | 0.000 | 0.000 |
| 2.5%Perc.: | 0.618 | 0.002 | 0.002 | 0.004 | 0.105 | 2.5%Perc.: | 0.621 | 0.005 | 0.001 | 0.014 | 0.003 | 0.000 | 0.000 |
| Median: | 0.747 | 0.058 | 0.020 | 0.044 | 0.125 | Median: | 0.756 | 0.035 | 0.021 | 0.095 | 0.081 | 0.003 | 0.001 |
| Mean: | 0.736 | 0.070 | 0.024 | 0.045 | 0.125 | Mean: | 0.741 | 0.057 | 0.023 | 0.098 | 0.074 | 0.006 | 0.001 |
| Mode: | 0.762 | 0.026 | 0.010 | 0.041 | 0.123 | Mode: | 0.763 | 0.024 | 0.009 | 0.091 | 0.095 | 0.002 | 0.000 |
| 97.5%Perc.: | 0.804 | 0.190 | 0.067 | 0.082 | 0.145 | 97.5%Perc.: | 0.798 | 0.181 | 0.061 | 0.193 | 0.138 | 0.026 | 0.003 |
| Max.: | 0.818 | 0.270 | 0.084 | 0.099 | 0.151 | Max.: | 0.828 | 0.276 | 0.084 | 0.199 | 0.144 | 0.038 | 0.004 |

| IB model A | | | IB model C | | | | |
|---|---|---|---|---|---|---|---|
| | Sd | b | | Sd | b | pb | Sb |
| Min. | 26778.0 | 0.18 | Min. | 690.3 | 1.616 | 0.136 | 0.009 |
| Weighted 2.5% | 26876.1 | 0.18 | Weighted 2.5% | 1151.4 | 1.899 | 0.139 | 0.010 |
| Weighted Median | 27028.0 | 0.18 | Weighted Median | 4081.7 | 2.954 | 0.144 | 0.011 |
| Weighted Mean | 27016.5 | 0.18 | Weighted Mean | 6625.8 | 3.064 | 0.144 | 0.011 |
| Weighted Mode | 27062.6 | 0.18 | Weighted Mode | 2800.6 | 2.527 | 0.143 | 0.011 |
| Weighted 97.5% | 27104.0 | 0.18 | Weighted 97.5% | 22538.1 | 4.838 | 0.147 | 0.012 |
| Max. | 27110.3 | 0.18 | Max. | 33708.0 | 8.298 | 0.147 | 0.014 |

| IB model DN | | | | | | IB model D | | | | | | | |
|---|---|---|---|---|---|---|---|---|---|---|---|---|---|
| | p1 (S=-2000) | p2 (S=-200) | p3 (S=-20) | p4 (S=-2) | p5 (S=0) | | p1 (S=-2000) | p2 (S=-200) | p3 (S=-20) | p4 (S=-2) | p5 (S=0) | p6 (S=2) | p7 (S=20) |
| Min.: | 0.663 | 0.001 | 0.000 | 0.008 | 0.099 | Min.: | 0.595 | 0.001 | 0.000 | 0.002 | 0.001 | 0.000 | 0.000 |
| 2.5%Perc.: | 0.675 | 0.003 | 0.001 | 0.011 | 0.106 | 2.5%Perc.: | 0.646 | 0.004 | 0.001 | 0.004 | 0.007 | 0.000 | 0.000 |
| Median: | 0.776 | 0.038 | 0.014 | 0.044 | 0.123 | Median: | 0.767 | 0.038 | 0.015 | 0.052 | 0.103 | 0.003 | 0.001 |
| Mean: | 0.767 | 0.049 | 0.016 | 0.045 | 0.123 | Mean: | 0.757 | 0.057 | 0.018 | 0.061 | 0.098 | 0.008 | 0.001 |
| Mode: | 0.788 | 0.022 | 0.006 | 0.045 | 0.122 | Mode: | 0.769 | 0.023 | 0.007 | 0.033 | 0.118 | 0.002 | 0.000 |
| 97.5%Perc.: | 0.817 | 0.140 | 0.047 | 0.077 | 0.138 | 97.5%Perc.: | 0.827 | 0.165 | 0.049 | 0.152 | 0.150 | 0.031 | 0.003 |
| Max.: | 0.827 | 0.167 | 0.060 | 0.093 | 0.141 | Max.: | 0.833 | 0.238 | 0.060 | 0.187 | 0.157 | 0.038 | 0.003 |

| LW model A | | | LW model C | | | | |
|---|---|---|---|---|---|---|---|
| | Sd | b | | Sd | b | pb | Sb |
| Min. | 20175.3 | 0.19 | Min. | 700.6 | 0.174 | 0.122 | 0.005 |
| Weighted 2.5% | 20234.6 | 0.19 | Weighted 2.5% | 1379.6 | 0.340 | 0.130 | 0.008 |
| Weighted Median | 20358.6 | 0.19 | Weighted Median | 4025.4 | 0.979 | 0.137 | 0.015 |
| Weighted Mean | 20351.1 | 0.19 | Weighted Mean | 6563.3 | 1.940 | 0.137 | 0.017 |
| Weighted Mode | 20394.4 | 0.19 | Weighted Mode | 2555.2 | 0.648 | 0.138 | 0.012 |
| Weighted 97.5% | 20412.3 | 0.19 | Weighted 97.5% | 25361.1 | 7.251 | 0.141 | 0.035 |
| Max. | 20419.5 | 0.19 | Max. | 47559.9 | 12.189 | 0.143 | 0.059 |

| LW model DN | | | | | | LW model D | | | | | | | |
|---|---|---|---|---|---|---|---|---|---|---|---|---|---|
| | p1 (S=-2000) | p2 (S=-200) | p3 (S=-20) | p4 (S=-2) | p5 (S=0) | | p1 (S=-2000) | p2 (S=-200) | p3 (S=-20) | p4 (S=-2) | p5 (S=0) | p6 (S=2) | p7 (S=20) |
| Min.: | 0.558 | 0.001 | 0.000 | 0.000 | 0.101 | Min.: | 0.609 | 0.000 | 0.000 | 0.001 | 0.000 | 0.000 | 0.000 |
| 2.5%Perc.: | 0.637 | 0.001 | 0.001 | 0.002 | 0.110 | 2.5%Perc.: | 0.683 | 0.002 | 0.000 | 0.009 | 0.003 | 0.000 | 0.000 |
| Median: | 0.785 | 0.043 | 0.017 | 0.024 | 0.125 | Median: | 0.776 | 0.038 | 0.014 | 0.076 | 0.077 | 0.004 | 0.001 |
| Mean: | 0.772 | 0.058 | 0.019 | 0.027 | 0.125 | Mean: | 0.771 | 0.050 | 0.019 | 0.076 | 0.073 | 0.010 | 0.001 |
| Mode: | 0.799 | 0.022 | 0.015 | 0.022 | 0.129 | Mode: | 0.779 | 0.025 | 0.007 | 0.084 | 0.096 | 0.002 | 0.000 |
| 97.5%Perc.: | 0.833 | 0.215 | 0.050 | 0.061 | 0.138 | 97.5%Perc.: | 0.832 | 0.164 | 0.057 | 0.149 | 0.131 | 0.036 | 0.003 |
| Max.: | 0.844 | 0.299 | 0.073 | 0.084 | 0.139 | Max.: | 0.839 | 0.246 | 0.067 | 0.173 | 0.139 | 0.043 | 0.004 |

B

**EXCLUSIVE VARIANTS: Posterior distributions for each parameter and breed**

| WB mode lA | | | WB model C | | | | |
|---|---|---|---|---|---|---|---|
| | Sd | b | | Sd | b | pb | Sb |
| Min. | 32109.8 | 0.17 | Min. | 1986.5 | 0.289 | 0.152 | 0.001 |
| Weighted 2.5% | 32756.3 | 0.17 | Weighted 2.5% | 2435.8 | 0.337 | 0.153 | 0.001 |
| Weighted Median | 33473.0 | 0.17 | Weighted Median | 3563.1 | 0.762 | 0.153 | 0.002 |
| Weighted Mean | 33447.9 | 0.17 | Weighted Mean | 3694.8 | 0.821 | 0.153 | 0.002 |
| Weighted Mode | 33726.1 | 0.17 | Weighted Mode | 2978.2 | 0.457 | 0.153 | 0.001 |
| Weighted 97.5% | 33931.8 | 0.17 | Weighted 97.5% | 5467.8 | 1.585 | 0.154 | 0.004 |
| Max. | 34057.8 | 0.17 | Max. | 6187.9 | 2.073 | 0.154 | 0.006 |

| WB model DN | | | | | | WB model D | | | | | | | |
|---|---|---|---|---|---|---|---|---|---|---|---|---|---|
| | p1 (S=-2000) | p2 (S=-200) | p3 (S=-20) | p4 (S=-2) | p5 (S=0) | | p1 (S=-2000) | p2 (S=-200) | p3 (S=-20) | p4 (S=-2) | p5 (S=0) | p6 (S=2) | p7 (S=20) |
| Min.: | 0.538 | 0.000 | 0.000 | 0.019 | 0.078 | Min.: | 0.574 | 0.003 | 0.000 | 0.013 | 0.000 | 0.000 | 0.000 |
| 2.5%Perc.: | 0.586 | 0.001 | 0.001 | 0.045 | 0.092 | 2.5%Perc.: | 0.617 | 0.004 | 0.000 | 0.031 | 0.004 | 0.000 | 0.000 |
| Median: | 0.736 | 0.040 | 0.015 | 0.091 | 0.114 | Median: | 0.713 | 0.052 | 0.016 | 0.146 | 0.059 | 0.003 | 0.001 |
| Mean: | 0.723 | 0.055 | 0.018 | 0.091 | 0.113 | Mean: | 0.709 | 0.064 | 0.020 | 0.135 | 0.065 | 0.006 | 0.001 |
| Mode: | 0.750 | 0.016 | 0.009 | 0.083 | 0.117 | Mode: | 0.744 | 0.030 | 0.008 | 0.158 | 0.041 | 0.002 | 0.001 |
| 97.5%Perc.: | 0.786 | 0.198 | 0.059 | 0.130 | 0.131 | 97.5%Perc.: | 0.775 | 0.177 | 0.062 | 0.222 | 0.154 | 0.030 | 0.004 |
| Max.: | 0.799 | 0.259 | 0.075 | 0.147 | 0.141 | Max.: | 0.807 | 0.212 | 0.093 | 0.239 | 0.159 | 0.041 | 0.004 |

| IB model A | | | IB model C | | | | |
|---|---|---|---|---|---|---|---|
| | Sd | b | | Sd | b | pb | Sb |
| Min. | 511.1 | 0.21 | Min. | 196.0 | 35.092 | 0.150 | 0.026 |
| Weighted 2.5% | 1004.5 | 0.21 | Weighted 2.5% | 213.6 | 36.337 | 0.150 | 0.026 |
| Weighted Median | 1683.4 | 0.22 | Weighted Median | 254.6 | 44.307 | 0.151 | 0.033 |
| Weighted Mean | 1834.4 | 0.22 | Weighted Mean | 253.9 | 44.577 | 0.151 | 0.038 |
| Weighted Mode | 1491.0 | 0.22 | Weighted Mode | 266.0 | 47.910 | 0.151 | 0.030 |
| Weighted 97.5% | 3440.9 | 0.22 | Weighted 97.5% | 290.2 | 54.864 | 0.151 | 0.070 |
| Max. | 5041.9 | 0.22 | Max. | 300.3 | 63.558 | 0.151 | 0.138 |

| IB model DN | | | | | | IB model D | | | | | | | |
|---|---|---|---|---|---|---|---|---|---|---|---|---|---|
| | p1 (S=-2000) | p2 (S=-200) | p3 (S=-20) | p4 (S=-2) | p5 (S=0) | | p1 (S=-2000) | p2 (S=-200) | p3 (S=-20) | p4 (S=-2) | p5 (S=0) | p6 (S=2) | p7 (S=20) |
| Min.: | 0.298 | 0.000 | 0.001 | 0.115 | 0.011 | Min.: | 0.351 | 0.003 | 0.000 | 0.003 | 0.000 | 0.000 | 0.000 |
| 2.5%Perc.: | 0.406 | 0.002 | 0.002 | 0.148 | 0.029 | 2.5%Perc.: | 0.399 | 0.004 | 0.001 | 0.006 | 0.003 | 0.000 | 0.000 |
| Median: | 0.601 | 0.066 | 0.020 | 0.220 | 0.075 | Median: | 0.614 | 0.088 | 0.026 | 0.129 | 0.125 | 0.004 | 0.001 |
| Mean: | 0.584 | 0.091 | 0.027 | 0.223 | 0.074 | Mean: | 0.599 | 0.101 | 0.028 | 0.142 | 0.119 | 0.009 | 0.001 |
| Mode: | 0.628 | 0.049 | 0.013 | 0.203 | 0.079 | Mode: | 0.635 | 0.030 | 0.029 | 0.067 | 0.174 | 0.002 | 0.000 |
| 97.5%Perc.: | 0.682 | 0.269 | 0.081 | 0.301 | 0.121 | 97.5%Perc.: | 0.710 | 0.286 | 0.074 | 0.302 | 0.230 | 0.047 | 0.003 |
| Max.: | 0.692 | 0.396 | 0.123 | 0.317 | 0.129 | Max.: | 0.727 | 0.379 | 0.100 | 0.322 | 0.251 | 0.061 | 0.004 |

| LW model A | | | LW model C | | | | |
|---|---|---|---|---|---|---|---|
| | Sd | b | | Sd | b | pb | Sb |
| Min. | 21184.4 | 0.19 | Min. | 840.2 | 0.230 | 0.104 | 0.006 |
| Weighted 2.5% | 21719.0 | 0.19 | Weighted 2.5% | 1377.1 | 0.347 | 0.119 | 0.008 |
| Weighted Median | 22087.1 | 0.19 | Weighted Median | 4573.4 | 0.678 | 0.134 | 0.021 |
| Weighted Mean | 22064.5 | 0.19 | Weighted Mean | 8311.6 | 1.046 | 0.133 | 0.027 |
| Weighted Mode | 22182.6 | 0.19 | Weighted Mode | 2801.2 | 0.536 | 0.135 | 0.015 |
| Weighted 97.5% | 22250.3 | 0.19 | Weighted 97.5% | 32854.0 | 2.988 | 0.140 | 0.075 |
| Max. | 22291.5 | 0.19 | Max. | 82204.2 | 6.814 | 0.142 | 0.120 |

| LW model DN | | | | | | LW model D | | | | | | | |
|---|---|---|---|---|---|---|---|---|---|---|---|---|---|
| | p1 (S=-2000) | p2 (S=-200) | p3 (S=-20) | p4 (S=-2) | p5 (S=0) | | p1 (S=-2000) | p2 (S=-200) | p3 (S=-20) | p4 (S=-2) | p5 (S=0) | p6 (S=2) | p7 (S=20) |
| Min.: | 0.515 | 0.001 | 0.000 | 0.000 | 0.097 | Min.: | 0.602 | 0.000 | 0.000 | 0.012 | 0.000 | 0.000 | 0.000 |
| 2.5%Perc.: | 0.592 | 0.002 | 0.002 | 0.001 | 0.106 | 2.5%Perc.: | 0.655 | 0.001 | 0.001 | 0.021 | 0.001 | 0.000 | 0.000 |
| Median: | 0.765 | 0.054 | 0.021 | 0.021 | 0.124 | Median: | 0.765 | 0.046 | 0.014 | 0.092 | 0.051 | 0.004 | 0.001 |
| Mean: | 0.751 | 0.077 | 0.025 | 0.025 | 0.123 | Mean: | 0.762 | 0.058 | 0.020 | 0.091 | 0.058 | 0.010 | 0.001 |
| Mode: | 0.779 | 0.040 | 0.016 | 0.012 | 0.125 | Mode: | 0.762 | 0.030 | 0.009 | 0.089 | 0.039 | 0.003 | 0.001 |
| 97.5%Perc.: | 0.827 | 0.250 | 0.068 | 0.061 | 0.135 | 97.5%Perc.: | 0.830 | 0.186 | 0.067 | 0.157 | 0.121 | 0.037 | 0.004 |
| Max.: | 0.832 | 0.334 | 0.081 | 0.082 | 0.139 | Max.: | 0.839 | 0.246 | 0.095 | 0.173 | 0.128 | 0.043 | 0.004 |

C

**SHARED VARIANTS: Posterior distributions for each parameter and breed**

| WB mode lA | | | WB model C | | | | |
|---|---|---|---|---|---|---|---|
| | Sd | b | | Sd | b | pb | Sb |
| Min. | 25100.9 | 0.18 | Min. | 1133.1 | 0.380 | 0.085 | 0.008 |
| Weighted 2.5% | 25111.5 | 0.18 | Weighted 2.5% | 1658.7 | 0.858 | 0.126 | 0.009 |
| Weighted Median | 25129.2 | 0.18 | Weighted Median | 4833.2 | 4.261 | 0.147 | 0.015 |
| Weighted Mean | 25128.1 | 0.18 | Weighted Mean | 5223.1 | 5.852 | 0.146 | 0.016 |
| Weighted Mode | 25134.5 | 0.18 | Weighted Mode | 3409.8 | 2.521 | 0.149 | 0.012 |
| Weighted 97.5% | 25137.3 | 0.18 | Weighted 97.5% | 11028.5 | 15.676 | 0.160 | 0.027 |
| Max. | 25138.9 | 0.18 | Max. | 13798.7 | 21.200 | 0.170 | 0.081 |

| WB model DN | | | | | | WB model D | | | | | | | |
|---|---|---|---|---|---|---|---|---|---|---|---|---|---|
| | p1 (S=-2000) | p2 (S=-200) | p3 (S=-20) | p4 (S=-2) | p5 (S=0) | | p1 (S=-2000) | p2 (S=-200) | p3 (S=-20) | p4 (S=-2) | p5 (S=0) | p6 (S=2) | p7 (S=20) |
| Min.: | 0.682 | 0.001 | 0.000 | 0.000 | 0.110 | Min.: | 0.608 | 0.001 | 0.000 | 0.004 | 0.000 | 0.000 | 0.000 |
| 2.5%Perc.: | 0.726 | 0.001 | 0.000 | 0.002 | 0.118 | 2.5%Perc.: | 0.680 | 0.003 | 0.001 | 0.005 | 0.012 | 0.000 | 0.000 |
| Median: | 0.796 | 0.035 | 0.011 | 0.020 | 0.133 | Median: | 0.775 | 0.038 | 0.015 | 0.058 | 0.093 | 0.006 | 0.001 |
| Mean: | 0.790 | 0.042 | 0.013 | 0.021 | 0.133 | Mean: | 0.773 | 0.052 | 0.017 | 0.061 | 0.085 | 0.012 | 0.001 |
| Mode: | 0.810 | 0.021 | 0.005 | 0.022 | 0.132 | Mode: | 0.774 | 0.024 | 0.006 | 0.030 | 0.102 | 0.002 | 0.000 |
| 97.5%Perc.: | 0.836 | 0.116 | 0.041 | 0.052 | 0.145 | 97.5%Perc.: | 0.833 | 0.152 | 0.049 | 0.139 | 0.138 | 0.038 | 0.003 |
| Max.: | 0.839 | 0.155 | 0.063 | 0.066 | 0.147 | Max.: | 0.840 | 0.246 | 0.060 | 0.152 | 0.145 | 0.055 | 0.004 |

| IB model A | | | IB model C | | | | |
|---|---|---|---|---|---|---|---|
| | Sd | b | | Sd | b | pb | Sb |
| Min. | 29518.6 | 0.18 | Min. | 538.2 | 4.021 | 0.102 | 0.003 |
| Weighted 2.5% | 29528.5 | 0.18 | Weighted 2.5% | 1563.5 | 6.283 | 0.134 | 0.004 |
| Weighted Median | 29541.7 | 0.18 | Weighted Median | 8388.8 | 11.719 | 0.148 | 0.005 |
| Weighted Mean | 29540.8 | 0.18 | Weighted Mean | 9042.2 | 11.758 | 0.147 | 0.005 |
| Weighted Mode | 29545.5 | 0.18 | Weighted Mode | 4177.3 | 11.227 | 0.149 | 0.005 |
| Weighted 97.5% | 29547.9 | 0.18 | Weighted 97.5% | 20166.1 | 17.189 | 0.154 | 0.007 |
| Max. | 29549.1 | 0.18 | Max. | 26451.4 | 19.856 | 0.158 | 0.009 |

| IB model DN | | | | | | IB model D | | | | | | | |
|---|---|---|---|---|---|---|---|---|---|---|---|---|---|
| | p1 (S=-2000) | p2 (S=-200) | p3 (S=-20) | p4 (S=-2) | p5 (S=0) | | p1 (S=-2000) | p2 (S=-200) | p3 (S=-20) | p4 (S=-2) | p5 (S=0) | p6 (S=2) | p7 (S=20) |
| Min.: | 0.731 | 0.001 | 0.000 | 0.001 | 0.110 | Min.: | 0.672 | 0.001 | 0.000 | 0.000 | 0.006 | 0.000 | 0.000 |
| 2.5%Perc.: | 0.738 | 0.002 | 0.000 | 0.002 | 0.117 | 2.5%Perc.: | 0.711 | 0.002 | 0.001 | 0.000 | 0.015 | 0.000 | 0.000 |
| Median: | 0.807 | 0.028 | 0.007 | 0.019 | 0.131 | Median: | 0.783 | 0.047 | 0.012 | 0.036 | 0.096 | 0.009 | 0.001 |
| Mean: | 0.804 | 0.035 | 0.010 | 0.020 | 0.131 | Mean: | 0.783 | 0.051 | 0.016 | 0.045 | 0.089 | 0.014 | 0.001 |
| Mode: | 0.811 | 0.015 | 0.004 | 0.020 | 0.131 | Mode: | 0.793 | 0.019 | 0.006 | 0.023 | 0.105 | 0.003 | 0.000 |
| 97.5%Perc.: | 0.844 | 0.112 | 0.036 | 0.044 | 0.144 | 97.5%Perc.: | 0.838 | 0.136 | 0.043 | 0.119 | 0.141 | 0.050 | 0.002 |
| Max.: | 0.851 | 0.119 | 0.045 | 0.066 | 0.147 | Max.: | 0.849 | 0.192 | 0.050 | 0.139 | 0.151 | 0.055 | 0.002 |

| LW model A | | | LW model C | | | | |
|---|---|---|---|---|---|---|---|
| | Sd | b | | Sd | b | pb | Sb |
| Min. | 24896.0 | 0.19 | Min. | 247.7 | 0.318 | 0.075 | 0.007 |
| Weighted 2.5% | 24908.1 | 0.19 | Weighted 2.5% | 638.1 | 0.867 | 0.115 | 0.010 |
| Weighted Median | 24931.0 | 0.19 | Weighted Median | 3691.9 | 4.706 | 0.139 | 0.018 |
| Weighted Mean | 24929.4 | 0.19 | Weighted Mean | 5169.6 | 5.573 | 0.137 | 0.020 |
| Weighted Mode | 24937.8 | 0.19 | Weighted Mode | 2347.6 | 2.190 | 0.142 | 0.014 |
| Weighted 97.5% | 24941.3 | 0.19 | Weighted 97.5% | 15849.4 | 13.612 | 0.152 | 0.041 |
| Max. | 24943.0 | 0.19 | Max. | 24797.2 | 17.044 | 0.161 | 0.073 |

| LW model DN | | | | | | LW model D | | | | | | | |
|---|---|---|---|---|---|---|---|---|---|---|---|---|---|
| | p1 (S=-2000) | p2 (S=-200) | p3 (S=-20) | p4 (S=-2) | p5 (S=0) | | p1 (S=-2000) | p2 (S=-200) | p3 (S=-20) | p4 (S=-2) | p5 (S=0) | p6 (S=2) | p7 (S=20) |
| Min.: | 0.707 | 0.001 | 0.000 | 0.001 | 0.110 | Min.: | 0.609 | 0.001 | 0.000 | 0.000 | 0.000 | 0.000 | 0.000 |
| 2.5%Perc.: | 0.739 | 0.002 | 0.000 | 0.002 | 0.115 | 2.5%Perc.: | 0.700 | 0.002 | 0.000 | 0.002 | 0.012 | 0.000 | 0.000 |
| Median: | 0.800 | 0.032 | 0.011 | 0.020 | 0.130 | Median: | 0.780 | 0.039 | 0.015 | 0.053 | 0.092 | 0.005 | 0.001 |
| Mean: | 0.798 | 0.038 | 0.014 | 0.022 | 0.129 | Mean: | 0.777 | 0.052 | 0.017 | 0.059 | 0.084 | 0.011 | 0.001 |
| Mode: | 0.801 | 0.014 | 0.005 | 0.018 | 0.130 | Mode: | 0.781 | 0.024 | 0.005 | 0.027 | 0.106 | 0.002 | 0.000 |
| 97.5%Perc.: | 0.841 | 0.110 | 0.041 | 0.045 | 0.138 | 97.5%Perc.: | 0.835 | 0.143 | 0.048 | 0.136 | 0.137 | 0.035 | 0.003 |
| Max.: | 0.851 | 0.142 | 0.045 | 0.066 | 0.141 | Max.: | 0.840 | 0.246 | 0.059 | 0.145 | 0.142 | 0.038 | 0.004 |

**Table S14.** Estimates of variation and divergence at synonymous and nonsynonymous sites and $\alpha$ values for those genes showing hallmarks of positive selection in previous studies for each pig population. (A) IB, (B) LW, (C) WB.

# A

| | | IB | | | | | | | | | | | | | |
|---|---|---|---|---|---|---|---|---|---|---|---|---|---|---|---|
| Gene | Function | Theta(Watterson) | | Theta(Tajima) | | Theta(Fu&Li) | | Theta(Fay&Wu) | | Divergence | | alpha(Fu&Li) | alpha(Tajima) | alpha(Watterson) | alpha(Fay&Wu) |
| | | Syn | nonSyn | Syn | nonSyn | Syn | nonSyn | Syn | nonSyn | Syn | nonSyn | | | | |
| NR6A1 | Body size | 0.00000 | 0.000443 | 0.000000 | 0.000251 | 0.000000 | 0.000000 | 0.000000 | 0.002255 | 0.012661 | 0.001128 | NA | -Inf | -Inf | -Inf |
| PLAG1 | Body size | 0.00000 | 0.000416 | 0.000000 | 0.000579 | 0.000000 | 0.000000 | 0.000000 | 0.000347 | 0.003497 | 0.000405 | NA | -Inf | -Inf | -Inf |
| LCORL | Body size | 0.00000 | 0.000000 | 0.000000 | 0.000000 | 0.000000 | 0.000000 | 0.000000 | 0.000000 | 0.000000 | 0.000000 | NA | NA | NA | NA |
| NR6A1 | Body size | 0.00000 | 0.000000 | 0.000000 | 0.000000 | 0.000000 | 0.000000 | 0.000000 | 0.000000 | 0.014874 | 0.000000 | NA | NA | NA | NA |
| KIT | Coat color | 0.00545 | 0.000325 | NA | NA | 0.000000 | 0.000460 | 0.007103 | 0.000174 | 0.011254 | 0.000690 | -Inf | NA | 0.027045 | 0.600922 |
| EDNRB | Coat color (Asian origin) | 0.00000 | 0.000000 | 0.000000 | 0.000000 | 0.000000 | 0.000000 | 0.000000 | 0.000000 | 0.003798 | 0.000000 | NA | NA | NA | NA |
| LRRTM4 | Behavior | 0.00000 | 0.000000 | 0.000000 | 0.000000 | 0.000000 | 0.000000 | 0.000000 | 0.000000 | 0.003528 | 0.000000 | NA | NA | NA | NA |
| LRRTM3 | Behavior | 0.00000 | 0.000000 | 0.000000 | 0.000000 | 0.000000 | 0.000000 | 0.000000 | 0.000000 | 0.002635 | 0.000000 | NA | NA | NA | NA |
| LRRTM1 | Behavior | NA | NA | NA | NA | NA | NA | NA | NA | NA | NA | NA | NA | NA | NA |
| PPP1R1B | Behavior | 0.00000 | 0.000000 | 0.000000 | 0.000000 | 0.000000 | 0.000000 | 0.000000 | 0.000000 | 0.000000 | 0.002932 | NA | NA | NA | NA |
| LEMD3 | Ear morphology | 0.00000 | 0.000000 | 0.000000 | 0.000000 | 0.000000 | 0.000000 | 0.000000 | 0.000000 | 0.000000 | 0.000000 | NA | NA | NA | NA |
| IGF2R | Lean growth | 0.00000 | 0.000000 | 0.000000 | 0.000000 | 0.000000 | 0.000000 | 0.000000 | 0.000000 | 0.012634 | 0.000647 | NA | NA | NA | NA |
| JMJD1C | Fertility | 0.00042 | 0.000246 | 0.000299 | 0.000212 | 0.000637 | 0.000180 | 0.000048 | 0.000046 | 0.001433 | 0.000117 | -2.461538 | -7.702232 | -6.156669 | -10.584534 |
| OSTN | Body composition | 0.00000 | 0.001173 | 0.000000 | 0.000664 | 0.000000 | 0.000000 | 0.000000 | 0.005974 | 0.000000 | 0.002987 | NA | NA | NA | NA |
| AHR | Litter size | 0.00113 | 0.000639 | 0.000567 | 0.000342 | 0.003401 | 0.001866 | 0.000052 | 0.000035 | 0.017291 | 0.002037 | -3.656489 | -4.122132 | -3.813611 | -4.742860 |

# B

| | | LW | | | | | | | | | | | | | |
|---|---|---|---|---|---|---|---|---|---|---|---|---|---|---|---|
| Gene | Function | Theta(Watterson) | | Theta(Tajima) | | Theta(Fu&Li) | | Theta(Fay&Wu) | | Divergence | | alpha(Fu&Li) | alpha(Tajima) | alpha(Watterson) | alpha(Fay&Wu) |
| | | Syn | nonSyn | Syn | nonSyn | Syn | nonSyn | Syn | nonSyn | Syn | nonSyn | | | | |
| NR6A1 | Body size | 0.001032 | 0.000601 | 0.000248 | 0.000401 | 0.004220 | 0.001253 | 0.000008 | 0.000060 | 0.012785 | 0.001478 | -1.568578 | -12.988761 | -4.037667 | -68.056291 |
| PLAG1 | Body size | 0.000000 | 0.000000 | 0.000000 | 0.000000 | 0.000000 | 0.000000 | 0.000000 | 0.000000 | 0.003497 | 0.000000 | NA | NA | NA | NA |
| LCORL | Body size | 0.000000 | 0.000000 | 0.000000 | 0.000000 | 0.000000 | 0.000000 | 0.000000 | 0.000000 | 0.000000 | 0.000000 | NA | NA | NA | NA |
| NR6A1 | Body size | 0.000000 | 0.001001 | 0.000000 | 0.000213 | 0.000000 | 0.004259 | 0.000000 | 0.000005 | 0.014874 | 0.000106 | -Inf | -Inf | -Inf | -Inf |
| KIT | Coat color | NA | NA | NA | NA | NA | NA | NA | NA | NA | NA | NA | NA | NA | NA |
| EDNRB | Coat color (Asian origin) | 0.000904 | 0.000273 | 0.000389 | 0.000058 | 0.000000 | 0.001162 | 0.000022 | 0.000001 | 0.003998 | 0.000029 | -Inf | -19.555605 | -40.591005 | -8.486638 |
| LRRTM4 | Behavior | 0.000000 | 0.000000 | 0.000000 | 0.000000 | 0.000000 | 0.000000 | 0.000000 | 0.000000 | 0.003528 | 0.000000 | NA | NA | NA | NA |
| LRRTM3 | Behavior | 0.000644 | 0.000000 | 0.000888 | 0.000000 | 0.000000 | 0.000000 | 0.003424 | 0.000000 | 0.002092 | 0.000000 | NA | NA | NA | NA |
| LRRTM1 | Behavior | NA | NA | NA | NA | NA | NA | NA | NA | NA | NA | NA | NA | NA | NA |
| PPP1R1B | Behavior | 0.000000 | 0.000698 | 0.000000 | 0.000801 | 0.000000 | 0.000000 | 0.000000 | 0.004271 | 0.000000 | 0.002469 | NA | NA | NA | NA |
| LEMD3 | Ear morphology | 0.000000 | 0.000000 | 0.000000 | 0.000000 | 0.000000 | 0.000000 | 0.000000 | 0.000000 | 0.000000 | 0.000000 | NA | NA | NA | NA |
| IGF2R | Lean growth | 0.005394 | 0.000346 | 0.003579 | 0.000148 | 0.004211 | 0.000647 | 0.010479 | 0.000009 | 0.013107 | 0.000723 | -1.787132 | 0.249789 | -0.162648 | 0.983804 |
| JMJD1C | Fertility | 0.004155 | 0.000698 | 0.002357 | 0.000511 | 0.007642 | 0.001079 | 0.001372 | 0.000122 | 0.002450 | 0.000308 | -0.123017 | -0.725981 | -0.336555 | 0.289882 |
| OSTN | Body composition | 0.000000 | 0.001570 | 0.000000 | 0.000341 | 0.000000 | 0.003319 | 0.000000 | 0.006467 | 0.000000 | 0.003315 | NA | NA | NA | NA |
| AHR | Litter size | 0.003939 | 0.001312 | 0.005898 | 0.001745 | 0.000000 | 0.000000 | 0.001790 | 0.002227 | 0.003736 | 0.001930 | NA | 0.427388 | 0.355550 | -1.407711 |

# C

| | | WB | | | | | | | | | | | | | |
|---|---|---|---|---|---|---|---|---|---|---|---|---|---|---|---|
| Gene | Function | Theta(Watterson) | | Theta(Tajima) | | Theta(Fu&Li) | | Theta(Fay&Wu) | | Divergence | | alpha(Fu&Li) | alpha(Tajima) | alpha(Watterson) | alpha(Fay&Wu) |
| | | Syn | nonSyn | Syn | nonSyn | Syn | nonSyn | Syn | nonSyn | Syn | nonSyn | | | | |
| NR6A1 | Body size | 0.005978 | 0.000601 | 0.002219 | 0.000270 | 0.021101 | 0.001253 | 0.000244 | 0.000022 | 0.013860 | 0.000142 | -4.800699 | -10.903500 | -8.822119 | -7.699543 |
| PLAG1 | Body size | 0.000000 | 0.000268 | 0.000000 | 0.000538 | 0.000000 | 0.000000 | 0.000000 | 0.000368 | 0.003497 | 0.000439 | NA | -Inf | -Inf | -Inf |
| LCORL | Body size | 0.002560 | 0.000368 | 0.000573 | 0.000078 | 0.010717 | 0.001564 | 0.000016 | 0.000002 | 0.000286 | 0.000039 | -0.068710 | 0.000005 | -0.051917 | 0.067810 |
| NR6A1 | Body size | 0.000000 | 0.000000 | 0.000000 | 0.000000 | 0.000000 | 0.000000 | 0.000000 | 0.000000 | 0.014874 | 0.000000 | NA | NA | NA | NA |
| KIT | Coat color | 0.004817 | 0.000449 | 0.004233 | 0.000386 | 0.003087 | 0.000000 | 0.011303 | 0.000894 | 0.010647 | 0.000621 | 1.000000 | -0.562920 | -0.596490 | -0.355432 |
| EDNRB | Coat color (Asian origin) | 0.000904 | 0.000000 | 0.000200 | 0.000000 | 0.003798 | 0.000000 | 0.000005 | 0.000000 | 0.003898 | 0.000000 | NA | NA | NA | NA |
| LRRTM4 | Behavior | 0.004200 | 0.000000 | 0.000930 | 0.000000 | 0.017641 | 0.000000 | 0.000025 | 0.000000 | 0.003993 | 0.000000 | NA | NA | NA | NA |
| LRRTM3 | Behavior | 0.000627 | 0.000392 | 0.000139 | 0.000135 | 0.000000 | 0.000812 | 0.005131 | 0.000007 | 0.002566 | 0.000069 | -Inf | -35.225275 | -22.188312 | 0.949770 |
| LRRTM1 | Behavior | 0.000000 | 0.000000 | 0.000000 | 0.000000 | 0.000000 | 0.000000 | 0.000000 | 0.000000 | 0.004374 | 0.000000 | NA | NA | NA | NA |
| PPP1R1B | Behavior | 0.000000 | 0.000717 | 0.000000 | 0.000172 | 0.000000 | 0.002932 | 0.000000 | 0.000005 | 0.000000 | 0.003019 | NA | NA | NA | NA |
| LEMD3 | Ear morphology | 0.000000 | 0.000000 | 0.000000 | 0.000000 | 0.000000 | 0.000000 | 0.000000 | 0.000000 | 0.000000 | 0.000000 | NA | NA | NA | NA |
| IGF2R | Lean growth | 0.001075 | 0.000170 | 0.000294 | 0.000050 | 0.004211 | 0.000647 | 0.000011 | 0.000002 | 0.012781 | 0.000672 | -1.922453 | -2.224774 | -2.001176 | -2.531273 |
| JMJD1C | Fertility | 0.003618 | 0.000910 | 0.002395 | 0.000821 | NA | NA | 0.000733 | 0.000286 | NA | NA | NA | -0.792304 | -0.314965 | -1.036534 |
| OSTN | Body composition | 0.002483 | 0.001570 | 0.001949 | 0.003402 | 0.000000 | 0.000000 | 0.000217 | 0.003581 | 0.001056 | 0.003402 | NA | 0.458305 | 0.803672 | -4.131299 |
| AHR | Litter size | 0.004804 | 0.000869 | 0.002405 | 0.000325 | 0.000000 | 0.000915 | NA | NA | 0.016325 | 0.001809 | -Inf | -0.219112 | -0.631894 | NA |

**Figure S1.** Scheme of the general historical processes used in forward simulations. A bottleneck event in the split between Wild and domestic populations was simulated for the second group of complex scenarios.

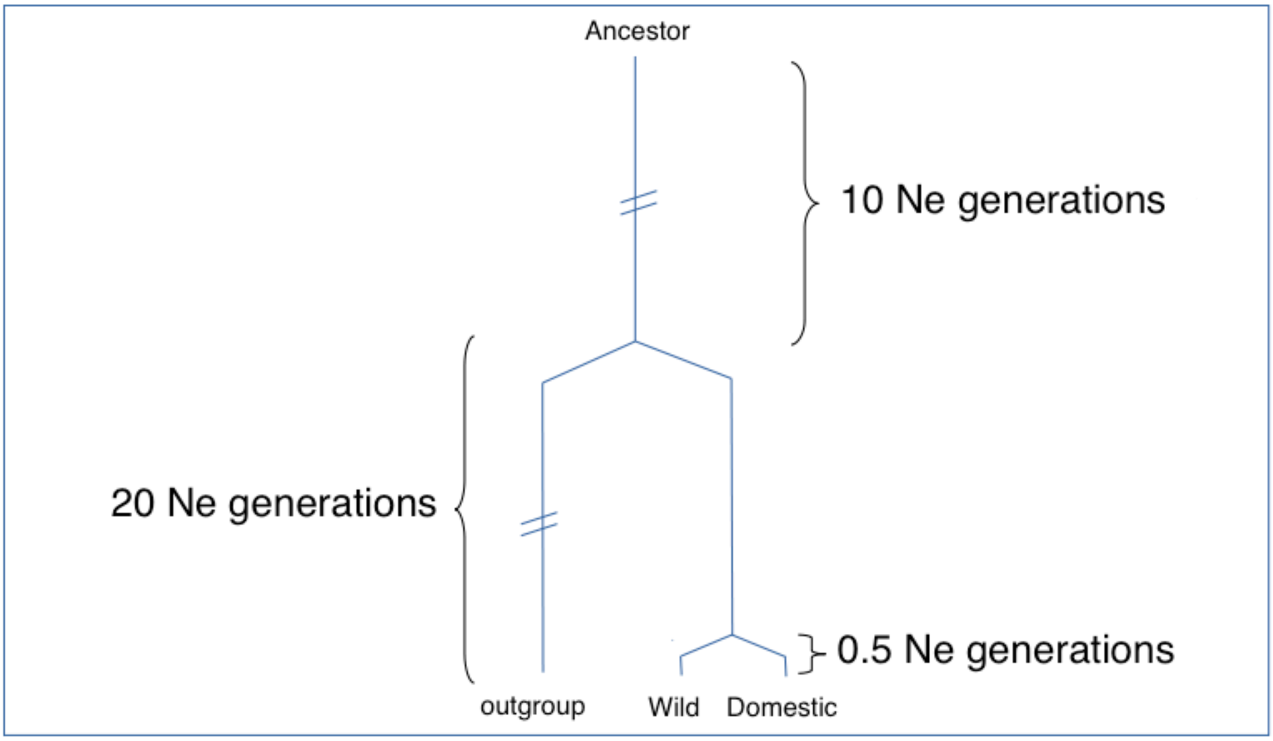

**Figure S2.** Cross validation of the parameters of the different ABC models. True values of parameters were obtained using polyDFE runs and were inferred using ABC using three different tolerances (0.05 -yellow-, 0.01 -orange- and 0.001 -red) and using the ratios of nonsynonymous versus synonymous variability as summary statistics. Relationship between true and estimated values for: (A) model A, (B) model C, (C) model DN, (D) model D (see Material and Methods for a complete description of model parameters).

***A***

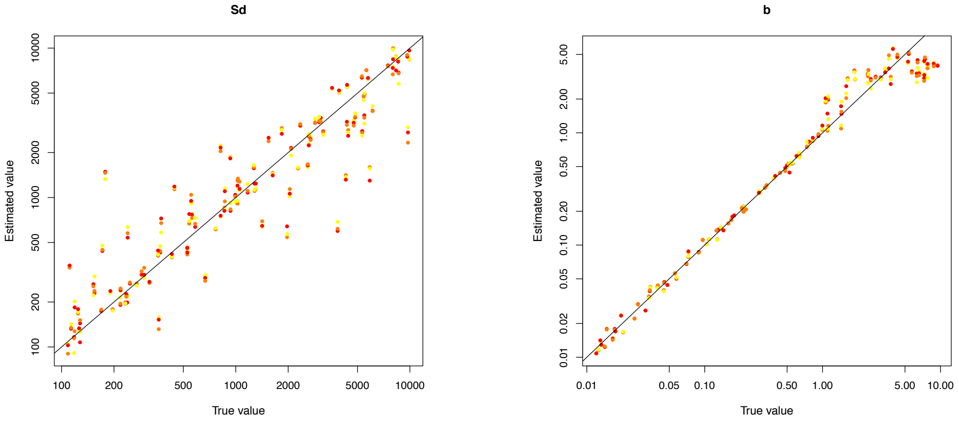


***B***

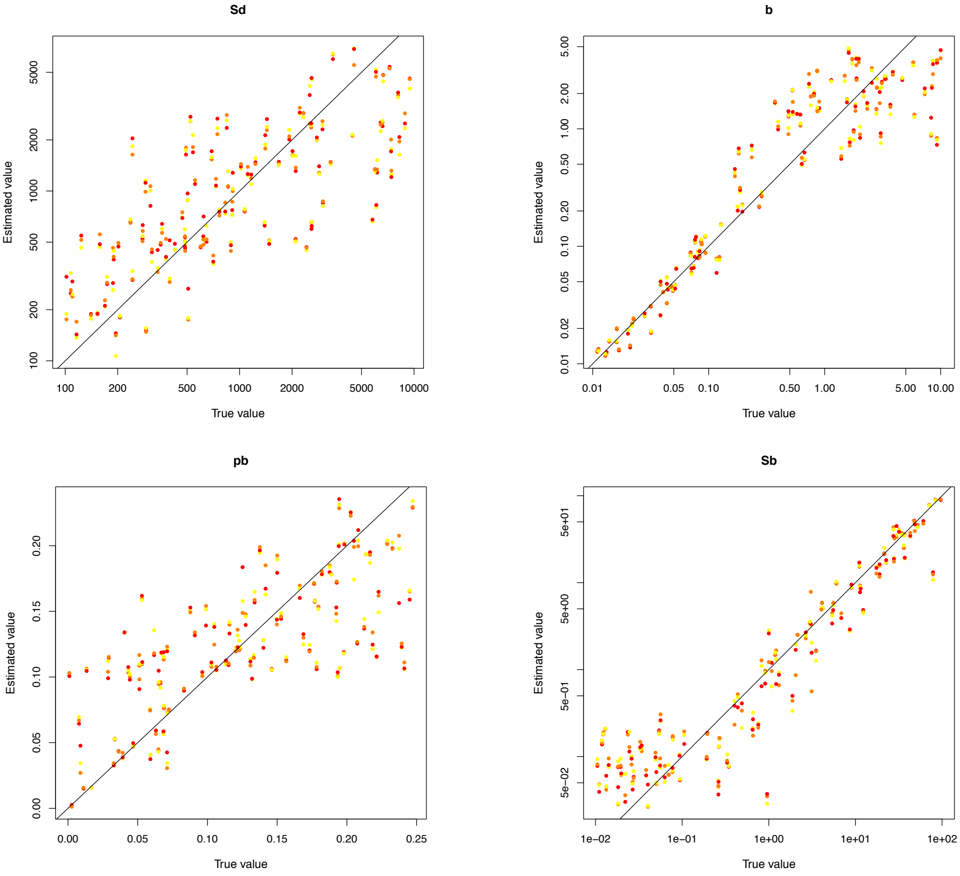

# C

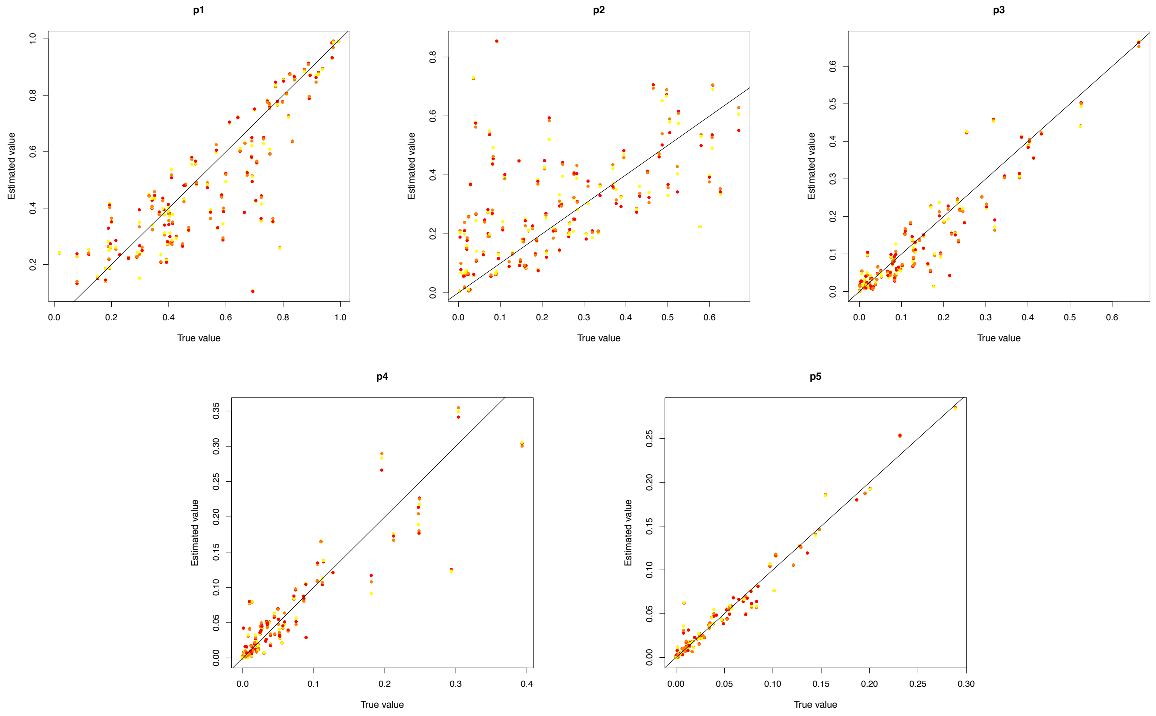
p1
p2
p3
p4
p5
Estimated value
True value

# D

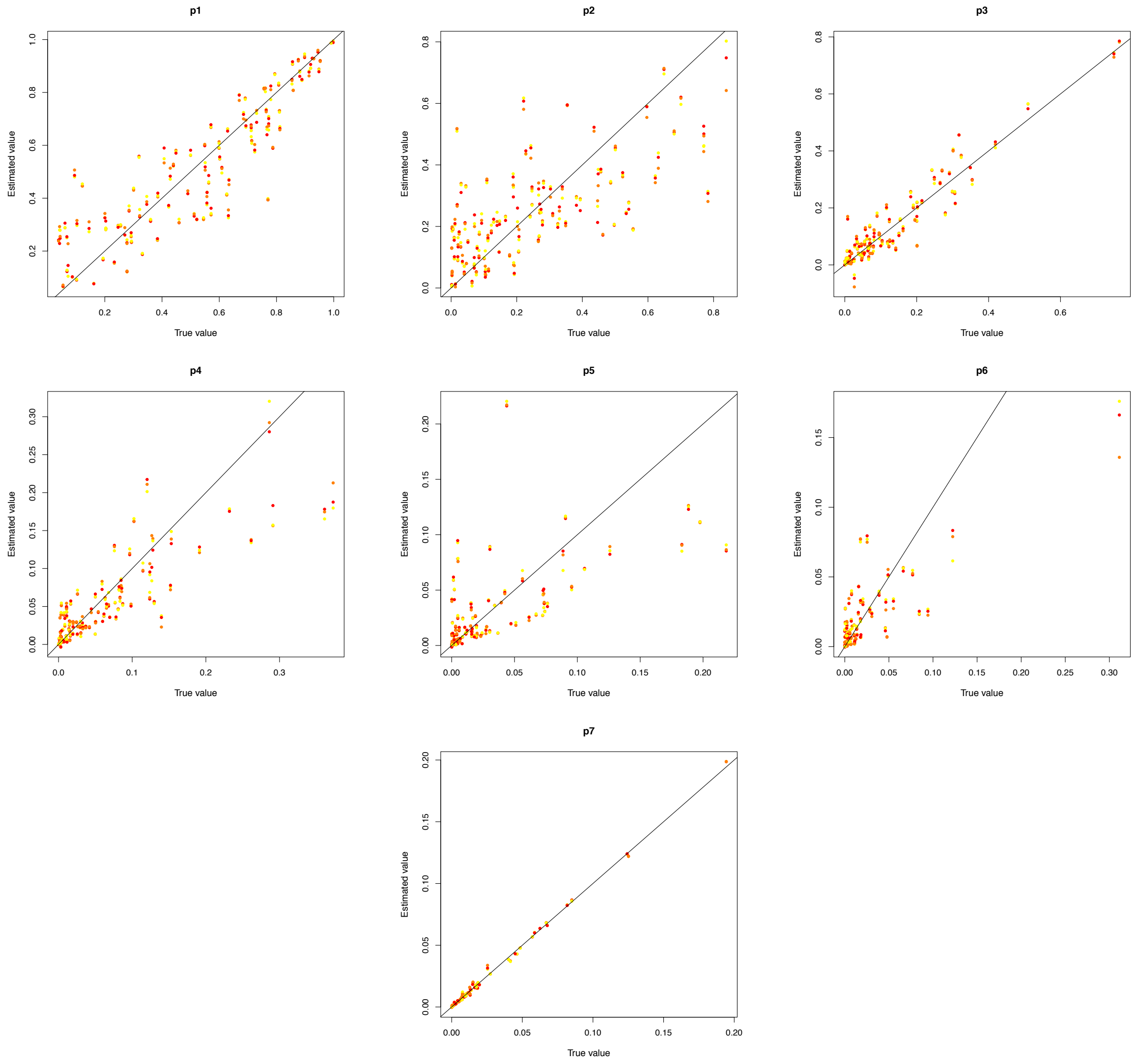
p1
p2
p3
p4
p5
p6
p7
Estimated value
True value

**Figure S3.** (A) Principal Component Analysis (PCA) using the total number of SNPs. (B) Admixture Analysis of population structure. Each K plot shows the different groups obtained and the degree of admixture between populations. Individuals are in the same order at each K value. (C) Coefficient of variation (left) and Evanno analysis (right) obtained with the most likely K value of K=2 (WB+IB, LW).

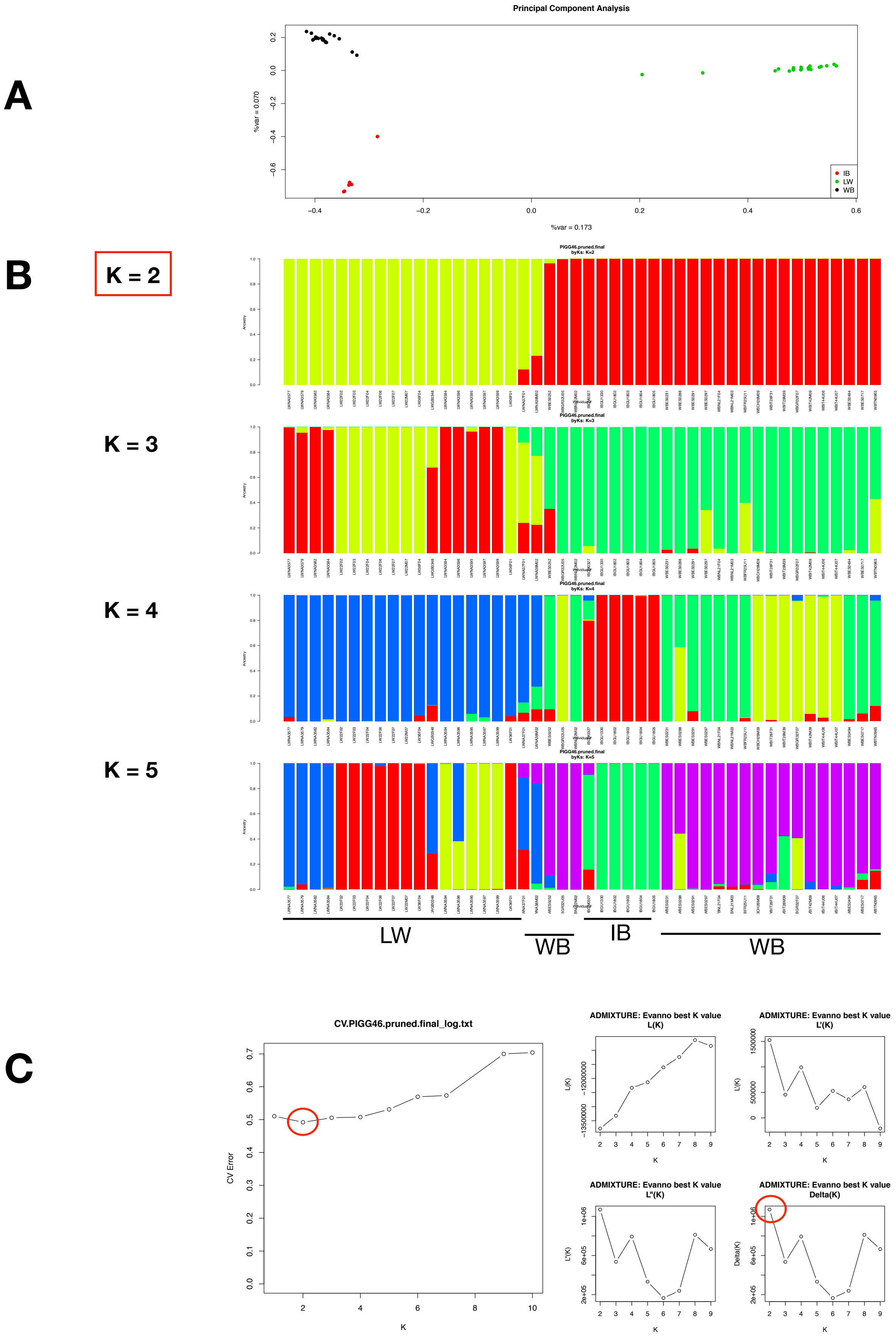

**Figure S4.** Average estimates of the network topology features (betweenness, in-degree and out-degree) for genes within pathways and grouping genes with positive and negative values of $\alpha$.

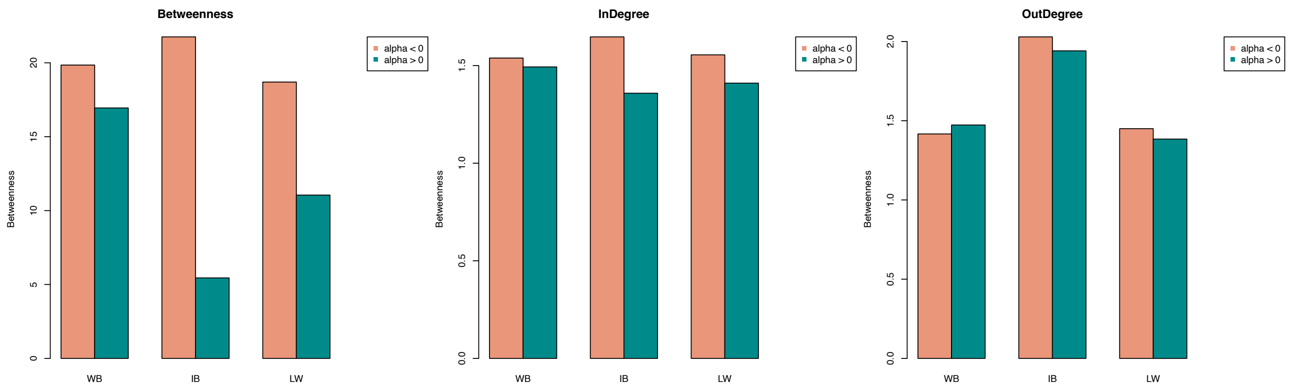

**Figure S5.** Distribution of the codon bias in the *Sus scrofa* genome. (A) Major Codon Usage (MCA), (B) Effective Number of Codons (Ncw). (C) Correlation of MCU versus positive estimates of $\alpha$ using Fu&Li, Watterson, Tajima and Fay&Wu variability estimates, respectively for Wild boar.

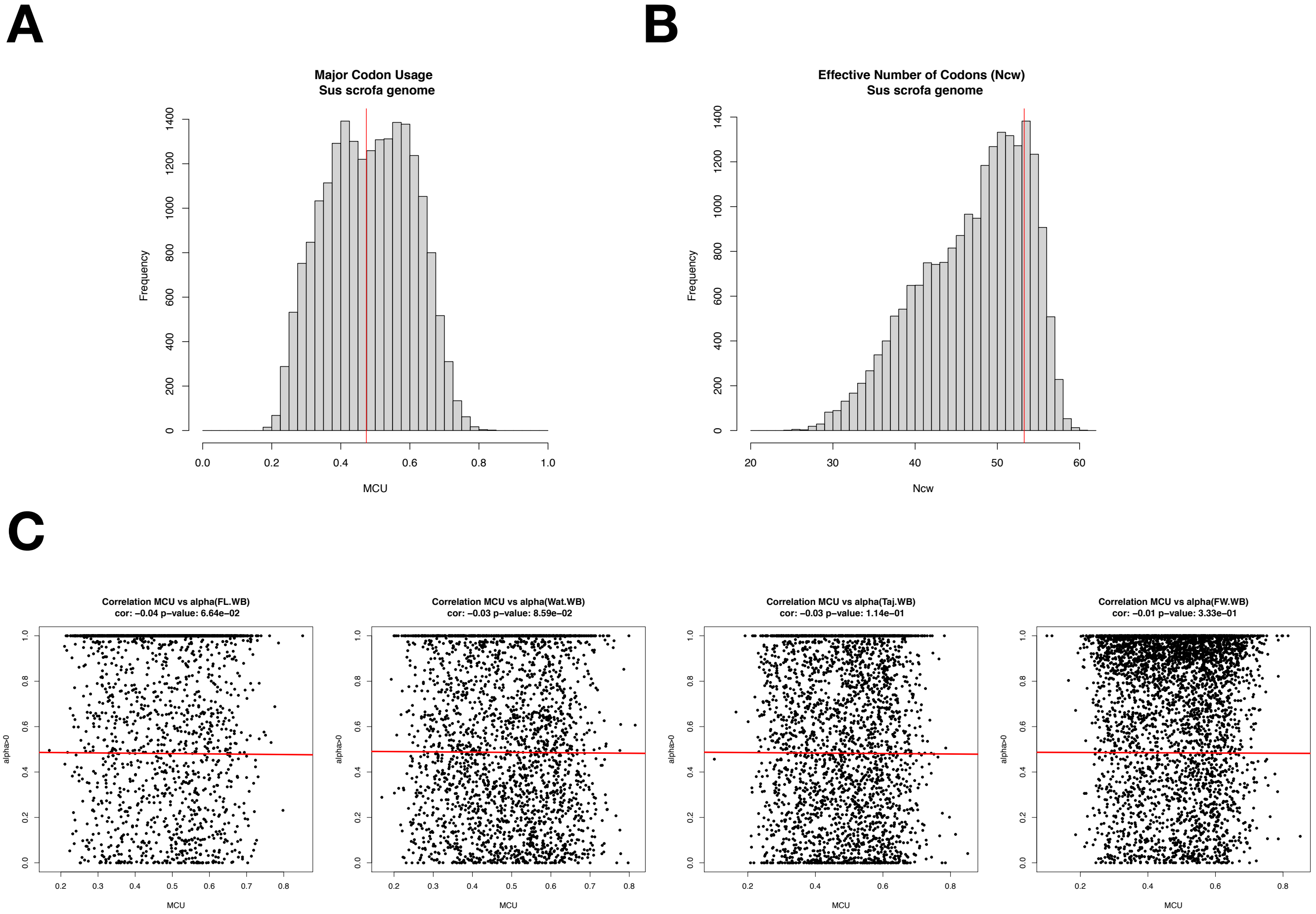

**Figure S6.** Site Frequency Spectrum and Asymptotic McDonald-Kreitman Test from a projection of whole data (around 25% of all variants; WB (n=38), IB (n=10) and LW (n=38)). (A) Whole Coding Positions (B) Exclusive Coding Positions. (C) Shared Coding Positions.

# A

### Wild Boar

#### Asymptotic McDonald–Kreitman Test: Results

**Analysis dataset:**

$d_0$ = 4189
$d$ = 1843
Input file = `SFS.WB.nsample38.txt`
$x$ interval = [0.100, 0.900]

**Plots:**

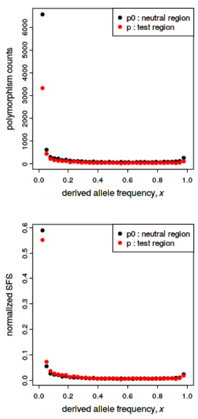


Fig. 1 [left]. Polymorphism levels in the test region ($p$, red points) and in the neutral reference region ($p_0$, black points), as a function of derived allele frequency $x$.

Fig. 2 [right]. Normalized site frequency spectra (SFS) in the test region (red points) and the neutral reference region (black points). This plot shows the data from Fig. 1, normalized such that $\Sigma_x\, p_0(x) = \Sigma_x\, p(x) = 1$ for purposes of comparison.

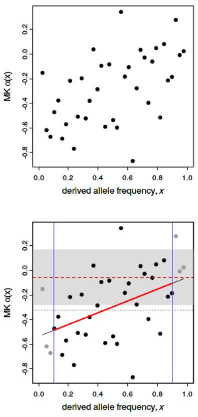


Fig. 3 [left]. McDonald–Kreitman $\alpha(x) = 1 - (d_0 / d)\,(p(x) / p_0(x))$ versus $x$.

Fig. 4 [right]. The asymptotic McDonald–Kreitman test results. This plot shows the data from Fig. 3, with fitting information superimposed. The blue vertical lines indicate the cutoff interval for the polymorphism data; points outside of the cutoff interval are plotted in gray, indicating that they were not used in the fit. The red curve shows the best fit to the data within the cutoff interval for a function $\alpha_{fit}(x) = a + bx$. The dashed red horizontal line shows the estimate of $\alpha_{asymptotic}$ from the fitted function, and the gray band indicates the 95% confidence interval around that estimate. Finally, the dotted gray horizontal line shows $\alpha_{original}$, the estimate from the original non-asymptotic McDonald–Kreitman test (also using only the data within the cutoff interval), for comparison; note that use of this value is not recommended.

**Fitted $\alpha(x)$:**

The linear model $\alpha(x) = a + bx$ was better (by AIC) than the exponential model, and is therefore reported here.

$a$ = -0.53703
$b$ = 0.48120
$c$ = NA

**Estimates of $\alpha$:**

The result of the asymptotic McDonald–Kreitman test is given by $\alpha_{asymptotic}$; this value is obtained by extrapolating the above fitted function to $x = 1$. The 95% confidence interval around the estimated value of $\alpha_{asymptotic}$ is also shown. The value of $\alpha_{original}$, from the original non-asymptotic McDonald–Kreitman test, is also given here for comparison but its use is not recommended. Both $\alpha$ estimates are derived from the polymorphism frequency data within the supplied cutoff interval for $x$.

$\alpha_{asymptotic}$ = -0.055831
95% CI(lower) = -0.28272
95% CI(upper) = 0.17106

$\alpha_{original}$ = -0.32330

### Iberian

#### Asymptotic McDonald–Kreitman Test: Results

**Analysis dataset:**

$d_0$ = 11017
$d$ = 4608
Input file = `SFS.IB.nsample10.txt`
$x$ interval = [0.100, 0.900]

**Plots:**

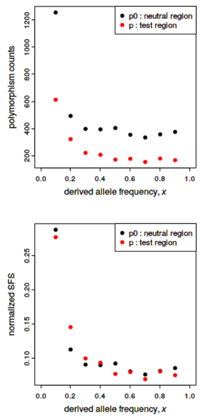


Fig. 1 [left]. Polymorphism levels in the test region ($p$, red points) and in the neutral reference region ($p_0$, black points), as a function of derived allele frequency $x$.

Fig. 2 [right]. Normalized site frequency spectra (SFS) in the test region (red points) and the neutral reference region (black points). This plot shows the data from Fig. 1, normalized such that $\Sigma_x\, p_0(x) = \Sigma_x\, p(x) = 1$ for purposes of comparison.

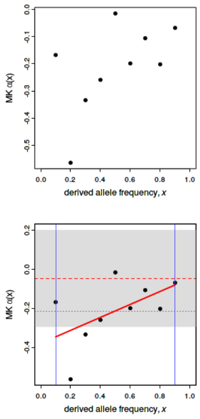


Fig. 3 [left]. McDonald–Kreitman $\alpha(x) = 1 - (d_0 / d)\,(p(x) / p_0(x))$ versus $x$.

Fig. 4 [right]. The asymptotic McDonald–Kreitman test results. This plot shows the data from Fig. 3, with fitting information superimposed. The blue vertical lines indicate the cutoff interval for the polymorphism data; points outside of the cutoff interval are plotted in gray, indicating that they were not used in the fit. The red curve shows the best fit to the data within the cutoff interval for a function $\alpha_{fit}(x) = a + bx$. The dashed red horizontal line shows the estimate of $\alpha_{asymptotic}$ from the fitted function, and the gray band indicates the 95% confidence interval around that estimate. Finally, the dotted gray horizontal line shows $\alpha_{original}$, the estimate from the original non-asymptotic McDonald–Kreitman test (also using only the data within the cutoff interval), for comparison; note that use of this value is not recommended.

**Fitted $\alpha(x)$:**

The linear model $\alpha(x) = a + bx$ was better (by AIC) than the exponential model, and is therefore reported here.

$a$ = -0.37908
$b$ = 0.33276
$c$ = NA

**Estimates of $\alpha$:**

The result of the asymptotic McDonald–Kreitman test is given by $\alpha_{asymptotic}$; this value is obtained by extrapolating the above fitted function to $x = 1$. The 95% confidence interval around the estimated value of $\alpha_{asymptotic}$ is also shown. The value of $\alpha_{original}$, from the original non-asymptotic McDonald–Kreitman test, is also given here for comparison but its use is not recommended. Both $\alpha$ estimates are derived from the polymorphism frequency data within the supplied cutoff interval for $x$.

$\alpha_{asymptotic}$ = -0.046322
95% CI(lower) = -0.29498
95% CI(upper) = 0.20234

$\alpha_{original}$ = -0.21374

### Largewhite

#### Asymptotic McDonald–Kreitman Test: Results

**Analysis dataset:**

$d_0$ = 3169
$d$ = 1397
Input file = `SFS.LW.nsample38.txt`
$x$ interval = [0.100, 0.900]

**Plots:**

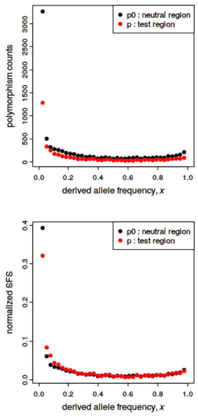


Fig. 1 [left]. Polymorphism levels in the test region ($p$, red points) and in the neutral reference region ($p_0$, black points), as a function of derived allele frequency $x$.

Fig. 2 [right]. Normalized site frequency spectra (SFS) in the test region (red points) and the neutral reference region (black points). This plot shows the data from Fig. 1, normalized such that $\Sigma_x\, p_0(x) = \Sigma_x\, p(x) = 1$ for purposes of comparison.

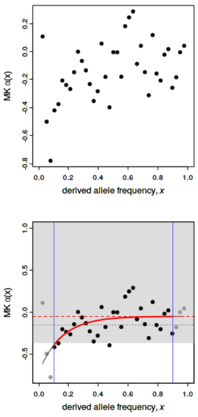


Fig. 3 [left]. McDonald–Kreitman $\alpha(x) = 1 - (d_0 / d)\,(p(x) / p_0(x))$ versus $x$.

Fig. 4 [right]. The asymptotic McDonald–Kreitman test results. This plot shows the data from Fig. 3, with fitting information superimposed. The blue vertical lines indicate the cutoff interval for the polymorphism data; points outside of the cutoff interval are plotted in gray, indicating that they were not used in the fit. The red curve shows the best fit to the data within the cutoff interval for a function $\alpha_{fit}(x) = a + b\,\exp(-cx)$. The dashed red horizontal line shows the estimate of $\alpha_{asymptotic}$ from the fitted function, and the gray band indicates the 95% confidence interval around that estimate. Finally, the dotted gray horizontal line shows $\alpha_{original}$, the estimate from the original non-asymptotic McDonald–Kreitman test (also using only the data within the cutoff interval), for comparison; note that use of this value is not recommended.

**Fitted $\alpha(x)$:**

The exponential model $\alpha(x) = a + b\,\exp(-cx)$ was better (by AIC) than the linear model, and is therefore reported here.

$a$ = -0.053984
$b$ = -0.66985
$c$ = 6.64684

**Estimates of $\alpha$:**

The result of the asymptotic McDonald–Kreitman test is given by $\alpha_{asymptotic}$; this value is obtained by extrapolating the above fitted function to $x = 1$. The 95% confidence interval around the estimated value of $\alpha_{asymptotic}$ is also shown. The value of $\alpha_{original}$, from the original non-asymptotic McDonald–Kreitman test, is also given here for comparison but its use is not recommended. Both $\alpha$ estimates are derived from the polymorphism frequency data within the supplied cutoff interval for $x$.

$\alpha_{asymptotic}$ = -0.054854
95% CI(lower) = -0.37087
95% CI(upper) = 7.01388

$\alpha_{original}$ = -0.16020

# B

### Wild Boar

#### Asymptotic McDonald–Kreitman Test: Results

**Analysis dataset:**

$d_0$ = 4189
$d$ = 1843
Input file = `SFS.WB.privateSNPs.nsample38.txt`
$x$ interval = [0.100, 0.900]

**Plots:**

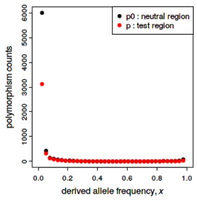


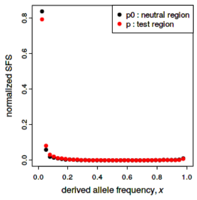


Fig. 1 [left]. Polymorphism levels in the test region ($p$, red points) and in the neutral reference region ($p_0$, black points), as a function of derived allele frequency $x$.

Fig. 2 [right]. Normalized site frequency spectra (SFS) in the test region (red points) and the neutral reference region (black points). This plot shows the data from Fig. 1, normalized such that $\Sigma_x\, p_0(x) = \Sigma_x\, p(x) = 1$ for purposes of comparison.

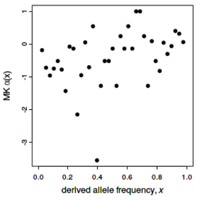


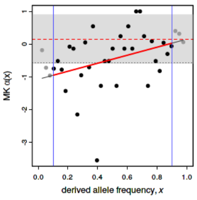


Fig. 3 [left]. McDonald–Kreitman $\alpha(x) = 1 - (d_0 / d)\,(p(x) / p_0(x))$ versus $x$.

Fig. 4 [right]. The asymptotic McDonald–Kreitman test results. This plot shows the data from Fig. 3, with fitting information superimposed. The blue vertical lines indicate the cutoff interval for the polymorphism data; points outside of the cutoff interval are plotted in gray, indicating that they were not used in the fit. The red curve shows the best fit to the data within the cutoff interval for a function $\alpha_{fit}(x) = a + bx$. The dashed red horizontal line shows the estimate of $\alpha_{asymptotic}$ from the fitted function, and the gray band indicates the 95% confidence interval around that estimate. Finally, the dotted gray horizontal line shows $\alpha_{original}$, the estimate from the original non-asymptotic

McDonald–Kreitman test (also using only the data within the cutoff interval), for comparison; note that use of this value is not recommended.

**Fitted $\alpha(x)$:**

The linear model $\alpha(x) = a + bx$ was better (by AIC) than the exponential model, and is therefore reported here.

$a$ = -1.07270
$b$ = 1.22816
$c$ = NA

**Estimates of $\alpha$:**

The result of the asymptotic McDonald–Kreitman test is given by $\alpha_{asymptotic}$; this value is obtained by extrapolating the above fitted function to $x = 1$. The 95% confidence interval around the estimated value of $\alpha_{asymptotic}$ is also shown. The value of $\alpha_{original}$, from the original non-asymptotic McDonald–Kreitman test, is also given here for comparison but its use is not recommended. Both $\alpha$ estimates are derived from the polymorphism frequency data within the supplied cutoff interval for $x$.

$\alpha_{asymptotic}$ = 0.15546
95% CI(lower) = -0.59857
95% CI(upper) = 0.90949

$\alpha_{original}$ = -0.56558

### Iberian

#### Asymptotic McDonald–Kreitman Test: Results

**Analysis dataset:**

$d_0$ = 11017
$d$ = 4608
Input file = `SFS.IB.privateSNPs.nsample10.txt`
$x$ interval = [0.100, 0.900]

**Plots:**

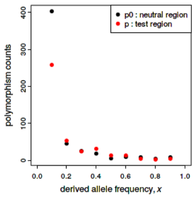


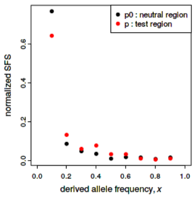


Fig. 1 [left]. Polymorphism levels in the test region ($p$, red points) and in the neutral reference region ($p_0$, black points), as a function of derived allele frequency $x$.

Fig. 2 [right]. Normalized site frequency spectra (SFS) in the test region (red points) and the neutral reference region (black points). This plot shows the data from Fig. 1, normalized such that $\Sigma_x\, p_0(x) = \Sigma_x\, p(x) = 1$ for purposes of comparison.

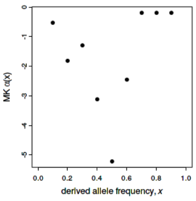


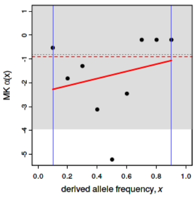


Fig. 3 [left]. McDonald–Kreitman $\alpha(x) = 1 - (d_0 / d)\,(p(x) / p_0(x))$ versus $x$.

Fig. 4 [right]. The asymptotic McDonald–Kreitman test results. This plot shows the data from Fig. 3, with fitting information superimposed. The blue vertical lines indicate the cutoff interval for the polymorphism data; points outside of the cutoff interval are plotted in gray, indicating that they were not used in the fit. The red curve shows the best fit to the data within the cutoff interval for a function $\alpha_{fit}(x) = a + bx$. The dashed red horizontal line shows the estimate of $\alpha_{asymptotic}$ from the fitted function, and the gray band indicates the 95% confidence interval around that estimate. Finally, the dotted gray horizontal line shows $\alpha_{original}$, the estimate from the original non-asymptotic

McDonald–Kreitman test (also using only the data within the cutoff interval), for comparison; note that use of this value is not recommended.

**Fitted $\alpha(x)$:**

The exponential fit failed to converge (usually because the data are not exponential in shape); the linear model is therefore reported here.

$a$ = -2.42384
$b$ = 1.51097
$c$ = NA

**Estimates of $\alpha$:**

The result of the asymptotic McDonald–Kreitman test is given by $\alpha_{asymptotic}$; this value is obtained by extrapolating the above fitted function to $x = 1$. The 95% confidence interval around the estimated value of $\alpha_{asymptotic}$ is also shown. The value of $\alpha_{original}$, from the original non-asymptotic McDonald–Kreitman test, is also given here for comparison but its use is not recommended. Both $\alpha$ estimates are derived from the polymorphism frequency data within the supplied cutoff interval for $x$.

$\alpha_{asymptotic}$ = -0.91286
95% CI(lower) = -3.95123
95% CI(upper) = 2.12550

$\alpha_{original}$ = -0.83070

### Largewhite

#### Asymptotic McDonald–Kreitman Test: Results

**Analysis dataset:**

$d_0$ = 3169
$d$ = 1397
Input file = `SFS.LW.privateSNPs.nsample38.txt`
$x$ interval = [0.100, 0.900]

**Plots:**

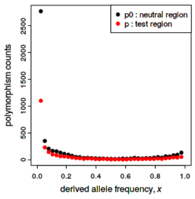


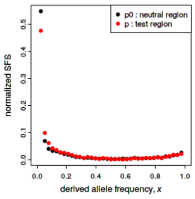


Fig. 1 [left]. Polymorphism levels in the test region ($p$, red points) and in the neutral reference region ($p_0$, black points), as a function of derived allele frequency $x$.

Fig. 2 [right]. Normalized site frequency spectra (SFS) in the test region (red points) and the neutral reference region (black points). This plot shows the data from Fig. 1, normalized such that $\Sigma_x\, p_0(x) = \Sigma_x\, p(x) = 1$ for purposes of comparison.

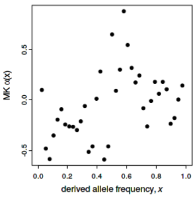


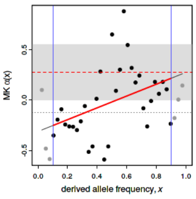


Fig. 3 [left]. McDonald–Kreitman $\alpha(x) = 1 - (d_0 / d)\,(p(x) / p_0(x))$ versus $x$.

Fig. 4 [right]. The asymptotic McDonald–Kreitman test results. This plot shows the data from Fig. 3, with fitting information superimposed. The blue vertical lines indicate the cutoff interval for the polymorphism data; points outside of the cutoff interval are plotted in gray, indicating that they were not used in the fit. The red curve shows the best fit to the data within the cutoff interval for a function $\alpha_{fit}(x) = a + bx$. The dashed red horizontal line shows the estimate of $\alpha_{asymptotic}$ from the fitted function, and the gray band indicates the 95% confidence interval around that estimate. Finally, the dotted gray horizontal line shows $\alpha_{original}$, the estimate from the original non-asymptotic

McDonald–Kreitman test (also using only the data within the cutoff interval), for comparison; note that use of this value is not recommended.

**Fitted $\alpha(x)$:**

The linear model $\alpha(x) = a + bx$ was better (by AIC) than the exponential model, and is therefore reported here.

$a$ = -0.31017
$b$ = 0.58668
$c$ = NA

**Estimates of $\alpha$:**

The result of the asymptotic McDonald–Kreitman test is given by $\alpha_{asymptotic}$; this value is obtained by extrapolating the above fitted function to $x = 1$. The 95% confidence interval around the estimated value of $\alpha_{asymptotic}$ is also shown. The value of $\alpha_{original}$, from the original non-asymptotic McDonald–Kreitman test, is also given here for comparison but its use is not recommended. Both $\alpha$ estimates are derived from the polymorphism frequency data within the supplied cutoff interval for $x$.

$\alpha_{asymptotic}$ = 0.27651
95% CI(lower) = -0.0012247
95% CI(upper) = 0.55424

$\alpha_{original}$ = -0.12391

# C

### Wild Boar

#### Asymptotic McDonald–Kreitman Test: Results

**Analysis dataset:**

$d_0$ = 4189
$d$ = 1843
Input file = `SFS.WB.allsharedSNPs.nsample38.txt`
$x$ interval = [0.100, 0.900]

**Plots:**

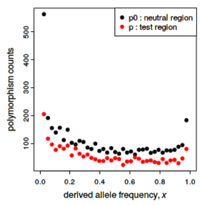


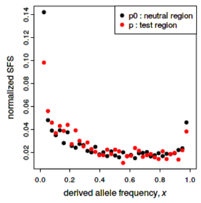


Fig. 1 [left]. Polymorphism levels in the test region ($p$, red points) and in the neutral reference region ($p_0$, black points), as a function of derived allele frequency $x$.

Fig. 2 [right]. Normalized site frequency spectra (SFS) in the test region (red points) and the neutral reference region (black points). This plot shows the data from Fig. 1, normalized such that $\Sigma_x\, p_0(x) = \Sigma_x\, p(x) = 1$ for purposes of comparison.

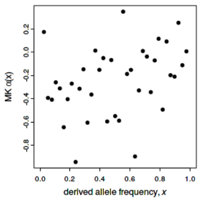


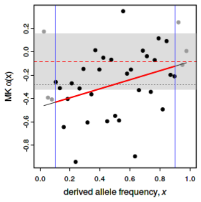


Fig. 3 [left]. McDonald–Kreitman $\alpha(x) = 1 - (d_0 / d)\ (p(x) / p_0(x))$ versus $x$.

Fig. 4 [right]. The asymptotic McDonald–Kreitman test results. This plot shows the data from Fig. 3, with fitting information superimposed. The blue vertical lines indicate the cutoff interval for the polymorphism data; points outside of the cutoff interval are plotted in gray, indicating that they were not used in the fit. The red curve shows the best fit to the data within the cutoff interval for a function $\alpha_{\mathrm{fit}}(x) = a + bx$. The dashed red horizontal line shows the estimate of $\alpha_{\mathrm{asymptotic}}$ from the fitted function, and the gray band indicates the 95% confidence interval around that estimate. Finally, the dotted gray horizontal line shows $\alpha_{\mathrm{original}}$, the estimate from the original non-asymptotic

McDonald–Kreitman test (also using only the data within the cutoff interval), for comparison; note that use of this value is not recommended.

**Fitted $\alpha(x)$:**

The linear model $\alpha(x) = a + bx$ was better (by AIC) than the exponential model, and is therefore reported here.

$a$ = -0.47093
$b$ = 0.38691
$c$ = NA

**Estimates of $\alpha$:**

The result of the asymptotic McDonald–Kreitman test is given by $\alpha_{\mathrm{asymptotic}}$; this value is obtained by extrapolating the above fitted function to $x = 1$. The 95% confidence interval around the estimated value of $\alpha_{\mathrm{asymptotic}}$ is also shown. The value of $\alpha_{\mathrm{original}}$, from the original non-asymptotic McDonald–Kreitman test, is also given here for comparison but its use is not recommended. Both $\alpha$ estimates are derived from the polymorphism frequency data within the supplied cutoff interval for $x$.

$\alpha_{\mathrm{asymptotic}}$ = -0.084028
95% CI(lower) = -0.32488
95% CI(upper) = 0.15682

$\alpha_{\mathrm{original}}$ = -0.27984

### Iberian

#### symptotic McDonald–Kreitman Test: Results

**alysis dataset:**

$d_0$ = 11017
$d$ = 4608
Input file = `SFS.IB.allsharedSNPs.nsample10.txt`
$x$ interval = [0.100, 0.900]

**ts:**

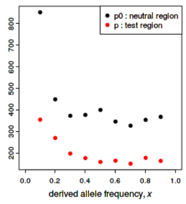


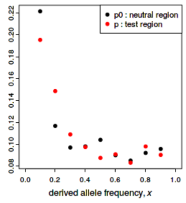


. 1 [left]. Polymorphism levels in the test region ($p$, red points) and in the neutral erence region ($p_0$, black points), as a function of derived allele frequency $x$.

. 2 [right]. Normalized site frequency spectra (SFS) in the test region (red points) l the neutral reference region (black points). This plot shows the data from Fig. 1, rmalized such that $\Sigma_x\, p_0(x) = \Sigma_x\, p(x) = 1$ for purposes of comparison.

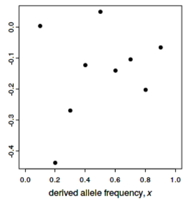


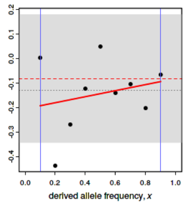


. 3 [left]. McDonald–Kreitman $\alpha(x) = 1 - (d_0 / d)\ (p(x) / p_0(x))$ versus $x$.

. 4 [right]. The asymptotic McDonald–Kreitman test results. This plot shows the a from Fig. 3, with fitting information superimposed. The blue vertical lines indicate cutoff interval for the polymorphism data; points outside of the cutoff interval are tted in gray, indicating that they were not used in the fit. The red curve shows the t fit to the data within the cutoff interval for a function $\alpha_{\mathrm{fit}}(x) = a + bx$. The dashed l horizontal line shows the estimate of $\alpha_{\mathrm{asymptotic}}$ from the fitted function, and the gray nd indicates the 95% confidence interval around that estimate. Finally, the dotted y horizontal line shows $\alpha_{\mathrm{original}}$, the estimate from the original non-asymptotic

:Donald–Kreitman test (also using only the data within the cutoff interval), for nparison; note that use of this value is not recommended.

**ted $\alpha(x)$:**

e exponential fit failed to converge (usually because the data are not exponential in ipe); the linear model is therefore reported here.

$a$ = -0.20491
$b$ = 0.12366
$c$ = NA

**imates of $\alpha$:**

e result of the asymptotic McDonald–Kreitman test is given by $\alpha_{\mathrm{asymptotic}}$; this value is ained by extrapolating the above fitted function to $x = 1$. The 95% confidence erval around the estimated value of $\alpha_{\mathrm{asymptotic}}$ is also shown. The value of $\alpha_{\mathrm{original}}$, m the original non-asymptotic McDonald–Kreitman test, is also given here for nparison but its use is not recommended. Both $\alpha$ estimates are derived from the lymorphism frequency data within the supplied cutoff interval for $x$.

$\alpha_{\mathrm{asymptotic}}$ = -0.081249
95% CI(lower) = -0.34396
95% CI(upper) = 0.18146

$\alpha_{\mathrm{original}}$ = -0.12953

### Largewhite

#### Asymptotic McDonald–Kreitman Test: Results

**Analysis dataset:**

$d_0$ = 3169
$d$ = 1397
Input file = `SFS.LW.allsharedSNPs.nsample38.txt`
$x$ interval = [0.100, 0.900]

**Plots:**

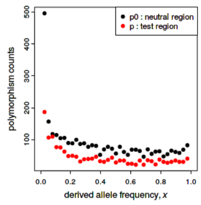


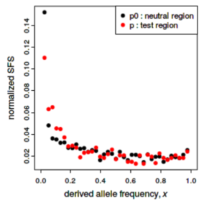


Fig. 1 [left]. Polymorphism levels in the test region ($p$, red points) and in the neutral reference region ($p_0$, black points), as a function of derived allele frequency $x$.

Fig. 2 [right]. Normalized site frequency spectra (SFS) in the test region (red points) and the neutral reference region (black points). This plot shows the data from Fig. 1, normalized such that $\Sigma_x\, p_0(x) = \Sigma_x\, p(x) = 1$ for purposes of comparison.

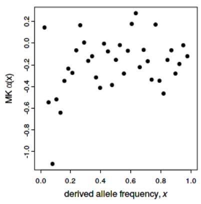


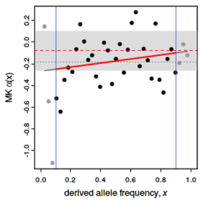


Fig. 3 [left]. McDonald–Kreitman $\alpha(x) = 1 - (d_0 / d)\ (p(x) / p_0(x))$ versus $x$.

Fig. 4 [right]. The asymptotic McDonald–Kreitman test results. This plot shows the data from Fig. 3, with fitting information superimposed. The blue vertical lines indicate the cutoff interval for the polymorphism data; points outside of the cutoff interval are plotted in gray, indicating that they were not used in the fit. The red curve shows the best fit to the data within the cutoff interval for a function $\alpha_{\mathrm{fit}}(x) = a + bx$. The dashed red horizontal line shows the estimate of $\alpha_{\mathrm{asymptotic}}$ from the fitted function, and the gray band indicates the 95% confidence interval around that estimate. Finally, the dotted gray horizontal line shows $\alpha_{\mathrm{original}}$, the estimate from the original non-asymptotic

McDonald–Kreitman test (also using only the data within the cutoff interval), for comparison; note that use of this value is not recommended.

**Fitted $\alpha(x)$:**

The linear model $\alpha(x) = a + bx$ was better (by AIC) than the exponential model, and is therefore reported here.

$a$ = -0.26768
$b$ = 0.18777
$c$ = NA

**Estimates of $\alpha$:**

The result of the asymptotic McDonald–Kreitman test is given by $\alpha_{\mathrm{asymptotic}}$; this value is obtained by extrapolating the above fitted function to $x = 1$. The 95% confidence interval around the estimated value of $\alpha_{\mathrm{asymptotic}}$ is also shown. The value of $\alpha_{\mathrm{original}}$, from the original non-asymptotic McDonald–Kreitman test, is also given here for comparison but its use is not recommended. Both $\alpha$ estimates are derived from the polymorphism frequency data within the supplied cutoff interval for $x$.

$\alpha_{\mathrm{asymptotic}}$ = -0.079914
95% CI(lower) = -0.26374
95% CI(upper) = 0.10391

$\alpha_{\mathrm{original}}$ = -0.18297

**Figure S7.** Estimates of $\alpha$ using different variability estimates considering all SNPs and for simulations under the SNM and Negative and Positive selection scenarios. The demographic history of pigs was simulated as a starting population of 10,000 diploid individuals and was left to evolve 100,000 generations to stabilize the mutations. After that, the population split in two populations of 10,000 individuals (corresponding to the observed divergence between *Sus scrofa* and the outgroup) and evolved 200,000 generations. 5,000 generations before present, one of the populations splits in two populations, which would correspond to domestic pigs and wild boars. In case of simulating reduction or expansion of the population size of the branch corresponding to domestic pigs, we will decrease (or increase) 10 times the number of individuals 4,000 generations before present. Every condition was run 100 times in order to have a distribution of the expected values. For each run, 100 sequences out of the 20,000 simulated sequences for each population were randomly sampled 10 times, eventually having 1,000 samples for each evolutionary model. General parameters were: mutation rate: 2.5e-7, recombination rate: 1.17e-8, 10.000 positions for each coding region, with 2/3 of positions as nonsynonymous and 1/3 as synonymous. Additivity ($h = 0.5$) was assumed for all mutations.

OBSERVED PATTERNS α TOTAL

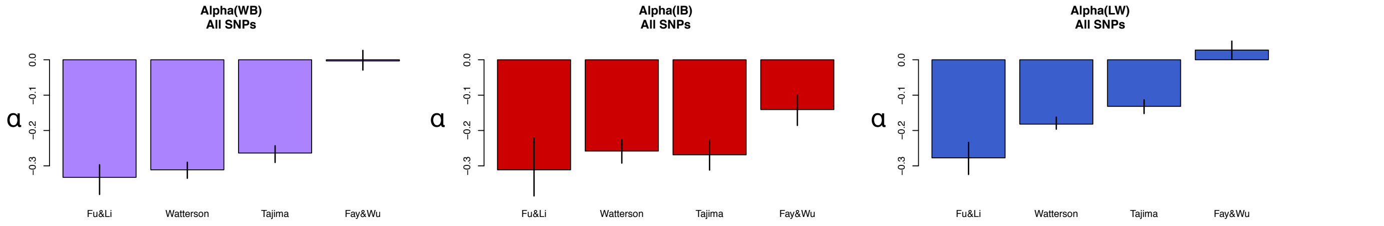


SIMULATED PATTERNS α TOTAL

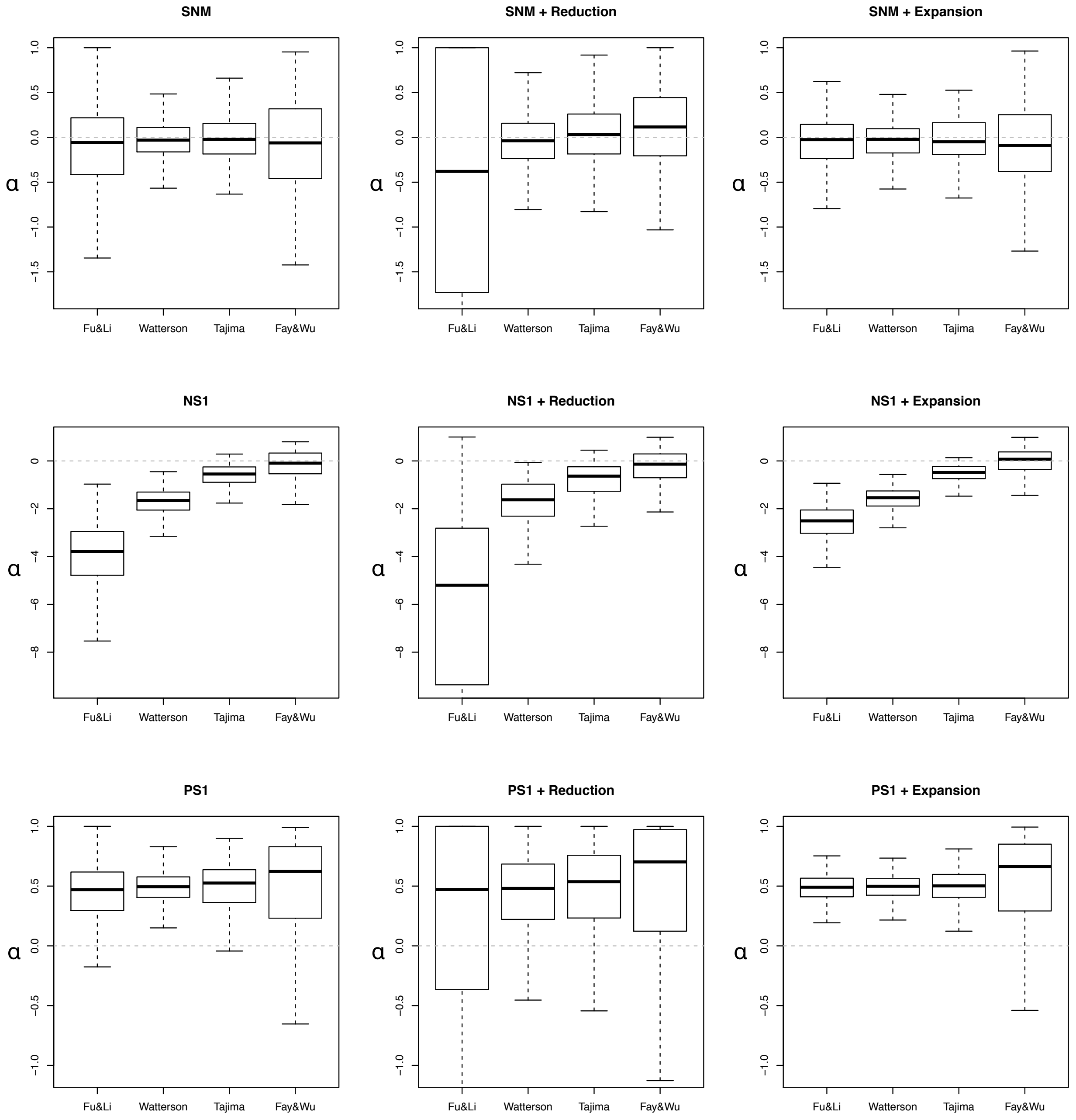

**Figure S8.** Estimates of *R_βγ* between Wild and Domestic populations using different variability estimates and considering Total SNPs and for simulation under the SNM, and Negative and Positive selection scenarios.

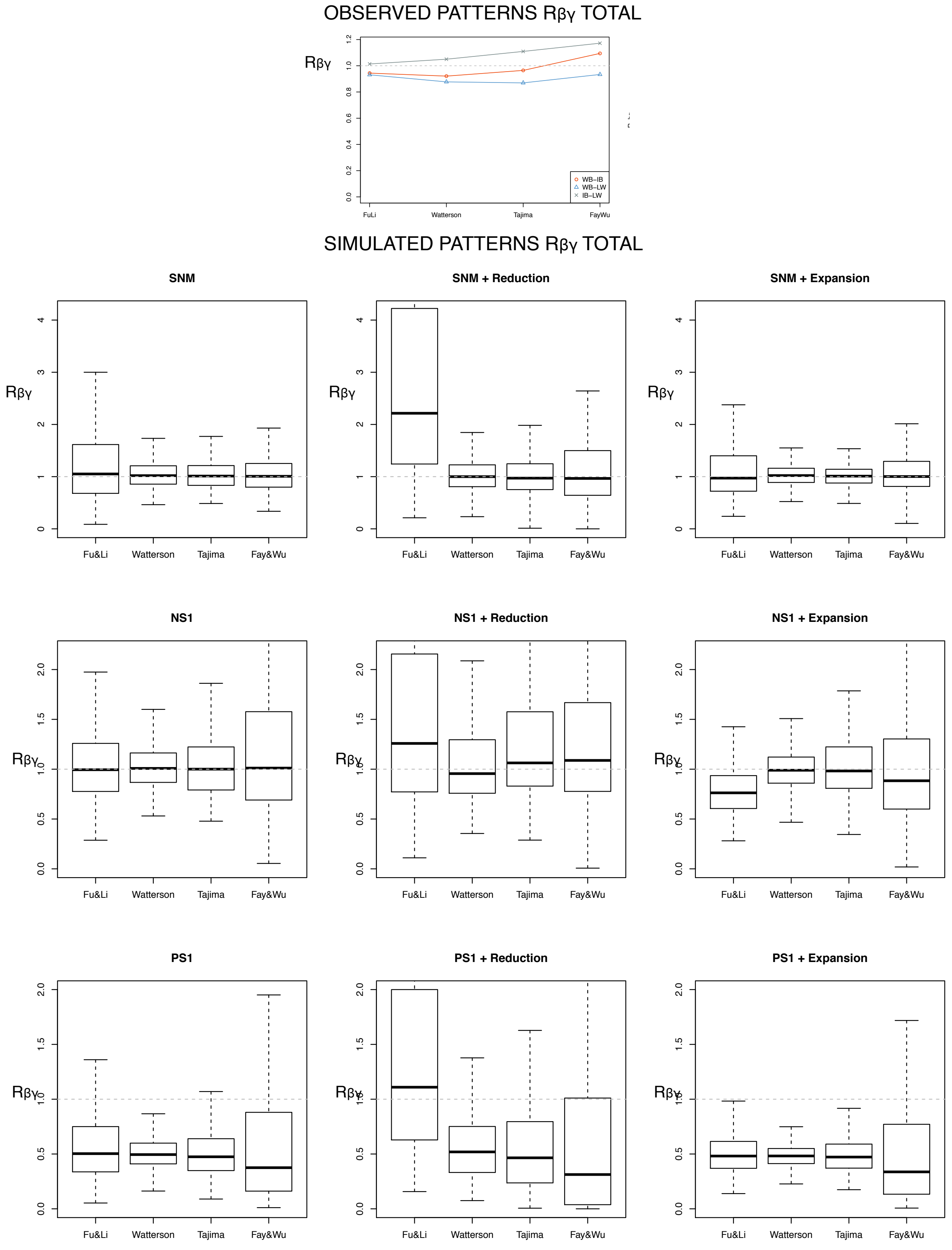

**Figure S9.** Estimates of $\alpha$ considering **Total** SNPs for the Domestic population. A bottleneck was simulated after the split of Wild and Domestic populations (0.1x reduction of population size during 0.1 Ne generations, where Ne is the ancestral population size;  see Table S3B for a description of the parameter values used in these simulations). The different scenarios were simulated using different combinations of positive and negative selection coefficients and considering **no migration** from Wild to Domestic populations and **no change** in the population size after the bottleneck.

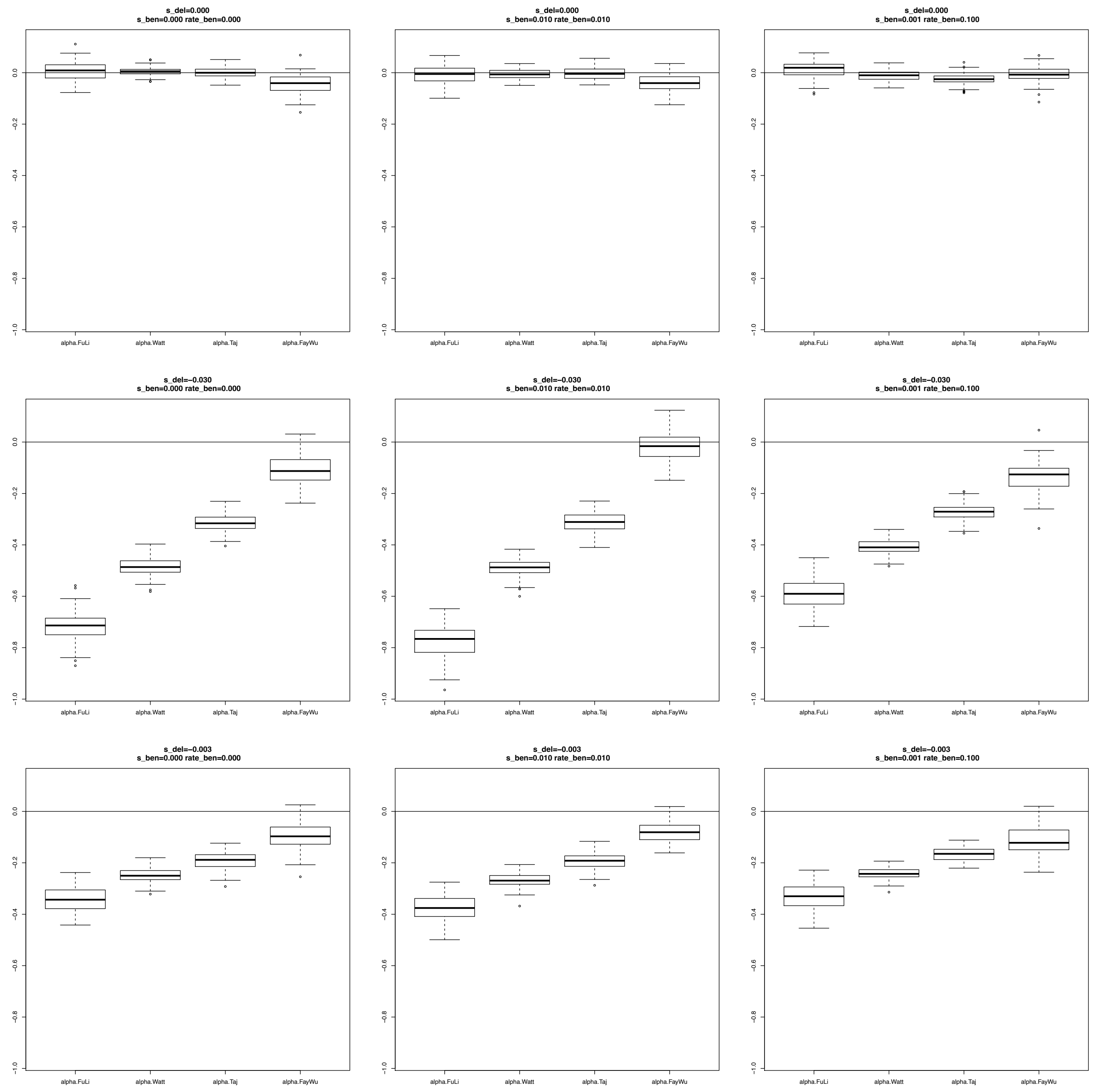

**Figure S10.** Estimates of $\alpha$ considering **Total** SNPs for the Domestic population. A bottleneck was simulated after the split of Wild and Domestic populations (0.1x reduction of population size during 0.1 Ne generations, where Ne is the ancestral population size; see Table S3B for a description of the parameter values used in these simulations). The different scenarios were simulated using different combinations of positive and negative selection coefficients considering **5% migrants** from Wild to Domestic populations and **no change** in the population size after the bottleneck.

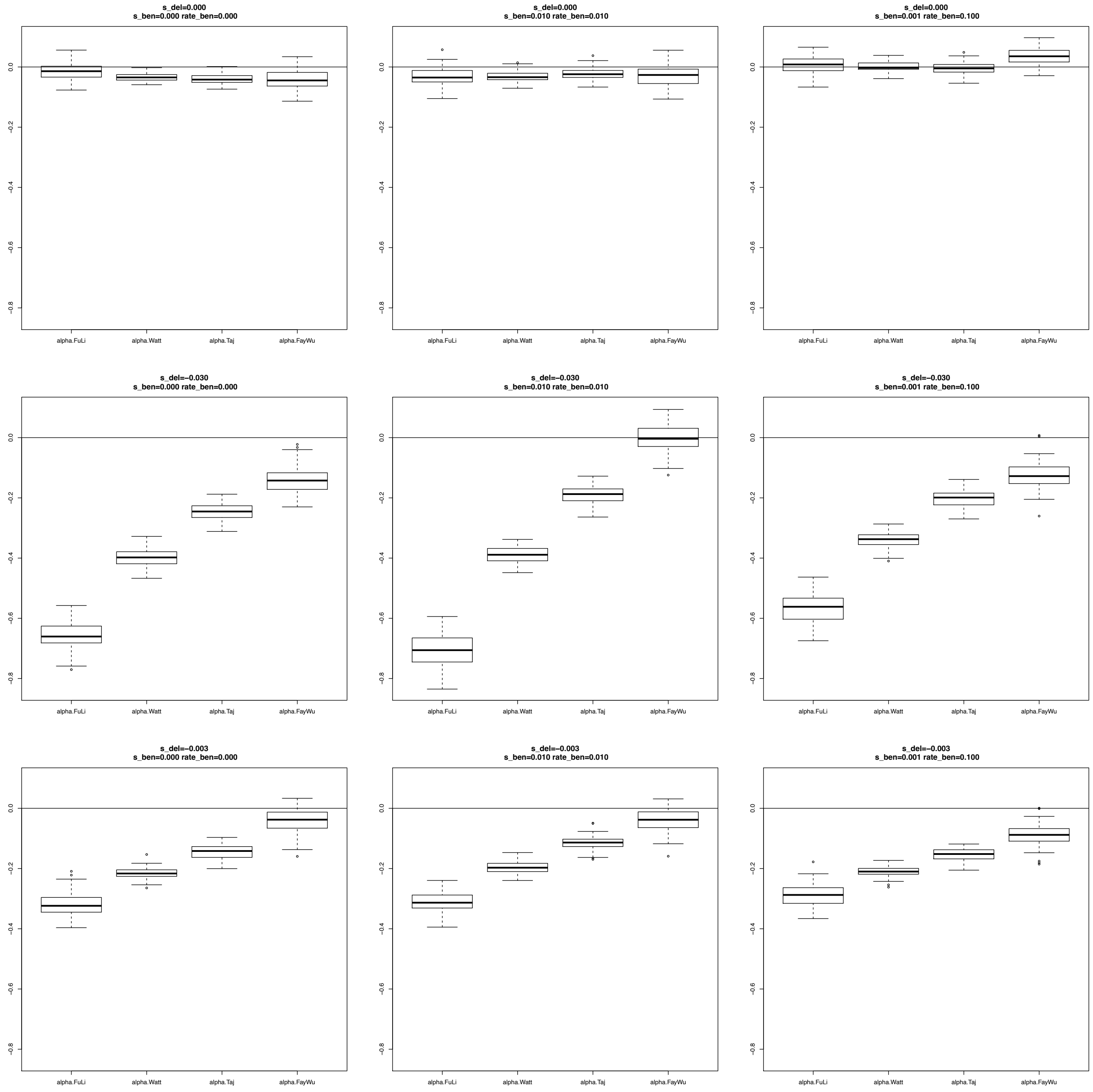

**Figure S11**. Estimates of $\alpha$ considering **Total** SNPs for the Domestic population. A bottleneck was simulated after the split of Wild and Domestic populations (0.1x reduction of population size during 0.1 Ne generations, where Ne is the ancestral population size; see Table S3B for a description of the parameter values used in these simulations). The different scenarios were simulated using different combinations of positive and negative selection coefficients considering **no migration** from Wild to Domestic populations and an increase (**Expansion)** of the population size after bottleneck.

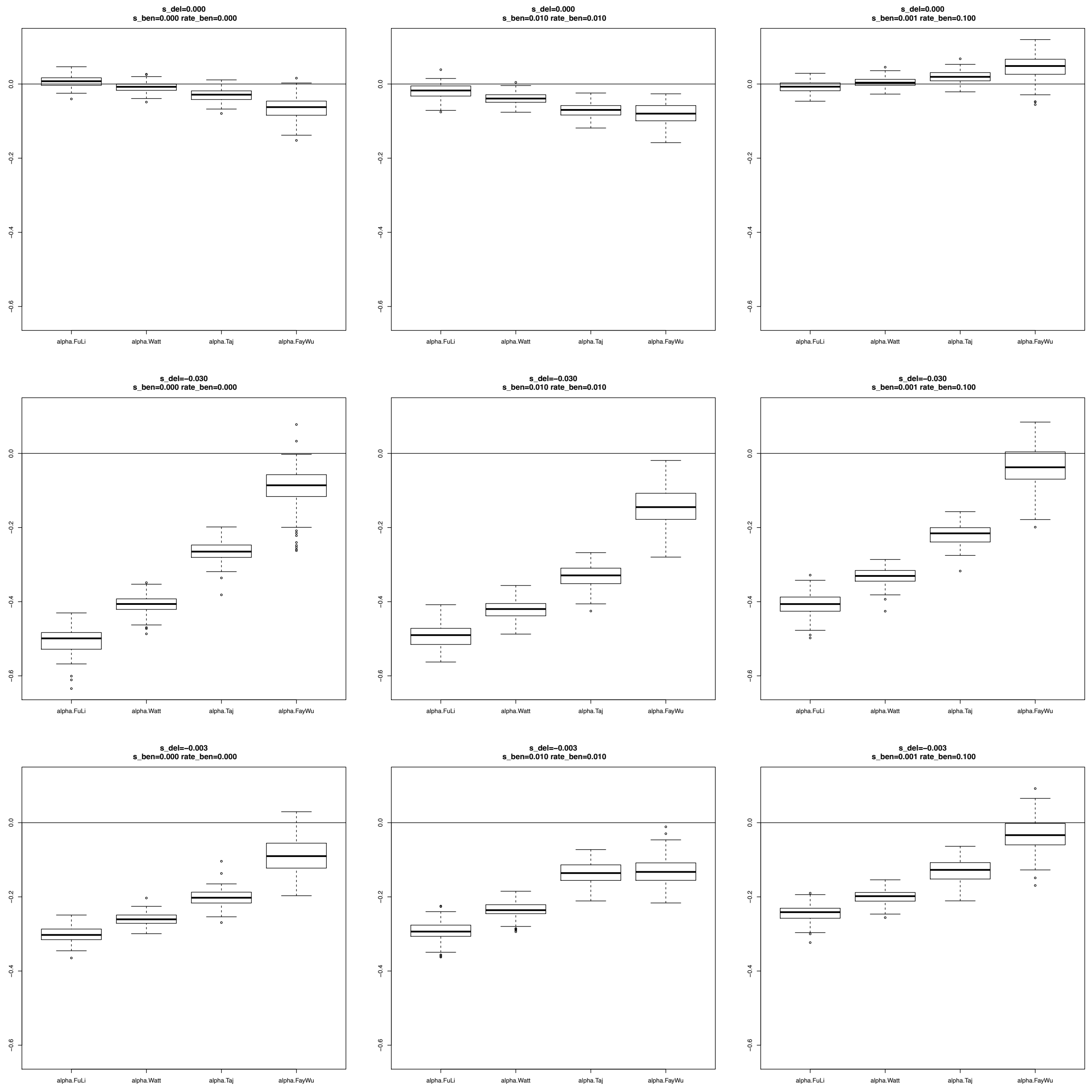

**Figure S12**. Estimates of $\alpha$ considering **Total** SNPs for the Domestic population. A bottleneck was simulated after the split of Wild and Domestic populations (0.1x reduction of population size during 0.1 Ne generations, where Ne is the ancestral population size; see Table S3B for a description of the parameter values used in these simulations). The different scenarios were simulated using different combinations of positive and negative selection coefficients considering **5% migrants** from Wild to Domestic populations and an increase (**Expansion)** of the population size after the bottleneck.

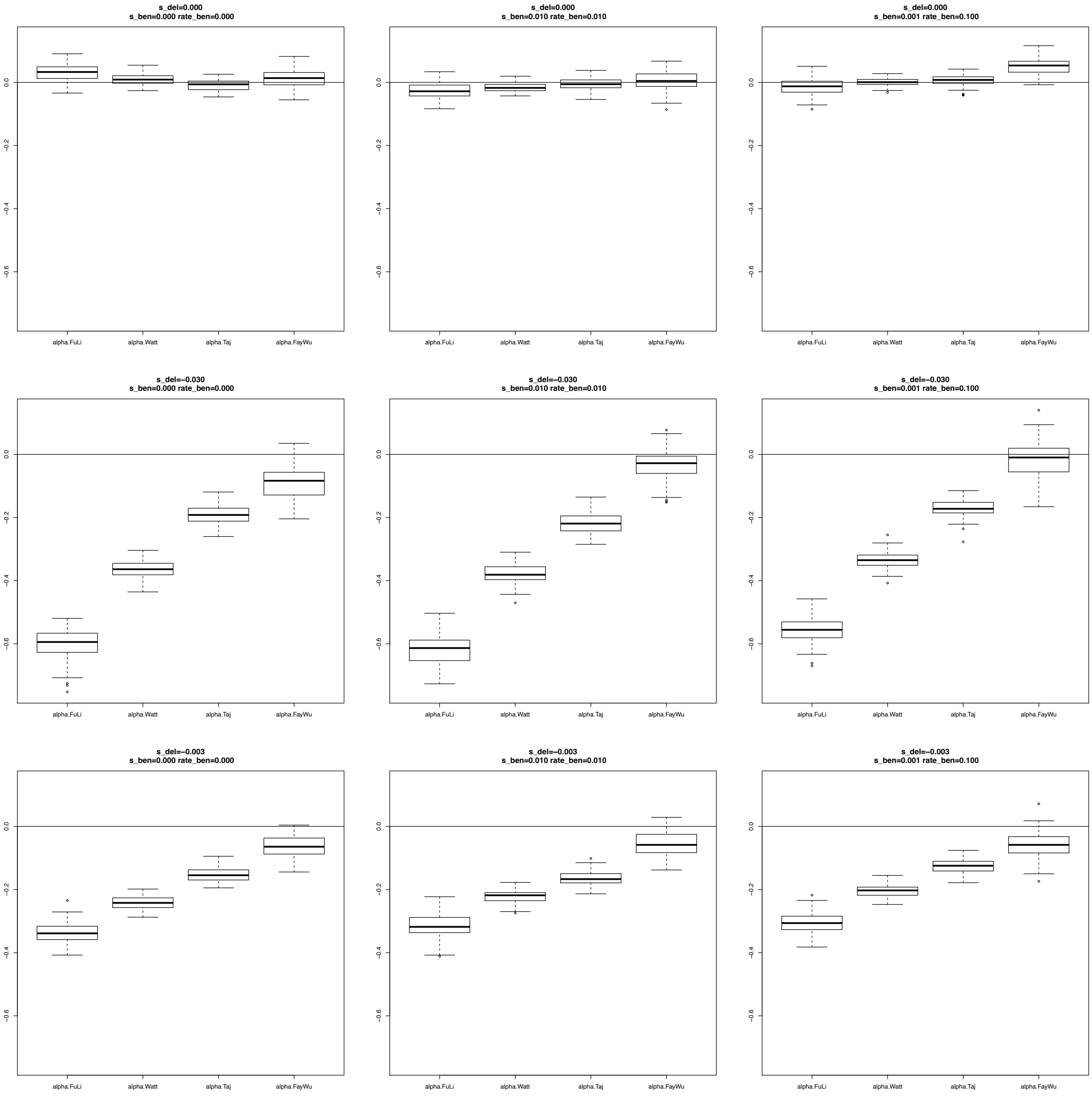

**Figure S13**. Estimates of $\alpha$ considering **Total** SNPs for the Domestic population. A bottleneck was simulated after the split of Wild and Domestic populations (0.1x reduction of population size during 0.1 Ne generations, where Ne is the ancestral population size; see Table S3B for a description of the parameter values used in these simulations). The different scenarios were simulated using different combinations of positive and negative selection coefficients considering **no migration** from Wild to Domestic populations and a decrease (**Reduction)** of the population size after the bottleneck.

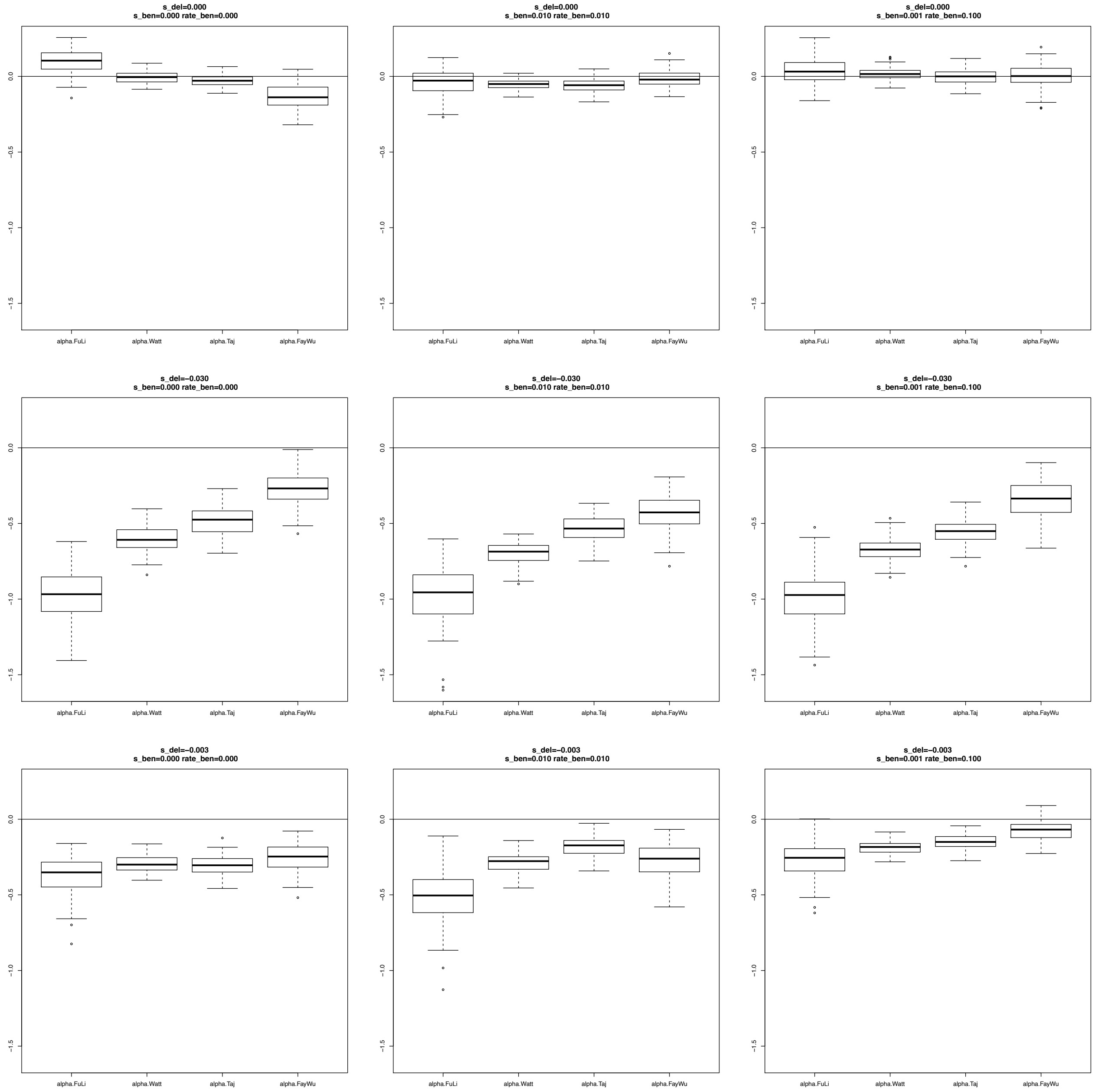

**Figure S14**. Estimates of $\alpha$ considering Total SNPs for the Domestic population. A bottleneck was simulated after the split of Wild and Domestic populations (0.1x reduction of population size during 0.1 Ne generations, where Ne is the ancestral population size; see Table S3B for a description of the parameter values used in these simulations). The different scenarios were simulated using different combinations of positive and negative selection coefficients considering **5% migrants** from Wild to Domestic populations and a decrease (**Reduction** ) of the population size after the bottleneck.

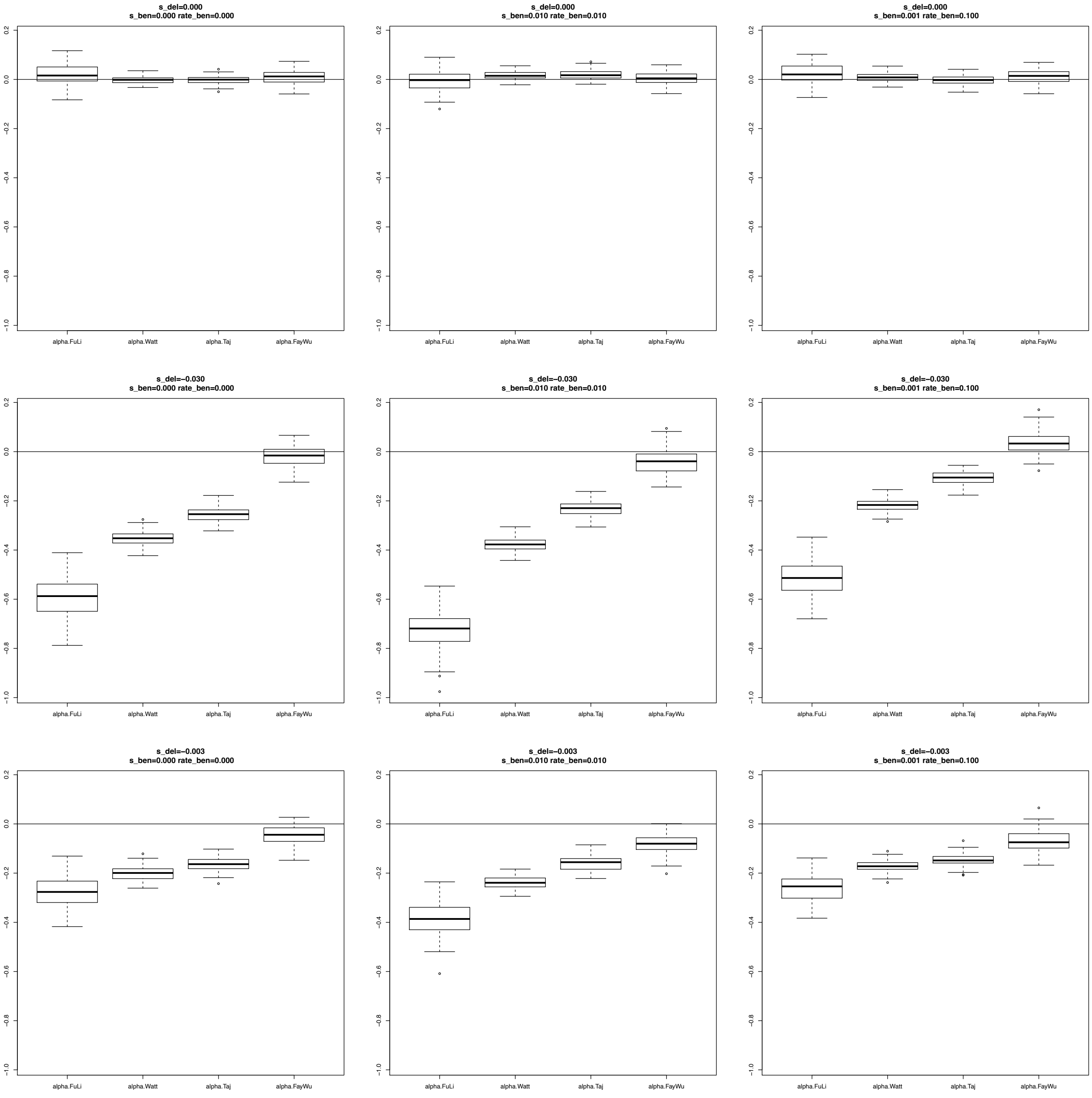

**Figure S15**. Estimates of $R\_\beta\gamma$ considering **Total** SNPs for the Domestic population. A bottleneck was simulated after the split of Wild and Domestic populations (0.1x reduction of population size during 0.1 Ne generations, where Ne is the ancestral population size; see Table S3B for a description of the parameter values used in these simulations). The different scenarios were simulated using different combinations of positive and negative selection coefficients considering **no migration** from Wild to Domestic populations and **no change** in the population size after the bottleneck.

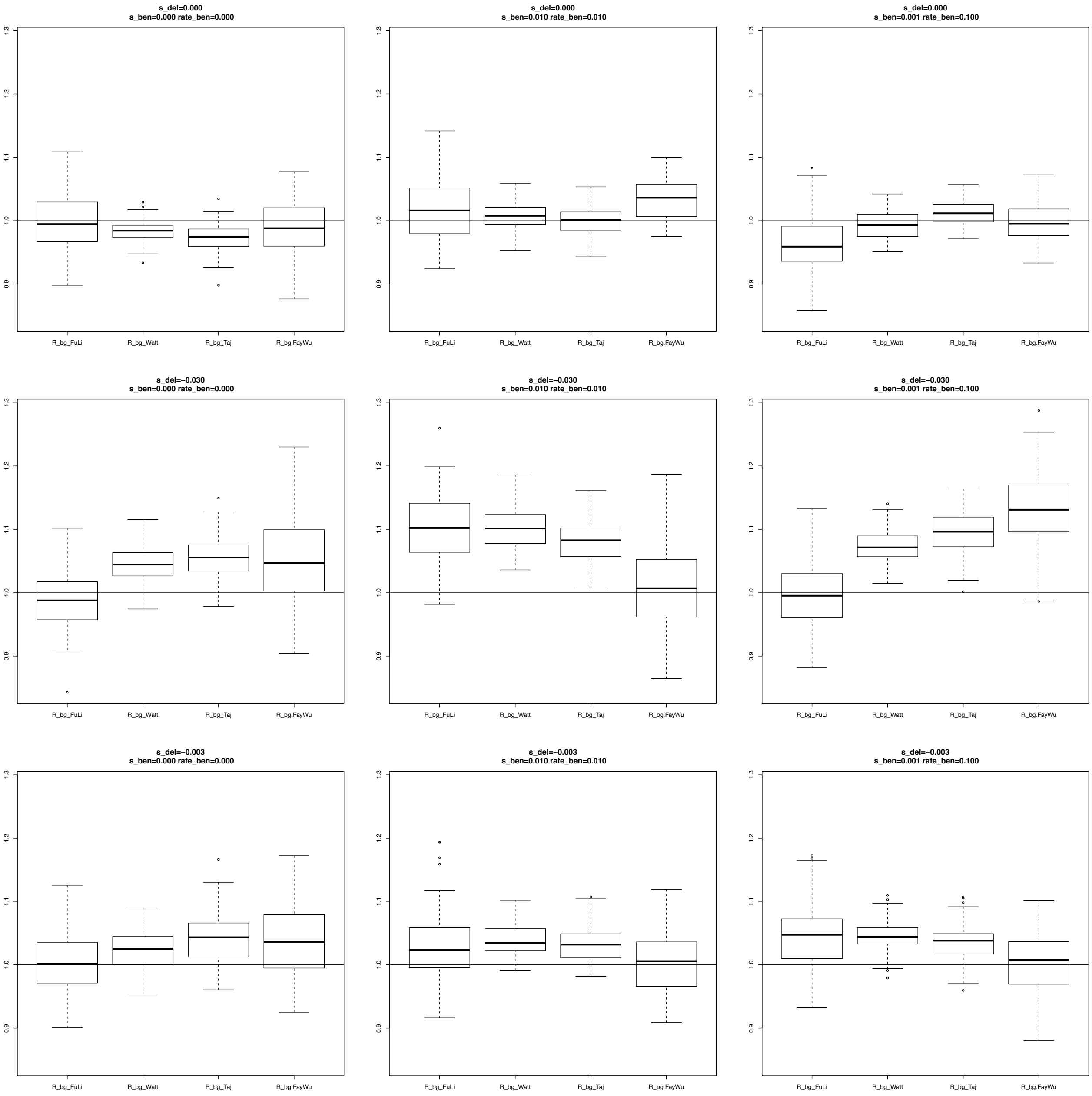

**Figure S16**. Estimates of *R_βγ* considering **Total** SNPs for the Domestic population. A bottleneck was simulated after the split of Wild and Domestic populations (0.1x reduction of population size during 0.1 Ne generations, where Ne is the ancestral population size; see Table S3B for a description of the parameter values used in these simulations). The different scenarios were simulated using different combinations of positive and negative selection coefficients considering **5% migrants** from Wild to Domestic populations and **no change** in the population size after the bottleneck.

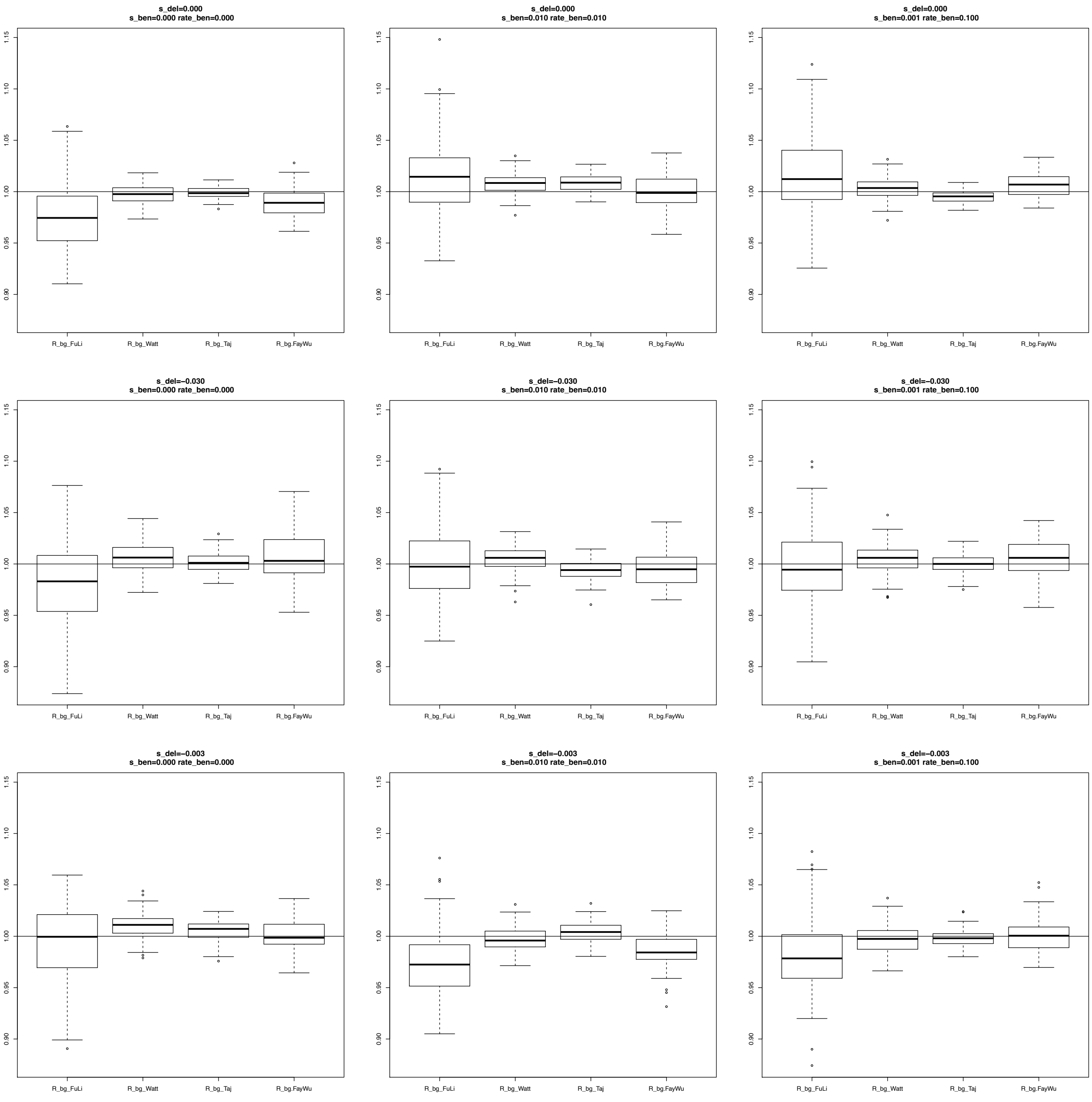

**Figure S17**. Estimates of *R_βγ* considering **Total** SNPs for the Domestic population. A bottleneck was simulated after the split of Wild and Domestic populations (0.1x reduction of population size during 0.1 Ne generations, where Ne is the ancestral population size; see Table S3B for a description of the parameter values used in these simulations). The different scenarios were simulated different combinations of positive and negative selection coefficients considering **no migration** from Wild to Domestic populations and an increase (**Expansion)** of the population size after the bottleneck.

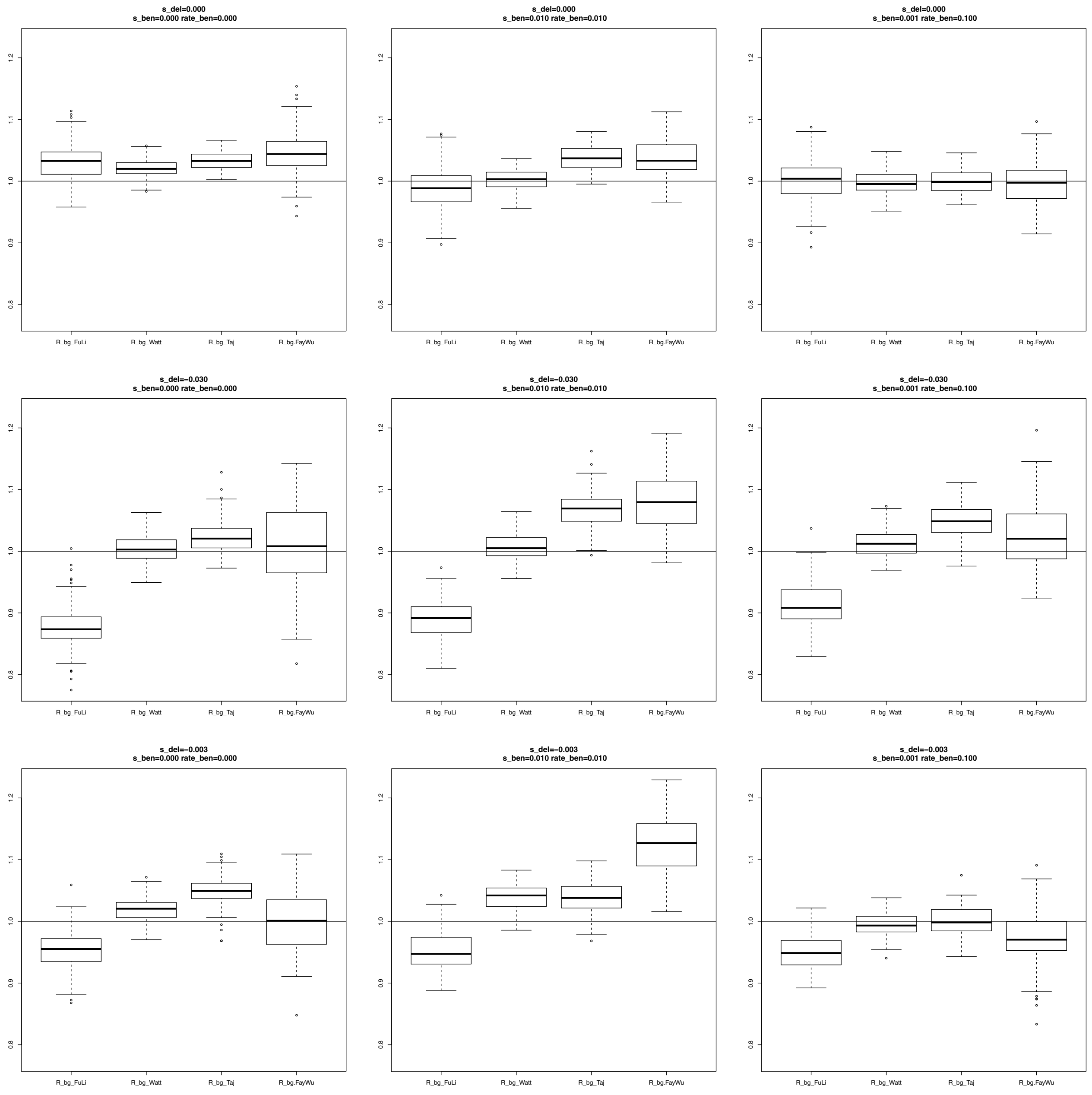

**Figure S18**. Estimates of $R\_\beta\gamma$ considering **Total** SNPs for the Domestic population. A bottleneck was simulated after the split of Wild and Domestic populations (0.1x reduction of population size during 0.1 Ne generations, where Ne is the ancestral population size; see Table S3B for a description of the parameter values used in these simulations). The different scenarios were simulated using different combinations of positive and negative selection coefficients considering **5% migrants** from Wild to Domestic populations and an increase (**Expansion)** of the population size after the bottleneck.

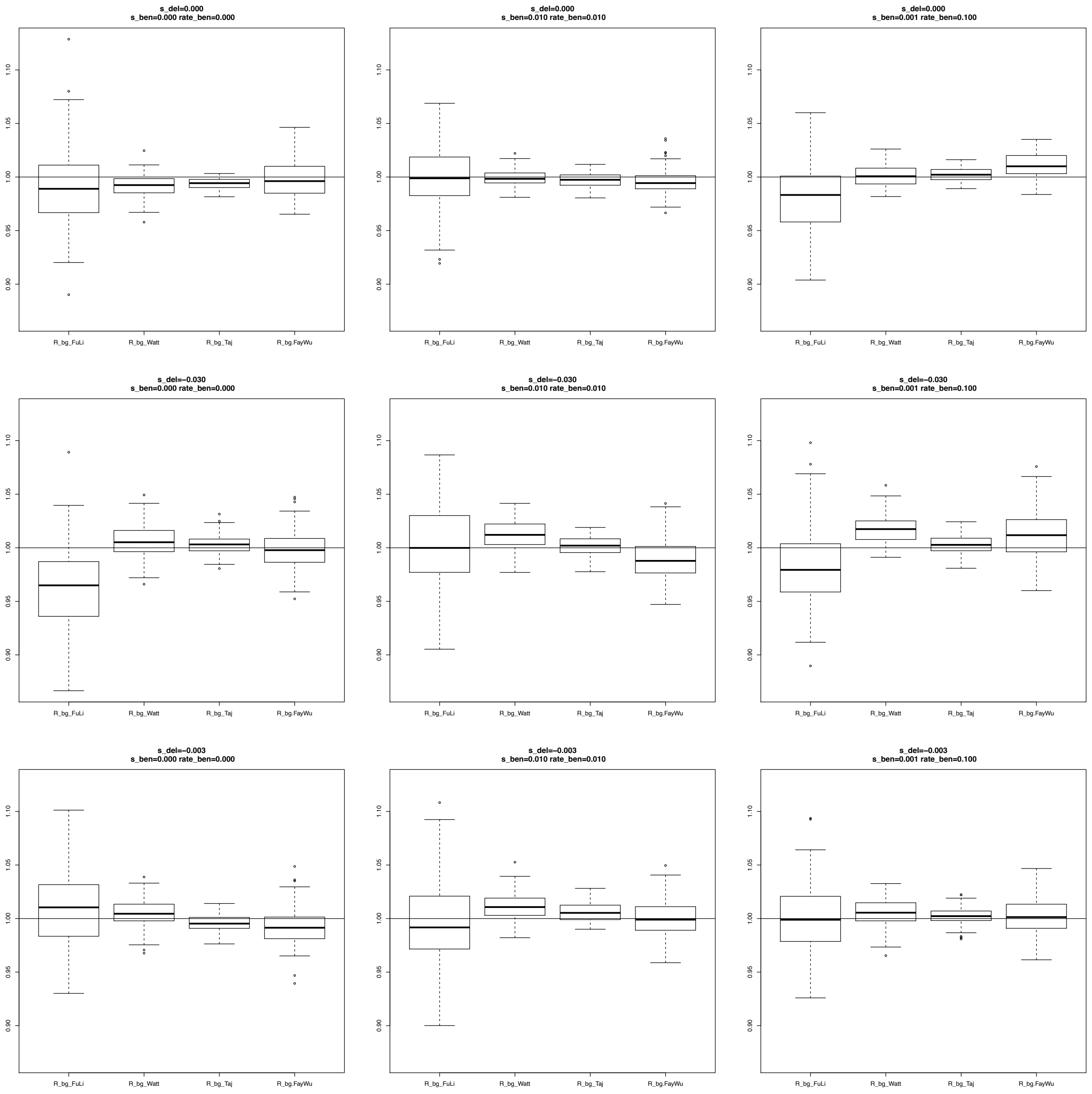

**Figure S19**. Estimates of $R\_\beta\gamma$ considering **Total** SNPs for the Domestic population. A bottleneck was simulated after the split of Wild and Domestic populations (0.1x reduction of population size during 0.1 Ne generations, where Ne is the ancestral population size; see Table S3B for a description of the parameter values used in these simulations). The different scenarios were simulated using different combinations of positive and negative selection coefficients considering **no migration** from Wild to Domestic populations and a decrease (**Reduction)** of the population size after the bottleneck.

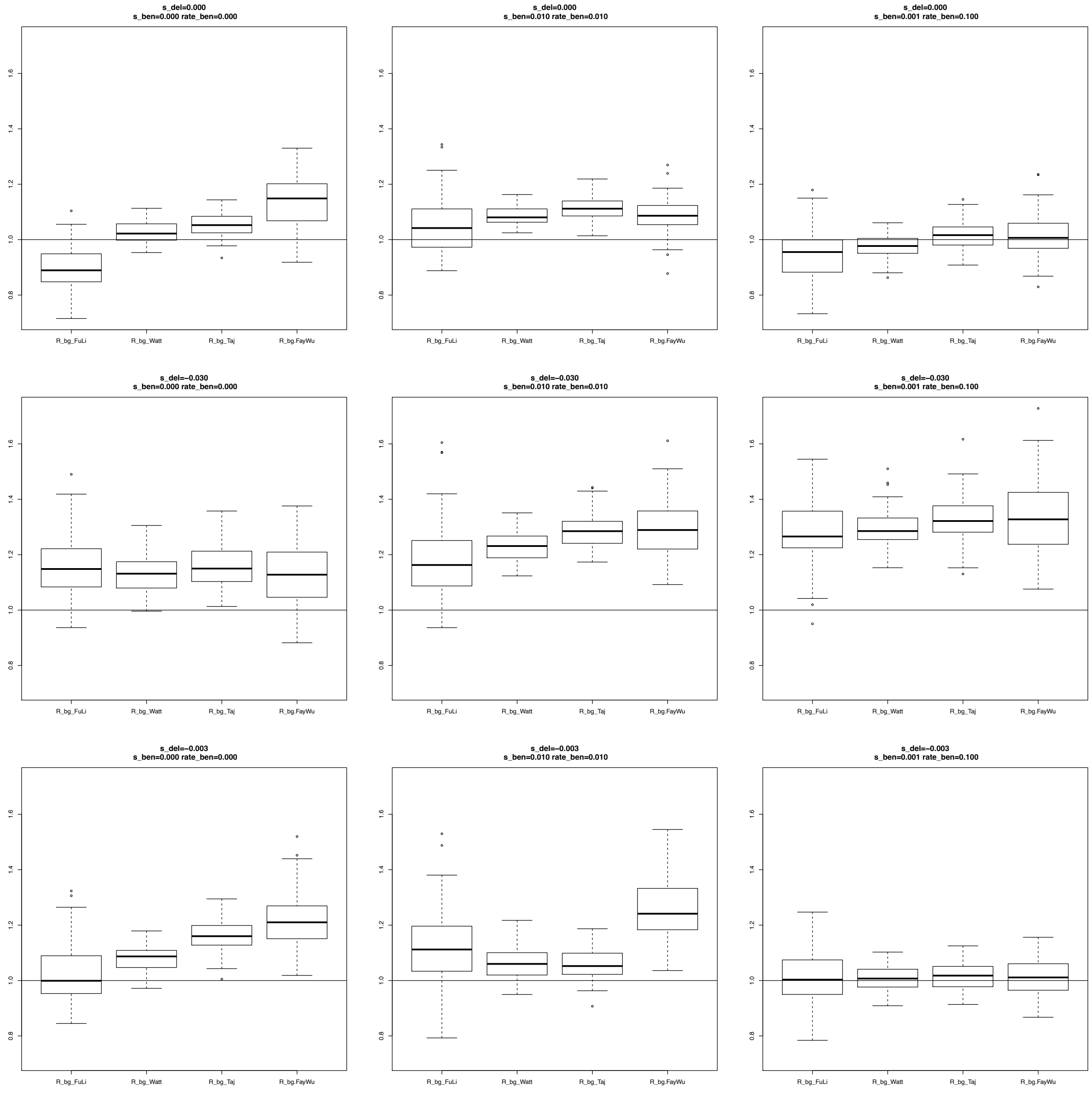

**Figure S20**. Estimates of $R\_\beta\gamma$ considering **Total** SNPs for the Domestic population. A bottleneck was simulated after the split of Wild-Domestic populations (0.1x reduction of population size during 0.1 Ne generations, where Ne is the ancestral population size; see Table S3B for a description of the parameter values used in these simulations). The different scenarios were simulated different combinations of positive and negative selection coefficients considering **5% migrants** from Wild to Domestic populations and a decrease (**Reduction)** of the population size after the bottleneck.

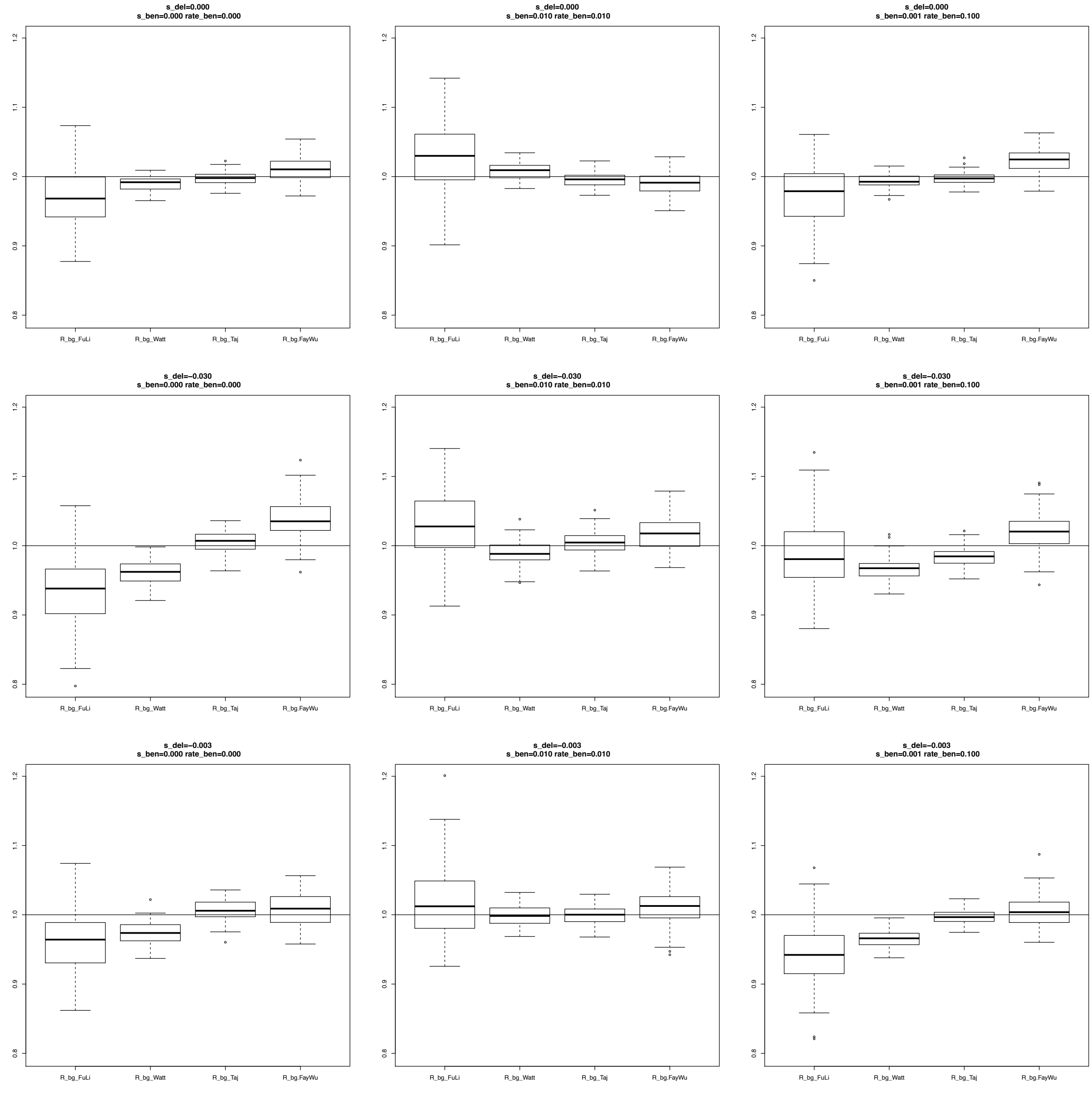

**Figure S21.** Estimates of $\alpha$ using different variability estimates for **Exclusive** SNPs for simulations under the SNM and Negative and Positive selection scenarios.

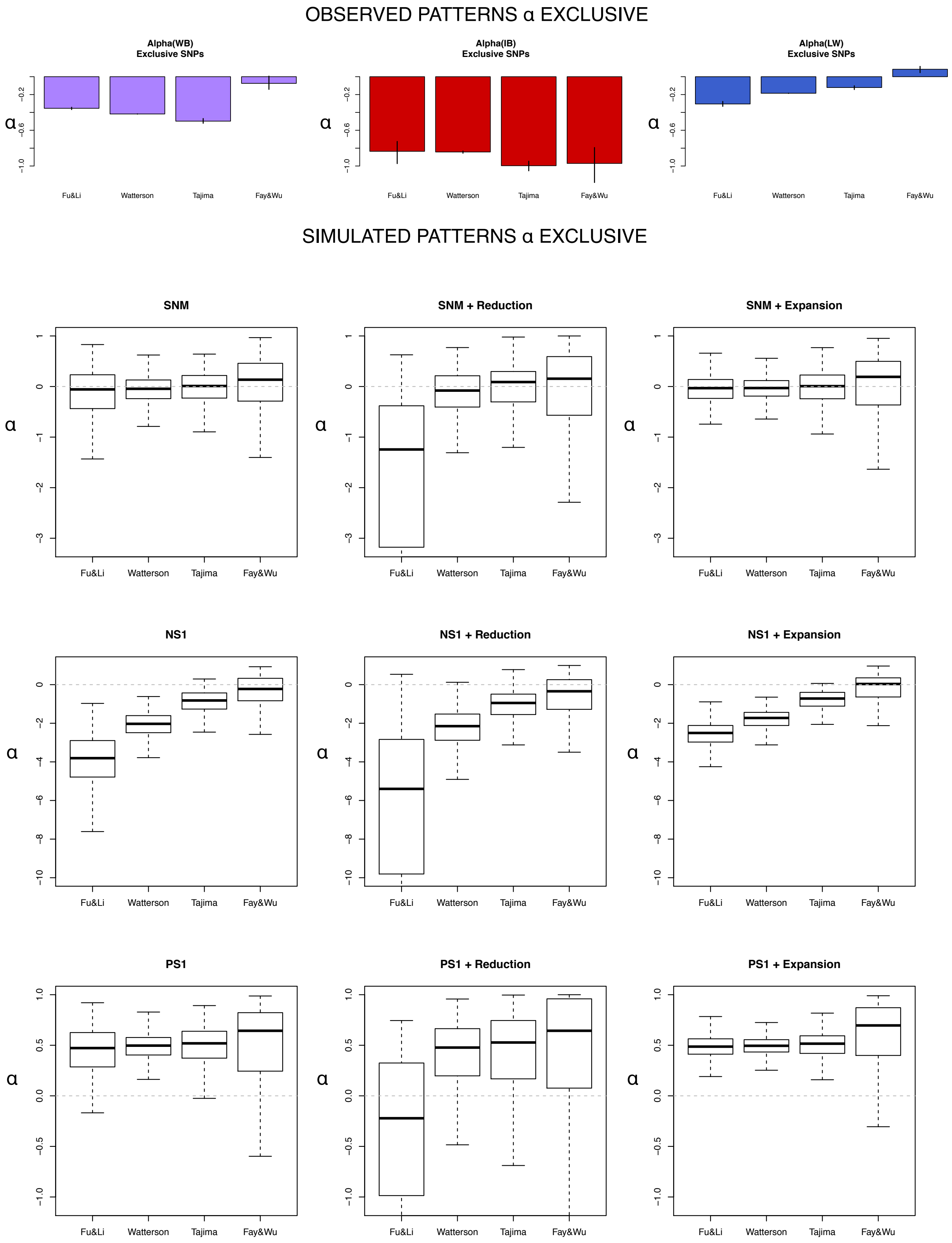

**Figure S22.** Estimates of *R_βγ* using different variability estimates for **Exclusive** SNPs for simulations under the SNM, and Negative and Positive selection scenarios.

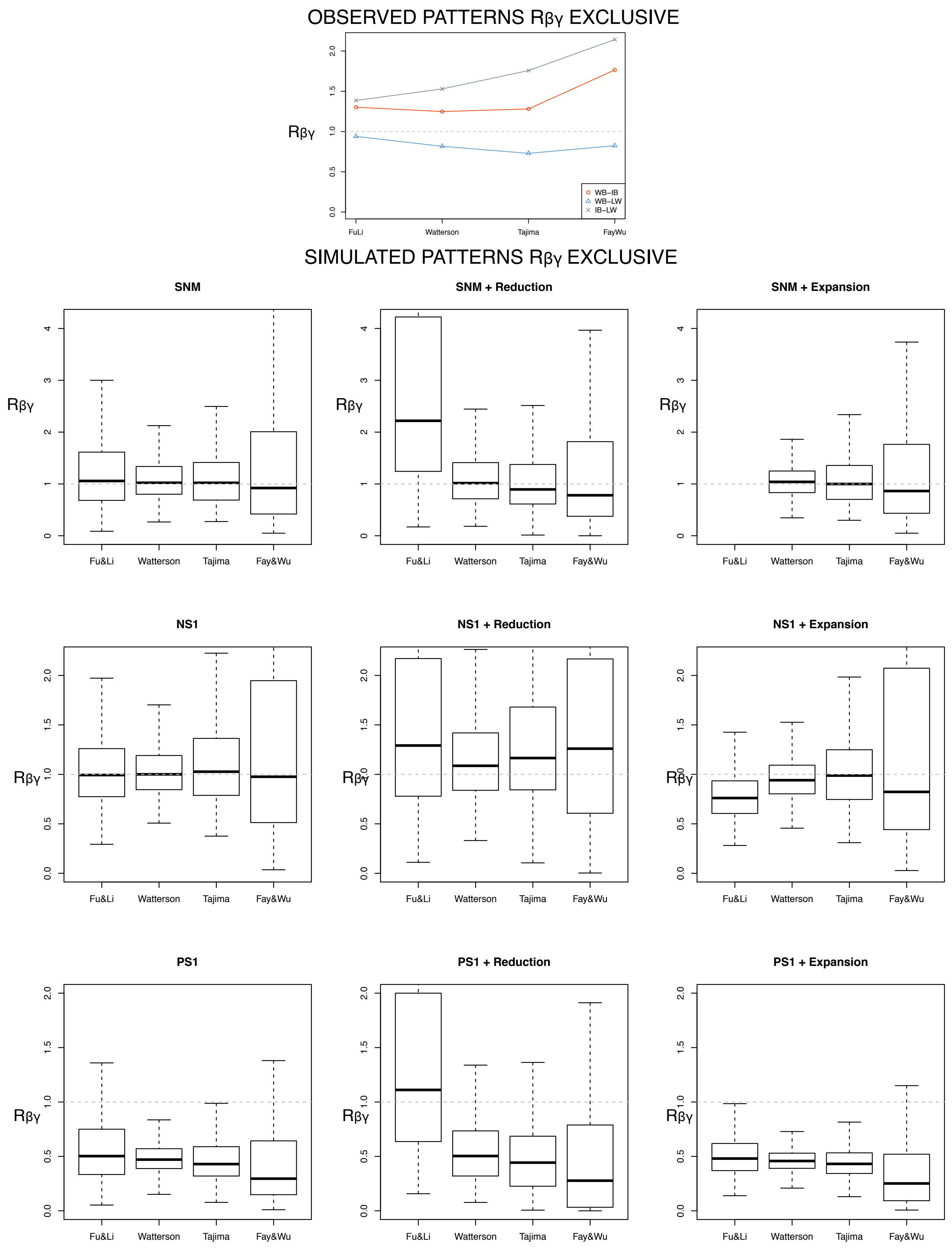

**Figure S23**. Estimates of $\alpha$ considering **Exclusive** SNPs for the Domestic population. A bottleneck was simulated after the split of Wild and Domestic populations (0.1x reduction of population size during 0.1 Ne generations, where Ne is the ancestral population size; see Table S3B for a description of the parameter values used in these simulations). The different scenarios were simulated using different combinations of positive and negative selection coefficients considering **no migration** from Wild to Domestic populations and **no change** in the population size after the bottleneck.

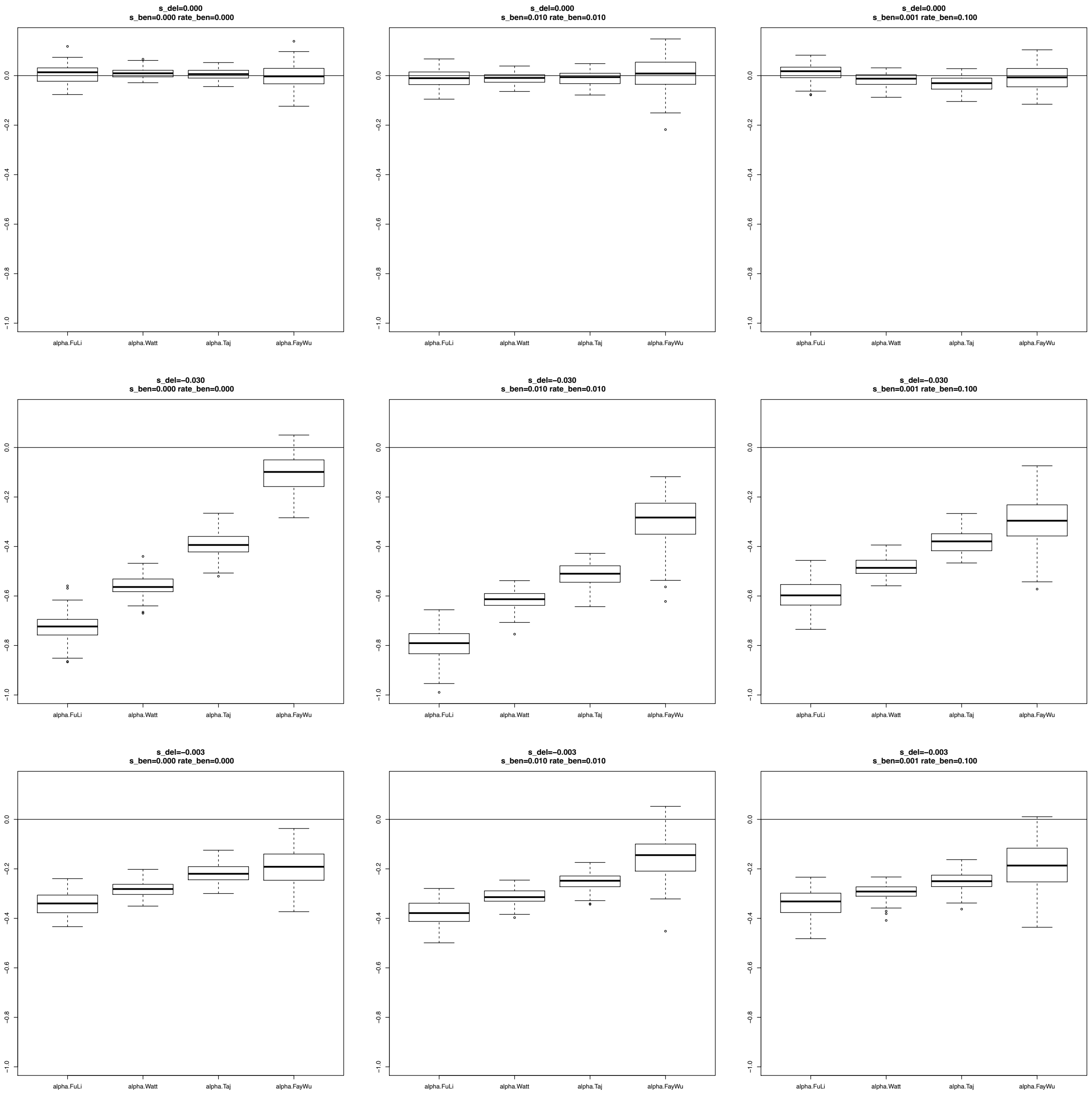

**Figure S24**. Estimates of $\alpha$ considering **Exclusive** SNPs for the Domestic population. A bottleneck was simulated after the split of Wild and Domestic populations (0.1x reduction of population size during 0.1 Ne generations, where Ne is the ancestral population size; see Table S3B for a description of the parameter values used in these simulations). The different scenarios were simulated using different combinations of positive and negative selection coefficients considering **5% migrants** from Wild to Domestic populations and **no change** in the population size after the bottleneck.

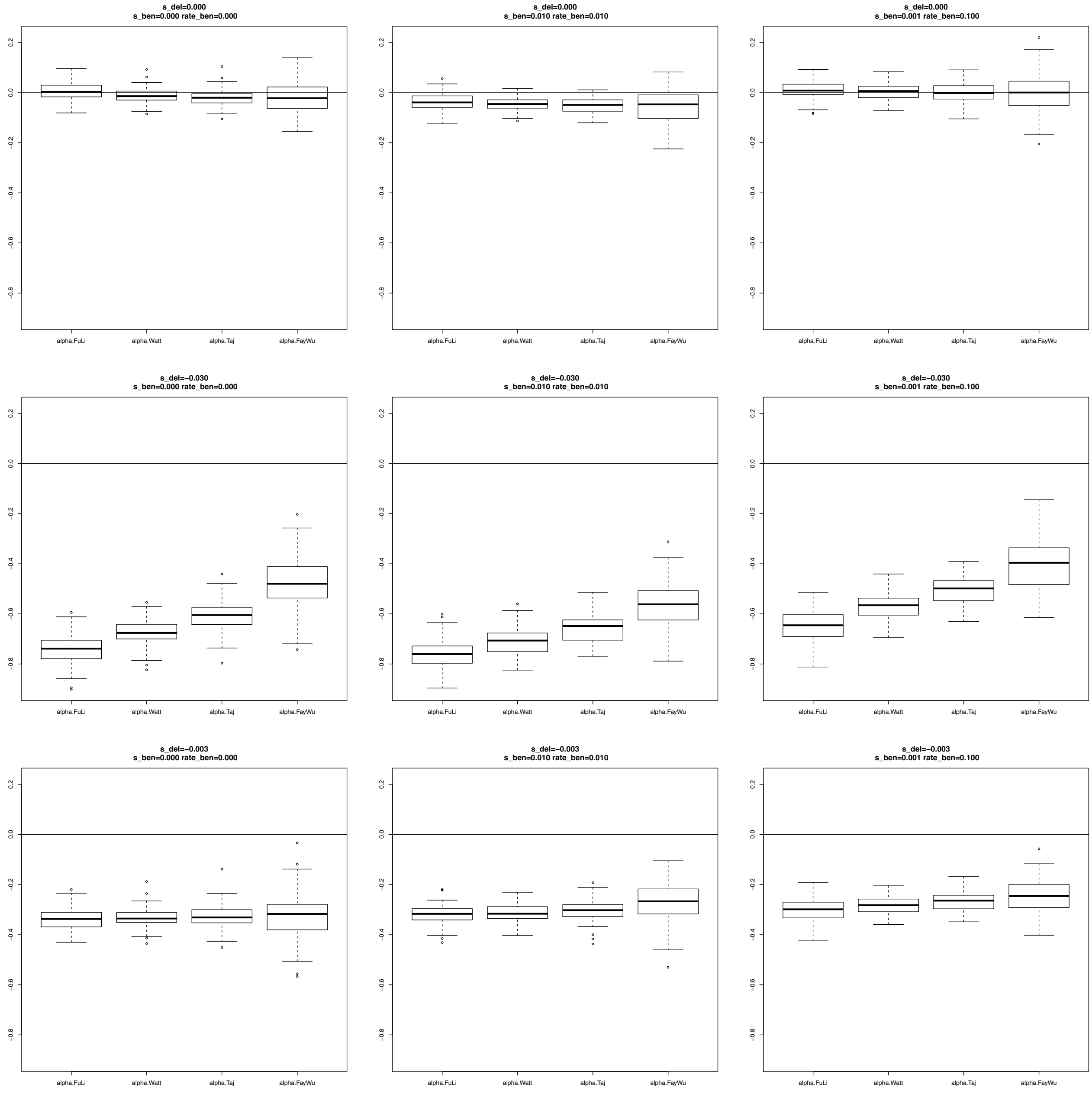

**Figure S25.** Estimates of $\alpha$ considering **Exclusive** SNPs for the Domestic population. A bottleneck was simulated after the split of Wild and Domestic populations (0.1x reduction of population size during 0.1 Ne generations, where Ne is the ancestral population size; see Table S3B for a description of the parameter values used in these simulations). The different scenarios were simulated using different combinations of positive and negative selection coefficients considering **no migration** from Wild to Domestic populations and an increase (**Expansion)** of the population size after the bottleneck.

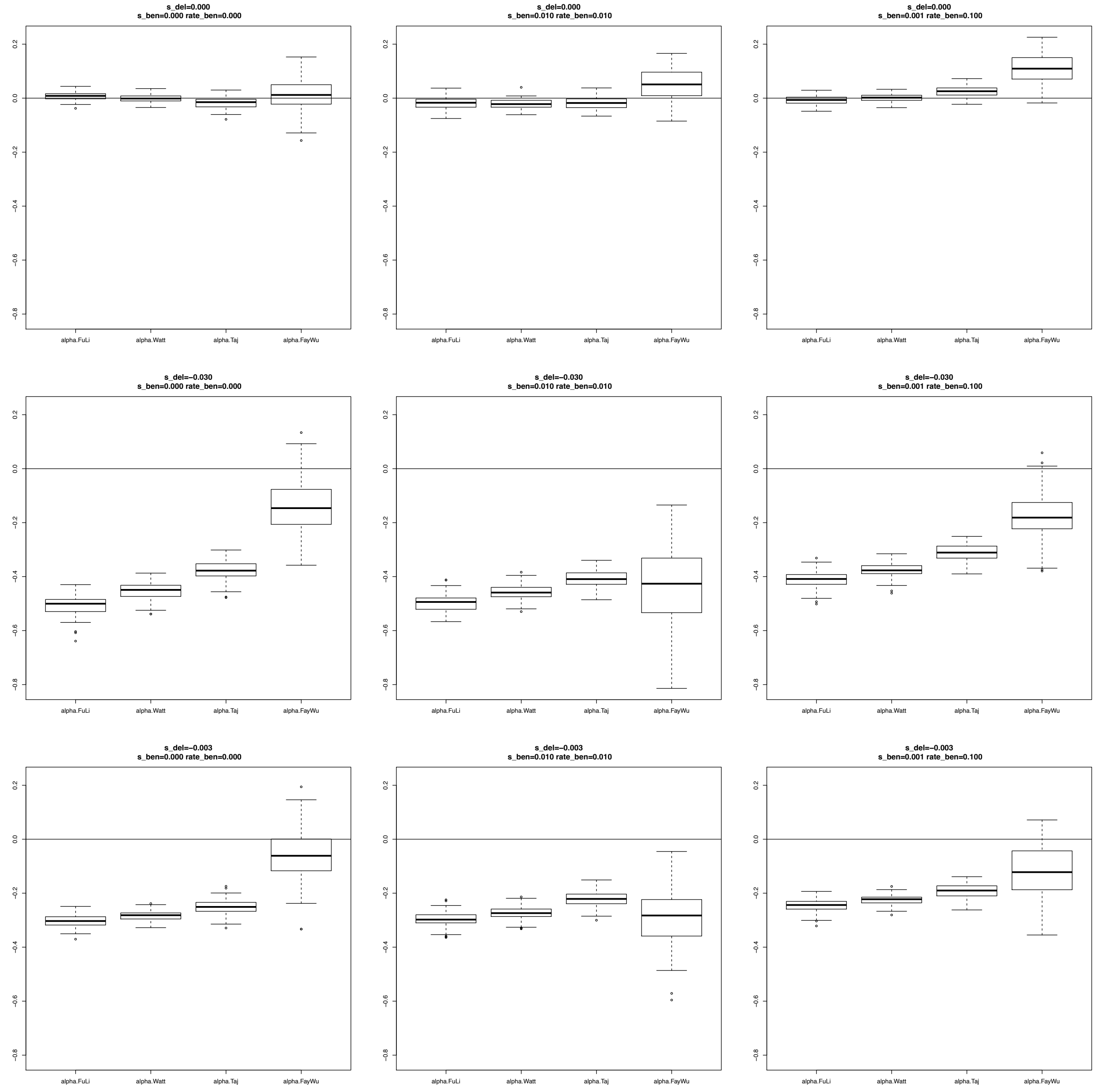

**Figure S26.** Estimates of $\alpha$ considering **Exclusive** SNPs for the Domestic population. A bottleneck was simulated after the split of Wild and Domestic populations (0.1x reduction of population size during 0.1 Ne generations, where Ne is the ancestral population size; see Table S3B for a description of the parameter values used in these simulations). The different scenarios were simulated using different combinations of positive and negative selection coefficients considering **5% migrants** from Wild to Domestic populations and an increase (**Expansion)** of the population size after the bottleneck.

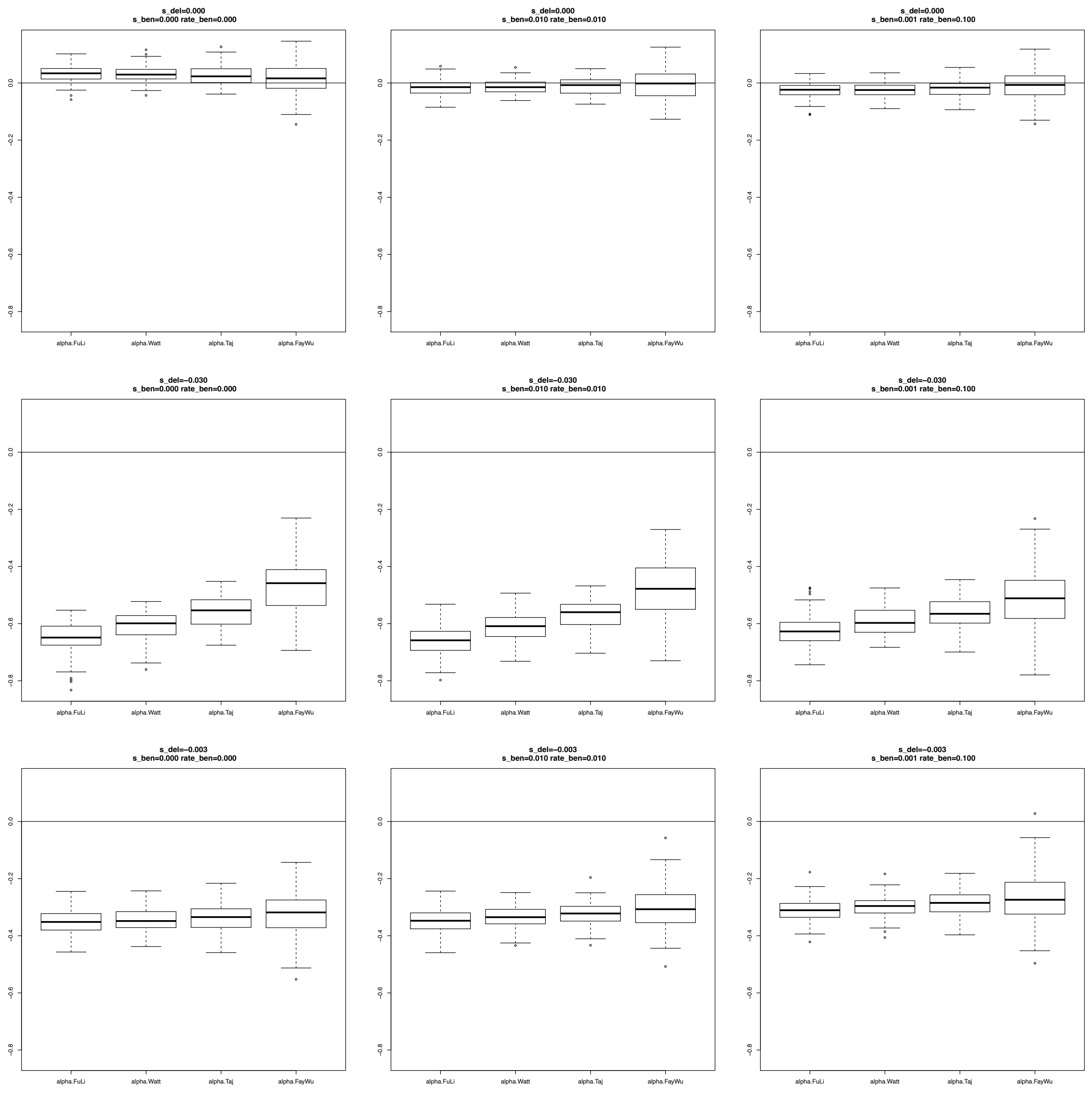

**Figure S27**. Estimates of $\alpha$ considering **Exclusive** SNPs for the Domestic population. A bottleneck was simulated after the split of Wild and Domestic populations (0.1x reduction of population size during 0.1 Ne generations, where Ne is the ancestral population size; see Table S3B for a description of the parameter values used in these simulations). The different scenarios were simulated using different combinations of positive and negative selection coefficients considering **no migration** from Wild to Domestic populations and a decrease (**Reduction)** of the population size after the bottleneck.

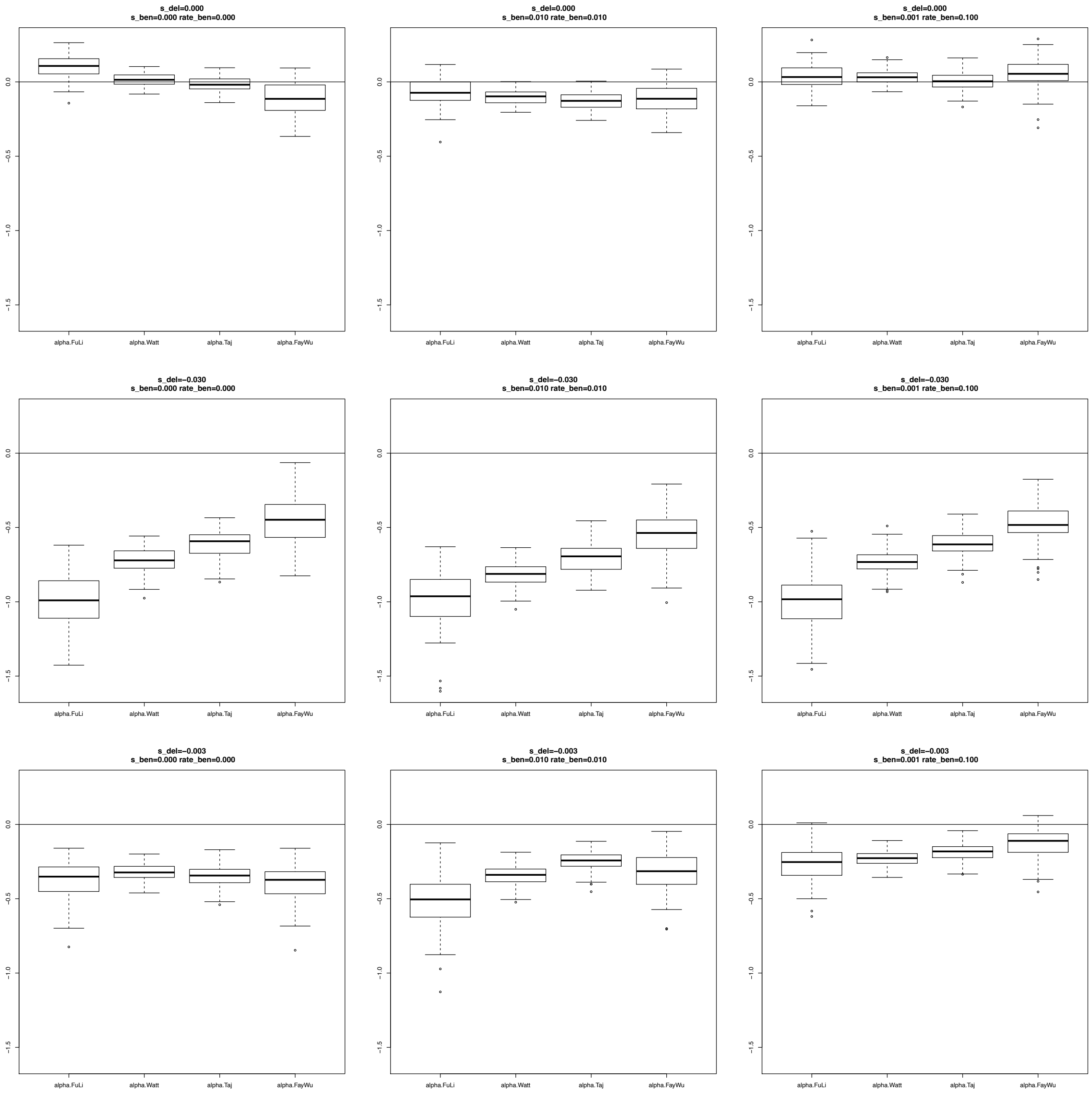

**Figure S28**. Estimates of $\alpha$ considering **Exclusive** SNPs for the Domestic population. A bottleneck was simulated after the split of Wild and Domestic populations (0.1x reduction of population size during 0.1 Ne generations, where Ne is the ancestral population size; see Table S3B for a description of the parameter values used in these simulations). The different scenarios were simulated using different combinations of positive and negative selection coefficients considering **5% migrants** from Wild to Domestic populations and a decrease **(Reduction)** of the population size after the bottleneck.

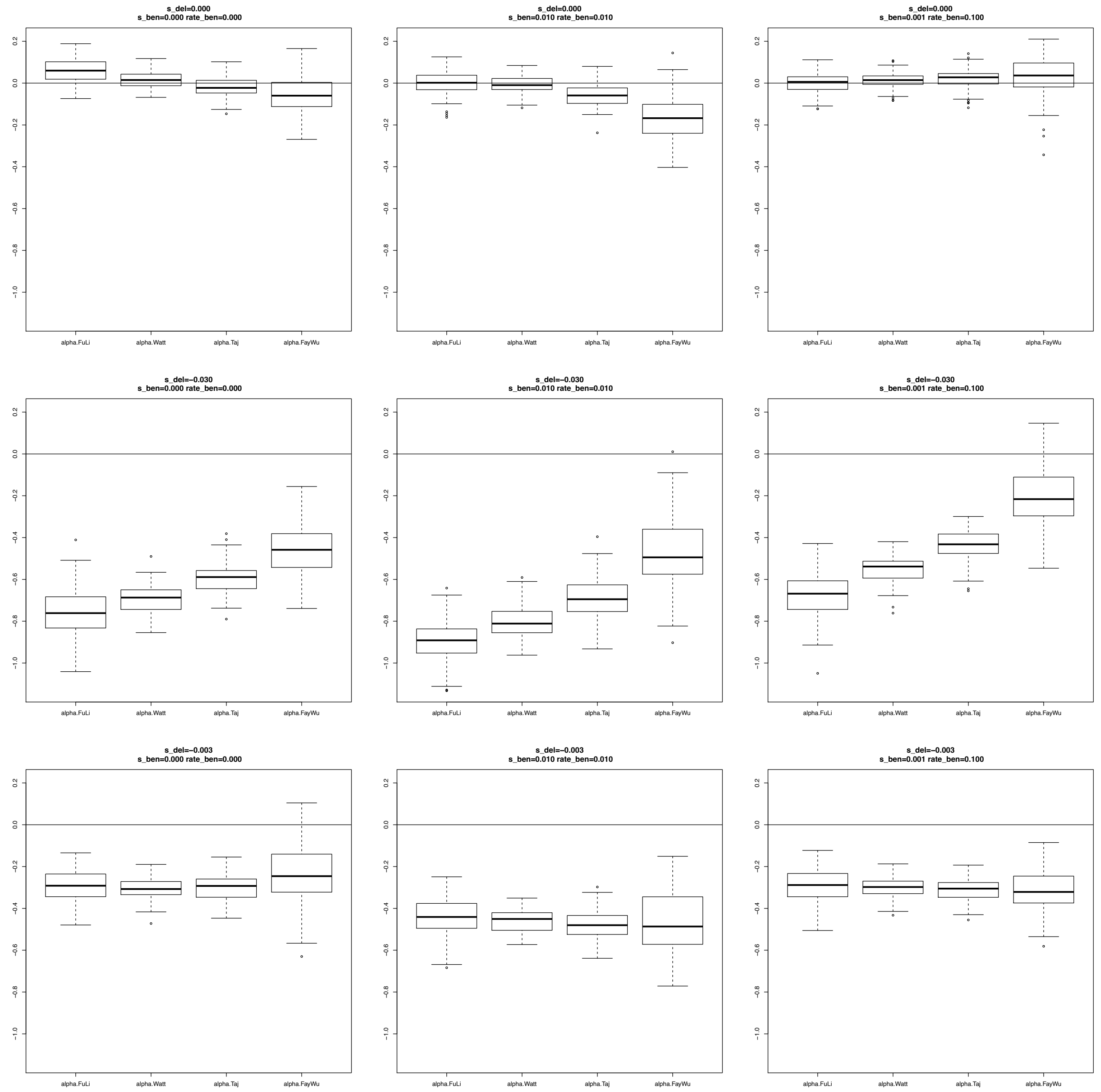

**Figure S29**. Estimates of $R\_\beta\gamma$ considering **Exclusive** SNPs for the Domestic population. A bottleneck was simulated after the split of Wild and Domestic populations (0.1x reduction of population size during 0.1 Ne generations, where Ne is the ancestral population size; see Table S3B for a description of the parameter values used in these simulations). The different scenarios were simulated using different combinations of positive and negative selection coefficients considering **no migration** from Wild to Domestic populations and **no change** in the population size after the bottleneck.

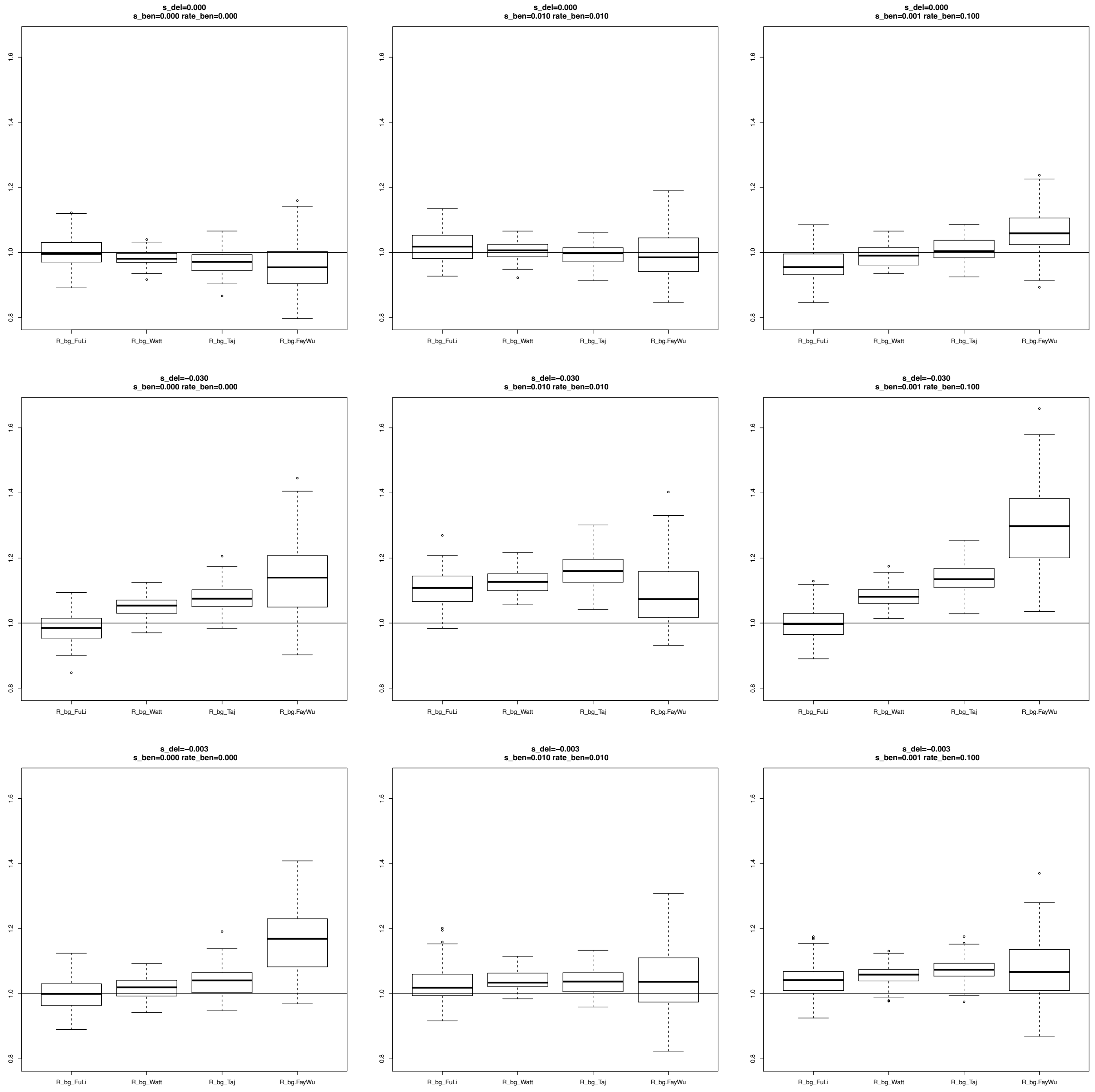

**Figure S30**. Estimates of $R\_\beta\gamma$ considering **Exclusive** SNPs for the Domestic population. A bottleneck was simulated after the split of Wild and Domestic populations (0.1x reduction of population size during 0.1 Ne generations, where Ne is the ancestral population size; see Table S3B for a description of the parameter values used in these simulations). The different scenarios were simulated using different combinations of positive and negative selection coefficients considering **5% migrants** from Wild to Domestic populations and **no change** in the population size after the bottleneck.

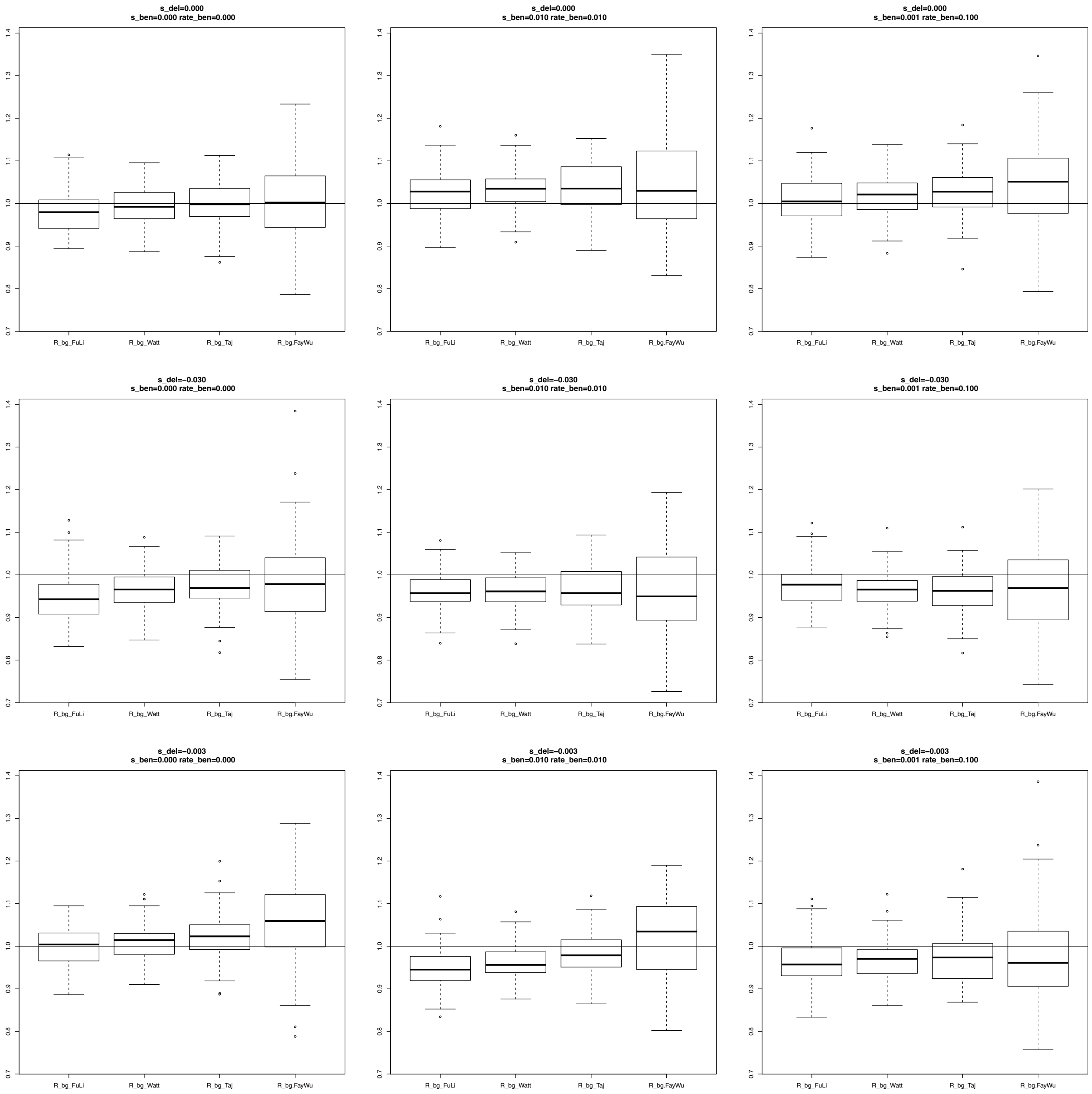

**Figure S31**. Estimates of $R\_\beta\gamma$ considering **Exclusive** SNPs for the Domestic population. A bottleneck was simulated after the split of Wild and Domestic populations (0.1x reduction of population size during 0.1 Ne generations, where Ne is the ancestral population size; see Table S3B for a description of the parameter values used in these simulations). The different scenarios were simulated using different combinations of positive and negative selection coefficients considering **no migration** from Wild to Domestic populations and an increase (**Expansion)** of the population size after the bottleneck.

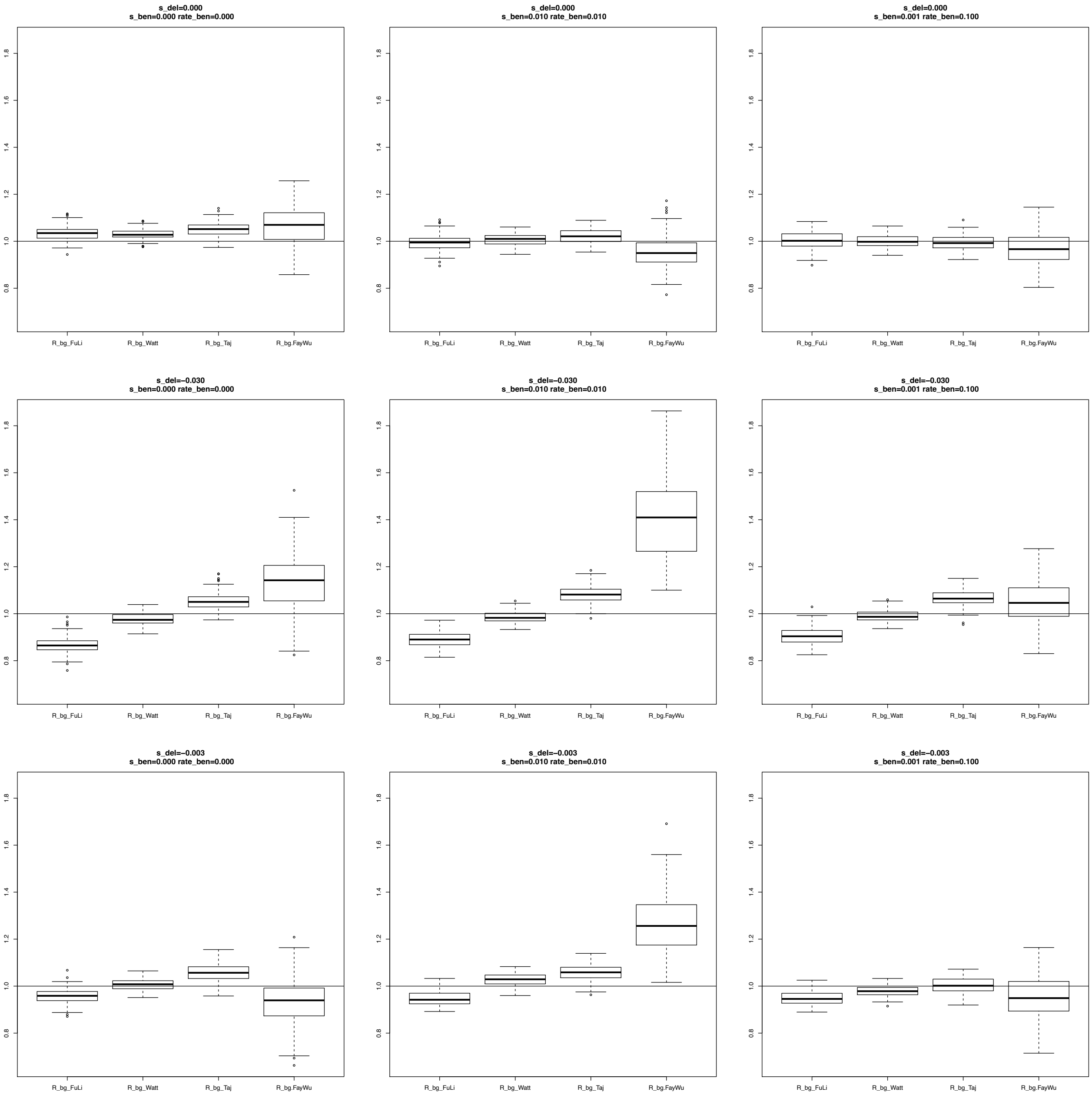

***F*****igure S32**. Estimates of $R\_\beta\gamma$ considering **Exclusive** SNPs for the Domestic population. A bottleneck was simulated after the split of Wild and Domestic populations (0.1x reduction of population size during 0.1 Ne generations, where Ne is the ancestral population size; see Table S3B for a description of the parameter values used in these simulations). The different scenarios were simulated using different combinations of positive and negative selection coefficients considering **5% migrants** from Wild to Domestic populations and an increase (**Expansion)** of the population size after the bottleneck.

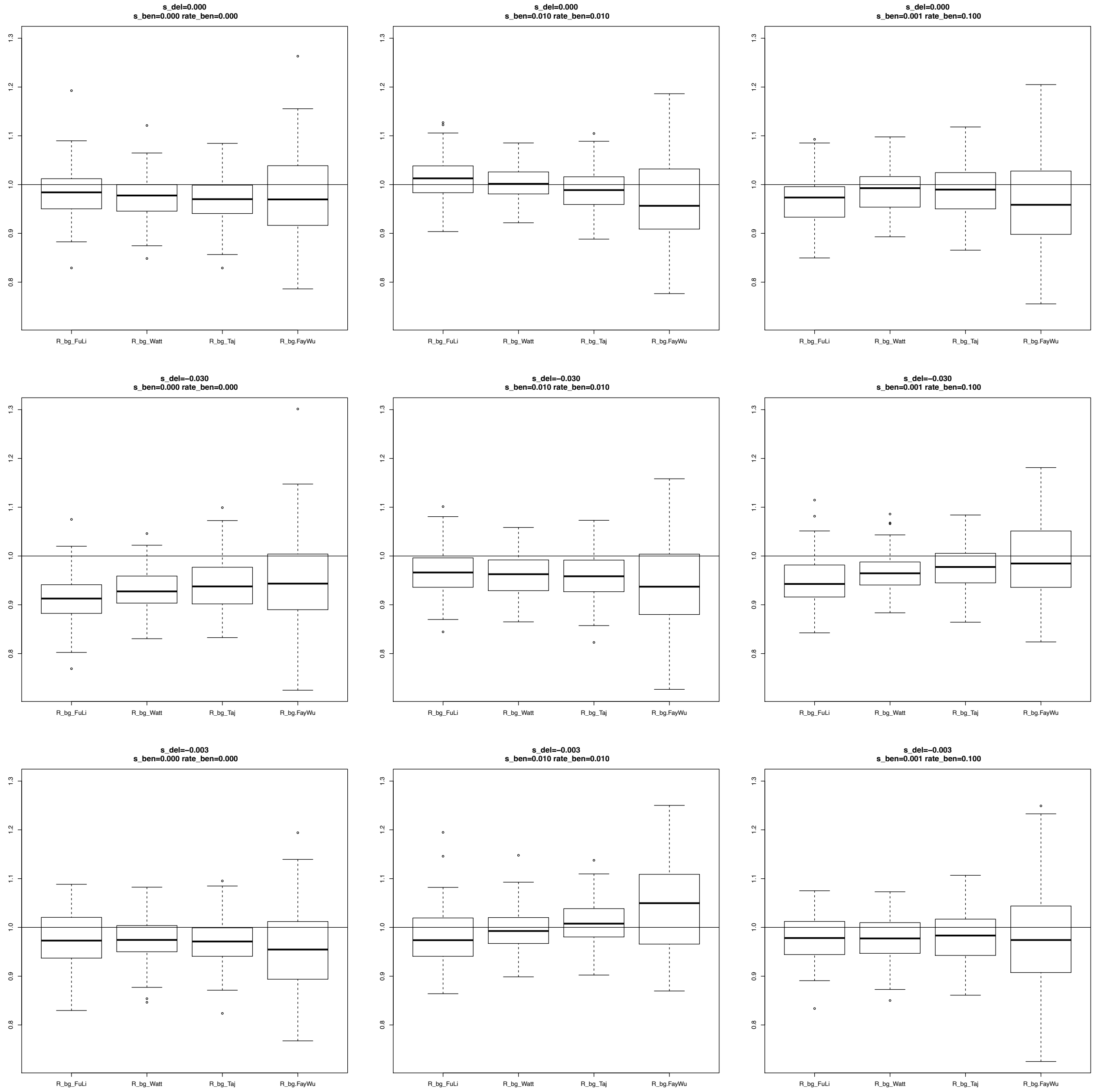

**Figure S33**. Estimates of $R\_\beta\gamma$ considering **Exclusive** SNPs for the Domestic population. A bottleneck was simulated after the split of Wild and Domestic populations (0.1x reduction of population size during 0.1 Ne generations, where Ne is the ancestral population size; see Table S3B for a description of the parameter values used in these simulations). The different scenarios were simulated using different combinations of positive and negative selection coefficients considering **no migration** from Wild to Domestic populations and a decrease (**Reduction)** of the population size after the bottleneck.

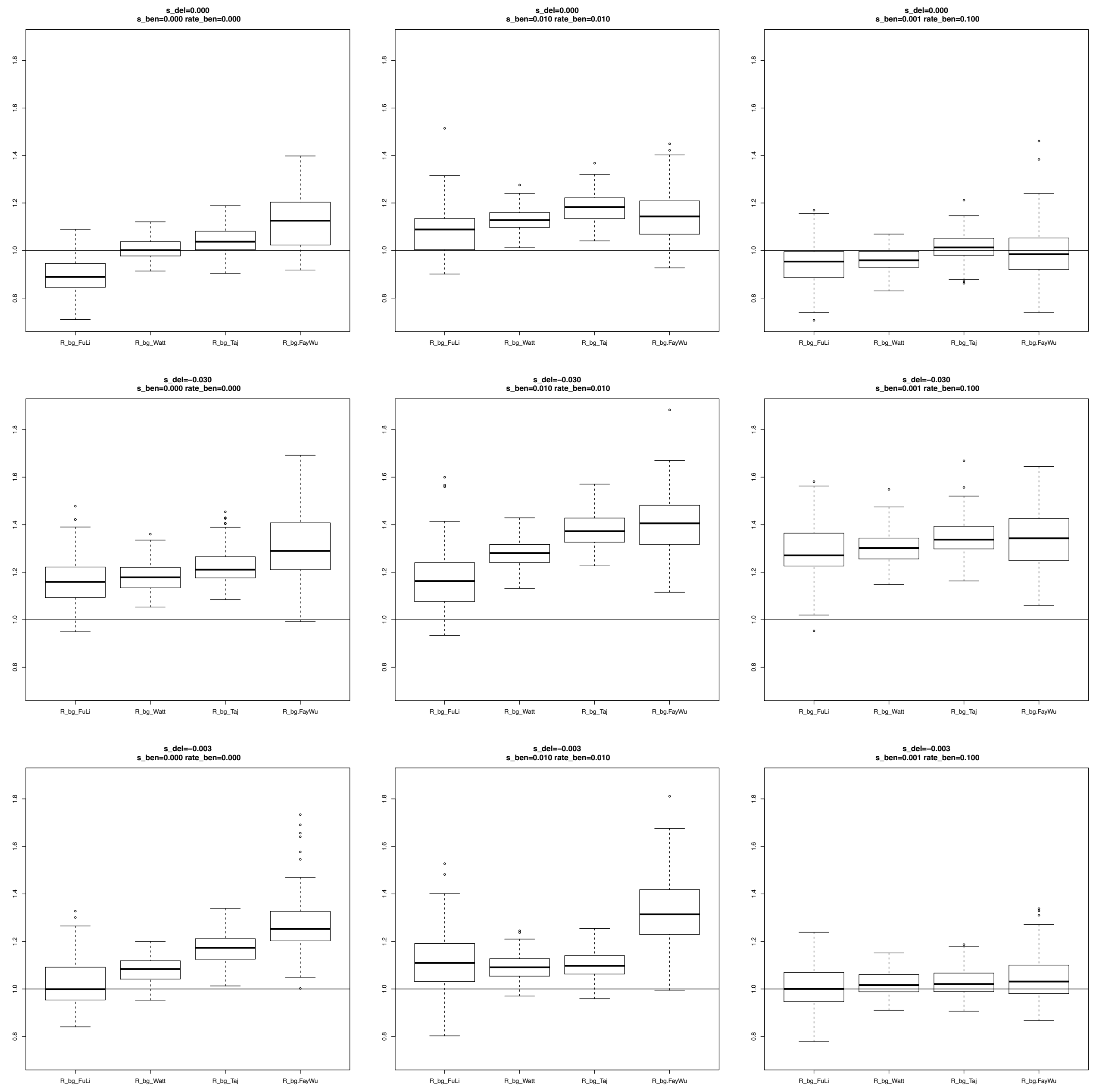

**Figure S34**. Estimates of $R\_\beta\gamma$ considering **Exclusive** SNPs for the Domestic population. A bottleneck was simulated after the split of Wild and Domestic populations (0.1x reduction of population size during 0.1 Ne generations, where Ne is the ancestral population size; see Table S3B for a description of the parameter values used in these simulations). The different scenarios were simulated using different combinations of positive and negative selection coefficients considering **5% migrants** from Wild to Domestic populations and a decrease (**Reduction)** of the population size after the bottleneck.

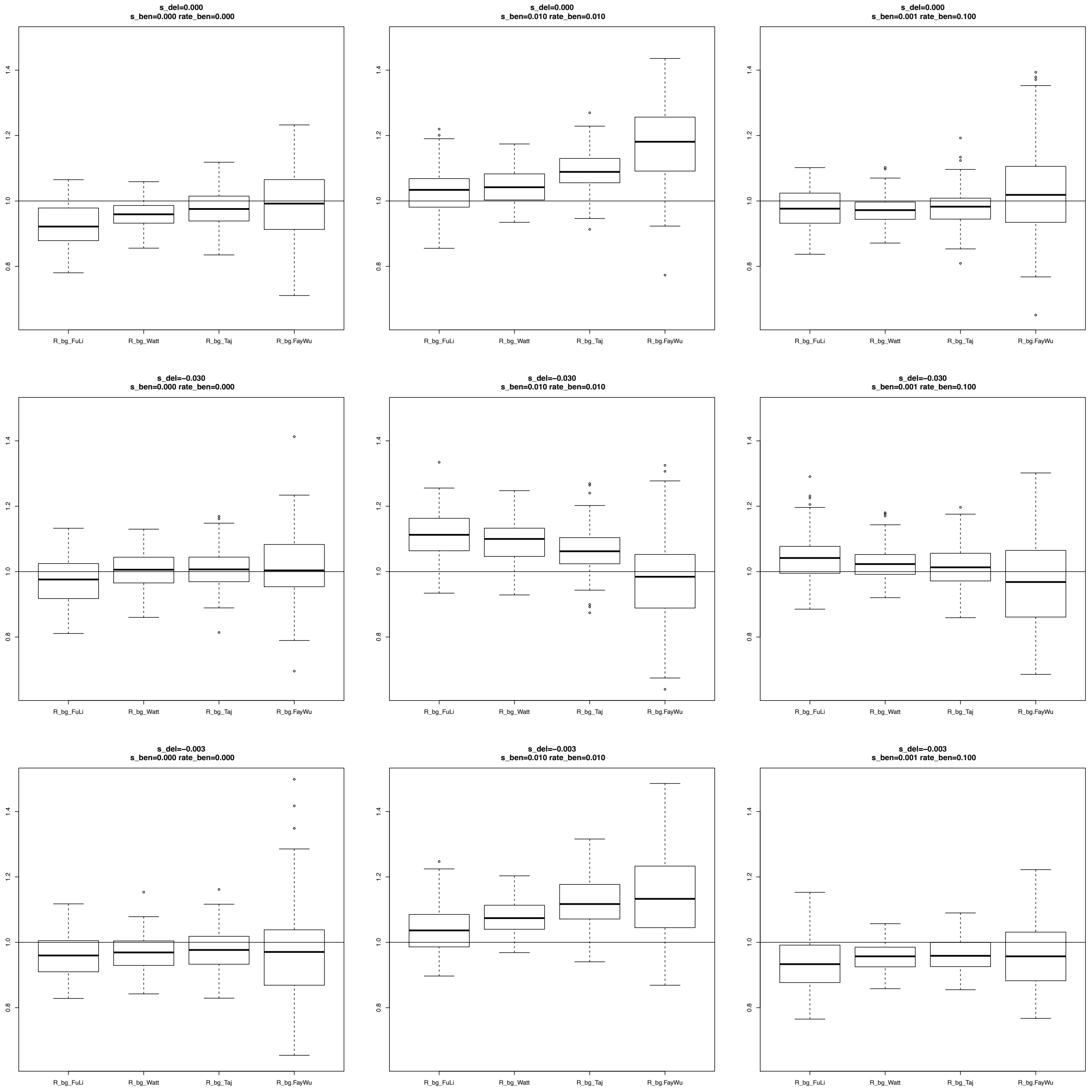

**Figure S35.** Estimates of $\alpha$ using different variability estimates for **Shared** SNPs for simulations under the SNM and under Negative and Positive selection scenarios.

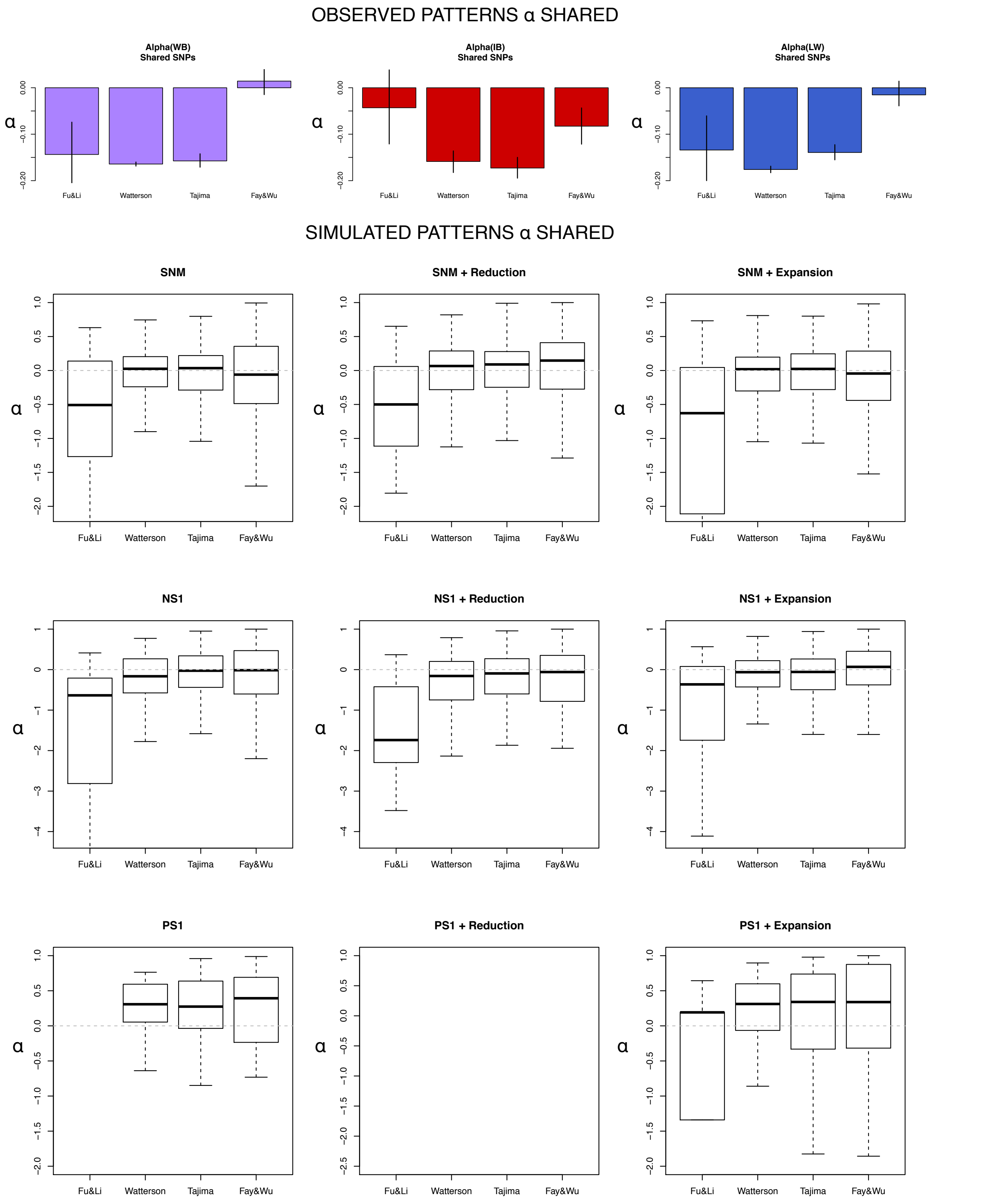

**Figure S36.** Estimates of *R_βγ* using different variability estimates for **Exclusive** SNPs for simulations under the SNM and Negative and Positive selection scenarios. Blank plots indicates that not information was obtained from simulations.

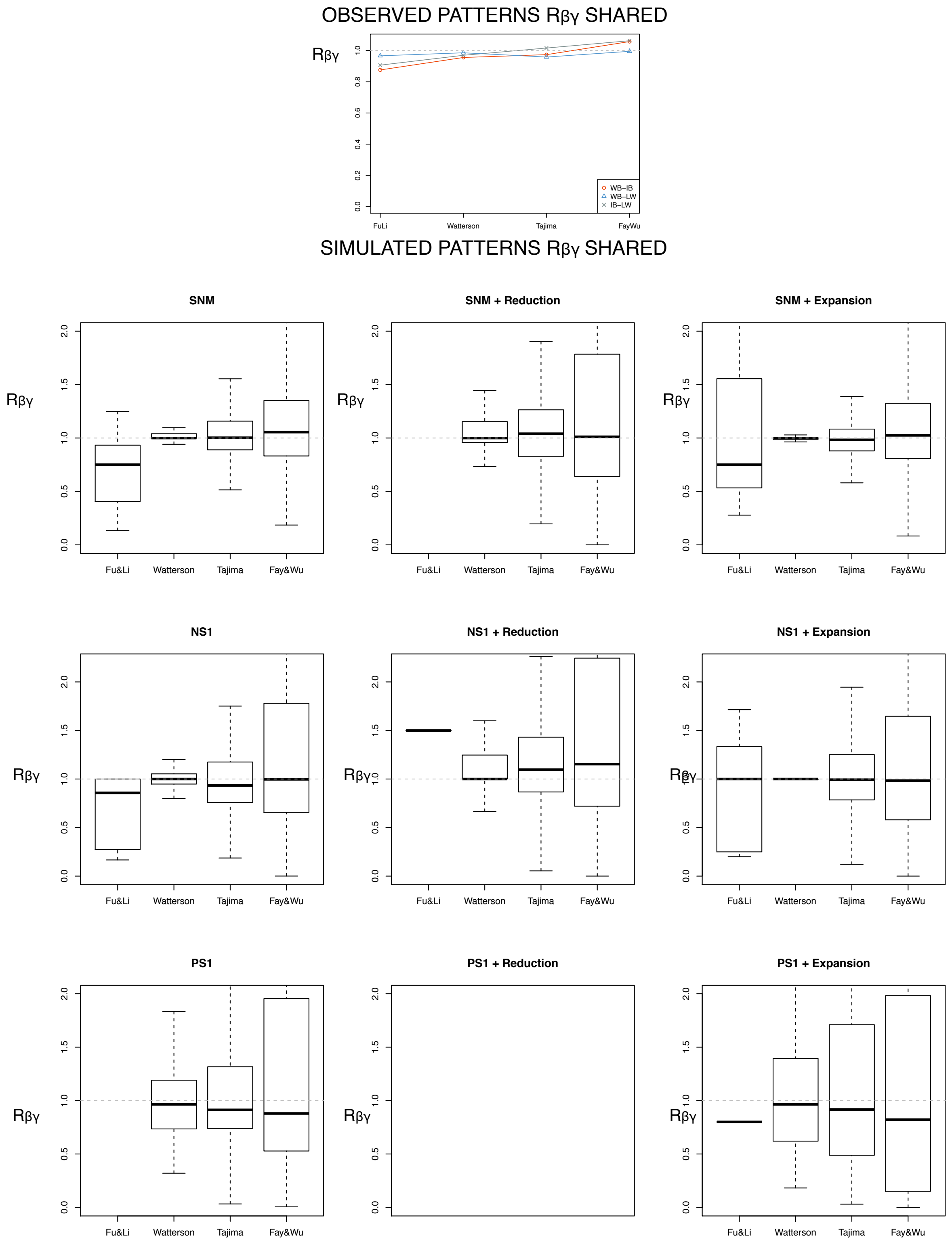

**Figure S37**. Estimates of $\alpha$ considering **Shared** SNPs for the Domestic population. In this second group of simulations a bottleneck was simulated after the split of Wild-Domestic populations (0.1x reduction of population size during 0.1 Ne generations, where Ne is the ancestral population size). Table S3B shows the parameter values used in these simulations. Here are shown the results of for scenarios with different combinations of positive and negative selection considering **no migration** from Wild to Domestic and **no change** in the population size after bottleneck.

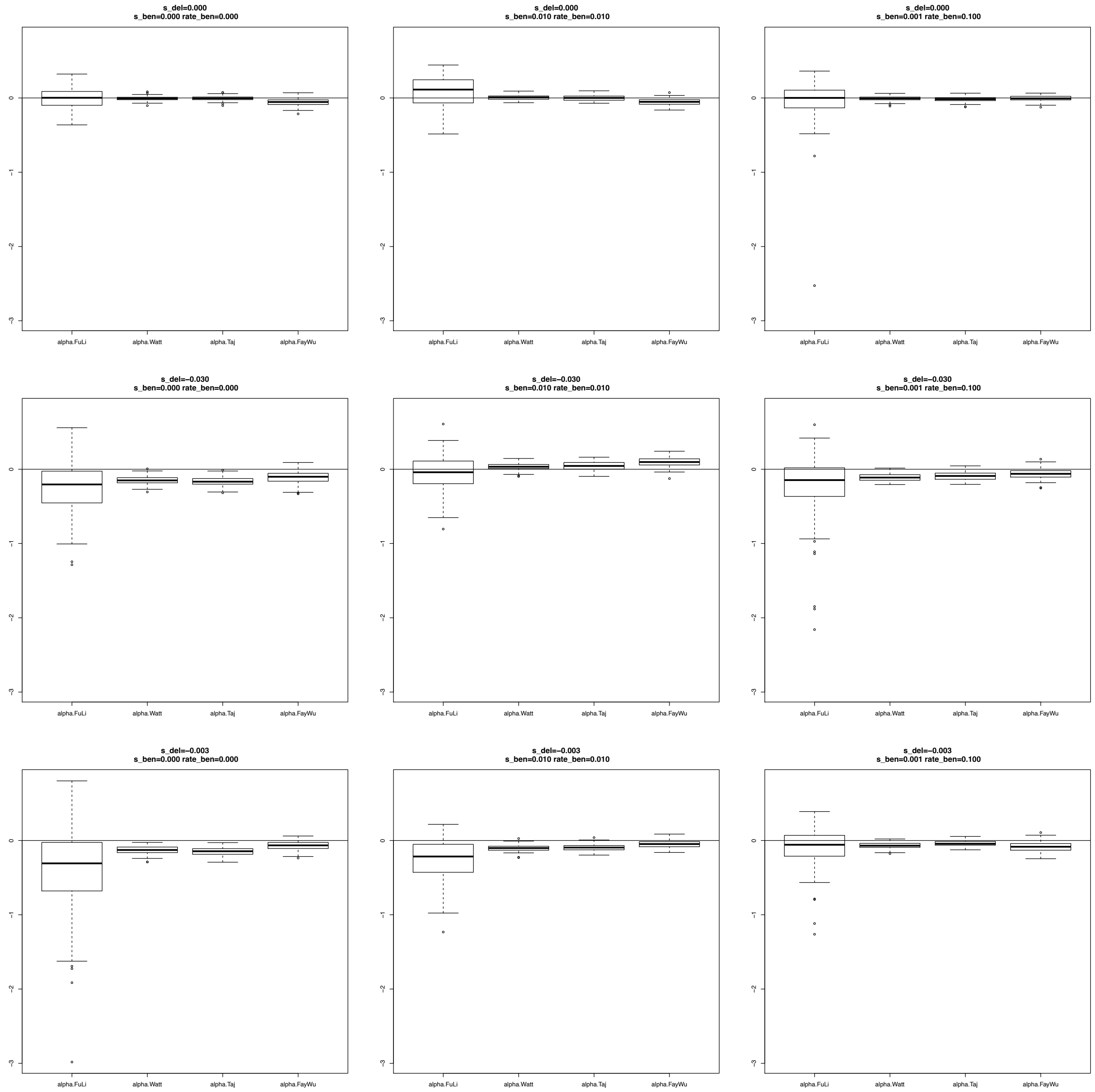

**Figure S38**. Estimates of $\alpha$ considering **Shared** SNPs for the Domestic population. A bottleneck was simulated after the split of Wild and Domestic populations (0.1x reduction of population size during 0.1 Ne generations, where Ne is the ancestral population size; see Table S3B for a description of the parameter values used in these simulations). The different scenarios were simulated using different combinations of positive and negative selection coefficients considering **5% migrants** from Wild to Domestic populations and **no change** in the population size after the bottleneck.

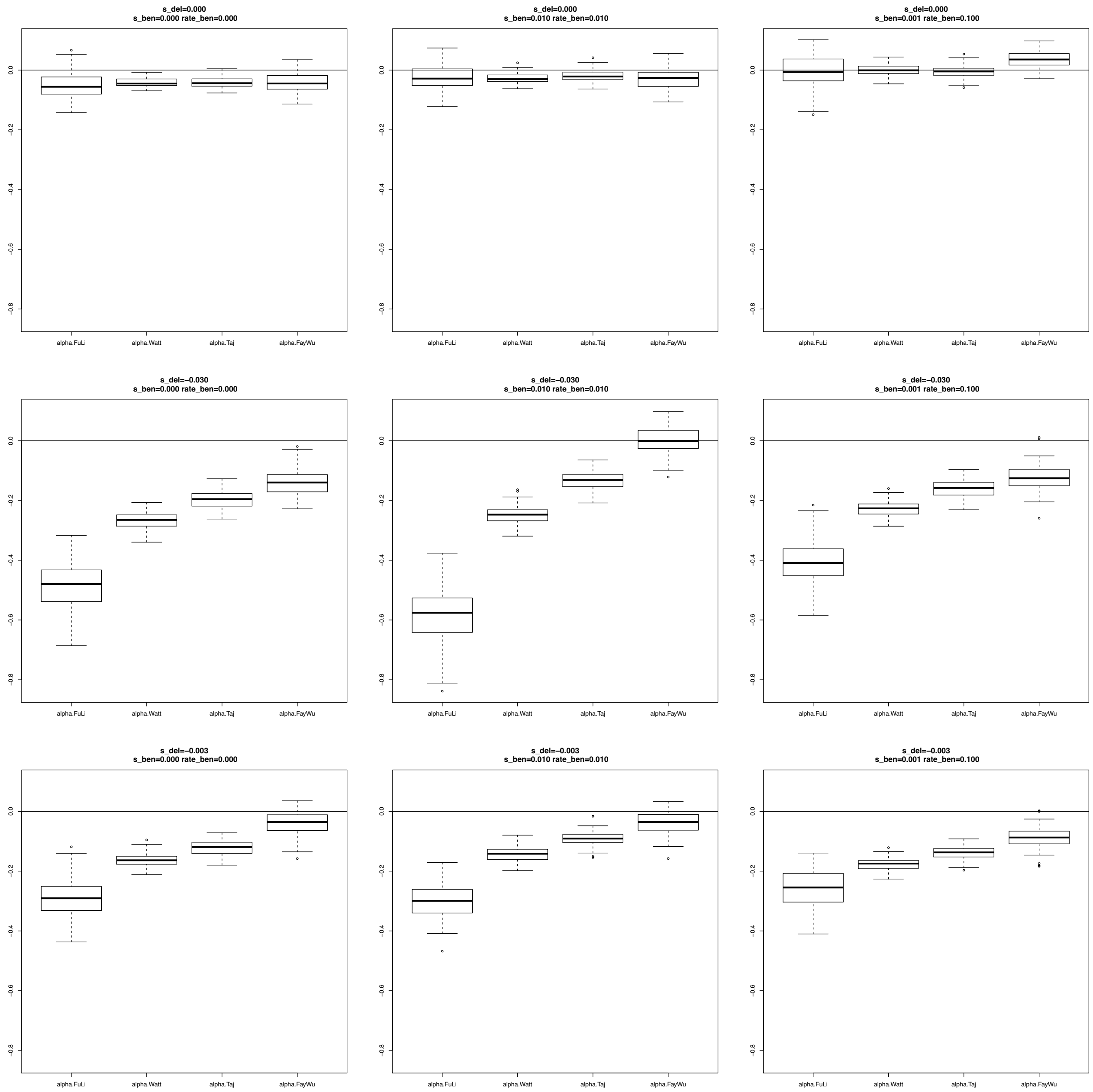

**Figure S39.** Estimates of $\alpha$ considering **Shared** SNPs for the Domestic population. A bottleneck was simulated after the split of Wild and Domestic populations (0.1x reduction of population size during 0.1 Ne generations, where Ne is the ancestral population size; see Table S3B for a description of the parameter values used in these simulations). The different scenarios were simulated using different combinations of positive and negative selection coefficients considering **no migration** from Wild to Domestic and ask increase (**Expansion)** of the population size after the bottleneck.

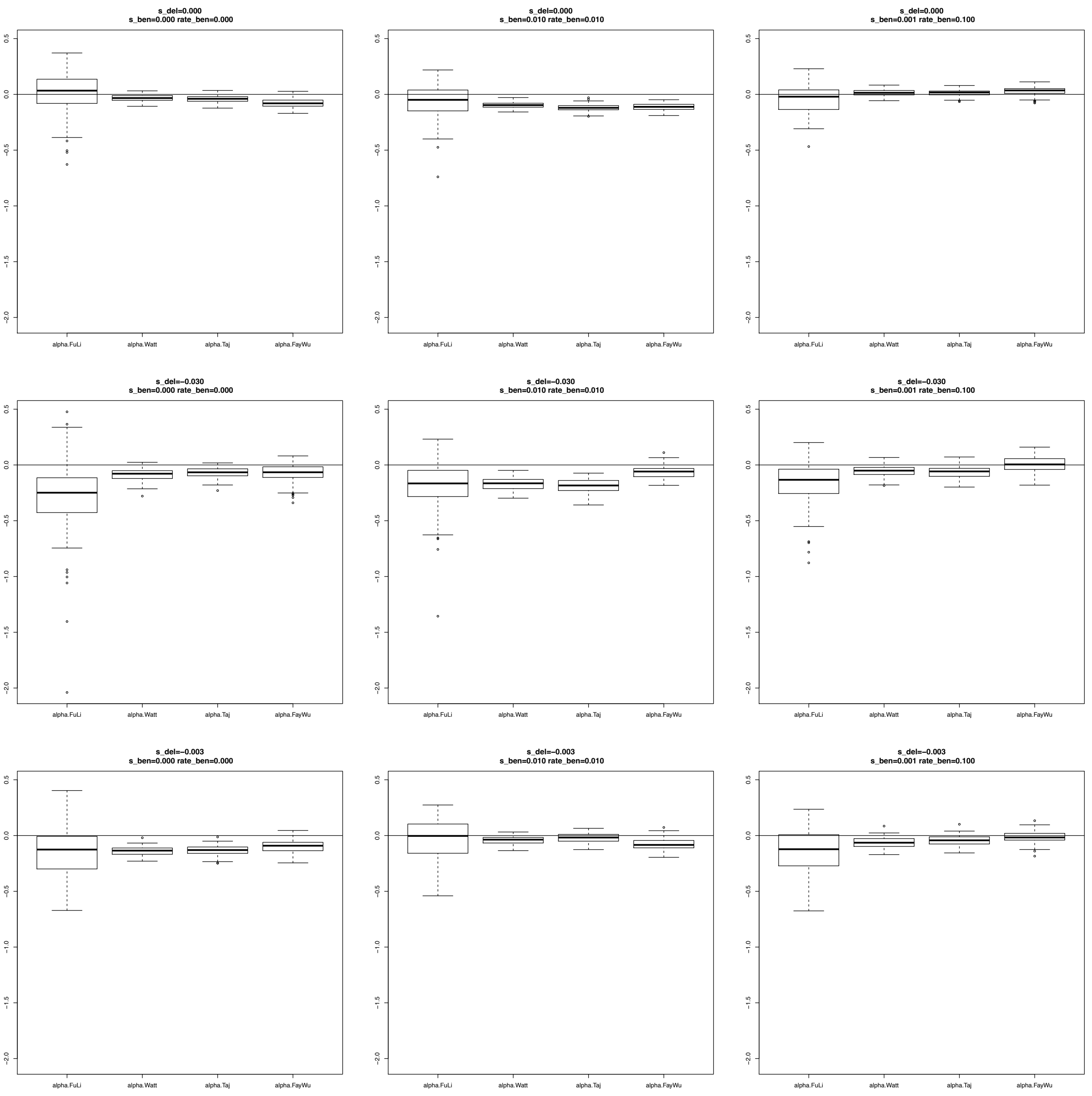

**Figure S40**. Estimates of $\alpha$ considering **Shared** SNPs for the Domestic population. A bottleneck was simulated after the split of Wild and Domestic populations (0.1x reduction of population size during 0.1 Ne generations, where Ne is the ancestral population size; see Table S3B for a description of the parameter values used in these simulations). The different scenarios were simulated using different combinations of positive and negative selection coefficients considering **5% migrants** from Wild to Domestic populations and an increase (**Expansion)** of the population size after the bottleneck.

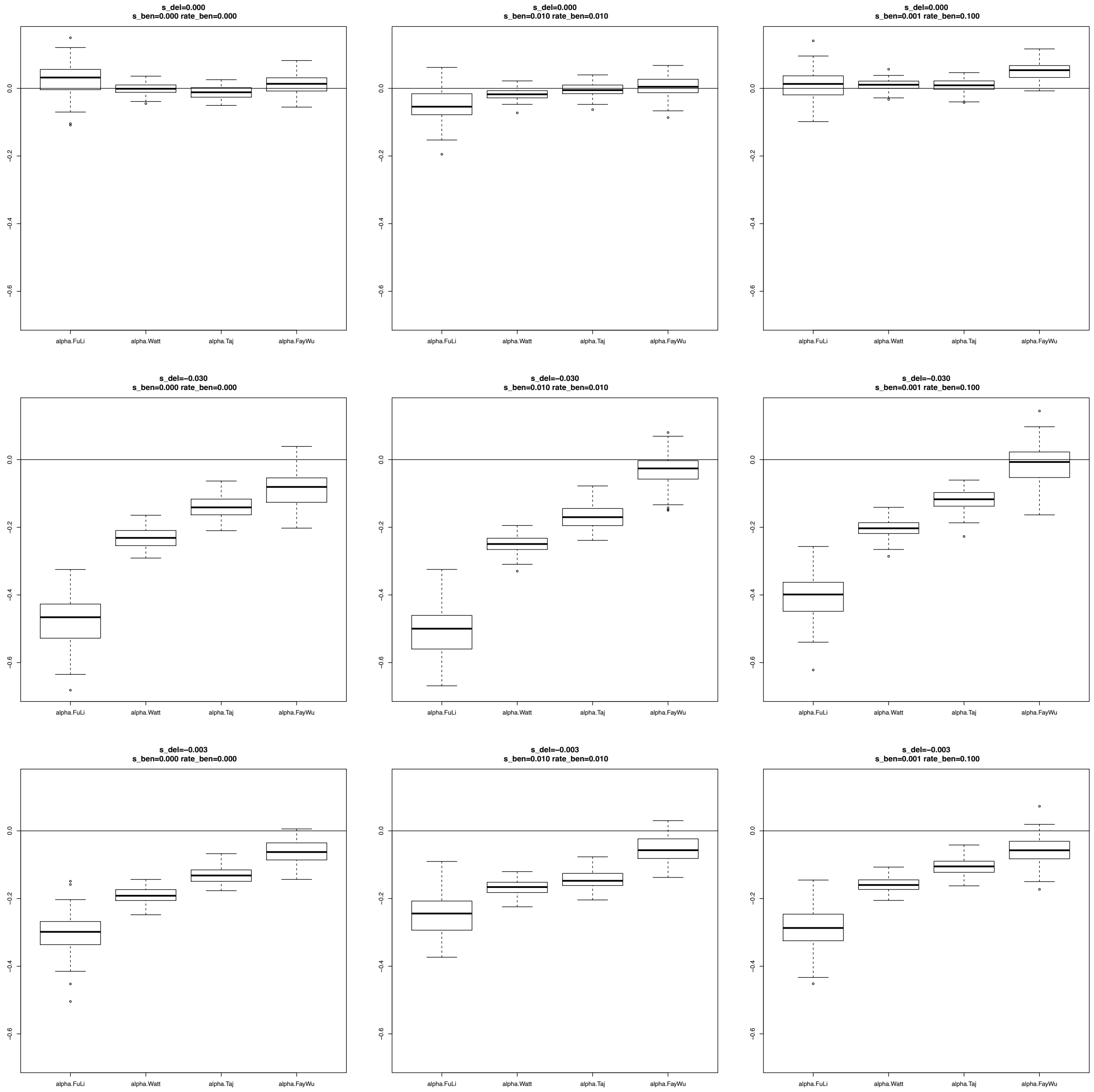

**Figure S41**. Estimates of $\alpha$ considering **Shared** SNPs for the Domestic population. A bottleneck was simulated after the split of Wild and Domestic populations (0.1x reduction of population size during 0.1 Ne generations, where Ne is the ancestral population size; see Table S3B for a description of the parameter values used in these simulations). The different scenarios were simulated using different combinations of positive and negative selection coefficients considering **no migration** from Wild to Domestic populations and a decrease (**Reduction)** of the population size after the bottleneck.

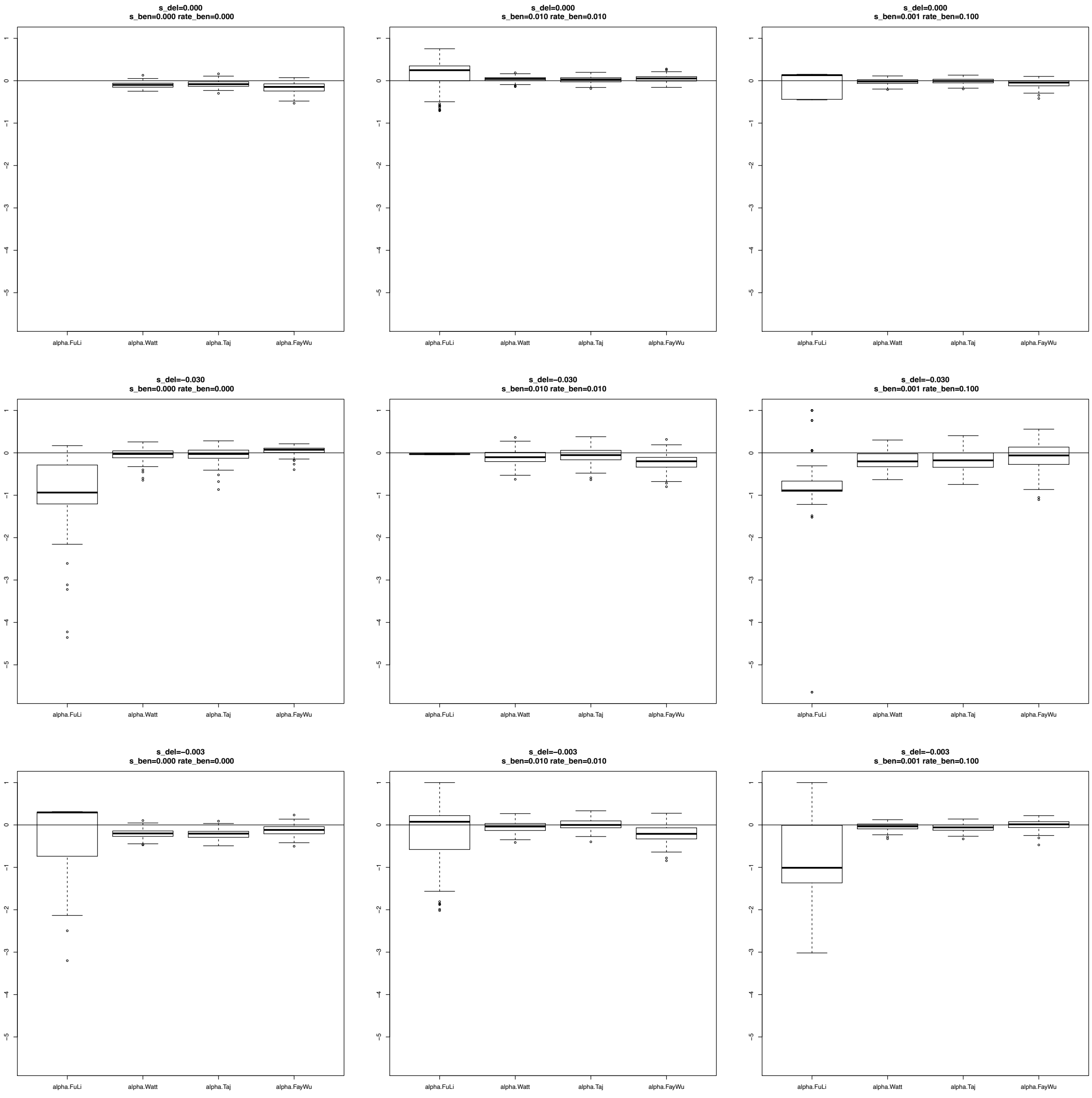

**Figure S42**. Estimates of $\alpha$ considering **Shared** SNPs for the Domestic population. A bottleneck was simulated after the split of Wild and Domestic populations (0.1x reduction of population size during 0.1 Ne generations, where Ne is the ancestral population size; see Table S3B for a description of the parameter values used in these simulations). The different scenarios were simulated using different combinations of positive and negative selection coefficients considering **5% migrants** from Wild to Domestic populations and a decrease (**Reduction)** of the population size after the bottleneck.

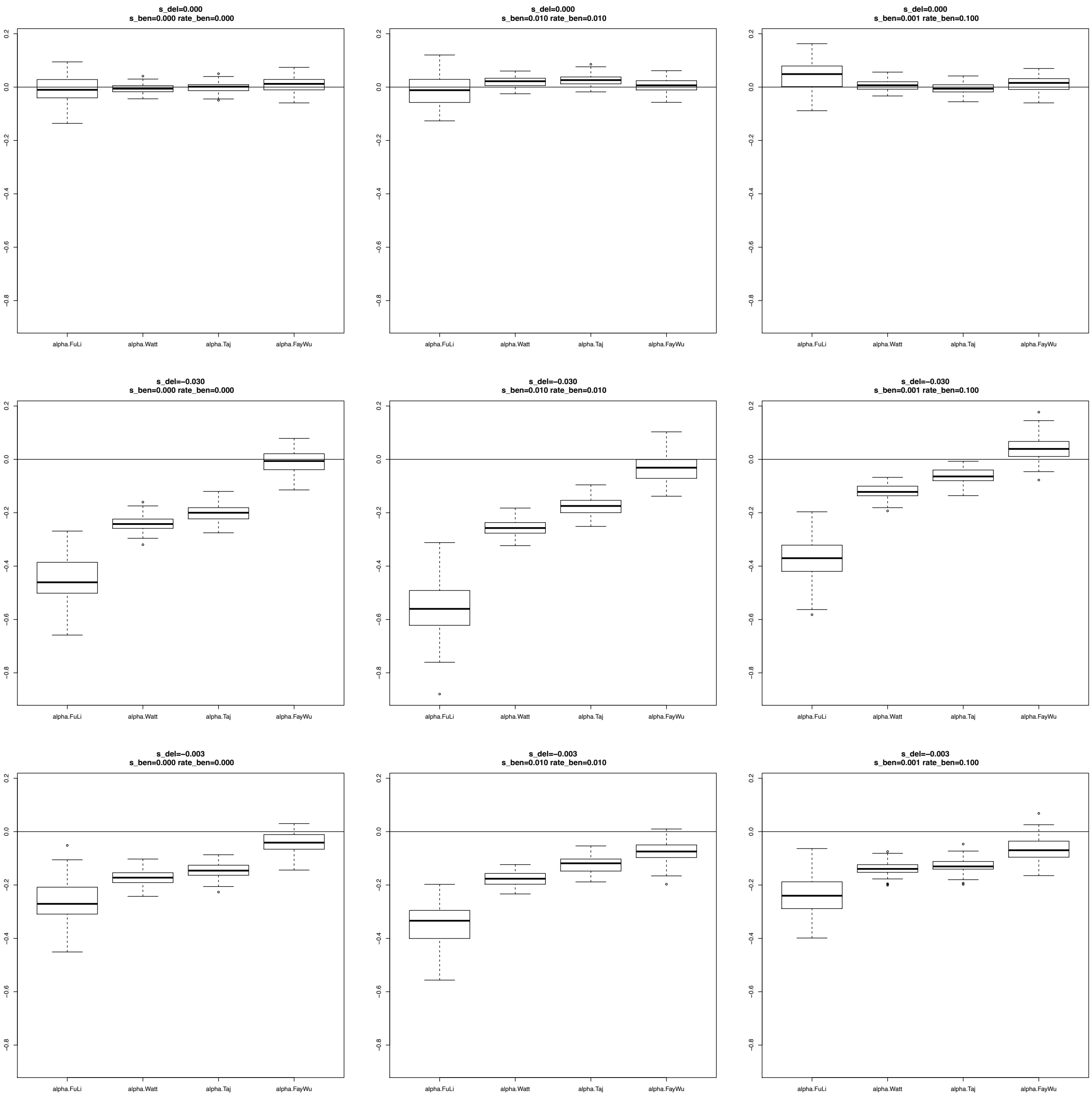

**Figure S43**. Estimates of $R\_\beta\gamma$ considering **Shared** SNPs for the Domestic population. A bottleneck was simulated after the split of Wild and Domestic populations (0.1x reduction of population size during 0.1 Ne generations, where Ne is the ancestral population size; see Table S3B for a description of the parameter values used in these simulations). The different scenarios were simulated using different combinations of positive and negative selection coefficients considering **no migration** from Wild to Domestic populations and **no change** in the population size after the bottleneck.

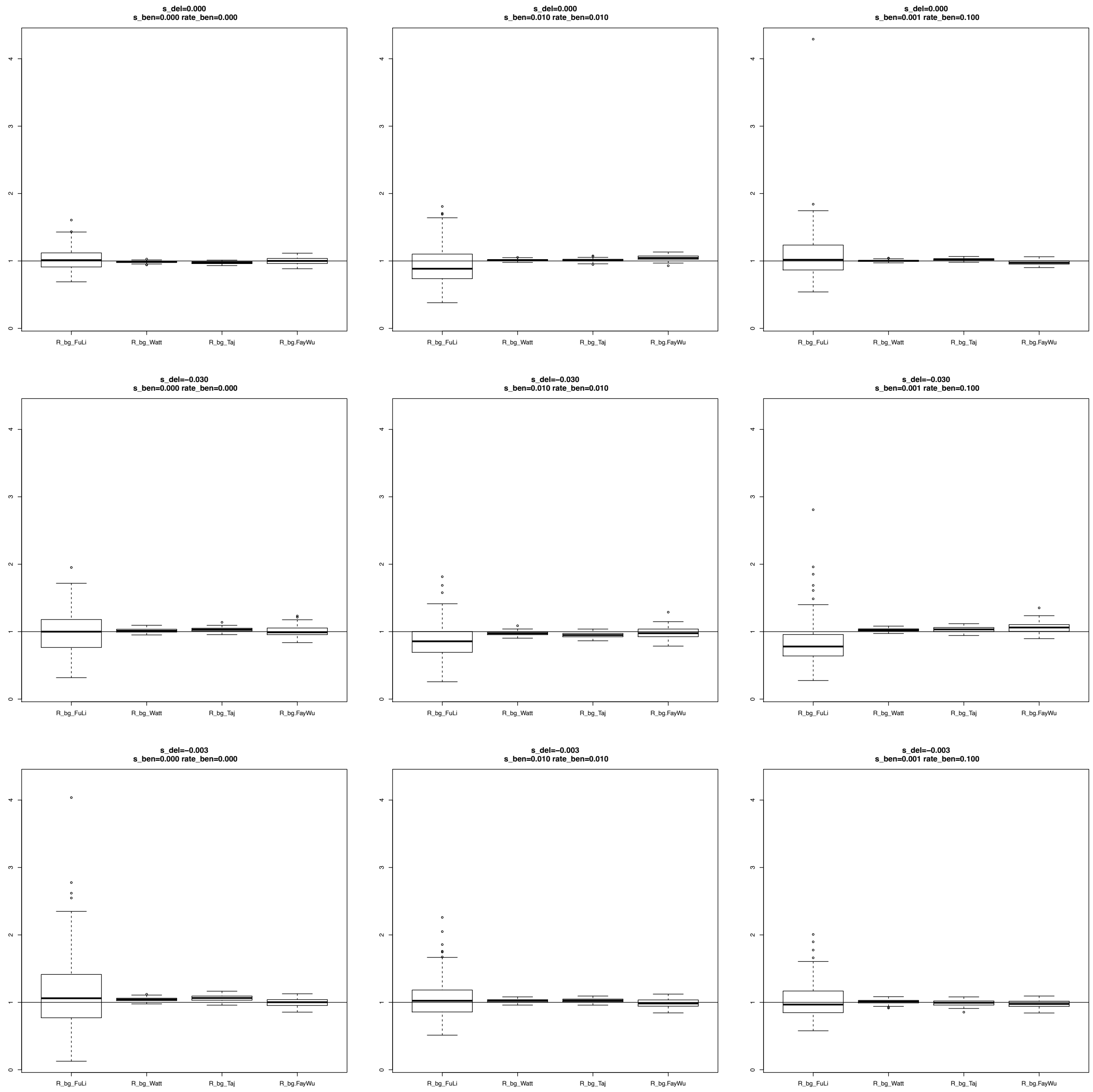

**Figure S44**. Estimates of $R\_\beta\gamma$ considering **Shared** SNPs for the Domestic population. A bottleneck was simulated after the split of Wild and Domestic populations (0.1x reduction of population size during 0.1 Ne generations, where Ne is the ancestral population size; see Table S3B for a description of the parameter values used in these simulations). The different scenarios were simulated using different combinations of positive and negative selection coefficients considering **5% migrants** from Wild to Domestic populations and **no change** in the population size after the bottleneck.

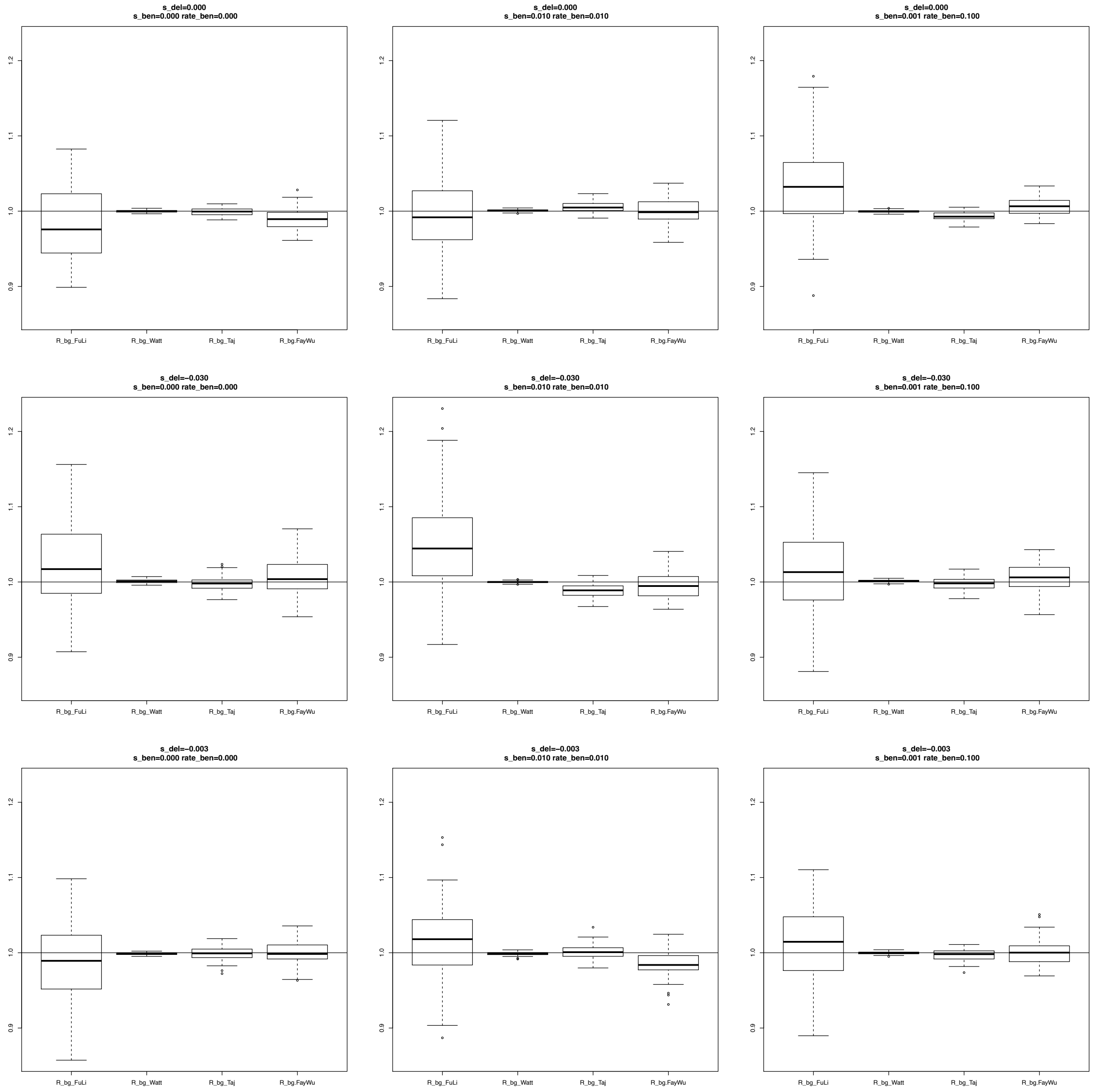

**Figure S45**. Estimates of $R\_\beta\gamma$ considering **Shared** SNPs for the Domestic population. A bottleneck was simulated after the split of Wild and Domestic populations (0.1x reduction of population size during 0.1 Ne generations, where Ne is the ancestral population size; see Table S3B for a description of the parameter values used in these simulations). The different scenarios were simulated using different combinations of positive and negative selection coefficients considering **no migration** from Wild to Domestic populations and an increase (**Expansion)** of the population size after the bottleneck.

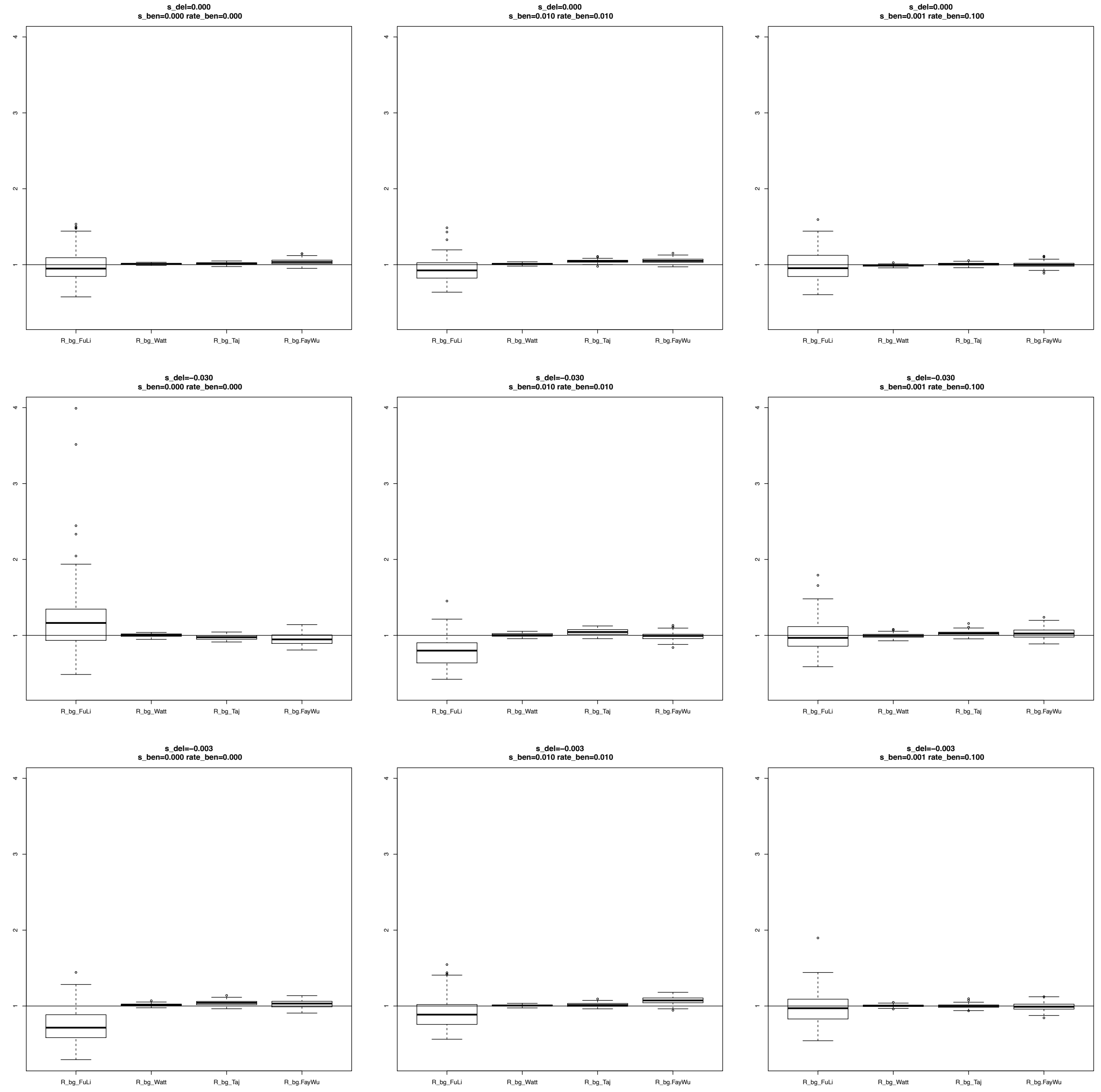

**Figure S46**. Estimates of $R\_\beta\gamma$ considering **Shared** SNPs for the Domestic population. A bottleneck was simulated after the split of Wild and Domestic populations (0.1x reduction of population size during 0.1 Ne generations, where Ne is the ancestral population size; see Table S3B for a description of the parameter values used in these simulations). The different scenarios were simulated using different combinations of positive and negative selection coefficients considering **5% migrants** from Wild to Domestic population and an increase (**Expansion)** of the population size after the bottleneck.

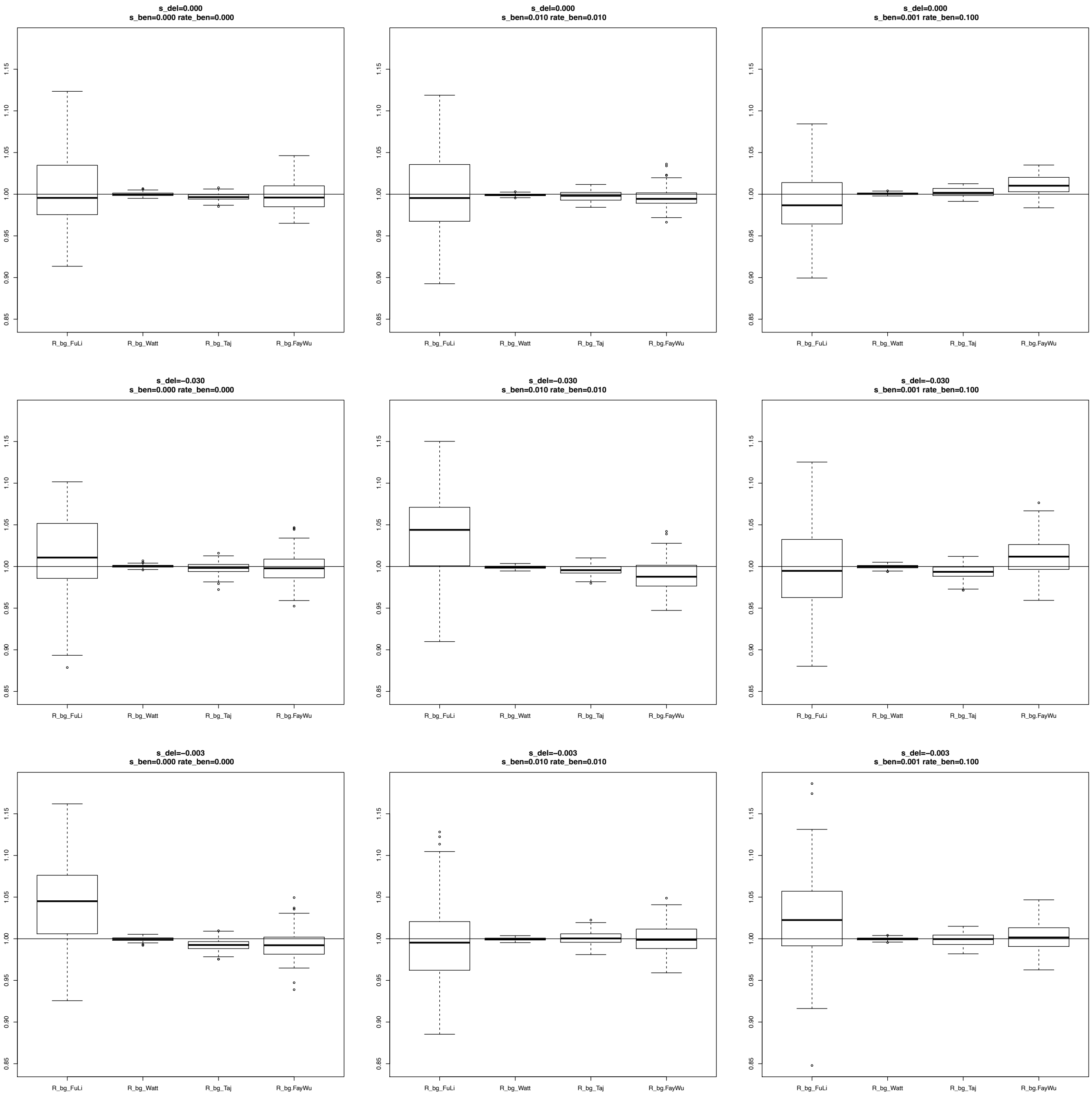

**Figure S47**. Estimates of $R\_\beta\gamma$ considering **Shared** SNPs for the Domestic population. A bottleneck was simulated after the split of Wild and Domestic populations (0.1x reduction of population size during 0.1 Ne generations, where Ne is the ancestral population size; see Table S3B for a description of the parameter values used in these simulations). The different scenarios were simulated using different combinations of positive and negative selection coefficients considering **no migration** from Wild to Domestic populations and a decrease (**Reduction)** of the population size after the bottleneck.

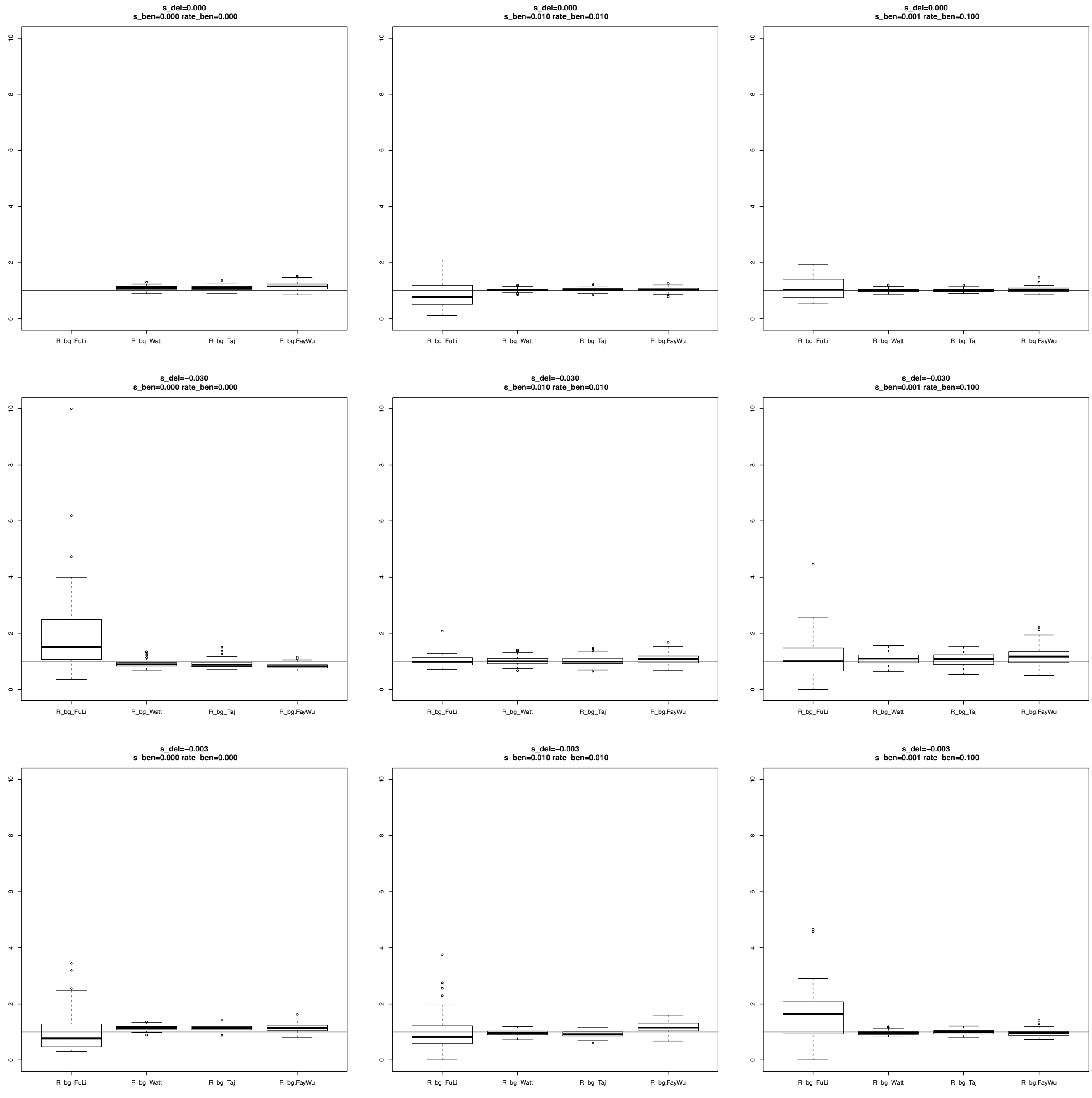

**Figure S48**. Estimates of $R\_\beta\gamma$ considering **Shared** SNPs for the Domestic population. A bottleneck was simulated after the split of Wild and Domestic populations (0.1x reduction of population size during 0.1 Ne generations, where Ne is the ancestral population size; see Table S3B for a description of the parameter values used in these simulations). The different scenarios were simulated using different combinations of positive and negative selection coefficients considering **5% migrants** from Wild to Domestic populations and a decrease (**Reduction)** of the population size after the bottleneck.

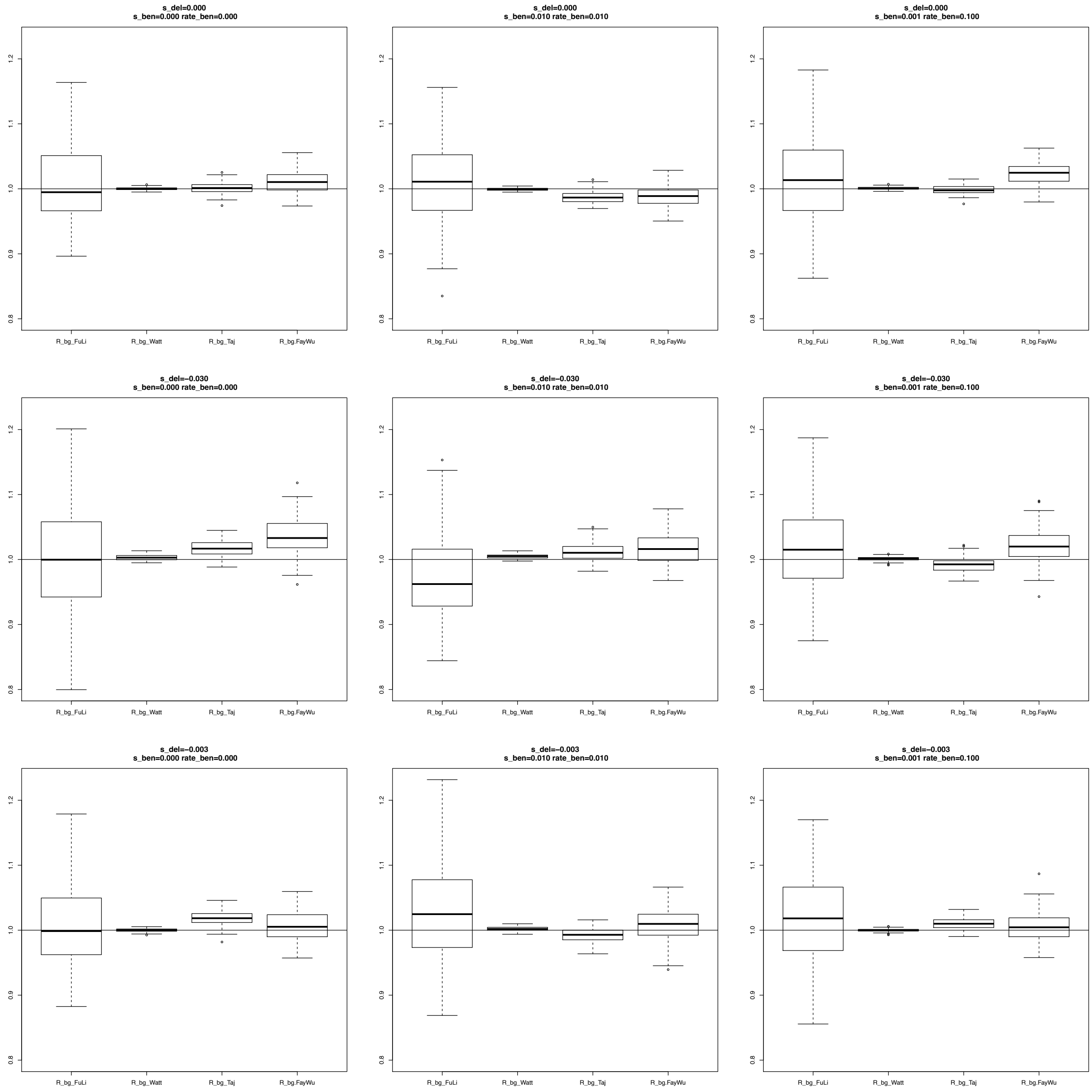
